# Co-opting the bacterial lipoprotein pathway for the biosynthesis of lipidated macrocyclic peptides

**DOI:** 10.1101/2025.10.31.685832

**Authors:** Jeff Y. Chen, Lingyang Zhu, Kevin Y. Zhang, Deborah A. Berthold, Wilfred A. van der Donk

## Abstract

Ribosomally synthesized and post-translationally modified peptides (RiPPs) are structurally diverse natural products that possess a range of bioactivities, often acting as antibiotics, antifungals, or metallophores. In RiPP biosynthesis, different modifying enzymes install an array of chemical motifs onto a precursor peptide. A recently described RiPP-modifying enzyme, ChrH, catalyzes a remarkably complex reaction on its precursor peptide that results in a macrocycle, heterocycle, and *S-*methyl group. By leveraging comparative genomics, we demonstrate that the products from a subfamily of enzymes related to ChrH display unexpected structural diversity, including the production of unmethylated macrocyclic congeners and C-terminally modified proteins over 30 kDa in size. Several of these precursors contain a signal peptide, sending them for downstream maturation by the bacterial lipoprotein biosynthetic pathway. Like bacterial lipoproteins, such peptides are modified by addition of a diacylglycerol (DAG) group to the N-terminal cysteine residue along with acylation of the N-terminal amine. Genome mining reveals that these RiPP-lipoprotein hybrids, which we term DAG-RiPPs, are widespread across bacterial phyla and are likely involved in different biological roles. Together, these results highlight a novel maturation paradigm for membrane-bound RiPPs and lay the foundation for the discovery and bioengineering of other RiPP-lipoprotein hybrids.

**Significance:** Ribosomally synthesized and post-translationally modified peptides (RiPPs) are a superfamily of natural products that display antibiotic, antifungal, anticancer, and metal-binding activities. Their biosynthesis typically follows a common logic in which modifying enzymes install chemical motifs onto a precursor peptide, followed by proteolytic processing and export from the cell. Herein, we describe the discovery and biochemical characterization of a new class of lipid-RiPP hybrid products. These RiPPs contain a signal peptide that exploits the endogenous bacterial lipoprotein biosynthesis pathway for lipidation, membrane localization, and potential secretion. Genome mining shows that these lipid-peptide hybrids are widespread across bacterial phyla.

## Introduction

Ribosomally synthesized and post-translationally modified peptides (RiPPs) are a class of structurally diverse natural products possessing a wide range of biological activities (1-5). A rapidly growing number of modifying enzymes have been shown to install various post-translational modifications onto their respective precursor peptides (6, 7). One such class of enzymes is the multinuclear non-heme iron-dependent oxidative enzyme family (MNIO, formerly DUF692/UPF0276), whose members catalyze diverse and challenging peptide modifications (8-10). MNIOs can act on different amino acid residues, such as cysteine, asparagine, aspartate, and phenylalanine. In the biosynthesis of the copper-binding peptide methanobactin, the MNIO MbnB (with its partner protein MbnC) converts a cysteine residue to an oxazolone-thioamide (11), and HvfBC has been reported to install the same modification on the conserved EGKCG motif of copper-binding oxazolins (12) (**SI Appendix, Fig. S1**). In alternative strategies, BufBC converts a cysteine residue to a 5-thiooxazole in the biosynthesis of the widespread bufferin peptides (13-15), and TglH (with its partner protein TglI) excises the β-carbon from a C-terminal cysteine to form norcysteine (16). ChrH performs another remarkable reaction in which a conserved motif featuring two cysteines is converted to a product that contains a macrocycle and a hydantoin heterocycle, and that is *S-*methylated (17) (**Fig. 1a**). In a diversion of reactions involving Cys residues, three other MNIOs, ApyH, PflD and MovX, act on aspartate and asparagine residues (18-20) (**SI Appendix, Fig. S1**), and PbsC was shown to perform *ortho*-hydroxylation of phenylalanine in the biosynthesis of the biphenomycin class of antibiotics (21). While MNIOs catalyze widely diverse reactions, they share a common TIM-barrel fold that houses two to three irons in its active site (22-24), and use molecular oxygen to perform a four-electron oxidation on their substrate in most cases (8).

**Fig. 1.**
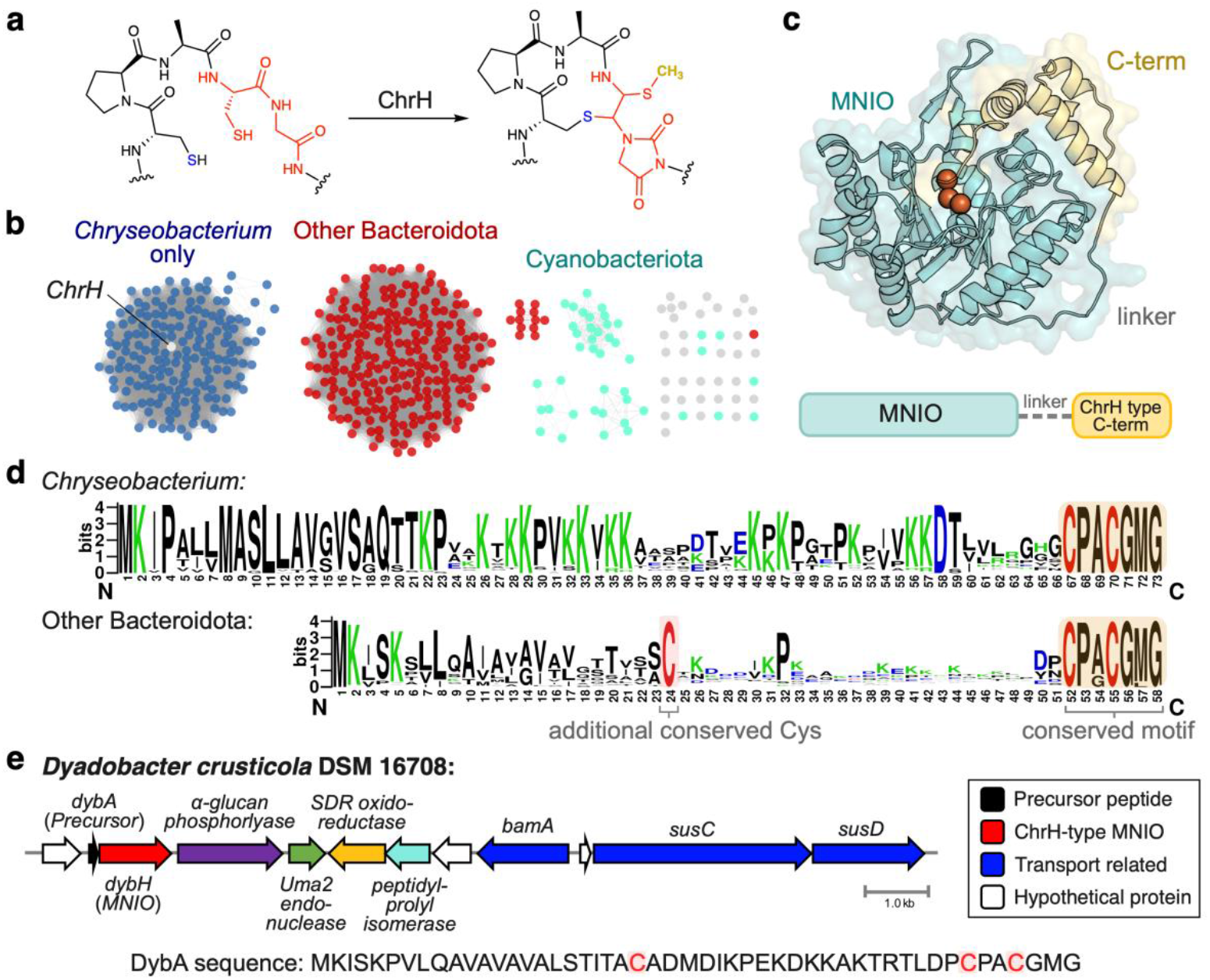
Bioinformatic analysis of the ChrH-like subfamily of MNIOs. **a** Reaction catalyzed by ChrH on its substrate peptide ChrA to install a macrocycle, hydantoin (imidazolidinedione) heterocycle, and an *S*-methyl group shown in bold gold font. Only the CPACG sequence from the ChrA peptide is shown. **b** SSN of protein sequences in the ChrH subfamily, showing division into two clusters, one enriched specifically in *Chryseobacterium* (blue), and one containing several Bacteroidota genera (red). Smaller clusters contain members from Cyanobacteria (cyan) and other phyla. The SSN was generated using an alignment score of 110 and e-value of 10^−5^. The full list of proteins can be found in **SI Appendix, Table S2** and Dataset 2. **c** Structural model of ChrH predicted by Alphafold 3 showing the overall fold of ChrH-family enzymes. Three iron atoms are modelled and represented as orange spheres. ChrH family enzymes contain the core MNIO domain in addition to an approximately 7.8 kDa C-terminal helical bundle joined by a linker. In Cyanobacteria, the C-terminal domain is encoded as a separate protein. **d** Sequence logo of ChrA-like peptides from *Chryseobacterium* and other Bacteroidota genera, showing shared conservation of the CPACGMG motif. However, ChrA-like peptides from *Chryseobacterium* are longer and lack an additional conserved cysteine. **e** Biosynthetic gene cluster from a representative species encoding the Bacteroidetes-type precursors, *Dyadobacter crusticola* DSM 16708. The gene for the precursor peptide DybA is located upstream of the gene encoding the MNIO DybH. Below: the amino acid sequence of DybA with conserved cysteines shown in red.

The biosynthetic gene cluster (BGC) encoding ChrH is enriched in the genus *Chryseobacterium*, but related members of this subfamily are also found in the Bacteroidota and Cyanobacteria phyla (8). Thus, we hypothesized that comparative genomics could facilitate the discovery of new products. We identified ChrH homologs across various gram-negative phyla including Bacteroidota, Cyanobacteria, and Myxococcota. Multiple sequence alignments show that the precursor peptides in the majority of other Bacteroidota genera contain an additional highly conserved cysteine residue. A representative member from this group was selected for analysis, DybA from *Dyadobacter crusticola*, which is modified by the MNIO DybH. Heterologous coexpression in *E. coli* and structural characterization showed that modified DybA is a macrocyclic congener of the ChrA product that instead contains a free, unmethylated thiol. Characterization of other homologs demonstrate a surprising substrate tolerance of the ChrH/DybH-subfamily, with homologs from Cyanobacteria modifying precursor proteins over 34 kDa in size. Additionally, bioinformatics and biochemical experiments show that several of these RiPPs contain an N-terminal signal sequence, resulting in lipidation of the modified peptide via the ubiquitous bacterial lipoprotein biosynthesis pathway. We demonstrate that the RiPP product of the lipoprotein pathway features the additional conserved Cys as the N-terminal residue, and that this Cys carries a diacylglycerol (DAG) group on its sulfur and an acyl moiety on its amino group. Bioinformatic analyses show that these RiPP-lipoprotein hybrids are widespread across other subfamilies of MNIOs. We suggest to call these products DAG-RiPPs to distinguish them from the growing group of lipidated RiPPs that are *N-*acylated (25).

## Results

### Phylogenetic analysis of ChrH-like gene clusters

To analyze the ChrH-like subfamily in detail, we first generated a sequence similarity network (SSN) (26) of ChrH homologs (**Fig. 1b**). We observed a clear division of the subfamily into two main clusters. One cluster is populated by members of the *Chryseobacterium* genus, which includes the previously characterized ChrH (17). The other cluster contains members of other Bacteroidota genera (mostly *Dyadobacter, Chitinophaga*, and *Hymenobacter*). Finally, smaller clusters are populated by other gram-negative bacterial phyla, including Cyanobacteriota, Myxococcota, and Planctomycetota.

Structural prediction of ChrH by Alphafold 3 (27) shows that in addition to the core TIM barrel that is present in all MNIOs, ChrH contains an additional C-terminal helical bundle (**Fig. 1c**). This additional C-terminal domain is present across all members of the ChrH family (in **Fig. 1b**), but is absent in other characterized MNIOs such as MbnB and TglH (23, 24). In the Cyanobacteriota MNIOs, this C-terminal domain is structurally conserved, but encoded as a separate polypeptide, suggesting fusion of the two domains at some point during evolution (**SI Appendix, Fig. S2**).

Analysis of the precursor peptides associated with these MNIOs showed that they contain the highly conserved CPxCGxG motif at the end of the peptide across the entire subfamily, with CPACGMG being the most frequently observed sequence (**Fig. 1d**). However, comparison of ChrA-like precursors from *Chryseobacterium* with precursors from other Bacteroidota clades show key differences. The latter are shorter and contain an additional, completely conserved cysteine residue in the middle of the peptide (**Fig. 1d**). Given the proclivity of MNIOs to act on Cys residues (8), we hypothesized that the additional cysteine could be modified or affect the reactivity of the MNIO.

Co-occurrence analyses also support differences between the BGCs of *Chryseobacterium* versus other Bacteroidota genera (**SI Appendix, Table S1**). Such analysis shows that whereas the ChrH-type enzymes are part of an operon with highly conserved genes, the Bacteroidota enzyme families appear to be encoded in BGCs that often consist of the precursor peptide and MNIO, with other, mostly transport related genes at lower conservation. Notably, these BGCs do not encode an obvious conserved partner protein used for substrate binding that is encoded nearby for most characterized MNIOs (8).

To characterize the modification performed by the divergent cluster in the ChrH subfamily, we selected a representative member, DybH, from *Dyadobacter crusticola* DSM 16708, isolated from soil crusts in Colorado (28) (**Fig. 1e**). DybH has an amino acid sequence identity of 40% compared to ChrH, and the associated precursor peptide DybA contains the third conserved cysteine in addition to the CPxCGxG motif. The *dyb* BGC also lacks a gene for an obvious partner protein.

### Coexpression of DybA and DybH yields a − 4 Da modification

We first heterologously co-expressed the genes encoding the His-tagged precursor peptide DybA and DybH in *E. coli*. Purification of the peptide (which we will refer to as DybAH heretoforth) and analysis by matrix-assisted laser desorption/ionization time-of-flight mass spectrometry (MALDI-TOF MS) showed a − 4 Da mass shift relative to the unmodified precursor peptide (**Fig. 2a**). As forecast by the architecture of the BGC, DybH was active without the need for a partner protein. The − 4 Da shift resulting from co-expression with DybH is different from that observed for ChrH, which results in an overall +10 Da mass shift (17).

**Fig. 2.**
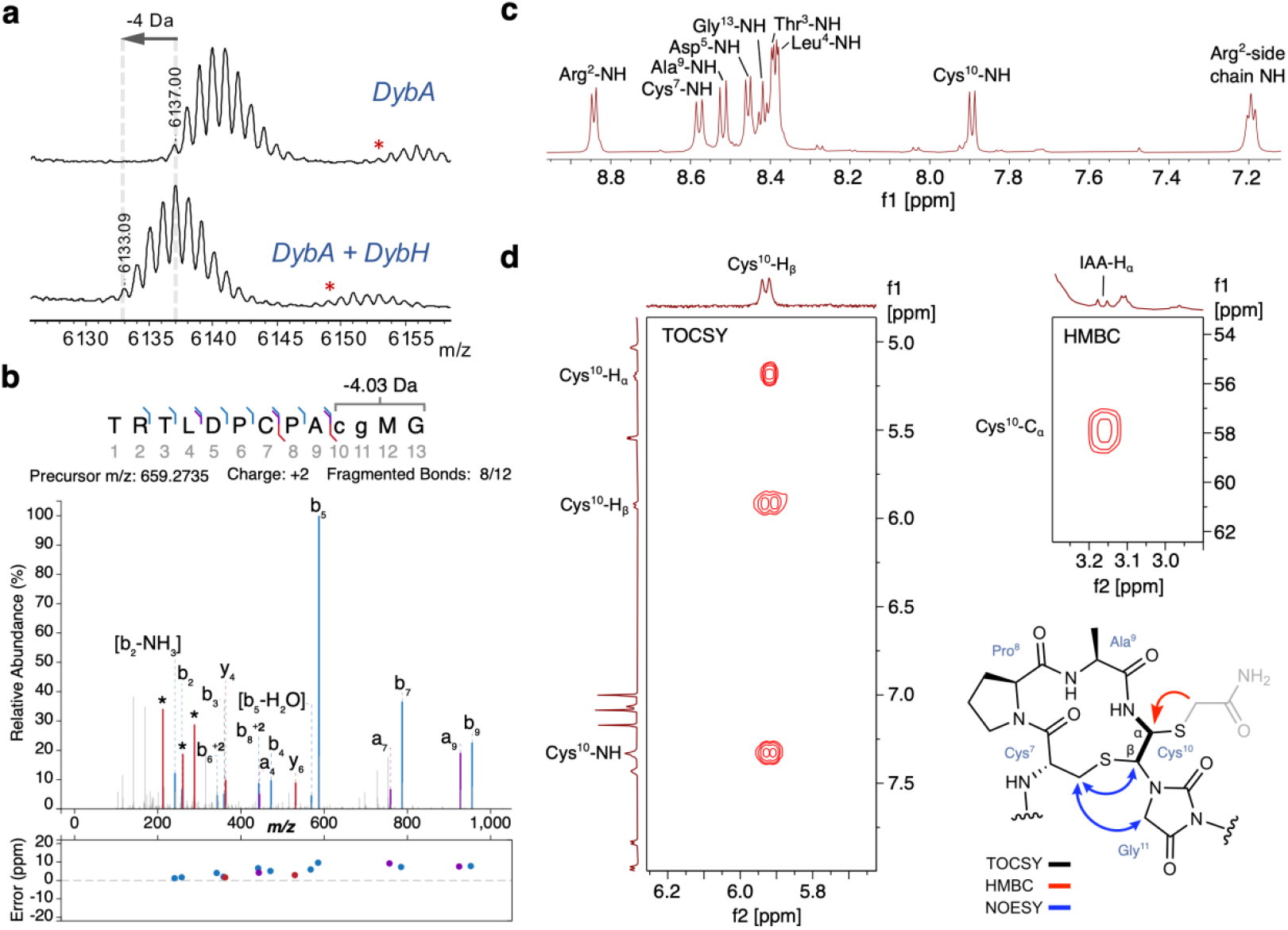
DybH installs a macrocycle and heterocycle onto DybA. **a** MALDI-TOF MS of DybA expressed alone or coexpressed with DybH in *E. coli*. M+H peaks are labelled. Expected monoisotopic [M+H]^+^ mass of unmodified peptide: 6137.12 Da (**SI Appendix, Table S3**). Red asterisks denote peptide with oxidized methionine (+16 Da). **b** LC-MS/MS spectrum of LysC-digested, DybH-modified peptide annotated by IPSA. The input sequence entered the − 4 Da modification to the positions indicated with lower-case letters. Asterisks denote modified y-ion fragmentations (see **SI Appendix, Fig. S3**). **c** ^1^H NMR spectrum of the peptide in panel b showing eight amide N-H protons (and an N-H from an Arg side chain), indicating two missing signals. **d** Left: diagnostic TOCSY signals for the DybAH product showing a new modified spin system with three protons. Right: key HMBC correlation between the methylene proton of the carbamidomethylene group (in grey) and the α-carbon of the thio-hemiaminal. The proposed structure of DybAH is shown (IAA-alkylated) based on diagnostic TOCSY, HMBC, and NOESY signals. Residue numbers are based on the fragment obtained after LysC cleavage. Full spectra and assignments in **SI Appendix, Figs S6-15** and **Tables S4-5**.

Digestion of the full-length peptide to a 13-mer peptide by LysC endoproteinase was followed by LC-MS/MS analysis. The fragmentation data was analyzed using the Interactive Peptide Spectral Annotator (IPSA) (29), which generates predicted peptide fragments based on user-entered modifications. The analysis localized the − 4.0313 Da modification to the C-terminal CPACGMG motif (**Fig. 2b**). This mass change is consistent with a loss of 4 hydrogens. Several observed fragments could be assigned to modified *y*-ions resulting from the final four amino acids (**SI Appendix, Fig. S3**). These fragments suggest that whereas the product is distinct from the product formed by ChrH (**Fig. 1a**), similarities in structure do exist in the DybH product. Reaction with the thiol-specific alkylating reagent iodoacetamide (IAA) resulted in one adduct to the 13-mer peptide, indicating the presence of one free thiol, with the second thiol that was present in the substrate now modified such that it is not reactive towards electrophiles (**SI Appendix, Fig. S4**).

UV-vis characterization of the modified peptide did not reveal an absorbance feature, suggesting that aromatic functional groups observed in other MNIO-modified peptides such as methanobactins or bufferins were not formed (11, 13) (**SI Appendix, Fig. S5**). Together, the presence of a free thiol, a − 4 Da modification, and the observed fragments by tandem MS suggest that the product could be the non-methylated congener of modified ChrA.

### DybAH analysis by NMR spectroscopy shows macrocycle and heterocycle formation

We used NMR spectroscopy to determine the structure of the modified, LysC-digested DybAH. The ^1^H NMR spectrum in 90% H_2_O:10% D_2_O showed the disappearance of the two amide N-H signals for methionine and glycine in the conserved CPACGMG motif (**Fig. 2c, SI Appendix, Fig. S6**). Furthermore, TOCSY and HSQC data showed a coupled spin system with three protons associated with the former Cys10 (numbering based on the peptide obtained by LysC digestion, **Fig. 2d, SI Appendix, Fig. S7 and 8**). The former Cys10 β-proton at 5.81 ppm is shifted far downfield from a typical Cys β-proton. We observed an NOE from this β-proton of former Cys10 to the β-protons of Cys7 (**SI Appendix, Fig. S9**). An NOE cross peak was also observed between the β-protons of Cys7 and the α-protons of Gly11. Collectively, these observations are similar to previous observations for the ChrA product and suggest formation of two thiohemiaminals (17) (**Fig. 2d**). In H_2_O/D_2_O, the observed peaks for both thiohemiaminal protons (Cys^10^ H_α_ and H_β_) were broad, which was also observed for modified ChrA. Therefore, we alkylated the peptide with IAA and recorded a spectrum in DMSO-d6. The broad peak corresponding to the former Cys^10^ H_α_ sharpened to a triplet, as expected by being split by the former Cys^10^ NH and H_β_ (**SI Appendix, Fig. S10-13**). These spectra confirmed that the β-proton of former Cys10 is a CH group in the product instead of a CH_2_ group in the substrate (**SI Appendix, Fig. S11**). Analysis of the IAA-alkylated peptide also revealed the position of the free thiol in the enzymatic product because we observed a cross peak in the HMBC spectrum between the α-protons of the carbamidomethylene originating from IAA (3.16 ppm) and the α-carbon (57.3 ppm) of the former Cys10 (**Fig. 2d, and SI Appendix, Fig. S14**). These data together support the structure of modified DybA as having a macrocycle and hydantoin heterocycle (**Fig. 2d, SI Appendix, Fig. S15**). While the stereochemistry of the thiohemiaminals is predicted to be *trans* based on the J-coupling of 11.8 Hz, additional crystallographic studies of the modified peptide are needed to unambiguously establish the absolute stereochemistry.

### In vitro reconstitution of DybH activity

We next purified His-tagged DybH aerobically. DybH was yellow in color, unlike ChrH, which is purple when purified aerobically (17). DybH co-purified with approximately 1.8-2.2 irons per protomer under the expression conditions used in this study, as determined by the ferene S colorimetric assay (30). The UV-Vis spectrum of DybH showed a shoulder at 350 nm, which is also present in other MNIOs and non-heme iron-containing enzymes (12) (**SI Appendix, Fig. S16**).

Incubation of DybA with aerobically prepared DybH resulted in production of the − 4 Da product and DybH performed multiple turnovers of substrate (**SI Appendix, Fig. S17**). These results confirm that a partner protein is not needed (as also observed with MovX (19)), and that DybH alone can perform the four-electron oxidation of its substrate. These data also suggest a mechanism consistent with that proposed for ChrH (17). The isolation of a congener lacking *S-* methylation supports a stepwise mechanism in which heterocyclization and macrocyclization are not concerted with the final *S*-methylation step (**SI Appendix, Fig. S18**).

### Modifications performed by homologs in Bacteroidota and Cyanobacteria

The different reactivity at the same CPACGMG motif observed in DybA versus ChrA prompted us to explore homologs in other clades with different gene architectures to investigate whether they produce the *S-*methylated or unmethylated product. An unrooted maximum-likelihood phylogenetic tree of the ChrH subfamily was generated (**Fig. 3a**). The tree shows separation of distinct clades, predominantly expanded in the Bacteroidota genera *Chryseobacterium, Chitinophaga, Dyadobacter*, and *Hymenobacter*, as well as the phyla Cyanobacteriota/Myxococcota. The associated BGCs not only encode ChrA-type and DybA-type precursor peptides, but also precursors that contain variations to the CPACGMG motif. For example, some precursors from Myxobacteria have a CPACGMMM motif that is not at the C-terminus, but instead embedded within the peptide (**Fig. 3a**). Furthermore, we observed that several Cyanobacteria BGCs encode putative precursor proteins over 30 kDa in size, which end with the CPACGMG motif at the C-terminus. These unusually large RiPP precursors are tetracopeptide repeat containing proteins (31), but otherwise do not contain any other obvious predicted domains by sequence or structure.

Given the absence of a ChrI homolog in the *dyb* BGC and the observed activity of DybH without such a partner protein, we revisited the requirement of ChrI for ChrH activity, which we had previously reported (17). Coexpression of ChrA with ChrH in *E. coli*, using the same expression system as used above for DybAH, resulted in formation of the +10 Da product even in the absence of ChrI (**Fig. 3b**). This discrepancy compared to our previous findings may be the result of a higher expressing plasmid construct used in this study. In contrast to the +10 Da product with ChrAH, coexpressions of ChsA with ChsH from *Chitinophaga sancti*, and HymA with HymH from *Hymenobacter gelipurpurascens* all showed production of the − 4 Da (unmethylated) product (**Fig. 3b**; for BGCs, see **SI Appendix, Fig. S19**). The C-terminal sequence of HymA ends with CP<u>G</u>CG<u>L</u>G, resulting in formation of a different macrocyclic congener. Coexpression of ArlAH, which contains an embedded CPACGMMM motif, showed a mixture of both the unmethylated and *S-*methylated product, further supporting that these two products are formed via the same route (**Fig. 3b**). The reasons for the methylations observed with ChrAH and ArlAH is currently not understood. We cannot rule out that a protein in *E. coli* may have specificity for methylating the ChrAH/ArlAH product and not the products from the other systems.

**Fig. 3.**
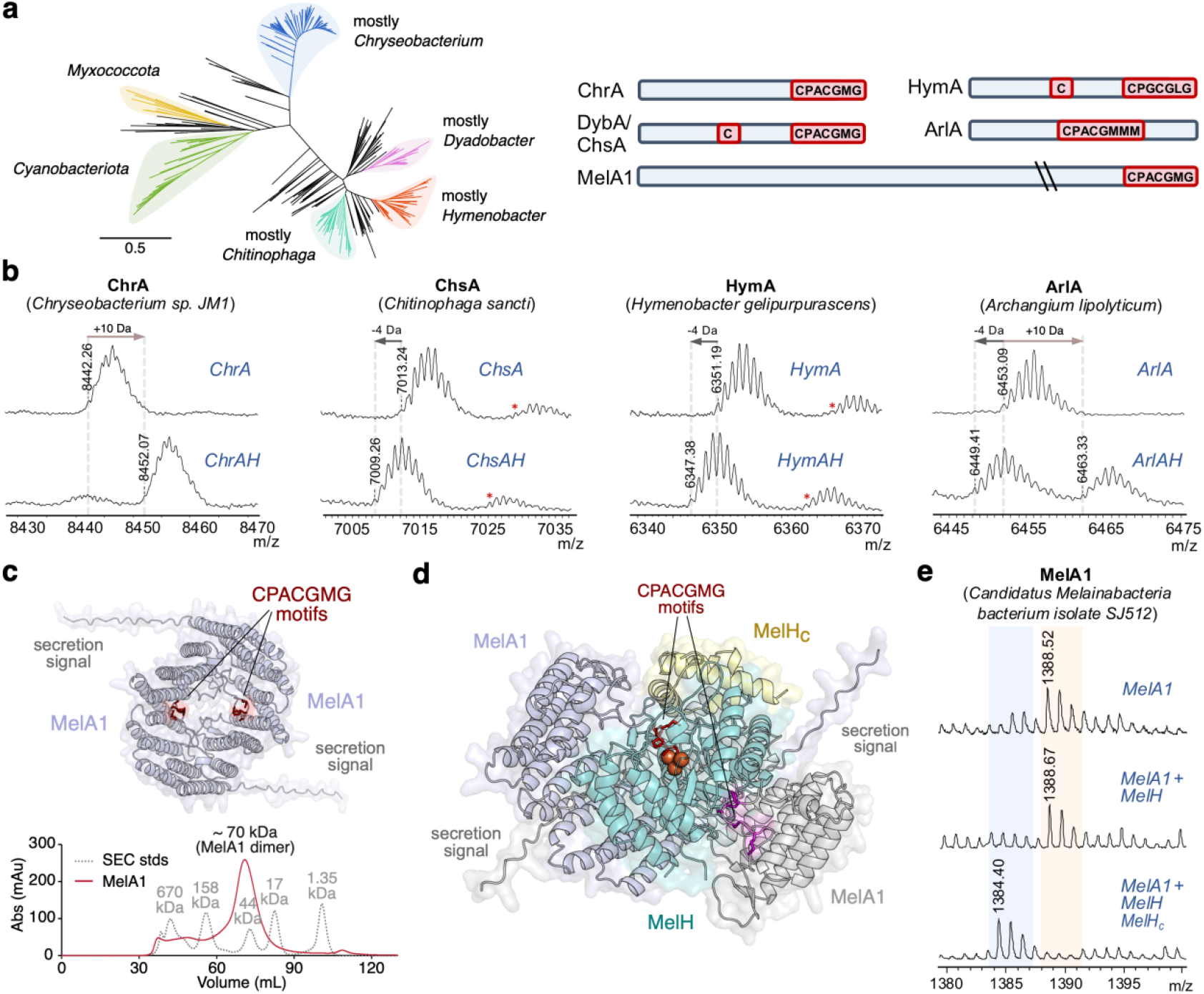
Characterization of ChrH homologs. **a** Left: Unrooted maximum-likelihood phylogenetic tree of the ChrH subfamily. Scale bar for branch length indicates number of amino acid substitutions per site. Right: Diagrams of representative substrates (not to scale), showing conserved cysteines and the conserved CPxCGxG motif in different contexts. **b** MALDI-TOF MS of peptides coexpressed with their respective MNIO from representative members from Bacteroidota clades (*Chryseobacterium, Chitinophaga, Hymenobacter*) and a member from Myxococcota (*Archangium*). Expected monoisotopic [M+H]^+^ m/z: ChrA, 8442.54 Da unmodified, 8452.52 Da modified (methylated); ChsA, 7013.45 Da unmodified, 7009.42 Da modified; HymA, 6351.22 Da unmodified, 6347.18 Da modified; ArlA, 6453.07 Da unmodified, 6449.04 Da modified. Red asterisks denote peptide with oxidized methionine (+16 Da). **c** Alphafold 3 model of a homodimer of MelA1 from Candidatus *Melainabacteria bacterium* isolate SJ512, showing the C-terminal CPACGMG motif in red. Below: size-exclusion chromatography (SEC) analysis of MelA1 indicates dimer formation. **d** Alphafold 3 model of (MelA1)_2_ in complex with MelH and MelH_c_. The CPACGMG motif (shown as red sticks) from one copy of MelA1 is positioned next to the triiron active site (shown as orange spheres). The second copy of MelA1 is shown in grey. **e** MALDI-TOF MS of the C-terminal fragment of MelA1 containing the CPACGMG motif (obtained by LysC-digestion) from experiments in which MelA1 was expressed alone, with MelH, or together with MelH and MelH_c_. Expected monoisotopic [M+H]^+^ mass of C-terminal fragment: 1388.62 Da unmodified, 1384.58 Da modified.

We next investigated the homologous gene clusters from Cyanobacteria that contain precursor proteins 30 to 40 kDa in size. One example is the *mel* cluster from *Melainabacterium bacterium* isolate SJ512 (**SI Appendix, Fig. S19**). The 34 kDa precursor protein MelA1 contains an N-terminal secretion signal peptide. The protein is predicted by Alphafold 3 to form a dimer, which was validated by size-exclusion chromatography (**Fig. 3c**). In this gene cluster, the ChrH-like C-terminal domain is separately encoded as a small protein that we call MelH_c_. Coexpression of His_6_-tagged MNIO (MelH) pulled down untagged MelH_c_ during purification, confirming that MelH_c_ forms a stable heterodimeric complex with MelH (**SI Appendix, Fig. S20**). We structurally modelled this complex together with the substrate MelA1 with Alphafold 3 (**Fig. 3d**). Complexes of MelH/H_c_ with the MelA1 dimer or its monomer produced similar predicted local distance difference test (pLDDT) values (**Fig. 3d** and **SI Appendix, Fig. S21**). Interestingly, both models predict that the CPACGMG motif from MelA1 is positioned into the active site of MelH in the complexes.

Coexpression in *E. coli* of the precursor protein MelA1 with the MNIO MelH did not yield any modification (**Fig. 3e**). However, coexpression of MelA1 with MelH and MelH_c_ led to production of the unmethylated (− 4 Da) product at the CPACGMG motif, further demonstrating the diversity of substrates processed by this MNIO subfamily. In turn, it suggests that the product structure can fulfill its currently unknown physiological function both at the end of a > 30 kDa secreted protein or as a small peptide.

### The product of the dyb BGC is a RiPP-lipoprotein hybrid

While we initially hypothesized that the internal cysteine in DybA and DybA-like precursor peptides could be a site for modification by the MNIO, we only observed modification at the CPACGMG motif. By analyzing the N-terminal sequence in more detail, we noticed a stretch of hydrophobic amino acids that precede the conserved cysteine (**Fig. 1d**). The signal peptide prediction program SignalP 6.0 predicts that this sequence is a substrate for signal peptidase II (SPII), which is involved in lipoprotein maturation (32) (**Fig. 4a**). Conversely, when we analyzed the N-terminus of ChrA and MelA1 by SignalP, we observed that they contain secretion signals that are substrates for signal peptidase I (SPI), which does not result in lipidation (**SI Appendix, Fig. S22**) (32, 33). In peptides that contain a lipoprotein signal, the conserved cysteine and the three preceding residues constitute the lipobox motif (34). The precursor peptides that contain the additional third Cys residue all contain canonical lipobox motifs, thus explaining why the cysteine is invariable at that position (**Fig. 4a**). Therefore, following DybA modification by DybH in the cytoplasm (since DybH does not contain any signal peptide), the modified peptide is predicted to be transported to the inner plasma membrane, where a diacylglycerol (DAG) group is appended to the Cys thiol (**Fig. 4b**). The signal peptide is then cleaved off by signal peptidase II, and another acyl chain is often added to the N-terminal amine of the Cys residue in gram-negative bacteria (34).

**Fig. 4.**
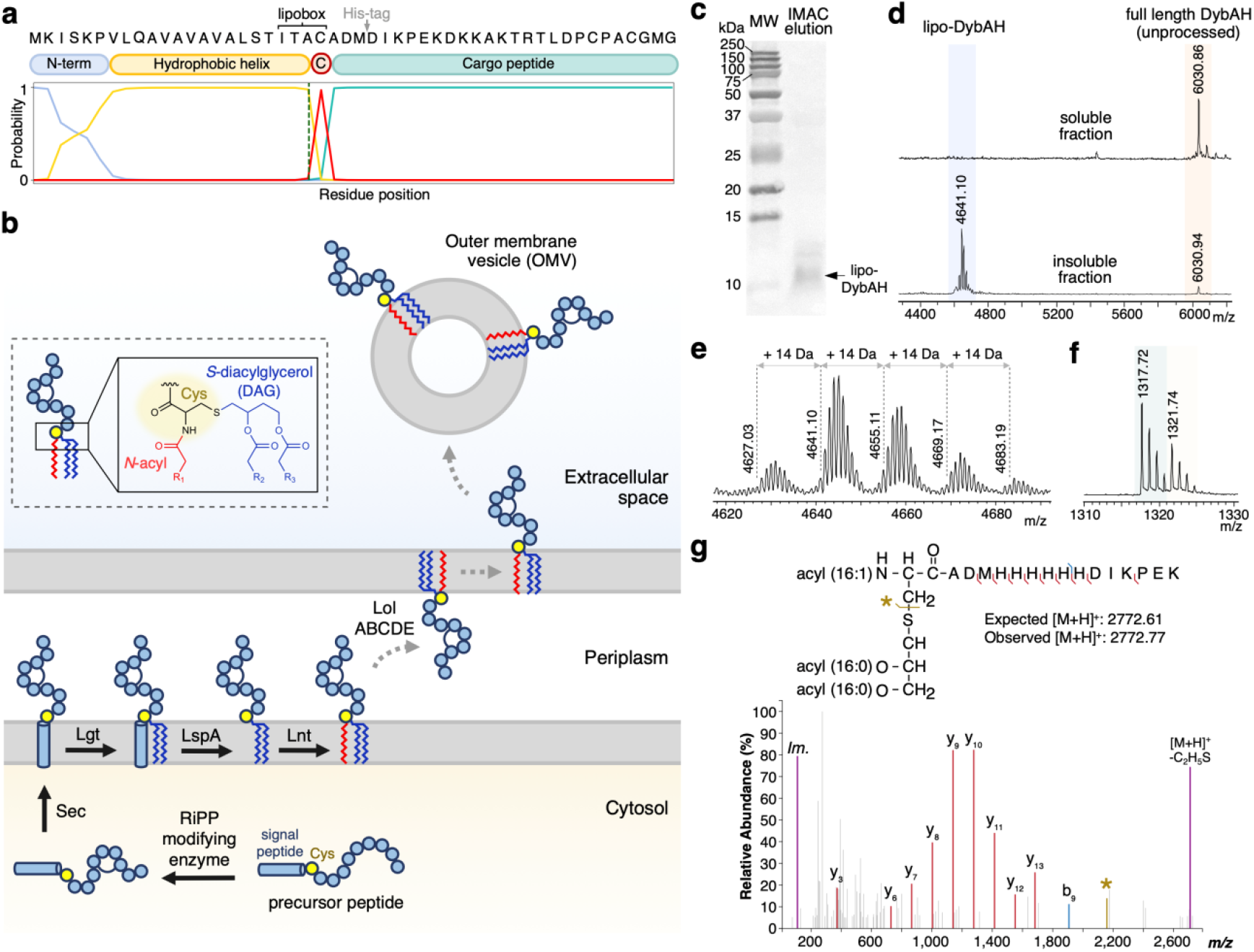
The product derived from *dybA* is a RiPP-lipoprotein. **a** SignalP 6.0 analysis of DybA showing the predicted N-terminal lipoprotein signal peptide. The graph shows the probability of each residue in the sequence as being part of the three subregions of the signal peptide system (the N-terminal positively charged region, the hydrophobic region, and the Cys to be modified) in addition to the predicted cleavage site (dashed line). A grey arrow shows the position of a 6xHis tag insertion in the construct that was used for heterologous expression of lipidated DybA. **b** Proposed biosynthetic pathway for DybA maturation via the gram-negative bacterial lipoprotein pathway and potential subcellular localizations. DybA is first modified by cytosolic DybH before modification with a diacylglyceryl group (blue) by prolipoprotein diacylglyceryl transferase (Lgt), signal peptide cleavage by lipoprotein signal peptidase (LspA/SPII), and *N*-acylation (red) by apolipoprotein N-acyltransferase (Lnt). Dashed arrows show potential transport of lipo-DybAH to the outer membrane and extracellular space. Inset: Structure of the modified N-terminal Cys in a triacylated lipoprotein. **c** Ponceau S stained nitrocellulose membrane showing the eluate from lipo-DybAH IMAC purification, following transfer from a 16.5% tris-tricine gel. An arrow indicates the band for lipo-DybAH, which was excised, digested by LysC, and analyzed by MS/MS. **d** MALDI-TOF MS of lipidated DybAH purified from *E. coli* by IMAC. The eluate precipitated and both the soluble (upper trace) and insoluble fraction (lower trace) were analyzed by MS. We observed full length DybA modified by DybH (but unprocessed by the lipoprotein pathway), in addition to mature, lipidated-DybAH. **e** Isolated lipo-DybA shows at least four differently lipidated species, with the major species containing C16:1, C16:0, C16:0 acyl chains. **f** LysC-digest of the total lipoprotein-enriched fraction shows that the lipidated species of DybA are partially unmodified (tan) and partially modified (teal) by DybH. The C-terminal DybA fragment upon LysC digest is shown. **g** MALDI-TOF tandem mass spectrum of the predominant 2772.77 Da N-terminal fragment obtained after excising the lipo-DybAH band and LysC digestion. These data show the triacylated form of lipo-DybAH [C16:1, C16:0, C16:0] when 2772.77 Da was selected as the parent ion. A fragment ion corresponding to loss of a C_2_H_5_S fragment from Met (47), and the immonium ion of histidine (*Im*.) were also detected.

The mature lipoprotein may have four different subcellular localizations: in the inner membrane, the outer membrane facing the periplasm, the outer membrane in a surface exposed manner, or in secreted outer membrane vesicles (OMVs) (**Fig. 4b**, dashed arrows). Inner membrane localization largely depends on a retention signal consisting of a negatively-charged residue immediately following the lipobox cysteine, at the +2 position (35). Without such a retention signal the majority of lipoproteins are transported to the outer membrane by the localization of lipoproteins (Lol) machinery (36). Furthermore, depending on the specific lipoprotein export signal, these products are surface exposed or secreted in the form of bacterial OMVs (37, 38).

To first confirm that the product of the *dyb* BGC is indeed a RiPP-derived lipoprotein, we removed the N-terminal His-tag to expose the putative signal peptide for processing. To facilitate purification, the His-tag was inserted between Met26 and Asp27 within the peptide sequence, which is after the lipoprotein signal peptide, but before the C-terminal CPACGMG motif to prevent interference with the modification by DybH (**SI Appendix, Fig. S23**). Following heterologous expression in *E. coli*, isolation of the total membrane fraction, solubilization, and purification by immobilized metal affinity chromatography (IMAC), a 16.5% tris-tricine gel showed the successful isolation of a small peptide (**Fig. 4c**). A precipitate formed in the IMAC eluate that was analyzed by MALDI-TOF MS. The spectrum showed a series of peaks that are consistent with lipidated forms of DybAH (**Fig. 4d**, lipo-DybAH). The observed ions were separated by 14 Da (CH_2_), consistent with the attachment of lipids with varying chain lengths (**Fig. 4e**). Digestion of this total lipoprotein fraction by LysC confirmed that the majority of the lipidated product is modified by DybH, showing the anticipated − 4 Da mass change in the C-terminal (non-lipidated) sequence (**Fig. 4f**). Conversely, analysis of the soluble fraction by MALDI-TOF MS showed enrichment of the DybH-modified full-length peptide (unprocessed by the lipoprotein machinery) (**Fig. 4d**).

To determine which lipids were attached to the *N*-acyl, *S*-diacylglyceryl-cysteine, the lipopeptide band from the 16.5% Tris-Tricine gel was transferred to a nitrocellulose membrane and excised for digestion with LysC (39) (**Fig. 4d**). Elution in a 2:1 (v/v) mixture of chloroform and methanol resulted in the digested N-terminal lipoprotein fragments. Tandem MS analysis of the predominant species by MALDI-TOF/TOF confirmed the formation of a triacylated lipoprotein that primarily contained a C16:1 acyl chain appended to the N-terminus of Cys, and a C16:0, C16:0-diacylglycerol group appended to the side chain of Cys (**Fig. 4g, SI Appendix, Fig. S24**). The lengths of the acyl chains are consistent with the lipids identified in previously characterized *E. coli* lipoproteins (40, 41). While these acyl chain lengths are affected by membrane lipid composition (42) (which likely differs in the native host *D. crusticola*), these results clearly show that DybA contains a signal peptide for lipidation via the ubiquitous bacterial lipoprotein biosynthesis pathway.

Analysis of the sequence logo of DybA-like peptides shows the presence of a ‘KDD’ signature at the +3 position and a short stretch of Asp residues (**Fig. 1d**), which have previously been shown in *Bacteroides* to be characteristic for cell surface exposed and OMV secreted lipoproteins, respectively (38, 43, 44). We therefore investigated whether heterologously expressed lipo-DybAH was present in secreted *E. coli* OMVs. OMVs were first isolated from the spent media by ultracentrifugation. Following lipoprotein extraction, we indeed detected lipo-DybAH in the OMVs (**SI Appendix, Fig. S25**). Cultivation and isolation from the native producing organisms will be required to confirm the prediction that these RiPP-lipoproteins are cell surface exposed and/or localized in OMVs, but members of the Bacteroidetes phylum and the Neisseria genus are known prolific producers of OMVs (45, 46).

### Bioinformatic mining for other RiPP-lipoprotein hybrids

A subset of bufferins (a group of MNIO-modified RiPPs) have recently been predicted to contain a signal peptide like the one in DybA suggesting they are lipidated (10). Therefore, we hypothesized that other bacterial RiPP-lipoprotein hybrids likely exist in nature that exploit the SPII pathway for lipidation (**Fig. 5a**). We developed a bioinformatic pipeline to facilitate the discovery of RiPP-lipoproteins, which we used here for the MNIO family (**Fig. 5b**). Two possible approaches can be envisioned for discovery of these hybrid products: either starting with lipoprotein prediction followed by RiPP prediction, or starting with RiPP prediction followed by lipoprotein prediction. The latter approach ensures that the resulting peptides are within or adjacent to RiPP BGCs (thus RiPP-related) and narrows down the initial list of putative lipoproteins significantly. The list of precursor peptides can be filtered based on the presence of a lipidation signal, and an SSN of the putative RiPP-lipoproteins can then be used to group hits with similar gene contexts.

**Fig. 5.**
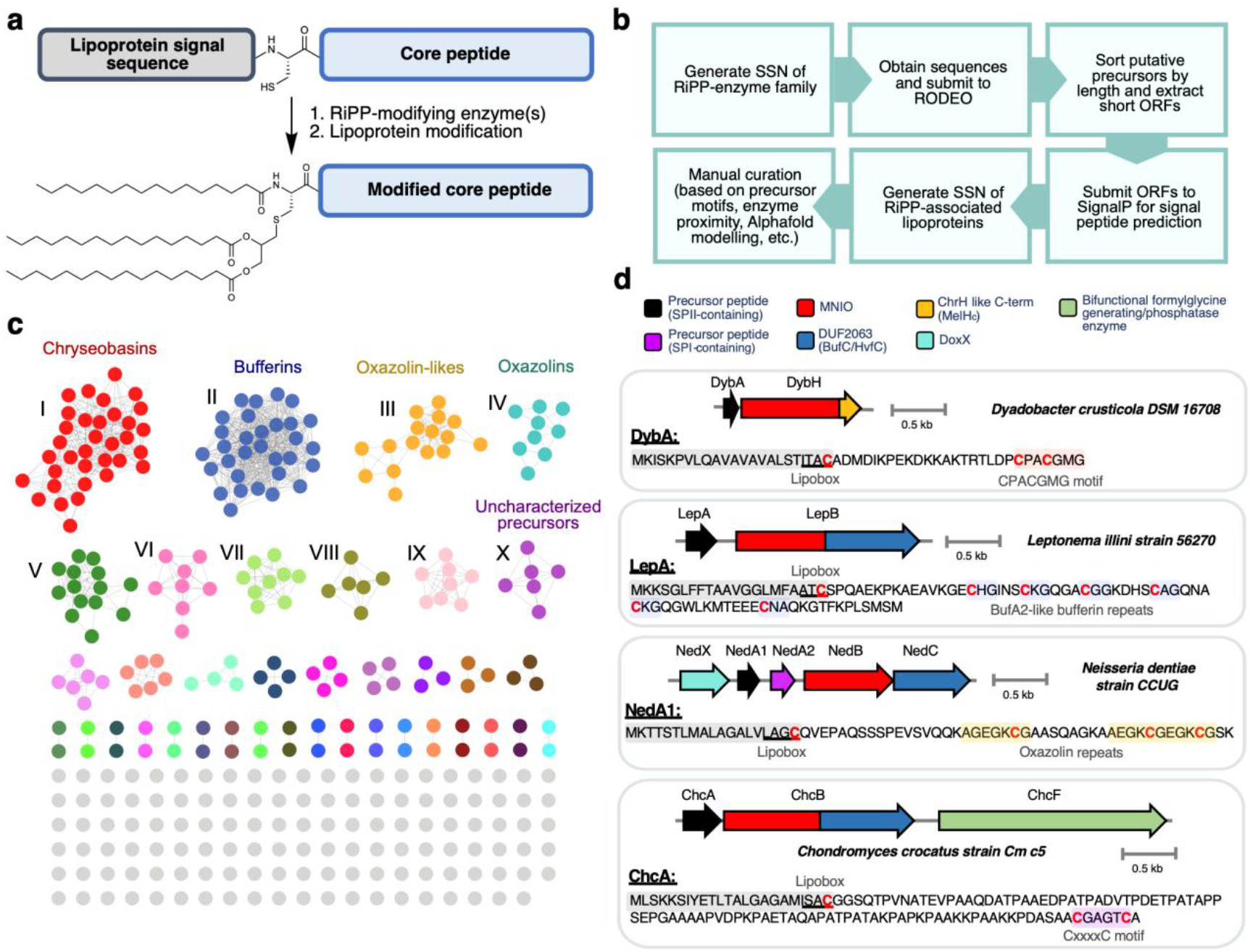
Genome mining for other RiPP-lipoproteins. **a** Schematic example of RiPP-lipoprotein maturation to the triacylated form (C16:0, C16:0, C16:0). **b** Bioinformatic pipeline for RiPP-directed discovery of RiPP-lipoproteins. During the SSN generation process, similar sequences can be filtered out using Uniref90 database or using a representative node network. **c** SSN of MNIO-associated lipoproteins (Dataset 2). Ten of the largest clusters are annotated I-X. Clusters V-IX correspond to putative non-RiPP lipoproteins that are found in (or adjacent to) MNIO-containing gene clusters. V: BON-domain containing protein in bufferin BGC. VI: LPS translocon maturation chaperone LptM in bufferin BGC. VII: Glycine zipper two transmembrane domain-containing protein in bufferin BGC. VIII: TssQ family T6SS-associated lipoprotein in bufferin BGC. IX: Hypothetical protein in bufferin BGC. The full list of sequences can be found in **SI Appendix, Table S6. d** Representative examples of predicted BGCs encoding RiPP-lipoproteins from the different precursor peptide clusters. The characterized RiPP families corresponding to chryseobasins, bufferins, and oxazolins all show examples of lipidation via the SPII pathway.

We utilized this approach to mine for precursor peptides that co-occur with MNIOs. From the local gene neighborhood of genes encoding MNIOs obtained from RODEO (48), ORFs were extracted and sorted based on length. A length of 50-160 amino acids was used as the cutoff for putative precursor peptides. These peptides were then analyzed for a lipoprotein signal peptide using SignalP 6.0 (32). An SSN of the putative RiPP-lipoproteins was then generated to group together related sequences (**Fig. 5c**).

As expected, chryseobasin-like peptides (the products of *chr*-like BGCs from Bacteroidota) represented one of the largest clusters of precursor peptides predicted to be lipidated (**Fig. 5c and d**, Cluster I). Additionally, a subset of the previously characterized copper-chelating peptide families (both bufferins (13) and oxazolins (12)) contained SPII lipoprotein signals. We also found a group of uncharacterized, putative precursor peptides that contain lipoprotein signals (**Fig. 5c and d**, Cluster X). These peptides contain a highly conserved CxxxxC motif that is the likely site for MNIO modification.

Our genome mining output predicted that NedA from *Neisseria dentiae* CCUG 53898 310003 (WP_085365104.1) is a lipidated, copper-binding peptide, as it contains both the SPII lipoprotein signal and the conserved repeated <u>EGKCG</u> motifs, previously shown to be modified to a copper-binding structure (12) (**Fig. 5d**). To confirm that NedA is a RiPP-lipoprotein, we heterologously expressed in *E. coli* the C-terminally His-tagged NedA with its modifying enzymes (the MNIO NedB and its substrate-binding partner protein NedC). Using the same lipoprotein purification method as for lipo-DybAH, we successfully isolated tri-acylated lipo-NedABC from the insoluble membrane fraction (**Fig. 6a and b**). Digestion with LysC, and MS/MS analysis of the major N-terminally lipidated fragment showed that the attached lipids are derived from palmitic and palmitoleic acids [16:1, 16:0, 16:0] (**Fig. 6c and SI Appendix, Fig. S26**). In addition, MS analysis of the core peptide showed a − 12 Da modification, which is consistent with a − 4 Da modification at each of the three <u>EGKCG</u> motifs by NedBC (**SI Appendix, Fig. S26**). Whether the modification at the <u>EGKCG</u> motif corresponds to formation of a thioamide-oxazolone or 5-thiooxazole will require more investigation given the different assignments made by other groups (12, 14, 15). Nevertheless, these experimental results confirm our initial bioinformatic prediction, showing that the product derived from NedA is a RiPP-lipoprotein. Since the related peptide MmrA from *Microbulbifer* sp. VAAF005 (which contains the conserved <u>EGKCG</u> repeats) is proposed to be involved in protection against copper toxicity (15), it is possible that lipo-NedABC sequesters copper at the membrane to prevent entry into the cell. Furthermore, copper-bound lipopeptides could be secreted from the cell in the form of OMVs.

**Fig. 6.**
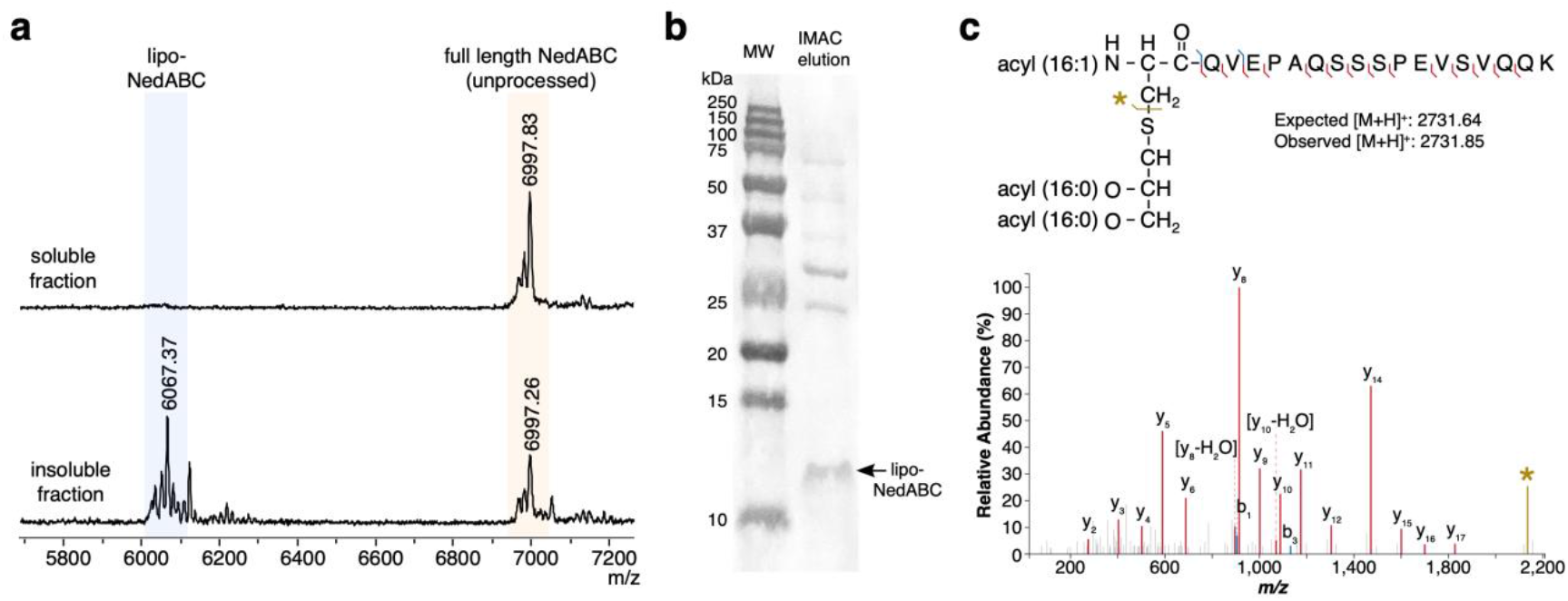
Heterologous production of lipo-NedABC in *E. coli*. **a** MALDI-TOF mass spectra of IMAC-purified C-terminally His-tagged NedA that was coexpressed with NedBC. Following IMAC purification, the soluble fraction (upper trace) and insoluble pellet (lower trace) that formed were analyzed by MS (average masses shown). We observed full length NedA modified by NedBC (but unprocessed by the lipoprotein pathway), in addition to mature, lipidated-NedA processed by NedB. **b** Ponceau S stained nitrocellulose membrane with the arrow pointing to the band corresponding to lipo-NedABC, which was excised for LysC digestion and MS/MS identification. The IMAC eluate was first run on a 16.5% tris-tricine gel, then transferred to the nitrocellulose membrane. **c** MALDI-TOF tandem mass spectrum of the excised lipo-NedABC nitrocellulose band following LysC digestion, showing the triacylated form of lipo-NedABC [C16:1, C16:0, C16:0].

These data show the application of the RiPP-lipoprotein mining approach for the MNIO family, but this method could be applied to any RiPP-modifying enzyme family. RiPPs most likely to be lipidated via this pathway involve short biosynthetic pathways that are leader-peptide independent or that use follower peptides (2). Conversely, RiPPs that rely on N-terminal cleavage or cyclization, such as lasso peptides and bottromycins, are likely incompatible with an N-terminal signal peptide (1). Aside from the MNIO-modified precursors, our preliminary bioinformatic analyses also show that a subset of autoinducing peptides (AIPs) (49) appear to contain canonical SPII signal peptides (**SI Appendix, Fig. S27**). In AIP biosynthesis, the thiol of a conserved internal cysteine typically cyclizes with the C-terminal carboxylate to generate a thiolactone macrocycle (catalyzed by the modifying enzyme AgrB) (50). We found examples of unusual AIP precursors that contain two cysteines, suggesting that the first cysteine is used as the glycerolipid attachment site, and the second for macrocyclization (**SI Appendix, Fig. S27, SI Appendix, Table S7**). However, not all predicted SPII-signal-containing AIPs contain two cysteine residues. Hence, whether these particular AIPs truly form lipoproteins (or whether the glycerolipid attachment site is blocked by formation of the thiolactone macrocycle) is presently not clear.

## Discussion

The MNIO family is a rich source for diverse post-translational modifications of peptides (8, 10). Here, we present a ChrH-like modification performed by MNIOs from Bacteroidota and Cyanobacteriota. The diverse substrates of this subfamily include secreted peptides, secreted proteins, and lipoproteins that share a C-terminal modification. The two previous examples of thiohemiaminals formed by MNIOs involved alkylation reactions of the thiol. In one example, the alkylation involved carboxymethylation by a dedicated carboxy-*S*-adenosyl methionine dependent enzyme encoded in the BGC (16, 51). In the second example, the MNIO itself appeared to methylate the thiol of the thiohemiaminal although methylation by a protein in *E. coli* cannot be ruled out (17). Conversely, DybH and its orthologs reported herein form the thiohemiaminal as a stable product that is not further alkylated based on the genes in their BGCs and the outcome of the co-expression experiments in *E. coli*. The function of the final products is currently still unclear.

Recognition of the minimal C-terminal motif likely provides relaxed purifying selection on the N-terminal sequence (which is typically constrained in RiPP biosynthesis by a mandatory leader peptide), thus explaining the broad structural diversity observed at the N-termini of these substrates. For instance, the motif is modified when present at the C-terminus of the protein MelA1.

As noted above, the biological function of the lipoprotein products is still unknown. In both Cyanobacteria and *Chryseobacterium*, these peptides are secreted via SPI, suggesting a periplasmic or extracellular function. In other Bacteroidota genera, the lipoproteins are membrane bound or could be released in OMVs. The BGC in *Spirosoma endophyticum* encodes a prolyl oligopeptidase with a secretion signal, which could cleave the lipidated peptide (**SI Appendix, Fig. S28**). Furthermore, the gene clusters frequently co-occur with or contain genes encoding SusC and SusD type importers. The SusCD import machinery is involved in the uptake of oligopeptides or oligosaccharides, and are commonly found in polysaccharide utilization domains (52). We also note that the macrocyclic ring of the highly conserved MNIO products reported here is structurally similar to the conserved B ring of lantibiotics such as nisin (53), epidermin (54), and mutacin 1140 (53-55) (**SI Appendix, Fig. S29**). The B-ring of these lantibiotics is part of the binding site of the pyrophosphate of the lipid II target (56). Future work will focus on elucidating the function of the lipoproteins.

The discovery of RiPP-lipoprotein hybrids demonstrates an example of specialized metabolism co-opting pathways involved in the general maturation of proteins. Our initial bioinformatic searches show that several products from other MNIO subfamilies are predicted to be lipidated via the lipoprotein pathway. These results also suggests that additional RiPP-lipoproteins, beyond those modified by MNIOs, may exist in nature. The observation here that lipidated DybA and NedA are modified by their RiPP maturation enzymes demonstrates that their post-translational modification (by DybH and NedBC respectively) is sufficiently fast to compete with export via the membrane bound lipoprotein pathway. In our identified examples of RiPP-lipoprotein BGCs, we did not encounter any MNIOs with secretion signals, supporting the biosynthetic order in which RiPPs are first modified by MNIOs in the cytoplasm, and then exported across the inner membrane and lipidated (**Fig. 4b**). Other classes of modifying enzymes could act on lipoproteins in the periplasm, but at present no evidence of this biosynthetic sequence has been reported.

Several characterized bacterial lipoproteins are immunogenic and potent activators of Toll-like receptors (42, 57). The use of RiPP-modifying enzymes could enable post-translational modifications of immunogenic lipoprotein adjuvants to expand their structural diversity beyond the canonical 20 amino acids and improve pharmacological stability.

## Methods

### Heterologous expression and purification of peptides

*E. coli* BL21 (DE3) cells were transformed with the appropriate plasmids and plated on selective media (**SI Appendix, Table S8**). Transformants were inoculated in LB and cultured overnight with appropriate antibiotic(s) at the following concentrations: 33 µg/mL chloramphenicol, 50 µg/mL streptomycin, 50 µg/mL kanamycin. The overnight culture was used to inoculate either 200 mL of LB (small scale) or 4 L of terrific broth (TB) media (large scale). The cultures were grown at 37 °C with shaking at 240 rpm until the OD_600_ reached 0.7-0.9. The flasks were cooled on ice for 20-30 min. For co-expressions with MNIOs, a freshly prepared 1000× stock solution of iron(II) sulfate heptahydrate and trisodium citrate was added to a final concentration of 350 μM and 1 mM, respectively. IPTG was added to 0.7 mM final concentration, and the flasks were placed back in the shaker at 18 °C, 220 rpm, for 18-20 h. The cells were harvested by centrifugation at 7000 g for 10 min, washed once with distilled (DI) water, then stored at − 80 °C until further use.

Cell pellets were thawed at room temperature, then an equal volume of DI water was added (containing 0.5× Pierce EDTA-free protease inhibitor). After stirring at 4 °C for 30 min, four volumes of denaturing lysis buffer (6 M guanidine hydrochloride, 20 mM sodium phosphate dibasic, 500 mM sodium chloride, 10 mM imidazole, pH 7.5) was added. The cell solution was sonicated until clear (10 s on, 10 s off cycle, 30 min total). The lysate was clarified by centrifugation at 40,000 g for 30 min. The supernatant was decanted into 0.5 mL (small scale) or 5 mL (large scale) of equilibrated Ni-NTA agarose beads, and nutated at 4 °C for 30 min. The beads were transferred to a column and washed sequentially with 20 column volumes (CV) of 4 M guanidine hydrochloride, 20 mM sodium phosphate dibasic, 300 mM sodium chloride, 30 mM imidazole, pH 7.5 followed by 20 CV of 20 mM sodium phosphate dibasic, 500 mM sodium chloride, 30 mM imidazole, pH 7.5. The peptide was eluted using 10 CV of 20 mM sodium phosphate dibasic, 500 mM sodium chloride, 500 mM imidazole, pH 7.5.

For small scale purifications, the IMAC eluate was directly desalted with a C18 Ziptip. For large scale purification, the eluate was desalted using a 2 g C18 Hypersep SPE column, and eluted in 50-60% acetonitrile with 0.1% trifluoroacetic acid (TFA). The desalted peptide was concentrated to dryness under reduced pressure. To obtain full length peptide, the residue was directly resuspended in water and injected onto an Agilent HPLC equipped with an analytical Grace Vydac C18 column (250 mm x 4.6 mm, 300 Å pore size, 5 μm particle size) at a flow rate of 1 mL/min with the following gradient: 2% MeCN in water with 0.1% TFA for 4 min, 2-50% MeCN in water with 0.1% TFA for 30 min, and a gradient of 50-90% MECN in water with 0.1% TFA over 15 min). Fractions containing the desired peptide, judged pure MALDI-TOF MS, were combined and lyophilized. The peptide was then stored at − 20 °C and resuspended in water when needed.

### Isolation of the N-terminally lipidated fragment for MALDI-TOF MS/MS analysis

The N-terminal lipid modification was confirmed by tandem mass spectrometry using a variation of a method described previously (39). The IMAC eluate was first run on a 16.5% tris-tricine SDS-PAGE gel (BioRad). Proteins were transferred to a 0.2 μm nitrocellulose membrane using a wet transfer method for 1 h at 110 mV. Ponceau S was used to visualize the peptide bands, which were excised and transferred to a protein low-bind microfuge tube (Eppendorf). The membrane pieces were fully destained with two washes of water and three washes of 50 mM ammonium bicarbonate, pH 7.8. Endoprotease LysC (10 μL of a 20 μg/μL solution in ammonium bicarbonate) was added to cover the nitrocellulose pieces, and the digest was incubated for 16 h. At the final hour of digestion, DTT was added to 0.1 mM. The liquid was removed, followed by sequential washings with 10 uL of 10%, 20% MeCN + 0.1% TFA, and 5 uL of and 80% MeCN + 0.1% TFA. Finally, 5 μL of a mixture of CHCl_3_:MeOH (2:1 v/v) was used to elute the lipopeptides from the nitrocellulose membrane. The mixture (1 μL) was mixed with 1 μL of a 15 mg/mL α-cyano-4-hydroxycinnamic acid (CHCA) matrix dissolved in CHCl_3_:MeOH (2:1 v/v) and spotted on a MALDI target plate. The synthetic lipoproteins Pam2CSK4 and Pam3CSK4 were used as standards to validate the efficacy of the method.

## Acknowledgments

We thank Dr. Chandrashekhar Padhi for assistance with LC-MS/MS experiments. This work was supported by a grant from the National Institutes of Health (R37 GM058822 to W.A.v.d.D.) and an award from the Natural Sciences and Engineering Research Council of Canada (PGS-D to J.Y.C). J.Y.C. is a fellow of a Chemistry-Biology Interface Training Grant (5T32-GM070421) from the National Institute of General Medical Sciences. A Bruker UltrafleXtreme mass spectrometer used was purchased with support from the National Institutes of Health (S10 RR027109).

