## Supporting information for "Co-opting the bacterial lipoprotein pathway for the biosynthesis of lipidated macrocyclic peptides"

**This PDF file includes:**

Supporting Materials and Methods
Figures S1 to S29
Tables S1 to S10
SI References

### Materials and Methods

### Materials and Methods

*Materials*

Primers and gene fragments were ordered from Integrated DNA Technologies and Twist Biosciences, respectively. Antibiotics, IPTG, DTT, TCEP, and lysozyme were purchased from GoldBio. LysC endoprotease was ordered from New England Biolabs (NEB). Deuterated NMR solvents were sourced from Cambridge Isotope Laboratories. Q5® High-Fidelity DNA Polymerase, NEBuilder® HiFi DNA Assembly, and DpnI restriction enzyme were purchased from NEB. Pam2CSK4 and Pam3CSK4 synthetic lipoprotein standards were purchased from InvivoGen. The MALDI matrices super-DHB and CHCA were obtained from Millipore Sigma. All accession numbers for proteins in this study are provided in **SI Appendix, Table S9**.

*Cloning*

All plasmid constructs were initially assembled by Gibson assembly of DNA fragments with PCR-amplified vector backbones (primers and gblocks listed in **SI Appendix, Table S10**). The genes *dybA* and *dybH* were directly amplified from the extracted gDNA of *Dyadobacter crusticola* strain DSM 16708. Prior to DNA extraction, *D. crusticola* was cultured in R2B media, shaking at 220 rpm at 28 °C to late mid-log phase. The other genes used in this study were obtained by synthetic gene fragments (**SI Appendix, Table S10**). Subsequent changes to the plasmids (site-directed mutagenesis, deletions, and short insertions) were performed by PCR with partially overlapping primers, and then digesting the template plasmid with Dpn1. The unpurified digestion reaction (2 µL) was directly used to transform *E. coli* 5α (NEB) chemically competent cells. The sequences of all plasmids were verified by Plasmidsaurus prior to transformation of the *E. coli* BL21 (DE3) expression strain.

*Co-expression experiments*

For co-expression experiments, *E. coli* BL21 DE3 cells were transformed with a pACYCDuet-1 plasmid encoding the His-tagged precursor peptides (ChrA, DybA, ChsA, HymA, Table S8) as well as a pCDFDuet-1 plasmid encoding the cognate non-tagged modification enzyme (ChrH, DybH, ChsH, HymH). For the experiments with MelH, His-tagged MelA1 was encoded in pACYCDuet-1 and either untagged MelH or untagged MelH and $MelH_c$ were encoded in pCDF-Duet-1. For ArlH, the His-tagged ArlA substrate and the untagged ArlH modifying enzyme were both encoded in pRSF-Duet-1. For expression conditions and peptide purification, see the main text.

*LysC digestion*

To obtain digested peptide, the lyophilized, desalted full length peptide was first resuspended in 1 mL of 10 mM Tris-HCl buffer pH 8, with 2 mM TCEP, to which 20 µL of LysC (0.1 µg/µL, NEB) was added. The digest was incubated at 37 °C overnight. The digest was then injected onto HPLC using the same conditions as above with the following modification to the gradient: 2% B for 4 min, then 2-30% B for 30 min, 30-90% B for 15 min). Fractions containing the desired peptide fragment were lyophilized.

*Heterologous expression and purification of DybH*

BL21 (DE3) cells were transformed with pET28a:His-TEV-DybH. Protein expression was carried out as with peptide expression described above. The cell pellet was resuspended in protein lysis buffer (50 mM HEPES, 200 mM NaCl, pH 7.6) containing 0.5x Pierce EDTA-free protease inhibitor, 2 mg/mL lysozyme, and 10 µg/mL DNAse I. Cells were stirred at 4 °C for 15 min, then lysed by sonication (10 s on, 10 s off cycle, 20 min total). Following centrifugation at 40,000 g for 40 min, the clarified lysate was loaded onto equilibrated HisPur™ Nickel-nitrilotriacetic acid resin, and washed with 20 CV of lysis buffer. The protein was eluted in the same buffer containing 300 mM imidazole. The fractions containing the most protein were pooled and immediately injected onto a HiLoad 16/60 Superdex 200 pg column equilibrated with storage buffer (25 mM HEPES, 200 mM NaCl, pH 7.5, 10% glycerol). Fractions containing purified protein, judged pure by SDS-PAGE, were pooled and concentrated to approximately 20 mg/mL, at which point they were flash frozen in liquid $N_2$ and stored at −80 °C.

*In vitro assays*

To 50 µL of assay buffer (25 mM HEPES, 200 mM NaCl, pH 7.6) was added DybA (75 µM), TCEP (1 mM) and DybH to a final concentration of 15 µM. The assay was incubated at room temperature for 1 h before the addition of acetonitrile to a final concentration of 10 % and TFA to 0.2 %. The reaction product was desalted by a C18 Zip tip and analyzed by MALDI-TOF MS.

*Iodoacetamide alkylation*

The peptide was resuspended in 50 mM Tris, pH 8.3. Dithiothreitol was added to 6 mM final concentration and the reaction was incubated at 55 °C for 10 min. Iodoacetamide was added to 20 mM final concentration, and the reaction was placed at RT in the dark for at least 1 h. The alkylated peptide was purified by a C18 Ziptip, or by C18 HPLC as described previously.

*LC-MS/MS*

Desalted samples were analyzed on an Agilent 6545B Q-TOF and 1290 Infinity II LC system using a Phenomenex Aeris C18-XB column (150 mm x 4.6 mm, 100 Å pore size, 2.6 µm particle size) at a flow rate of 0.6 mL/min, and the following gradient: 5% ($H_2O$ + 0.1% FA:MeCN + 0.1% FA) for 3 min, 5-70% for 12 min, 70-90% for 3 min. The collision energy was set to 25-30 mV and ions were detected in positive mode. Fragments were initially assigned by using the Interactive Peptide Spectral Annotator (IPSA) (1) and manual curation.

*MALDI-TOF MS and MS/MS*

Samples were desalted using a C18 Zip tip (0.6 µL bed volume) then eluted in 1 µL of 80% acetonitrile containing 0.1% TFA and 50 mg/mL of the MALDI matrix super-DHB. The eluate was spotted on a plate, dried, and analyzed on a Bruker UltrafleXtreme MALDI-TOF instrument using positive reflectron mode (RP) for peptides, or linear mode (LP) for proteins. Masses were calibrated using a set of peptide and protein standards (bradykinin, angiotensin II, ACTH clip 18-39, insulin, cytochrome C, and bovine serum albumin). MS/MS spectra were collected in LIFT positive mode.

*NMR spectroscopy*

Lyophilized digested peptide was resuspended in either 280 µL of 90% $H_2O$:10% $D_2O$ or 100% $D_2O$ and transferred to a $D_2O$ matched Shigemi tube, or in 100% DMSO-d6 in a DMSO-d6 matched Shigemi tube. The $^1H$ NMR spectrum was collected on a 750 MHz Agilent VNMRS with a 5 mm HCN probe. HSQC, HMBC, and NOESY data was collected on a 600 MHz Bruker NEO spectrometer with a 5 mm broadband Prodigy CryoProbe. Mnova was used for data analysis and chemical shifts were referenced to the solvent.

*Ferene S iron quantification assay*

Total iron in enzyme samples was quantified against a standardized iron solution (1000 mg/L Fe in 0.5 M $HNO_3$, Certipur® certified reference material) using the ferene S reagent, in a similar manner as described previously (2, 3). Briefly, to 300 µL of diluted enzyme sample and iron standards was added 300 µL of reagent A (155 mM SDS, 75 mM iron-free sodium acetate). The solution was thoroughly mixed, then 300 µL of reagent B (1 M sodium ascorbate, 8 mM sodium metabisulfite, 50 mM iron-free sodium acetate) was added. The solution was thoroughly mixed and incubated at 37 °C for 15 min. Then, 15 µL of reagent C (18 mM ferene S) was added, and the absorbance was read at 593 nm. The iron concentration in enzyme samples was determined using a standard curve of serial dilutions of the iron standard. The resulting iron concentration was normalized to the protein concentration (determined by the BCA assay) to yield the number of iron atoms per enzyme protomer.

*Phylogenetic and bioinformatic analyses*

Protein sequences were uploaded to MEGA11 (4) and the MUSCLE algorithm was used to align sequences. Using MEGA11, the aligned sequences were used to build a maximum likelihood phylogenetic tree using the Jones-Taylor-Thornton model and 90% partial deletion. For prediction of signal peptides in DybA or ChrA, individual sequences were submitted to the SignalP 6.0 server (5). For genome mining of MNIO-RiPP-lipoproteins, an SSN for the MNIO family (PF05114) was first constructed using the EFI-EST program (6, 7), against the UNIREF90 database. The alignment score was set to 30, and enzymes were filtered based on length from 220 to 50,000 amino acids (default maximum length). The MNIO sequences retained from this filter totalled 8227 sequences. The Uniprot accessions were converted to RefSeq Protein accessions (using the ID Mapping tool from Uniprot), resulting in 4360 converted sequences, which were submitted to RODEO (8). The predicted precursor open reading frames (ORFs) were sorted by length, and peptides between 50-160 amino acids in length starting with a methionine residue were selected. These peptides were submitted to SignalP 6.0 for signal sequence prediction (5). To generate an SSN of putative lipidated RiPPs, a list of all peptides with a predicted SPII signal sequence of over 95% confidence (339 total) were submitted to EFI-EST, using an alignment score of 1. Cytoscape 3.10.3 was used for network visualization, using yFiles Organic layout (9).

*Lipoprotein expression and purification*

Lipoprotein constructs were transformed and expressed in *E. coli* BL21 (DE3) cells, as described in the methods in the main text, with the exception that LB was adjusted to pH 8, and at induction, only IPTG was added to 0.7 mM, without additional iron supplement. The cells were incubated shaking at 30 °C for 12 h at 220 rpm.

For purification, cells were first resuspended in lysis buffer (25 mM HEPES, 300 mM NaCl, pH 7.5). Lysozyme (2 mg/mL lysozyme) and DNAse I (10 µg/mL) were added, and the mixture was incubated at room temperature for 5 min, then sonicated (10 s on, 10 s off cycle, 15 min total). Unbroken cells were removed by centrifugation at 7000 *g* for 8 min. To collect membranes, the supernatant was subjected to ultracentrifugation in a Beckman TL-100 ultracentrifuge at 100,000 *g* for 20 min. The supernatant was discarded, and the membrane was resuspended in 0.5× initial volume of buffer by pipetting up and down to homogenize. To the mixture was added *n*-octyl-β-d-glucopyranoside (OG) and Zwittergent 3-14 to 1% (w/v) and 0.5% (w/v) respectively. The addition of Zwittergent 3-14 has been previously shown to help solubilize lipoproteins from membranes (48). The mixture was incubated for 30 min nutating at RT. Insoluble precipitates that remained were removed by centrifugation at 100,000 *g* for 10 min. The supernatant was loaded onto 250 µL of Ni resin in a 5 mL tube and nutated at 4 °C for 1 h. The resin was transferred to a chromatography column and washed with lysis buffer containing 0.05% Zwittergent 3-14 and 30 mM imidazole. For the final 20 mL of wash and elution (containing 500 mM imidazole), no detergent was added. The elution was collected in 100-µL fractions, several of which formed precipitates (especially after freeze-thawing).

If the eluate precipitated (which can take several hours at 4 °C or freeze-thawing), then a pellet was obtained by centrifugation at 16,000 *g* for 10 min. The supernatant was removed, and the pellet was gently washed with 50 mM ammonium bicarbonate pH 7.8 twice to desalt the sample. To solubilize the pellet, 100 µL of 0.1% TFA was added, and the mixture was sonicated in a water bath. The sample (1 µL) was mixed with 1 µL of 80% acetonitrile containing 0.1% TFA and 50 mg/mL of super-DHB then spotted on a MALDI target plate.

If the eluate did not precipitate, it was desalted on a PD MiniTrap G-10 column (Cytiva) into milliQ water. The sample was concentrated using a SpeedVac vacuum concentrator, then (1 µL) was directly mixed with 1 µL of 80% acetonitrile containing 0.1% TFA and 50 mg/mL of super-DHB then spotted on a MALDI target plate.

*Isolation of outer membrane vesicles (OMVs)*

OMVs were isolated from *E. coli* as described previously (10). *E. coli* BL21 (DE3) cells heterologously expressing DybAH were grown in 1 L of LB media (pH 8) with shaking (240 rpm) at 37 °C, and induced at an OD of 0.6 by addition of IPTG to a final concentration of 0.7 mM. The culture was placed back in the incubator shaker at 37 °C for 18 h, shaking at 220 rpm. Cells were then removed from the supernatant by centrifugation at 7000 g for 15 min. To remove any remaining cells or large cellular debris in the supernatant, the supernatant was filtered through a 0.2 µm PES Nalgene™ Rapid-Flow™ Sterile Disposable Filter (Thermo Scientific™). The filtrate was then concentrated using 100 kDa MWCO Amicon centrifugal filters to a final volume of 3 mL. The concentrated supernatant was transferred to a polypropylene tube and ultracentrifuged at 160,000 g for 18 h at 4 °C. The next day, the supernatant was discarded and the pellet was gently washed with storage buffer (25 mM HEPES, 150 mM NaCl, pH 7.6) to yield a small translucent OMV pellet. To isolate lipoproteins from OMVs, the pellet was processed in the same way as the total membrane pellet, as described above.

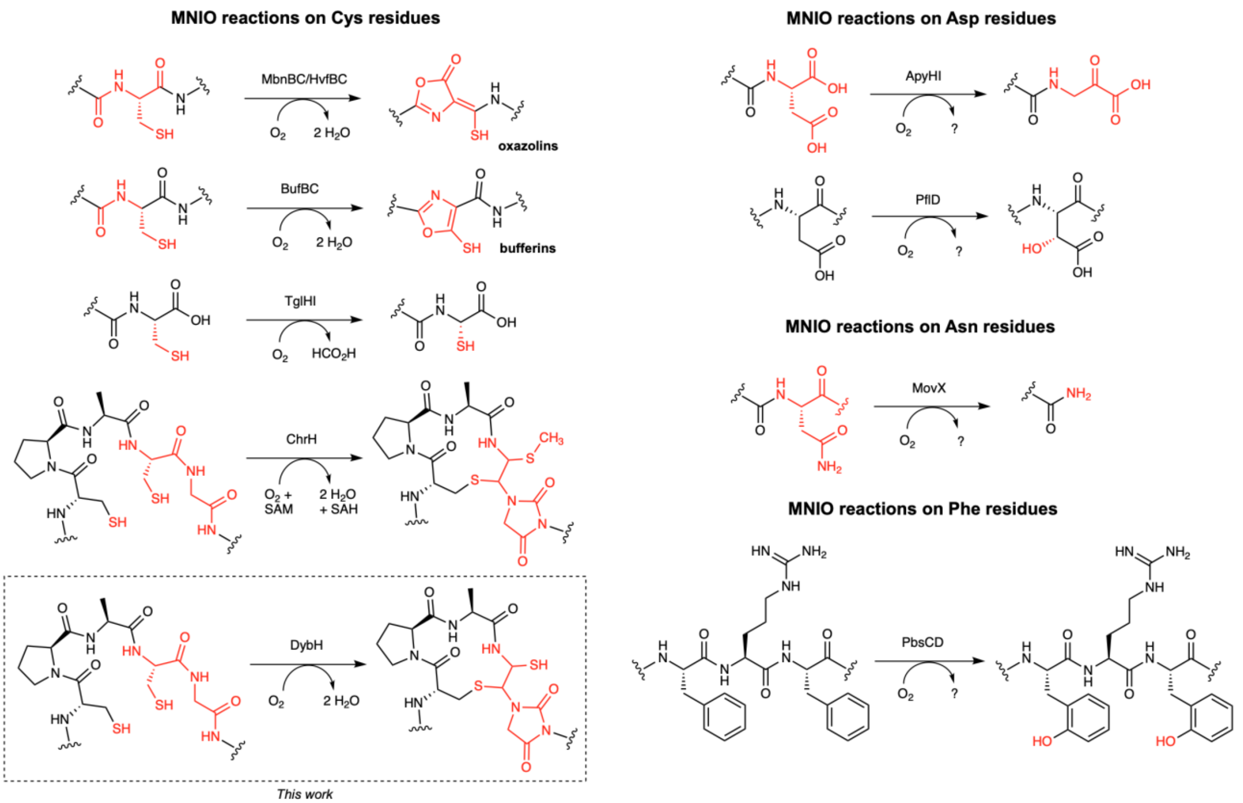

**Fig. S1. Characterized reactions catalyzed by multinuclear non-heme iron dependent oxidative enzymes (MNIOs).** Currently characterized MNIOs act on cysteine, aspartate, asparagine, and phenylalanine residues within conserved motifs in a peptide or protein sequence (11-20).

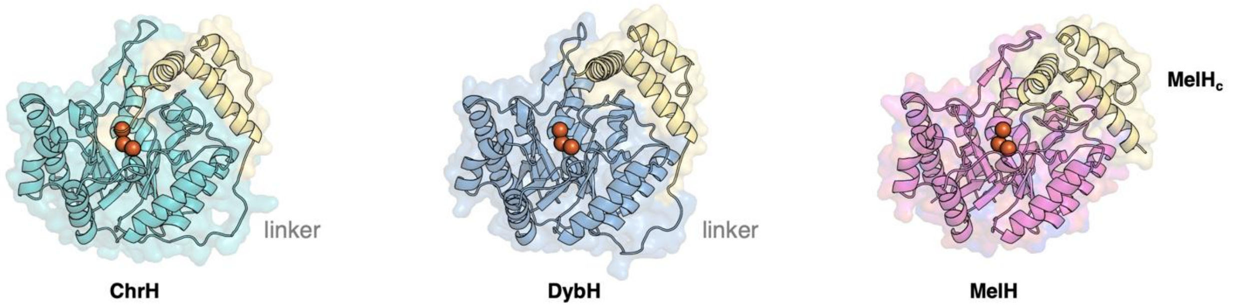

**Fig. S2. Alphafold3 models of ChrH, DybH and MelH with MelH$_c$.** Predicted structures showing conservation of the core MNIO TIM-barrel in addition to the C-terminal domain (yellow). In MelH, the C-terminal domain is encoded as a separate polypeptide (MelH$_c$). All proteins were modelled with three $Fe^{3+}$ ligands.

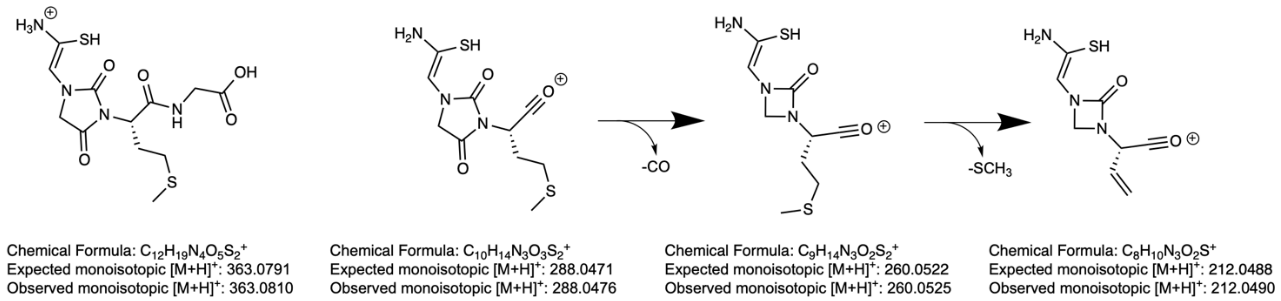

**Fig. S3. Assignment of fragmentation ions from modified DybAH observed in LC-MS/MS.** Potential modified y-ions observed at 30 eV collision energy of LysC-digested DybAH (Fig. 2b, main text). The exact mass and observed monoisotopic masses are shown below the structures.

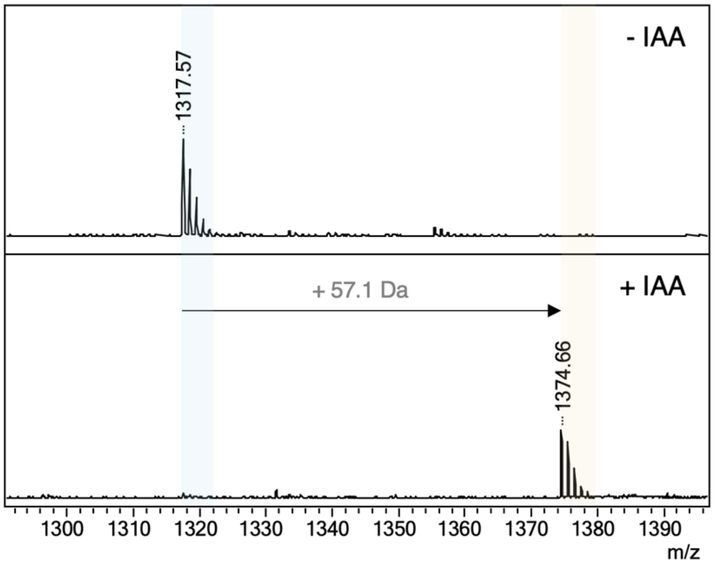

**Fig. S4. IAA-alkylation of LysC-digested DybAH.** MALDI-TOF MS data of LysC-digested DybAH before and after alkylation with iodoacetamide (IAA), showing a mass increase of 57 Da, corresponding to one carbamidomethylene adduct, thus suggesting that one free thiol remains in the product. The C-terminal LysC fragment corresponds to a 13-mer peptide (sequence: TRTLDPCPACGMG) modified by DybH. Expected monoisotopic $[M+H]^+$: 1317.54 Da. Expected monoisotopic $[M+H]^+$ of IAA-alkylated peptide: 1374.56 Da.

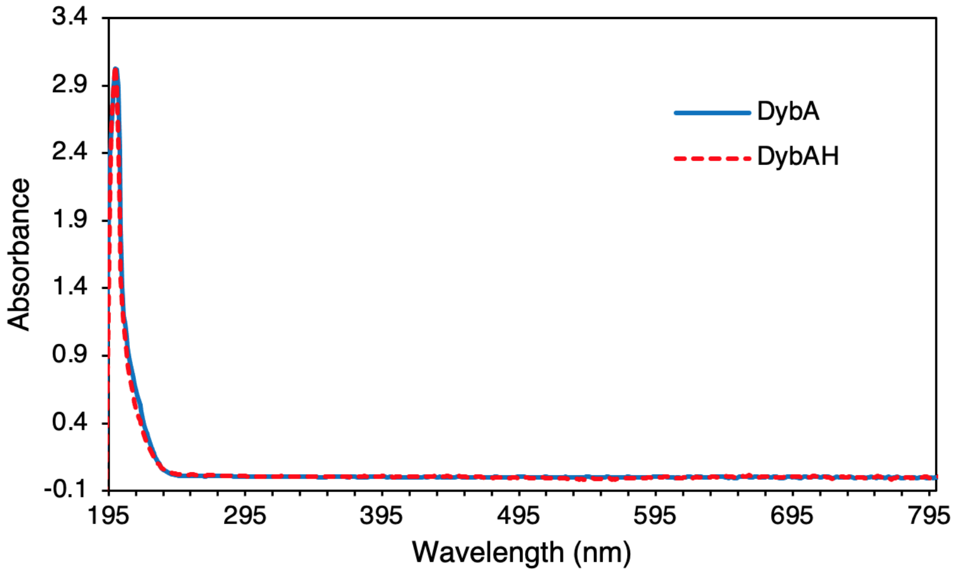

**Fig. S5. UV-Vis characterization of DybA and DybAH peptides.** UV-Vis spectra of unmodified DybA, and DybAH (DybA modified by DybH) at 100 μM in $H_2O$, showing no change in absorbance, suggesting the lack of an aromatic feature associated with the DybH modification. DybA does not contain any Trp and Tyr residues, hence the absence of a 280 nm peak.

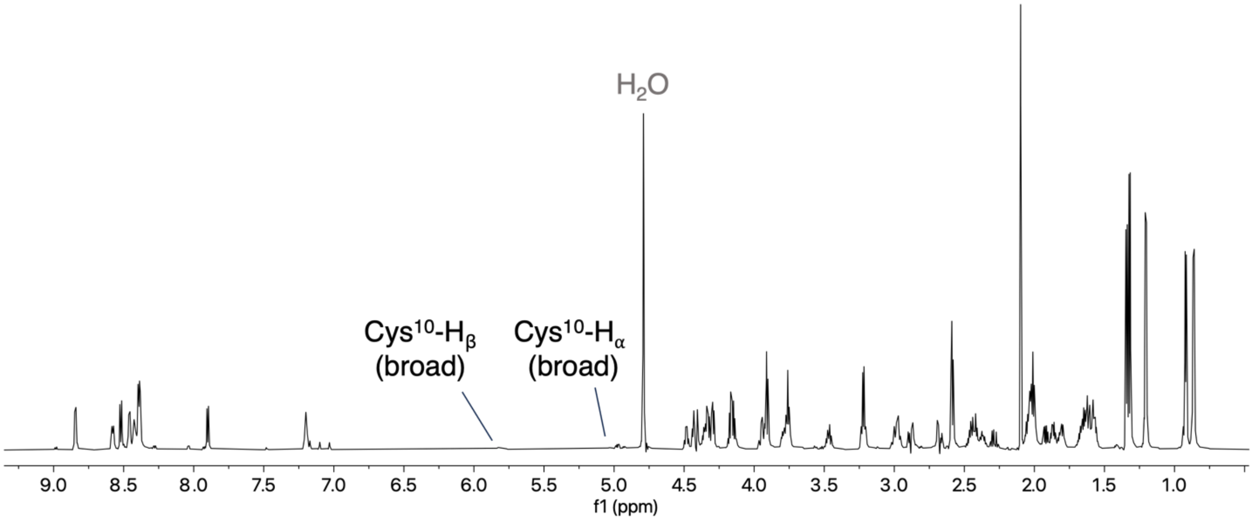

**Fig. S6. $^1$H NMR spectrum of LysC-digested DybAH in 90% $H_2O$:10% $D_2O$.** Data collected at 25 °C on an Agilent VNMRS 750-MHz NMR spectrometer equipped with a Varian 5 mm PFG triple resonance probe, with water suppression.

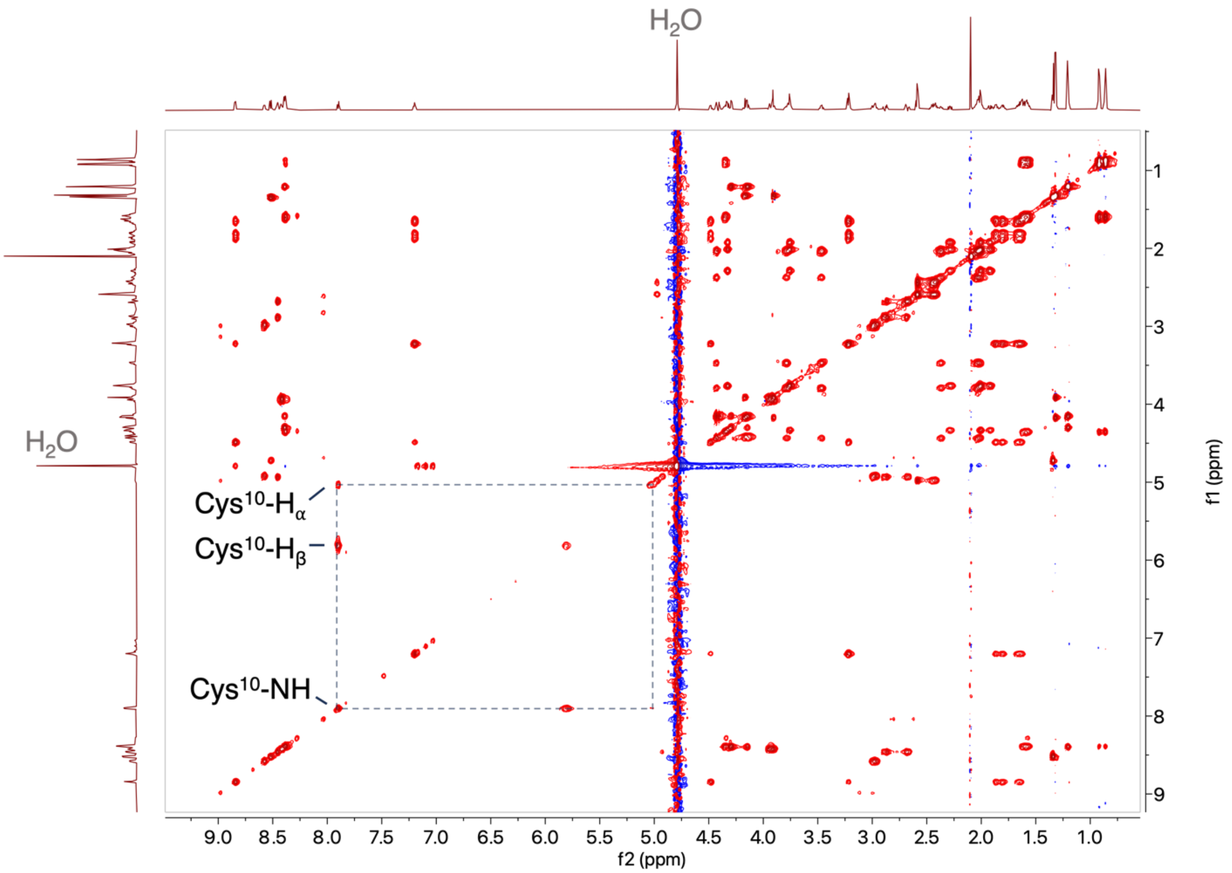

**Fig. S7. 2D $^{1}H$-$^{1}H$ TOCSY spectrum of LysC-digested DybAH in 90% $H_2O$:10% $D_2O$.** Data collected at 21 °C on a Bruker 600-MHz spectrometer equipped with a 5 mm CryoProbe Prodigy BBO. The signals of the Hα and Hβ of the former Cys10 that are shifted downfield compared to a normal Cys residue are indicated.

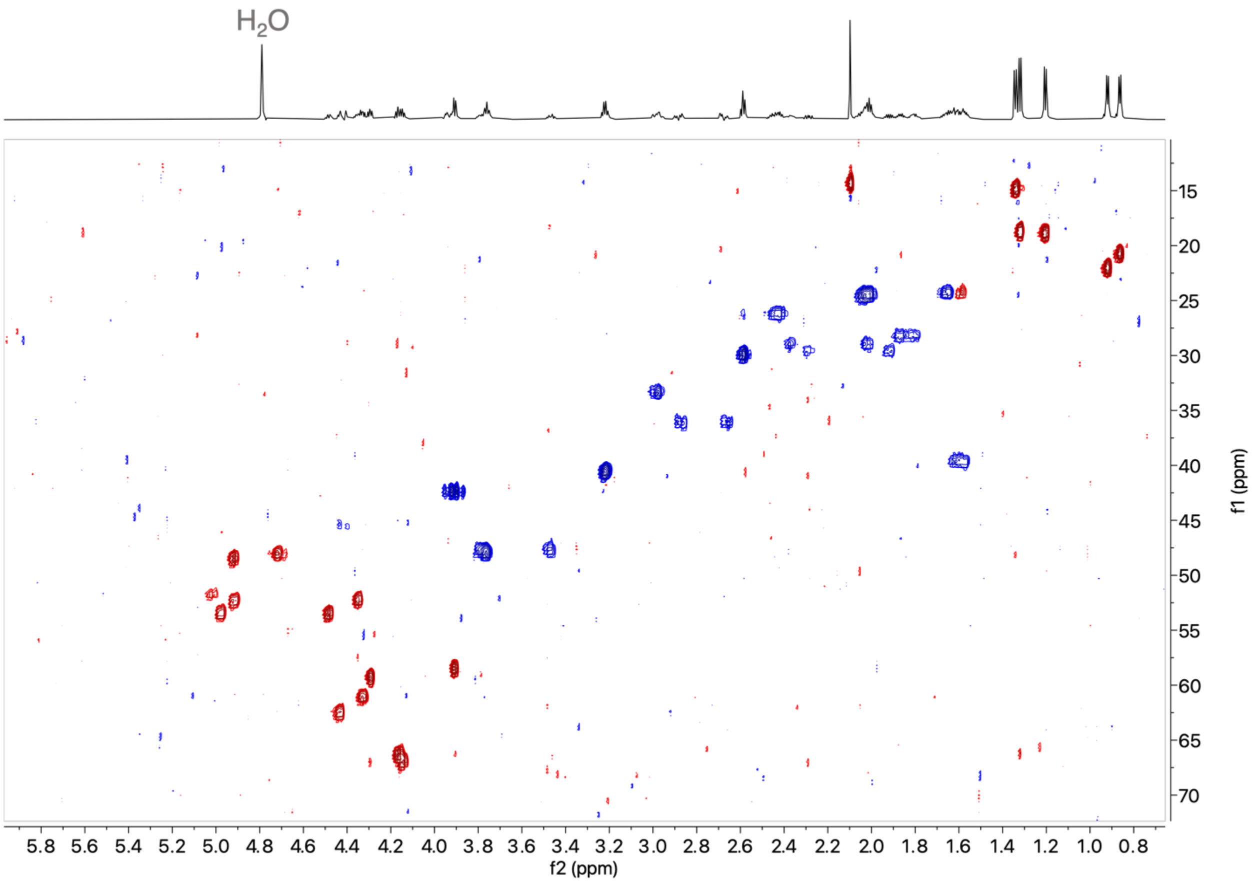

**Fig. S8. 2D $^1H$-$^{13}C$ HSQC spectrum of LysC-digested DybAH in $D_2O$.** Data collected at 21 °C on a Bruker 600-MHz spectrometer equipped with a 5 mm CryoProbe Prodigy BBO.

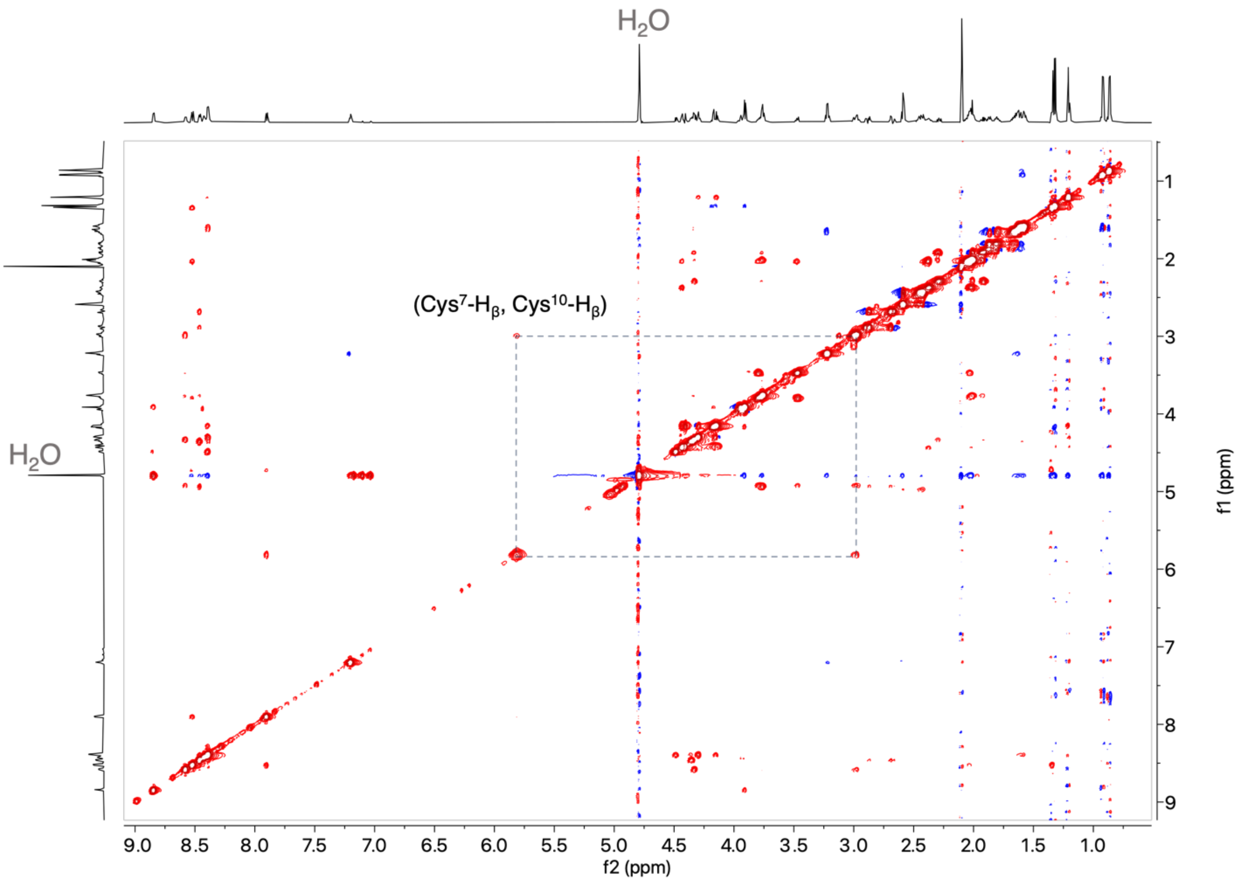

**Fig. S9. 2D $^1$H-$^1$H NOESY spectrum of LysC-digested DybAH in in 90% $H_2O$:10% $D_2O$.** Data collected at 25 °C on an Agilent VNMRS 750-MHz NMR spectrometer equipped with a Varian 5 mm PFG triple resonance probe, with water suppression. The NOE between the Hβ of the former Cys10 (5.81 ppm) and the Hβ of Cys7 (2.98 ppm) is indicated.

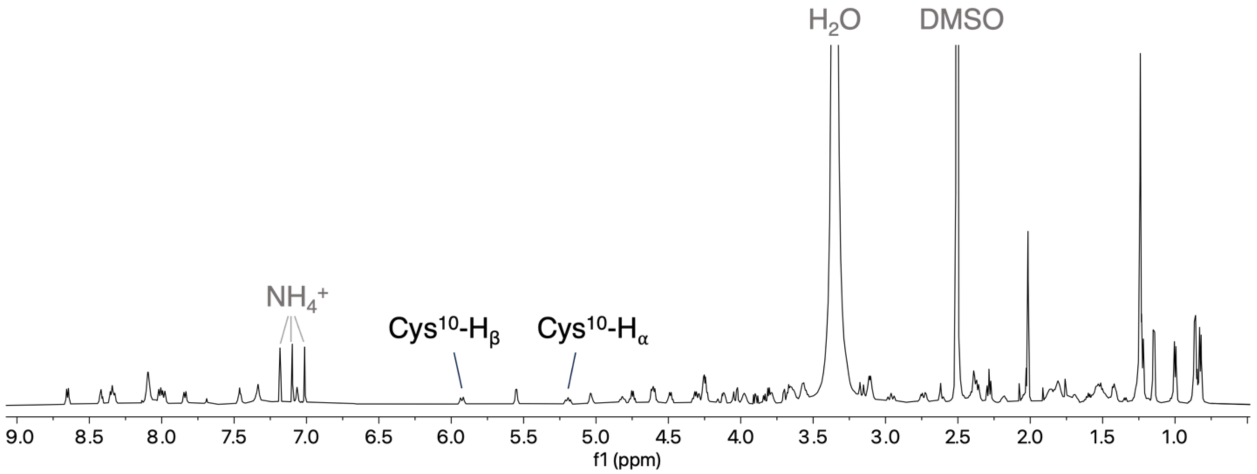

**Fig. S10. $^1$H NMR spectrum of IAA-alkylated, LysC-digested DybAH in DMSO-d6.** Data collected at 21 °C on a Bruker 600-MHz spectrometer equipped with a 5 mm CryoProbe Prodigy BBO.

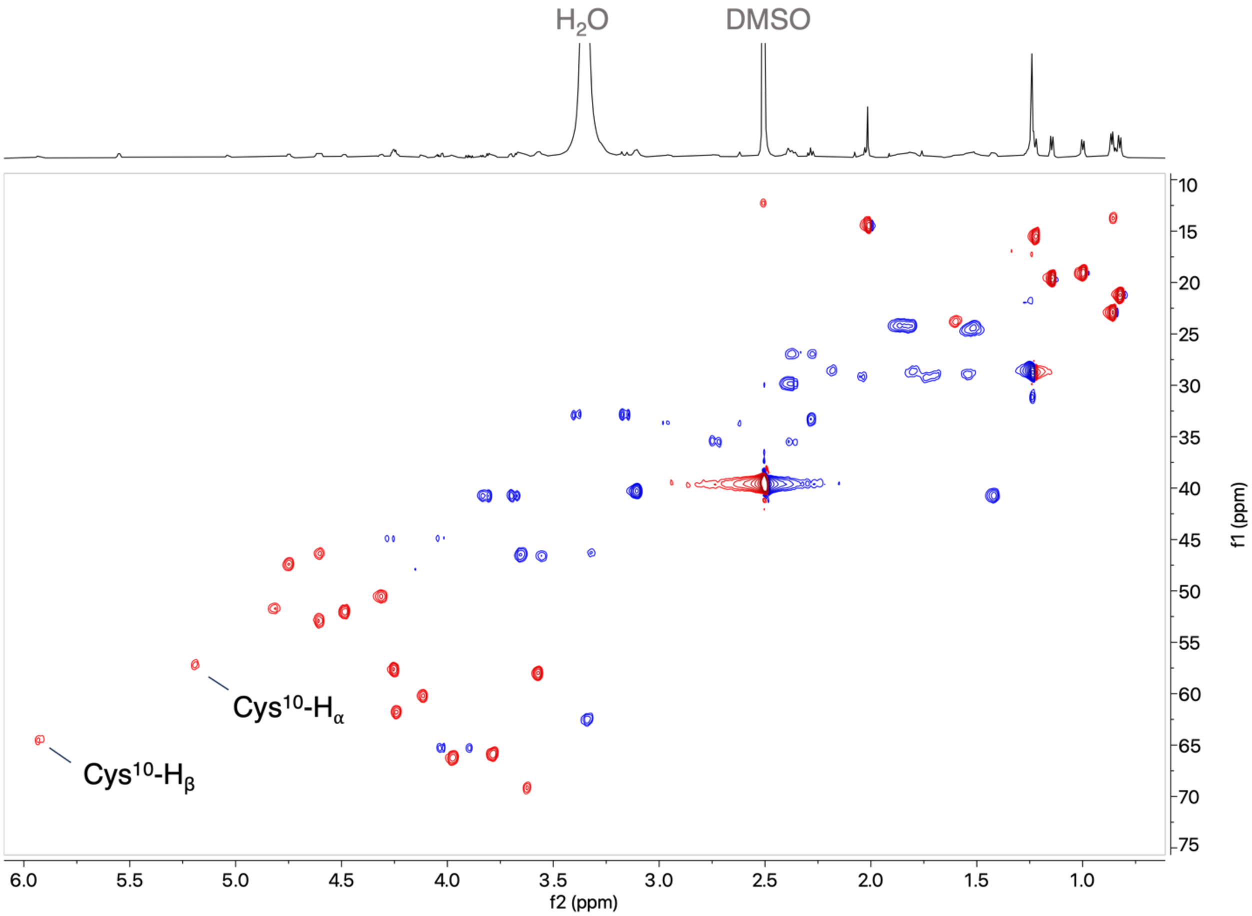

**Fig. S11. 2D $^1H$-$^{13}C$ HSQC spectrum of IAA-alkylated, LysC-digested DybAH in DMSO-d6.** Data collected at 21 °C on a Bruker 600-MHz spectrometer equipped with a 5 mm CryoProbe Prodigy BBO. Signals in red come from CH/$CH_3$ protons and signals in blue come from $CH_2$ groups.

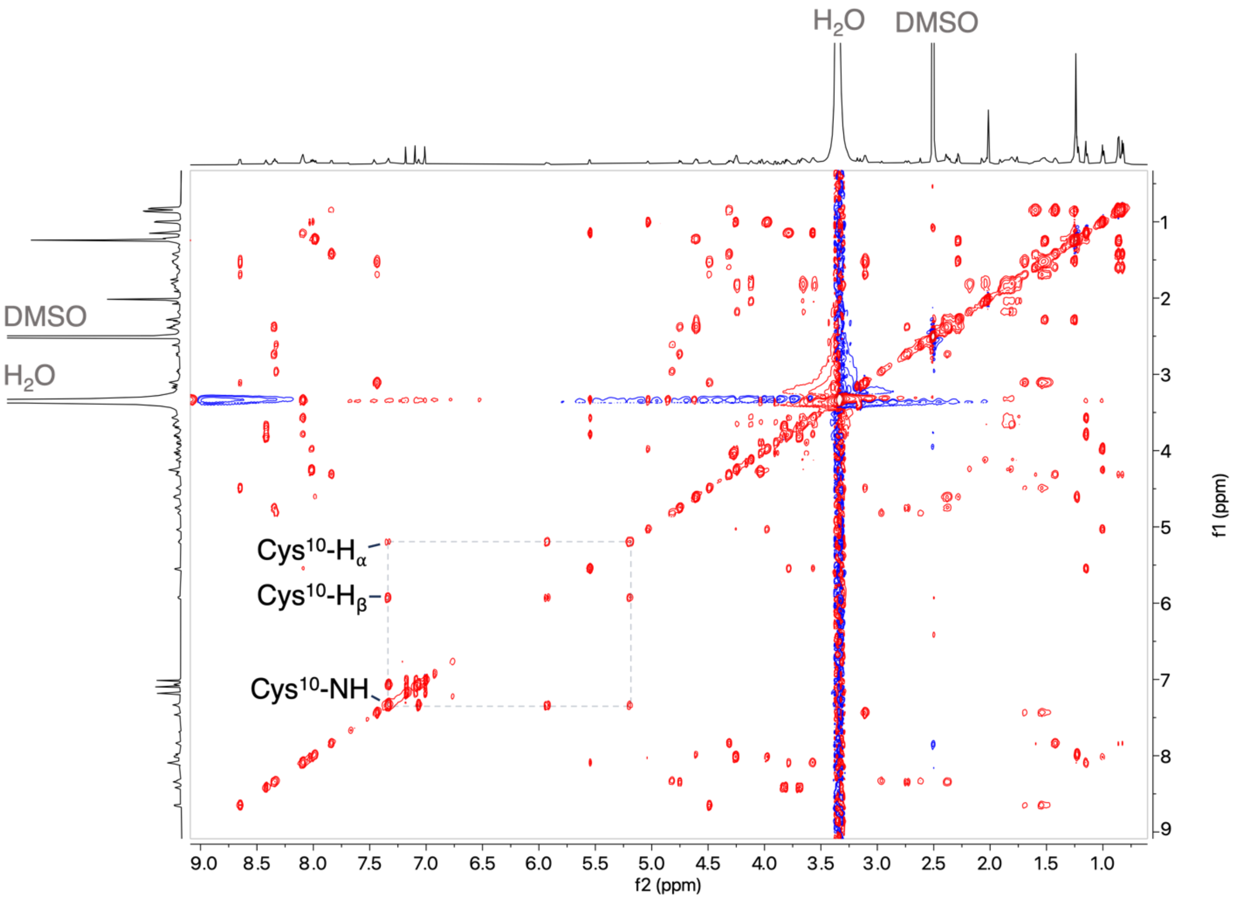

**Fig. S12. 2D $^1H$-$^1H$ TOCSY spectrum of IAA-alkylated, LysC-digested DybAH in DMSO-d6.** Data collected at 21 °C on a Bruker 600-MHz spectrometer equipped with a 5 mm CryoProbe Prodigy BBO.

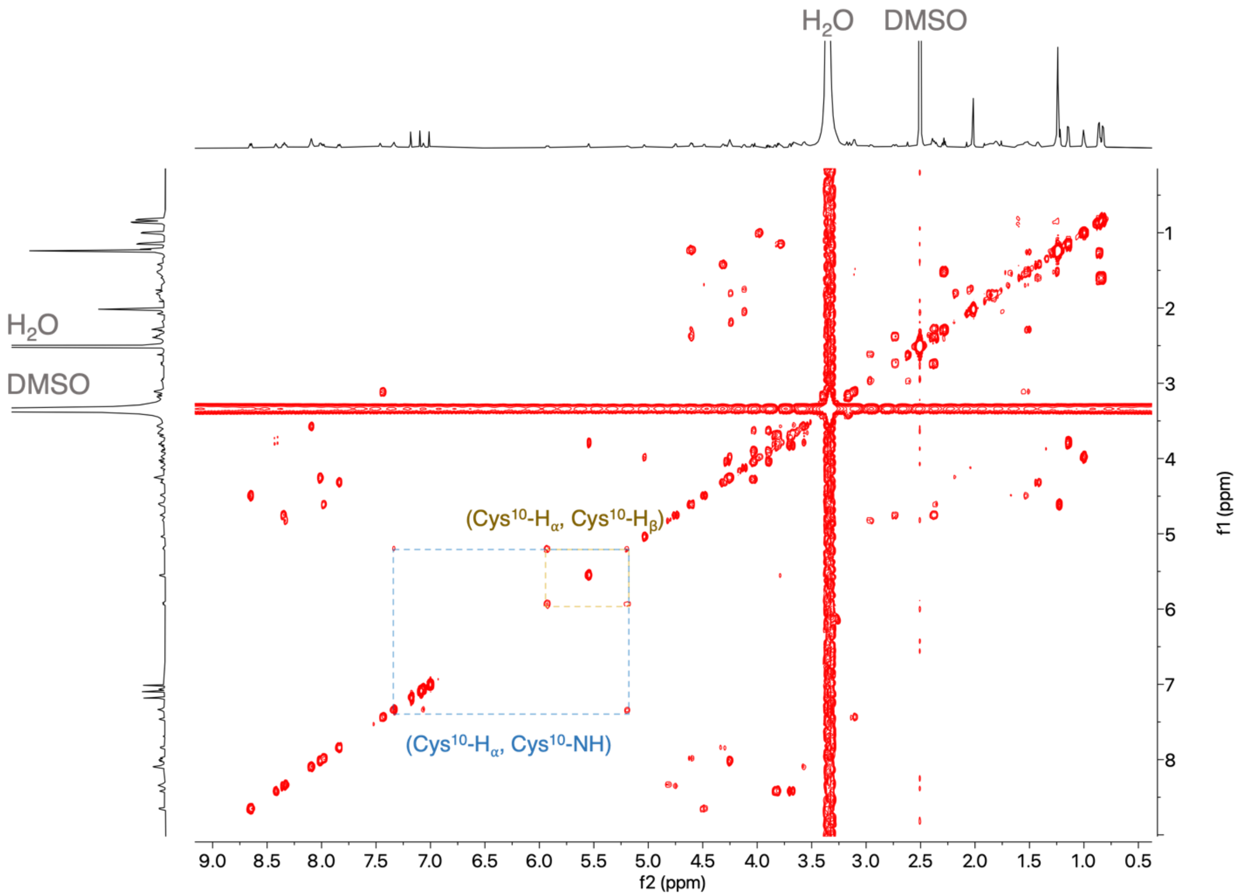

**Fig. S13. 2D $^1H$-$^1H$ COSY spectrum of IAA-alkylated, LysC-digested DybAH in DMSO-d6.** Cross peaks observed between the Cys10-NH with the Cys10-Cα support the assignment of the α- and β-carbons. Data collected at 21 °C on a Bruker 600-MHz spectrometer equipped with a 5 mm CryoProbe Prodigy BBO.

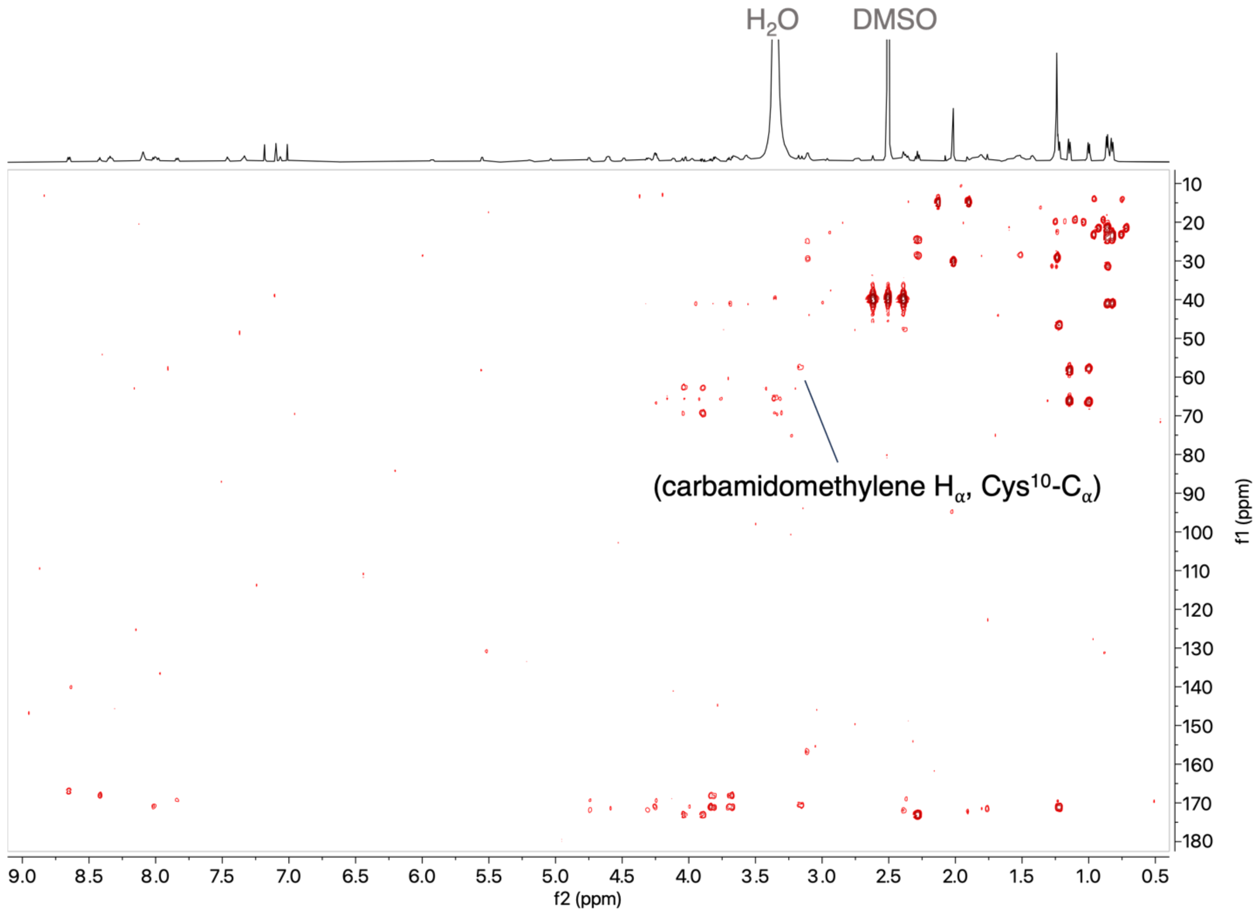

**Fig. S14. 2D $^{1}H$-$^{13}C$ HMBC spectrum of IAA-alkylated, LysC-digested DybAH in DMSO-d6.** Data collected at 21 °C on a Bruker 600-MHz spectrometer equipped with a 5 mm CryoProbe Prodigy BBO. The cross-peak between the carbamidomethylene protons originating from IAA and the Cα of the former Cys10 is indicated.

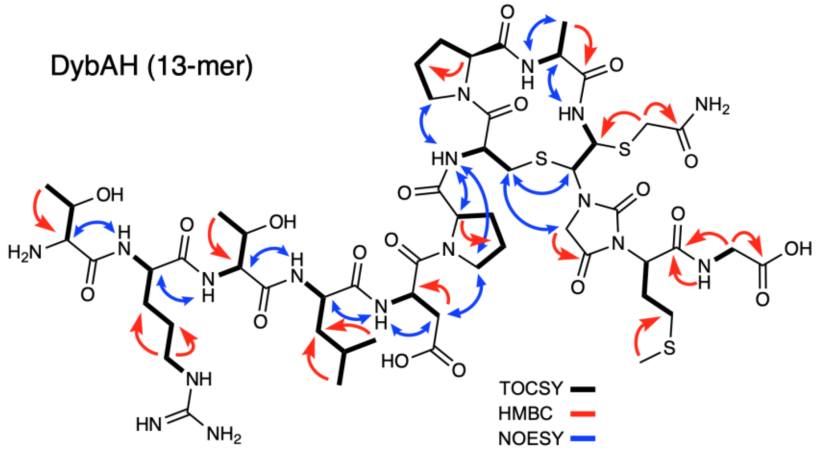

**Fig. S15. Full structure of IAA-alkylated, LysC-digested DybAH with NMR assignments.** Full chemical shift assignments can be found in Tables S4-S5.

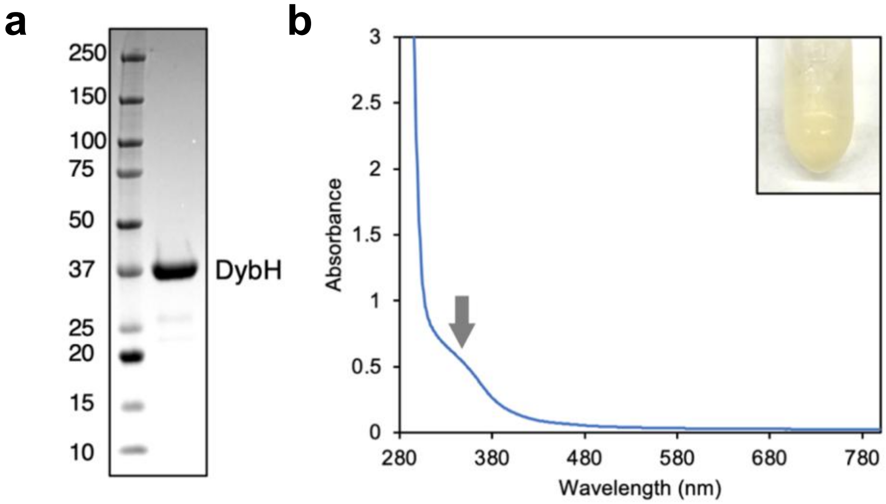

**Fig. S16. Heterologous expression and UV-Vis characterization of DybH.** (a) His-TEV-DybH following IMAC purification and desalting. The molecular masses of the proteins in the MW ladder (Bio-Rad Precision Plus Protein™ Kaleidoscope™ Prestained Protein Standard) are shown in kDa. (b) UV-Vis spectrum of aerobically isolated DybH, with the arrow showing an absorbance feature at approximately 350 nm. Inset: DybH enzyme at 100 μM concentration.

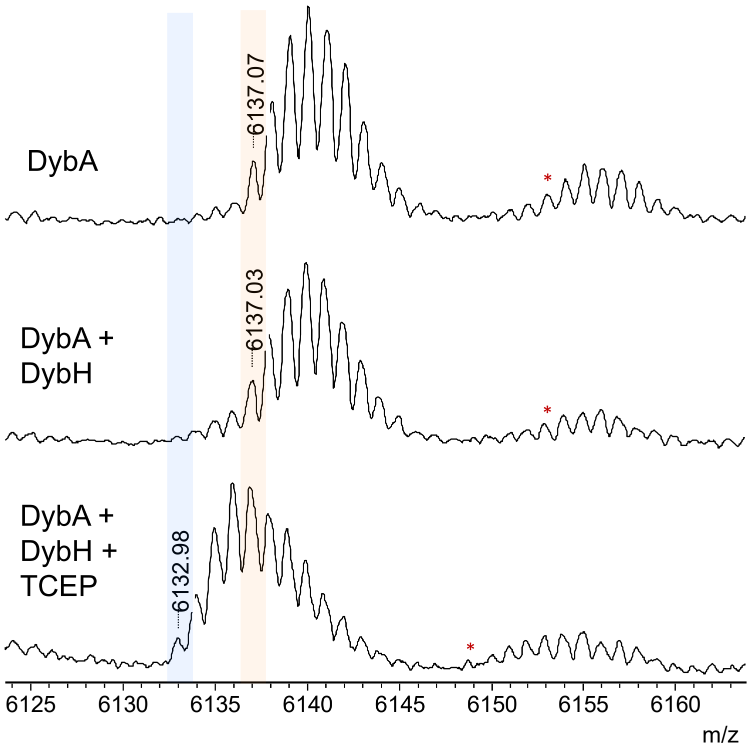

**Fig. S17. In vitro activity assay of DybH.** MALDI-TOF MS spectra of DybA alone, DybA with DybH, and DybA with DybH and TCEP. The assay concentrations were as follows: DybA (75 μM), DybH (15 μM), and TCEP (1 mM). TCEP is needed to keep the DybA thiols reduced. The reaction buffer was 25 mM HEPES, 200 mM NaCl, pH 7.6, and the reaction was performed at room temperature for 1 h before quenching the reaction to a final concentration of 10% acetonitrile and 0.2% trifluoroacetic acid. The samples were desalted with a C18 Ziptip prior to mass spectrometry analyses. Red asterisks correspond to DybA peptides with an oxidized methionine residue.

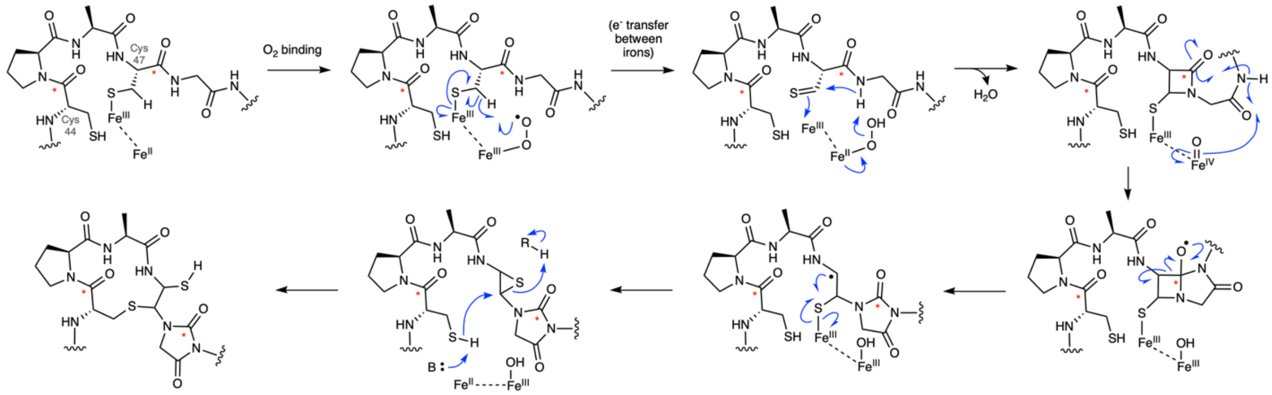

**Fig. S18. Proposed mechanism for DybH catalysis.** Following substrate binding of DybA, the reaction is proposed to be initiated by oxygen activation, and H-atom abstraction at the Cβ of Cys44, followed by a series of rearrangements. The mechanism is similar to that proposed for ChrH, with the exception of the final step, in which the ring opening of the episulfide results in addition of a proton instead of a methyl group from SAM (as in ChrH). Red asterisks show the fate of specific atoms based on $^{13}C$ labelling studies with ChrA (13).

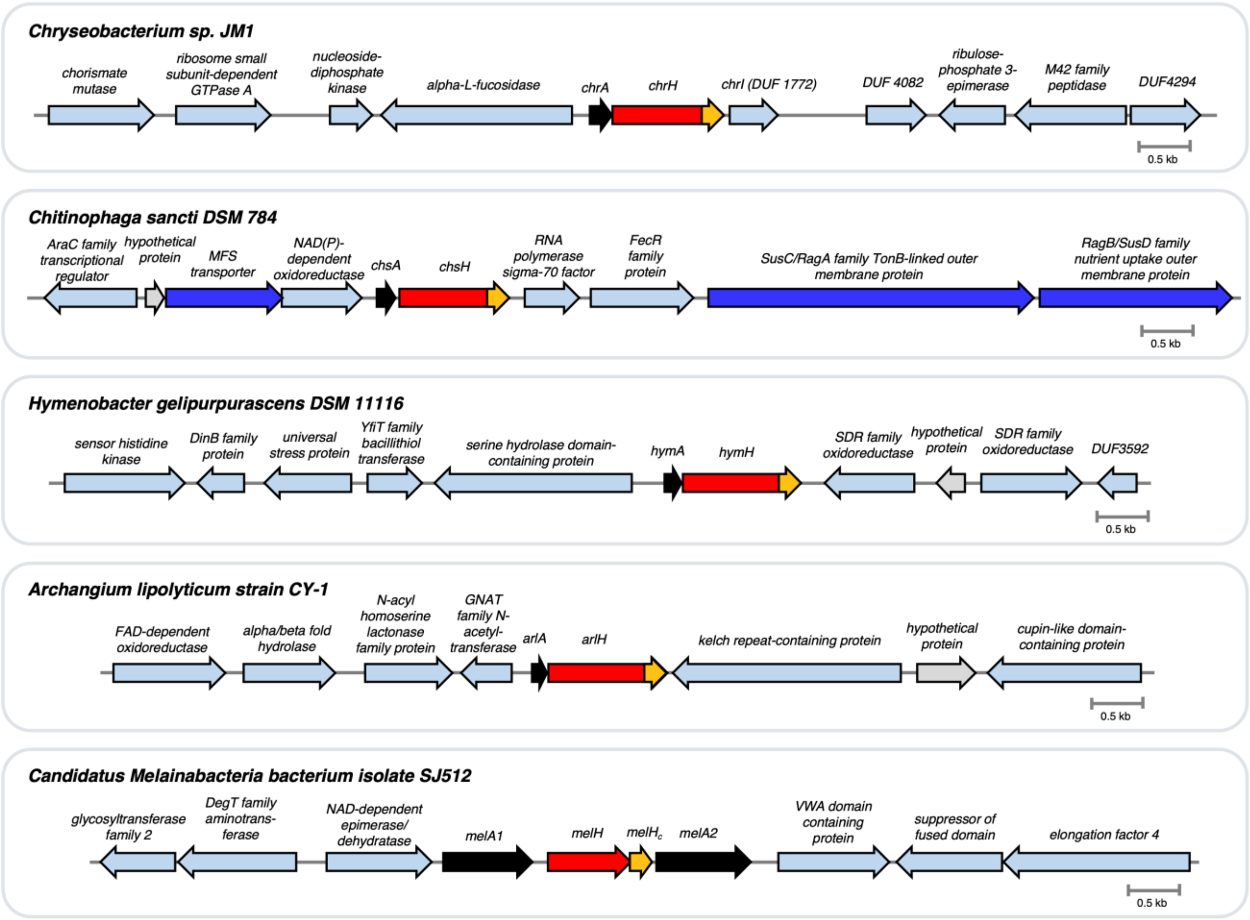

**Precursor sequences:**

**ChrA**: MKIPALVMASLLAVSVSGQTTKPVKKGTKSVKKVKKMDSPKTVKAEPTKVVKRDTILKHGGGCPACGMG
**ChsA**: MKLSKSLLSAIMIGIAVQTTVVSCSKDEQVKPKKADQANKQSESKPVDNPDSCPACGMG
**HymA**: MKLSQAILGAVLVGLTAQTTGCIKKSDPTPKEEQGKSGKKSPEIPYNCPGCGLG
**ArlA**: MNQNLREKATQLLDVASNTQGMKSSSPLPCPACGMMMALPESSPQLVTPATR
**MelA1**:
MKNKSAKVLTSLFALGLTTGAVPARADKAAVETAEASVKMFEQRSGTNNSSYTSLLMYLAGVYLANDREKDAEIIY
NKAIFNLKSKPDRKAEVPALMLNWAMLLASNKGASKEKAEKALSDGLKLANDLPAASKERINYLIGTINFYNVIGKPA
EKQARIKAADEHLATLEKNEKLNNEEITNVAANLVKLAEIQTFPMPVMRLRYQPPVFQVVPDNSPDKPNTVRAKEFK
SAEAFQLRAIKQYDRLPETVPWRIEAHRKLILWYRSLGQTKQEEFQIQQLGKIMHTTDRDKLFPQPAPCPACGMG

- Precursor
- MNIO
- ChrH-like C-term (MelH$_c$)
- Transport related
- Other protein function
- Hypothetical protein

**Fig. S19. BGCs and sequences of ChrA, ChsA, HymA, ArlA, MelA1.** Sequences of the precursor proteins/peptides are shown below.

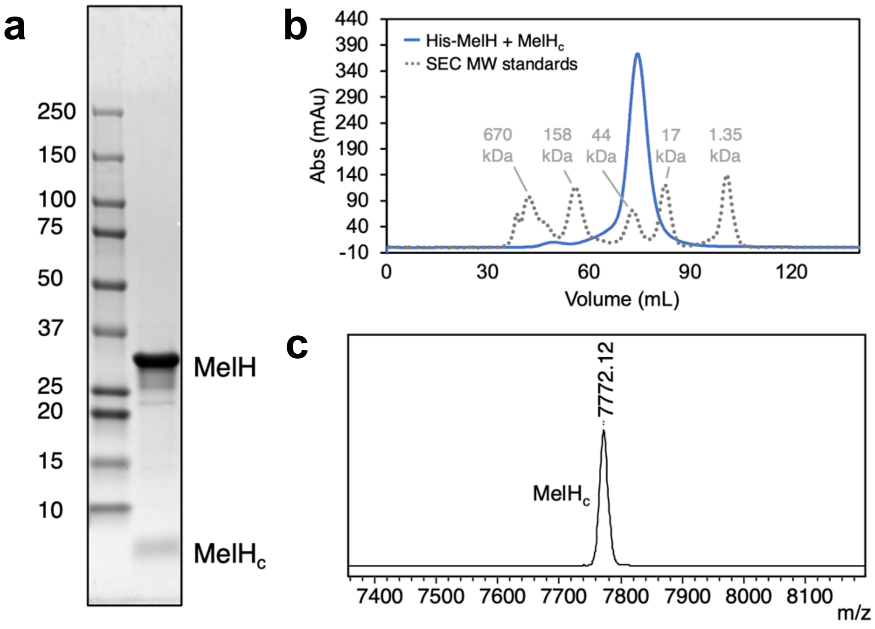

**Fig. S20. MelH and MelH$_c$ form a stable complex.** (a) SDS-PAGE analysis of the IMAC elution fraction of His$_6$-TEV-MelH coexpressed with untagged MelH$_c$. (b) Size exclusion chromatography of His$_6$-MelH with MelH$_c$ (solid line) and Bio-Rad SEC MW standards (dotted line). (c) MALDI-TOF MS spectrum (LP mode) of post-SEC purified MelH, showing that the additional gel band below 10 kDa corresponds to MelH$_c$ (7761.60 Da expected).

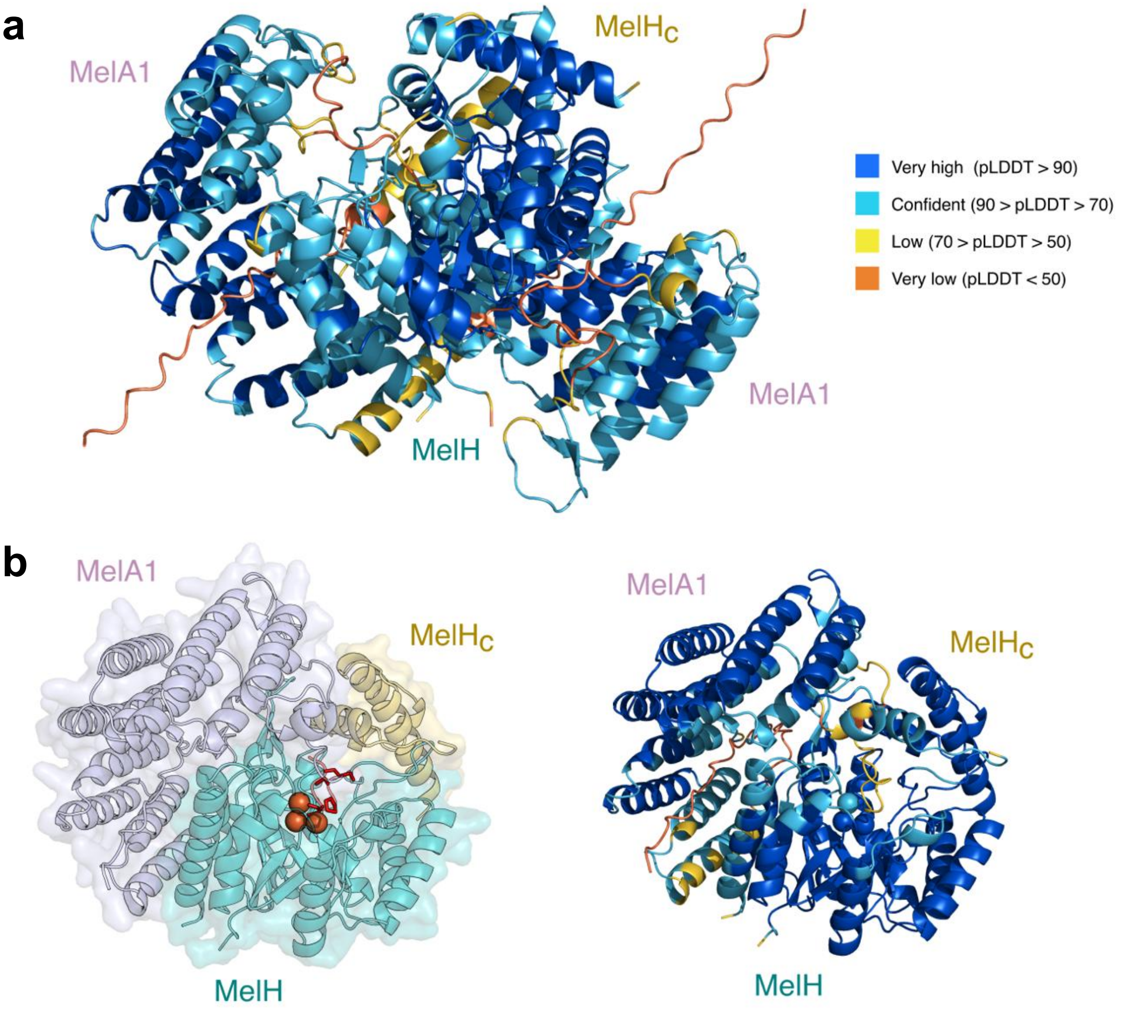

**Fig. S21. Structural prediction of the MelA1, MelH and $MelH_c$ complex.** (a) Alphafold3 predicted model of the MelA1 dimer complexed with MelH and $MelH_c$ in Fig. 3d of the main text, colored by predicted local distance difference test (pLDDT) values. Three $Fe^{3+}$ ions were modelled for MelH. (b) Alphafold3 predicted model of the MelA1 monomer complexed with MelH and $MelH_c$. Right: the structure colored by pLDDT values.

**a**

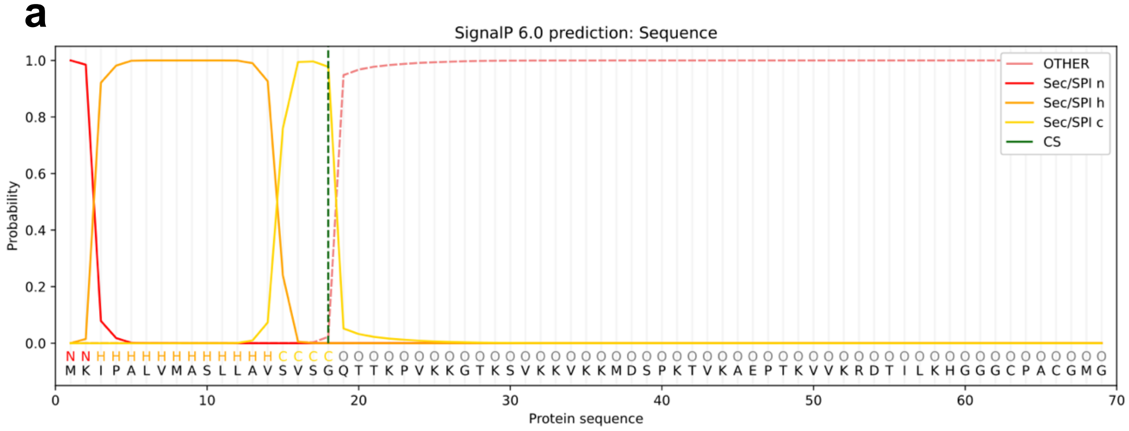

**b**

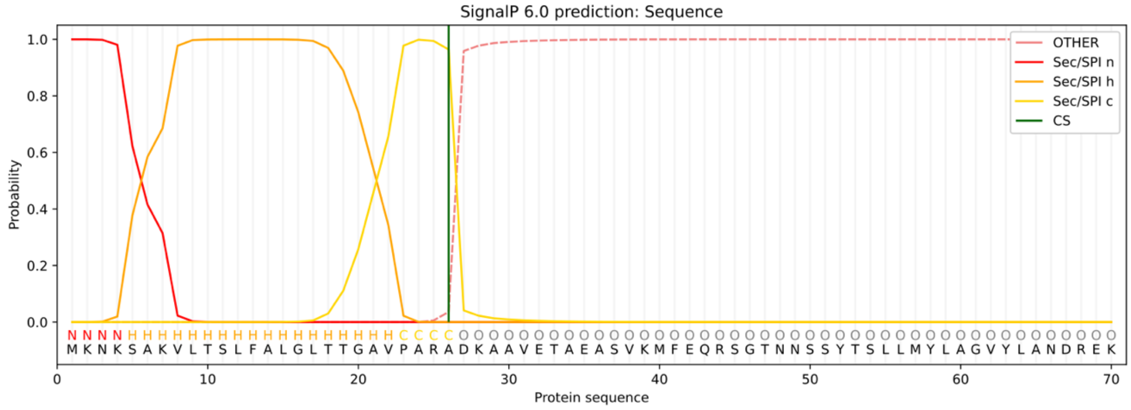

**Fig. S22. SignalP analysis of secreted peptides in the ChrA-like family.** SignalP 6.0 (5) graphs showing predicted signal peptidase I signal peptides for (a) ChrA and (b) MelA1. Only the first 70 amino acids are shown for MelA1.

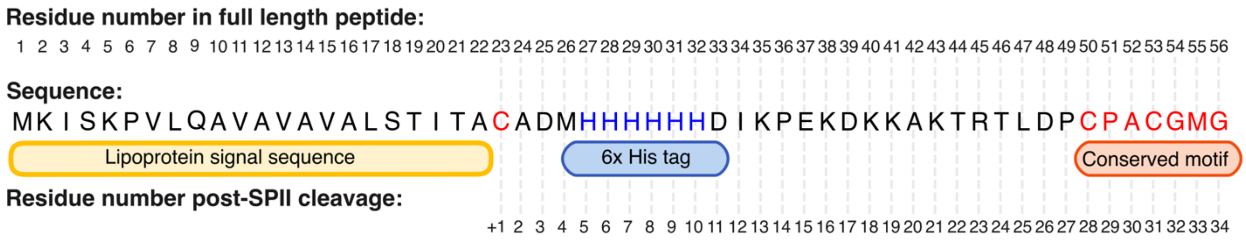

**Fig. S23. Sequence of DybAH construct with a 6xHis tag at residue 27.** The predicted lipoprotein signal sequence is shown in addition to the conserved CPACGMG motif. Following SPII cleavage, the His tag starts at the +5 position.

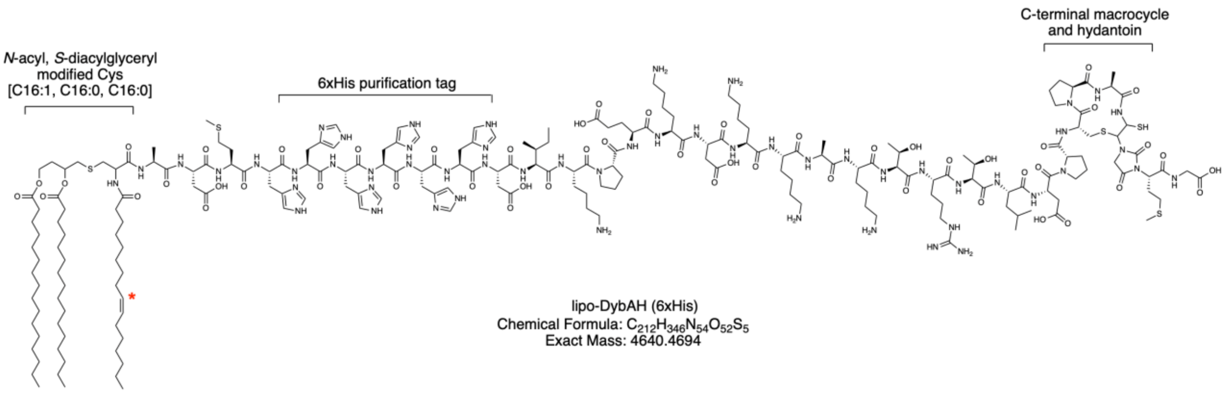

**Fig. S24. Full structure of lipo-DybAH with embedded 6xHis.** The 6xHis tag was inserted within the full length DybA sequence, corresponding to the +5 position after lipid modification (Cys is +1). The red asterisk indicates that the position and stereochemistry of the unsaturated C16:1 group was not experimentally determined in this study. However, based on prior work, the Δ9 isomer is the predominant positional isomer of palmitoleic acid phosphatidyl-ethanolamine (PE) in *E. coli*, which is the substrate for Lnt (apolipoprotein N-acyltransferase) (21, 22).

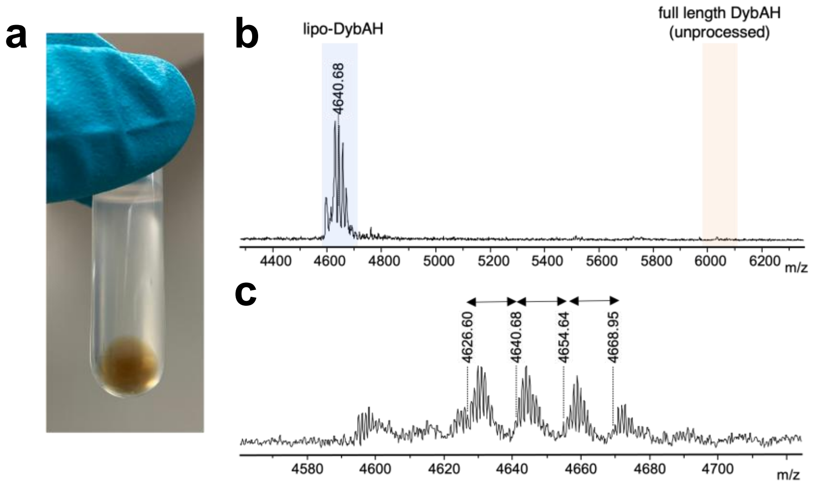

**Fig. S25. Heterologously expressed lipo-DybAH isolated from *E. coli* OMVs.** (a) Outer membrane vesicles (OMVs) were isolated from the spent media of *E. coli* cells that heterologously expressed lipo-DybAH. (b) MALDI-TOF MS of lipidated DybAH purified from *E. coli* OMVs by IMAC. The insoluble fraction of the eluate was analyzed by MS. (c) Magnified MALDI-TOF spectrum showing the lipo-DybA region containing at least four differently lipidated molecules.

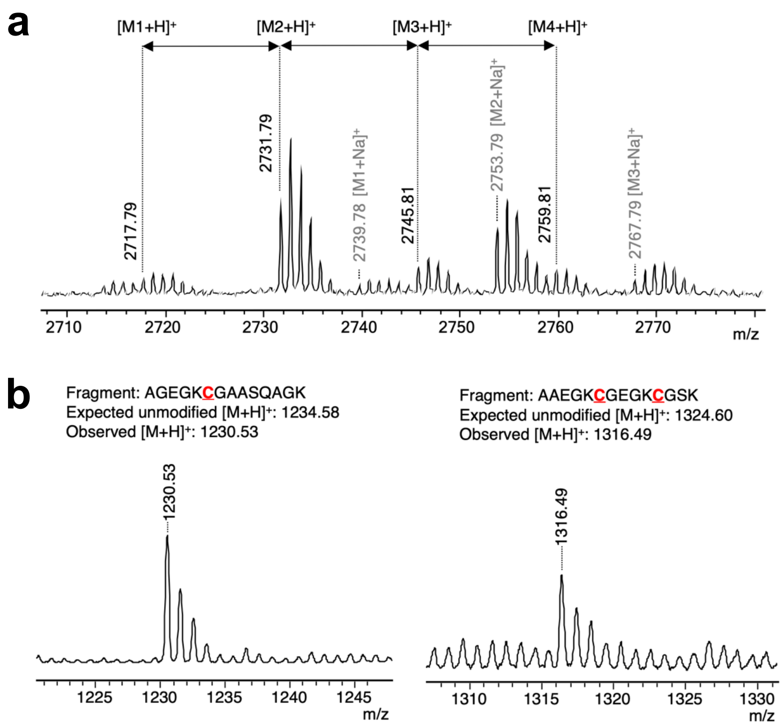

**Fig. S26. MALDI-TOF MS analysis of lipo-NedABC isolated from the insoluble membrane fraction.** (a) Isolated lipo-NedA post-LysC digest showing varying lipid lengths separated by 14 Da ($CH_2$) with the major lipidated species containing C16:1, C16:0, C16:0 acyl chains. (b) LysC-digest of the lipoprotein fraction showing that lipidated NedA is modified by NedBC, resulting in a −4 Da modification at each of the three conserved EGKCG motifs in the NedA peptide.

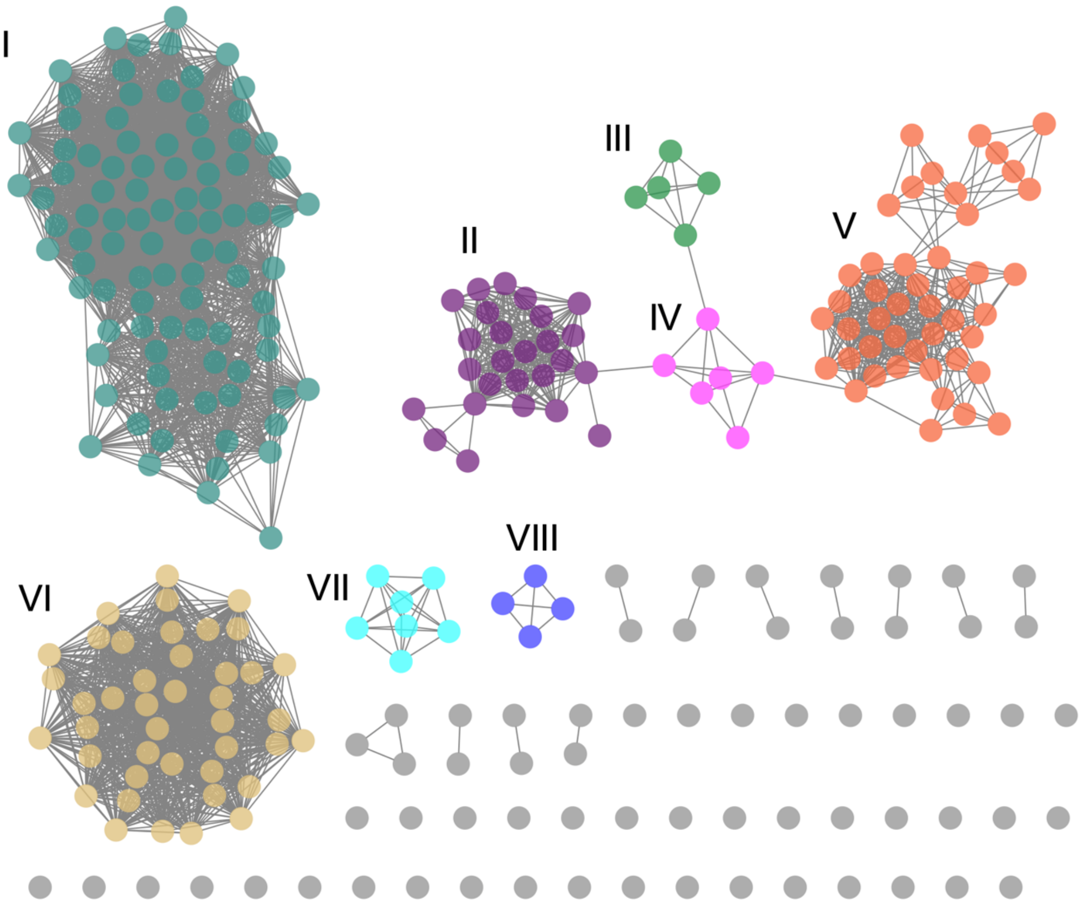

| Cluster | Annotation | Representative sequence |
|---|---|---|
| I | AIP | MKKALAIALSAIGAVLGTVATTGCVMVLFDEPDMPKSLID |
| II | AIP | MKNLMKIARKTSSVLAAMALVVATSSVASCCYHWFAQPVEPEELRKFVGEN |
| III | AIP | MKYRILNSMVKVLTLVAFVCAASPSHSNCYEPEKPASLR |
| IV | AIP | MKKKILMGIAAVATVFASVISTSACYFSFYQPEEPECLREE |
| V | AIP | MKNCKQYFLKTFAFLLSFLAISSVNSACAVLFGQEKEPDSLARYKKC |
| VI | LptM chaperone | MKKVLSILLLGAFCTLATGCGVKGPLYFPEKEQPQNTQKTQ |
| VII | TolC-subunit | MNNMNTSKIIACGATLLLSSCGIYTSYEPQTSVPENLYGEEVTEAVSK |
| VIII | AIP | MKKIMYQVCTKLAACAFLAAIVSVGTASCNGLYQPKMPDKLVK |

**Fig. S27. SSN of predicted auto-inducing peptides (AIPs) with SPII signaling peptides.** Representative sequences from the top 8 clusters are shown below the SSN, with cysteines in red. Clusters VI/VII correspond to lipoproteins predicted not to be AIPs. The full list of sequences in the SSN can be found in Table S7. Initial peptide sequence list of predicted AIPs obtained from Zhang *et al.* (2025) (23). The SSN was generated by EFI-EST (7). The Cytoscape file is provided as Dataset 3.

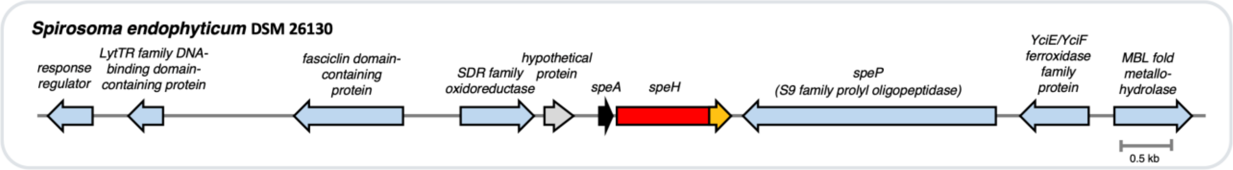

**SpeA**: MKLSKALLQTIAVAVAVTTLSSCDKGKVIDPKKDDTKQQIPYDCPGCGLG

**SpeP**:
MLHNLTTIRQSRWILLISWLAFSVASNAQDVPTYQTPPKALADLVTMPPTPTVRMSSKGDMMLIIEQPTVT
TIDELSQPELKLAGLRLNPANNGPSRVFYITGLKLKNLTDKEAKPIAGLPAKPQLSFIQWSPDQTKVAFAN
STDTQIEFYIVEVATATAKKVGSLALNATLGAPFQWLPDSKALIVKAVPAGRGPAPQLSRVPAGPTTQEN
VKGSRGQAPTFQDLLKNPVDEQQFAYYTTAQVVRMGLDGVATNIGPAGIILSAEPSPDGQYVMIETVHT
PFSYLVPVRRFPIKTEVYAVGGSLVKTLNDSPLQESVPYSRDGVATGPRDFNWRADTPASIYYTVAQDN
GDPKTKADIRDKVFLIDAPFSAQPKEIYAAQFRFETFDWGNESMALATERWWQNRKTITKTVNPTNWQT
ATLFDRSYEDRYTNPGQPDTKHNQYGREVLNLLPNNEILMVNGQGSSPEGDRPFVSLLNLKTKQTREL
WRSTAPLFERPIAVLDAAKQVILTTQETTEESPNYFVRNLKARIAPVQVTFFPHPYPQLKGIQKKQLRYKR
TDGVELTAMLYLPAGYKKEQGPLPTFLWAYPAEFKSKDAASQISGSPYQFGWNRPFWGAAAFVTMGYA
ILDNASIPIVGEGDKEPNDTYVEQLVSSAKAAIDEGVRLGVVDSSRVGVGGHSYGAFMTANLLTHSKLFK
GGIARSGAYNRTLTPFGFQNEQRSYWQAPDVYNNMSPFMNADKMKTPLLLVHGEADNNTGTFPIQSER
YYNALKGFGATTRLVFMPYESHHYDAKESLLHMLWEMNGWLDKYVKNPTAAGQANKAGKLGSGK

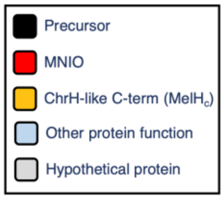

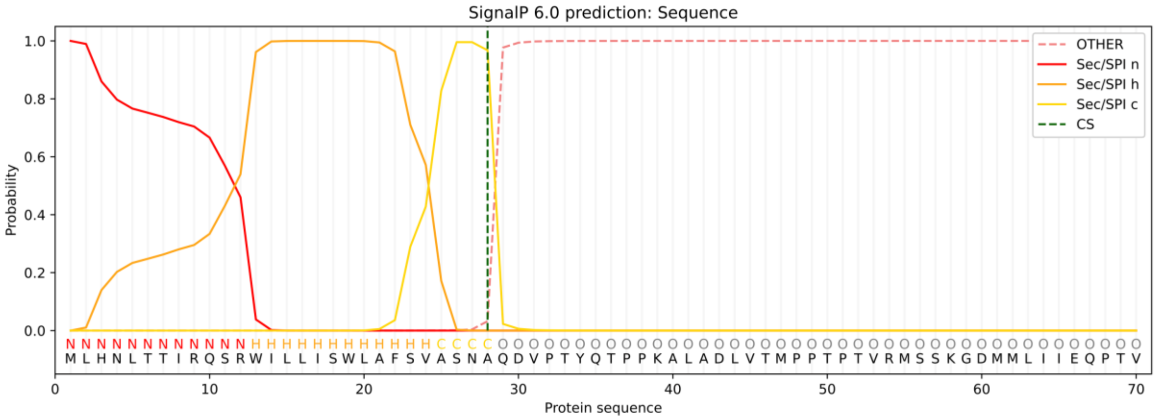

**Fig. S28.** ***S. endophyticum*** **BGC contains a secreted prolyl oligopeptidase.** (a) BGC of *S. endophyticum.* SpeA contains a SPII signal peptide for lipidation. SpeH is the ChrH/DybH-type MNIO and SpeP is a S9 family prolyl oligopeptidase. (b) SignalP 6.0 analysis of SpeP shows that the first 28 amino acids constitute a SPI secretion signal.

Ring B (in nisin, mutacin 1140, etc.) | Macrocycle in HymAH | Macrocycle in DybAH

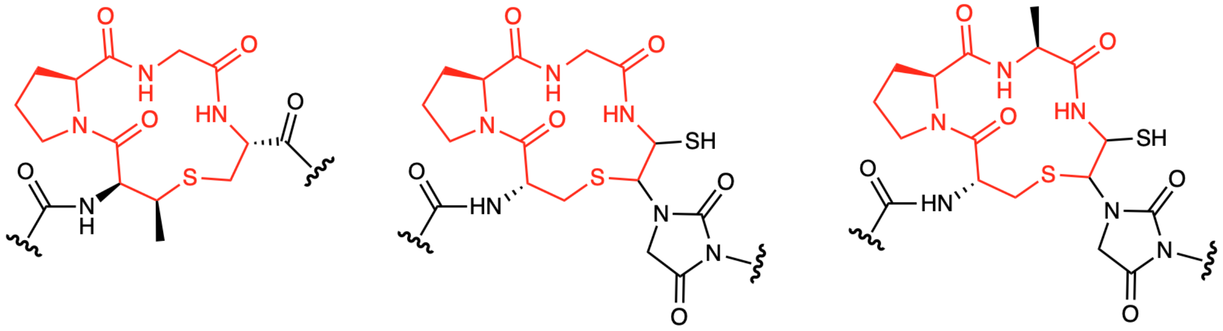

**Fig. S29. Structural comparison of the chryseobasin-type macrocycle to the B ring of nisin and mutacin 1140.** Structural similarities between the modified HymA and DybA macrocycle and the lantibiotics nisin and mutacin 1140 are shown in red bonds.

**Table S1. Co-occurring protein families with ChrH-like MNIOs.** The EFI-Genome Neighborhood Network Tool (GNN) was used to analyze frequency of co-occurrence (10 genes upstream and downstream) of the ChrH-like MNIOs. The top 10 co-occurring PFAM domains with corresponding SwissProt annotations are shown. Frequencies greater than 1 indicate multiple copies of the same PFAM within the gene neighborhood window.

| **ChrH-type BGCs** | | |
|---|---|---|
| **Frequency** | **PFAM** | **Description** |
| 1.52 | PF00254 | FKBP-type peptidyl-prolyl cis-trans isomerase |
| 0.95 | PF00834 | Ribulose-phosphate 3 epimerase family |
| 0.95 | PF05343 | M42 glutamyl aminopeptidase |
| 0.93 | PF00440 | Bacterial regulatory proteins, tetR family |
| 0.93 | PF00293 | NUDIX domain |
| 0.93 | PF00334 | Nucleoside diphosphate kinase |
| 0.93 | PF14127 | Domain of unknown function (DUF) 4294 |
| 0.92 | PF00793 | DAHP synthetase I family |
| 0.92 | PF01063 | Amino-transferase class IV |
| 0.92 | PF01817 | Chorismate mutase type II |
| **DybH-type BGCs** | | |
| **Frequency** | **PFAM** | **Description** |
| 0.41 | PF07715 | TonB-dependent Receptor Plug Domain |
| 0.38 | PF13715 | CarboxypepD_reg-like domain |
| 0.3 | PF00072 | Response regulator receiver domain |
| 0.26 | PF00593 | TonB dependent receptor-like, beta-barrel |
| 0.2 | PF08281 | Sigma-70, region 4 |
| 0.2 | PF12833 | Helix-turn-helix domain |
| 0.19 | PF04542 | Sigma-70 region 2 |
| 0.18 | PF07980 | SusD family |
| 0.18 | PF14322 | Starch-binding associating with outer membrane |
| 0.16 | PF02518 | Histidine kinase-, DNA gyrase B-, and HSP90-like ATPase |
| **MelH-type BGCs** | | |
| **Frequency** | **PFAM** | **Description** |
| 0.42 | PF02574 | Homocysteine S-methyltransferase |
| 0.28 | PF01370 | NAD dependent epimerase/dehydratase family |
| 0.28 | PF00535 | Glycosyl transferase family 2 |
| 0.28 | PF02321 | Outer membrane efflux protein |
| 0.28 | PF01041 | DegT/DnrJ/EryC1/StrS aminotransferase family |
| 0.28 | PF02219 | Methylenetetrahydrofolate reductase |
| 0.28 | PF05076 | Suppressor of fused protein (SUFU) |
| 0.28 | PF00009 | Elongation factor Tu GTP binding domain |
| 0.28 | PF00679 | Elongation factor G C-terminus |
| 0.28 | PF03144 | Elongation factor Tu domain 2 |

**Table S2. Accession IDs (NCBI) and sequences of ChrH-subfamily of MNIOs used in the SSN in Fig. 1b.** Because sequences and accession IDs are often removed in recent consolidations of databases, the sequences are presented.

| NCBI ID | Organism | Sequence |
|---|---|---|
| ACU59980.1 | Chitinophaga pinensis DSM 2588 DSM 2588 | MEESRIEAIEWSFDALYKVKEVPDWFRELLTAFSDENRLIGHGVFFSLFSGKWLPEQENWLKHLSQTADAFQFDHITEHFGFMTGKDFHQGAPLNIPYTAITLNIGR<br>DRLKRIYEACKRPVGLENLAFSYSLDEVKRHGAFLEQLLEPVNGFIILDLHNLYCQLRNFDVTFEEMIALYPLDRVREIHISGGSWEDSVATPSRSIRRDTHDDAVPTE<br>VFQLLEMTIPRCPQLKYVVLEQLGNGLETETSRQYFYNDFLRMQEIIQQHNDQADDTSTDPFLPLIPFSTDAVVEDMMLYQQQLELSAIFESSASYGEAMQSLSTSS<br>LANSDWRIEQWEPHMIETAVKIAQKWMK |
| WP_034736621.1 | Chryseobacteriu m indologenes | MMRKPLLGLSMMPEAEFVSAALPLLQNNSVEVLEWSFDTLFNTNEPDWLRDLLNFYAENNRLIGHGVYYSLFDAQWTDRQEIWLSRLKDEVSKRNYNHITEHFGF<br>MNTENFHQGVPLPVPLHPITLQIGQDRLQRLQDAVNIPVGIENLAFSFCLDDVKEQGVFLEKLIDNSGGFLILDLHNMYCQSCNFDVDIQEIIKLYPLQKVKEIHLSGGS<br>WQESVYGKKMIRRDTHDHTIPEEIFAILPFVLSHCENLEYIIIERLGHTIKTKEEEDDFFSDFMRVKTIIDSSAGHERQGDLWIKKEFELSKYPVEDPVLYEEQSRLTKLL<br>FDGTHVDSIKNENFHYFNTAGWDPEMIFTAQEIIKKWNPY |
| WP_027375286.1 | Chryseobacteriu m | MRKPLLGLSMMPEAEFVSAALPLLQNNSVEVLEWSFDTLFNTNEPDWLRDLLNFYAENDRLIGHGVYYSLFDAQWTDRQEIWLSRLKDEVSKRNYNHITEHFGFM<br>NTENFHQGVPLPVPLHPITLQIGQDRLQRLQDAVNIPVGIENLAFSFCLDDVKEQGVFLEKLIDNSGGFLILDLHNMYCQSCNFDVDIQEIIKLYPLQKVKEIHLSGGS<br>WQESVYGKKMIRRDTHDHTIPEEIFAILPFVLSHCENLEYIIIERLGHTIKTKEEEDDFFSDFMRVKTMIDSSAGHERQGDLWIKKEFELSKYPVEDPVLYEEQSRLTKL<br>LFDGTHVDSIKTENFHYFNTAGWDPEMIFTAQEIIKKWNPY |
| WP_012927702.1 | Spirosoma linguale | MSRLYSSLACNLDTNSLQASLPLFEAEKVQAIEWSFDTLFALGEMPDWFAELLRAYSQEGRLIGHGVFFSLFSGKWTPDQQQWLKQLKALSADFTFDHLTEHVGF<br>MTGEDFHKGAPISIPFTTSTLAIGRDRLLRLQDAGNCPVGLENLAVAYSLDDVKRQGDFLAQLLEAVNGFILLDLHNLYCQSQNFDLGIADIQALYPLDRVREIHISGG<br>SWVPSTVNPTKQIRRDTHDESVPAAVFHALQQVIGQCPNLKYVVLEQLGTGLTTDVSRQHFREDFYTMDALIEATNQLNSHSPINSFLPLSETSIPETPMENPLLNQ<br>QQTELSAILETATDYGQAQLFLNASSLANSDWNIENWQPEMLETALAIAQKWKDGLV |
| WP_013552756.1 | Cellulophaga algicola | MGNKKPKLGIAIAPSQKFLEAALPLFAEEKIEIIEWSFDTLKDEADEPAWLSLLLKEYGEKNRLLGHGVFYALLDANWSSRQENWLKKVRQETLSHKYCQISEHFGLM<br>SSANAHSGFPLPIPLSNPVLQIGIDRLKRLQATAQVDVGIENLALTANVADILEQGEFLFKLVNPVNGFVILDLHNIYCQSENFNIDMMTIIKSYPLSLVKELHISGGSWD<br>TDPTLSRKIRRDTHDGRIPEVILEILPEVLKICPFLAFIIFEKIEDSFLTEKDGIDFRADFQKIREIIDATSFSVTPKEKKQVAIILGPPVIDVELLQAQVALRESIALDTYQTN<br>SEWDKDMWKVATKLYEKWNN |
| WP_013687393.1 | Fluviicola taffensis | MSKEPLKIQSGVSCNLDNSLLQATLPLFAAEKVEAIEWSFDTLYKVREIPEWFVELLHAFGNEGRLIGHGVYFSLFSGKWTAEQQRWLDHLRKVSSNFTFNHITEHF<br>GFLTGEDFHKGAPISVPYTTETLNIGIDRLKRIQEACSCPVGLENLAFSYSLEEVQRQGEFLERLIEPVNGFIILDLHNLYCQLQNFSLDFETLIQAYPLDRVREIHISGG<br>SWESSGIHPQRTIRRDTHDDAVPEAVFELLKQTLPLCRNLQFVVLEQLGVALHTLESQQQFQTDFNLMQSIVNEFNAQNQSISNADFLPKEIFQLNEQIPESVTLHQ<br>QQLELSSILEHAPNYEEATRLLSNSSLAYSAWKTEAWSPYMLETALRIAQKWKQGW |
| WP_010467436.1 | Acaryochloris sp. CCMEE 5410 | MTPQIGLSLMPQPDFWQAAQPLFDASEVEVVEWSFDMGWGGVSLPDPLCSVLRQFSQRNGLLGHGVSYSLLSAQLDRRHWLACLQAECRDYSYRHISEHFGW<br>MATRNFARSAPLPMPMLPETLKLGCDRIQQFAEIAQVPVGLENLAFAFGLQDVRQQGEFLDQLLESVDGFLLLDLHNLYCQMHNFQLSATEILNLYPLARVRELHLS<br>GGSWSHHGTATIRRDTHDQAVPELVFELLKLALQRCSSVEAIILERMGNTLNSEQEQIQFRQDFTHIRQITDRFTIEART |
| XP_004338586.1 | Acanthamoeba castellanii str. Neff Neff | MEAEQAGGPWVGVSLMPHPQYRRAIQPLLDQHLIDAIEWSCDIGWGMGGRLPAWVVDLLDQFGRAGRLFGHGVMYSPTSAIWERRQAAWLRWLERECEERRF<br>GHITEHFGFMTAGGALEGCPLPLPFTPGAVEVGVDRLRRMHRAAGGATPVGLENLALAMGPADAADQGPFLAAVLERLGDQGVLLLDLHNLHCQIVNFSLSAQDAI<br>KRLACDPEGRPFRRDTHDGDVPPEIIDELLPLALRLCPNLRVVVLERLGETIRSEADAERLREDFCRIRSVVRDFASAQRAQSTAGAAAAADESAGGRATAEADHH<br>NNINATTATDDDDGGGDEAALESPPFEDEELGRYQAAFLRLFVRDLSPEQLLHQLRAQPEFAPFQKYVEGFDVRCVEVASVLVRKWVRFARPKAYESRWRRPRP<br>DSEGEEDEEEEEEEEYAEEDGEGGDGGECDVYESSEADE |
| WP_052346243.1 | Hymenobacter swuensis | MPTPTFPLPVEHPPVLATLACNLDADILAAAFPLLEAGRVAALEWSFDTLFAVEQVPEWFMALLRSYSTQGRLIGHGVYFSLLSGRWTAEQQQWLEQLRQLSAQVA<br>FDHVTEHFGFFTGENFHSGAPLPIPYTSSTLRLGQDRLRRVQAACGCPVGLENLAFAYSLAEVQRHGEFLAKLVAPVNGFLILDLHNVYCQLHNFSLSTESLLSAFP<br>LELVREIHISGGSWQPSGQVPGRQIRRDTHDEAVPEEVFQLLEYALPLCPNLKFVVLEQLGNALHTEASRTRFRADFSRMETLVQQRPQRPQNPIYQSFLPGPAGP<br>LTAAAAEDEQLHTQQRHLAHILETATSFAEARHQLRASPLAHTAWKIEDWEPHMLETAISIAQKWKQ |
| EFK35127.1 | Chryseobacteriu m gleum ATCC 35910 ATCC 35910 | MRKPMLGVSMMAEADFVSAILPLLQNNSIDVLEWSFDTLYHPNEPDWLRDLLNFYAENGRLIGHGVYYSLFDARWTERQDEWLKKLKEEVRLRNYNHITEHFGFM<br>NTENFHQGVPLPVSLHPKTLQIGKDRLCRLQEAVDIPVGIENLAFSFSVDDVKEQGVFLDQLTEDTNGFLILDLHNIYCQSCNFGVEMKEIVDLYPLNKVKEIHLSGGS<br>WQESAYGKKQVRRDTHDDVIPQDILSVLPSVMEQCQNLEYIIIERLGHTVKTEEEKVSFLNDFNTVRKIIEISDLKLAERNWMKKEMILPEKPLEDRVLYEEQSELTKL<br>LFENADAAVIKNHDFHYFKTENWDSEMILTAQNIIKKWNPY |
| WP_031530742.1 | Dyadobacter crusticola | MSQIFSTVACNLDAHILQASIPLFEMERVEAIEWSFDALFKVDVIPEWFTDFIKEFSDQNRLIGHGVYFSLFSGKWSAEQQKWLDQLSKLCDQFQFDHITEHFGFMT<br>GSDFHKGAPLGIPFNASTLAIGKDRLARIQNACNCPVGLENLAFSYSLDEVKRHGDFLAQLVESVNGFIILDLHNLYCQCHNFELDFIDFLSFYPLERVREMHISGGS<br>WDTQIPGRVVRRDTHDEAVPSEVFEYLRQAIPRCPNLKFVVMEQLGTALDTAEKQAGFQKDFLEMDRIVKSVGATLPDSVKNFQPLSQQVQIGPPIEDLTLHQQQM<br>QLSEILEKARDLHEAKNLLLASDLSNTDWHIERWEPHMLQTAMNIAQKWKAGF |
| WP_034727520.1 | Chryseobacteriu m sp. JM1 | MKKPLLGLAMMPEADFVSAILPLLQTQSVDVLEWSFDTLYDVEEPEWLSGLLDFYSDNSRLLGHGVYYSLFDARWMERQEIWLKKLKEEVKRRKYNHITEHFGFM<br>NTENFHQGVPLPVPLLPKTLQIGKDRLSRLQDAVEIPVGVENLAFSFSMDDVKEQGEFLDRLVEDIDGFLILDLHNIYCQSCNFKVDMMEIINLYPLEKVNEIHLSGGS<br>WQESAYGKKPVRRDTHDDRIPEEILNLLPEVLIQCHPEYVIIERLGHTLNTEVAKQIFFDDFNRVKKILELSDYPVGEEKIWNRQNTVHSKPVEDLLLYAEQTELTRLLF<br>DGNTTESIKNRDFHYFKPESWDEEMITTAQQIIKKWNPY |
| WP_050021162.1 | Chryseobacteriu m sp. P1-3 | MEKPFLGISMMAEADFLTAILPLLRDGSIDVLEWSFDTFYHAEEPDWLTRLLNFYSQNNRLLGHGVYYSLFDAKWTERQSMWLQKLKEEVQQRKYNHITEHFGFM<br>NTENFHQGVPLPVSLHPKTLEIGKDRLCRLQEVLEIPVGIENLAFSFSIDDVKQQGIFLEKLIEDTDGFLILDLHNIYCQACNFGLGIEEIVRLYPLEKVREIHLSGGSWQ<br>TSPYGKRRIRRDTHDDTVPAEILAVLPSVLSQCENLEYVIIERLGHTIQTETEAKDFLDDFLKVKAIIEASEWNSRHKEIKIKKKPECAGRPVEDEGLYREQTILTKLLFD<br>NVKADHFKDFDFQYFKTENWDPEMIRTAQNIIKKWNPY |
| KGP62404.1 | Legionella norrlandica | MPEQDYMDASRPLFESGDVDIIEWSFDMCWGRKIPQPCLDLLTKYSSQGCLLGHGVSFSILTAKRTKRQELWLQYLQNEVNQLNYQRISEHFGFLSTNTFIQGPPL<br>SMPFIPEMLEVGIKNLQQIKQIAKIPIGLENLGFAFCKQDVIEQGKFLTALLEAVDGFLLLDIHNIYCHMINFDMSLDEILDSLPLDRAKEMHISGGSFRSIEQNNRNIYCD<br>SHDAAIPQDLFLLLEKNLSRFSQIDSLIFERLGNTLSNHYEAAQFREDFMTLKSLIRR |
| WP_034751030.1 | Chryseobacteriu m vrystaatense | MKKPLLGLSMMPEADFVSAIVPLLQSNSVEVLEWSFDTFYDAEEPEWLSGLLNFYAGNERLIGHGVYYSLFDARWNDRQENWLQKLKEECKRRKYNHITEHFGFM<br>NTENFHQGVPLPVPLLPKTLQIGKDRLRRLQDAVEIPVGVENLAFSFSIDDVREQGHFLDMLVEDIDGFLILDLHNIYCQACNFQIDMQEIINLYPLGKVREIHLSGGS<br>WQESAYGRRPIRRDTHDESIPDEILKILPKVLSQCYNLEYIIIERLGNTLNNDTEKQIFFSDFNHVKEIIEVSVYPDIIRQDWKRTEKIYSEPVEDLLLHEEQTRLTKLLFD<br>GNNVESVKNGKFLYFKPENWDEEMIMTAQQIIRKWNPY |
| WP_045496050.1 | Chryseobacteriu m sp. StRB126 | MGKPLLGLSMMAEAEFVSAVLPLLQSNSVEVLEWSFDTFYNTEEPDWLSELLHFYSENNRLIGHGVYYSLFDARWTERQEIWLKKLKEEFRKRNYNHITEHFGFMN<br>TENFHQGVPLPVSLHSKTLEIGKDRLYRLQDAVQVPVGIENLAFSFSLDDVMEQGIFLNKLTEDTNGFLILDLHNIYCQSCNFGVEMQEIIKLYPLDKVKEIHLSGGSW<br>QESVYGKKLIRRDTHDDAVPEDILSVLPSVLSQCKNLEYIIIERLGNTLDAVEKKESFLDDFTQIKKIIDSSGAGTQNKDLWTKKEIRFSENPLEDMILHEEQTLLTKLLF<br>ENAGPASIKSHEFHYFKTKNWDPEMILTAQNIIKKWNPY |
| WP_044220505.1 | Phaeodactylibact er xiamenensis | MEYSGLQRPRATLACNLDTNLLRAAHPLLTAGQVEALEWSFDALFSFPEIPDWFTALIQAYSEEGRLVGHGVFFSLFSGRWSAEQQQWLDRLKALSEVYRFDHVT<br>EHFGFMTGADFHKGAPLGIPFTPQTLQLGRDRLQRISAAAGCPVGLENLAFAYTLDEVKRHGDFLEQLLEPINGFIILDLHNLYCQLHNFRQSAEELLRLMPLHRVRE<br>IHLSGGSWEPAAGQRNRQIRRDTHDGTVPQAVFDLLEKALPKCPNLKYVVLEQLGGALETPEQQRAFQADFRTMHGILKRQNHSGKTSAPANDFQPPAFTPSATP<br>RSDEALHQQQQALSEILENAASLRDARTQLSASPLAETAWNVENWEDDMLETARRIAQKWKGGWG |
| WP_040999004.1 | Chryseobacteriu m oranimense | MNKPLLGLAMMPEADFVSAILPVLQNNSVDVLEWSFDTFYDVEEPEWLSGLLDFYAENDRLLGHGVYYSLFDAQWTSRQKDWLKKLKEETKCRKYNHITEHFGFM<br>NTENFHQGVPLPVPLHAKTLQIGKDRLKRLQDTVEVPVGVENLAFSFSMDDVKEQGEFLEKLVEDIDGFLILDLHNIYCQSCNFEIDMQEIIRMYSLERVKEIHLSGGS<br>WQESAYGNKPVRRDTHDDNIPVEITDVLPEVFSQCPNLEYVIVERLGHTLDTEVKKQLFFDDFNKVKEIIEVSDFPAGITKAWNKRKGESSEPVEDLLLYEEQTKLTKI<br>LFEGNTVQSVKNETFHYFNPENWDKEMITTAQQIIKKWNPY |
| WP_027385220.1 | Chryseobacteriu m gregarium | MNDKPLLGLSMMADADFVSAVIPLLESDEIDILEWSFDTSYGADEPQWLTDLLNFYSENRRLLGHGVYYSLFDAKWTERQDYWLQQLKIETEKRNYNHITEHFGFM<br>NTENFHQGVPLPVPLHSEILKIGKDRLLRLQDVVEIPIGIENLAFSLSANDVREQGDFLSRLIEDIDGFLILDLHNLYCQALNFNMDIIELVNLYPLEKVKEIHLSGGSWE<br>NSVYTKNKRIRRDTHDEAIPLELFSVLPEVMARCHHLQYIIIERLGHTLEGESDKLNFFTDYKKVRGIIYNSKRKKPSPHQWSKKDIVLADPLENLLLFNDQKNLNKLLF<br>DSEDPERITNYPFNYFKTENWDLEMIDTAVQIVKKWNPY |
| WP_053328730.1 | Chryseobacteriu m gallinarum | MEKPFLGISMMAEADFLTAILPLLRDGSIDVLEWSFDTFYHAEEPDWLTRLLNFYSQNNRLLGHGVYYSLFDAKWTERQSMWLQKLKEEVQQRKYNHITEHFGFM<br>NTENFHQGVPLPVSLHPKTLEIGKDRLCRLQEVLEIPVGIENLAFSFSIDDVKQQGIFLEKLIEDTDGFLILDLHNIYCQACNFGLGIEEIVRLYPLEKVREIHLSGGSWQ<br>TSPYGKRRIRRDTHDDTVPAEILTVLPSVLSQCENLEYVIIERLGHTIQTETEAKDFLDDFLKVKAIIGASEWNSRHKEIKIKKKPECAGRPVEDEGLYREQTILTKLLFD<br>NVKADHFKDFDFQYFKTENWDPEMIRTAQNIIKKWNPY |
| WP_034698536.1 | Chryseobacteriu m sp. S0630 | MRKPMLGVSMMAEADFVSAILPLLQNNSIDVMEWSFDTLYHTNEPDWLRDLLNFYAENNRLIGHGVYYSLFDAKWTERQEKWLKKLKEEVSLRKYNHITEHFGFM<br>NTENFHQGVPLPVSLHSKTLQIGKDRLYRLQDAVNIPVGIENLAFSFSIDDVKEQGIFLDKLTEDTNGFLILDLHNIYCQSCNFELGMQEIIALYPLEKVKEIHLSGGSW<br>QDSVYGKKQVRRDTHDDIIPQDILSVLPSVLTQCGNLEYIIIERLGHTVKTEQERDDFLNDFNKVKDIIETSDFNGKEKSSWSKMEMKLSERPLEDLVLYEEQSRLTRL<br>LYDNANIETIKDLDFHYFKTEKWDSEMILTAQNIIKKWNPY |
| WP_046369149.1 | Flavihumibacter petaseus | MTTLAAVALNPDSNLLTAALPLLEEEMVDAIEWSFDAVDTTGRLPAYFHDLLQFYASHHRLIGHGIYFSAFSGKWTPDQQEWLSRLAAVCRQFPLEQVTEHFGFMT<br>GKDFHKGAPLNIPRNAATLAIGRDRLARIRQACNCPVGLENLAFSYSLEETRHHGAFLAELVEPVNGFIILDLHNCYCQAMNFSVDADRLLDTYPLHLVREIHISGGS<br>WEPFPDAPGGMVRRDTHDDSVPQEVFDLLQLALRRCPNVRYVTLEQLSTALETTSAQEQFCKDFRRMRGMMAIPEGERQQAASGVPQVDFHRPIPLLTGQIPVE<br>DMLLYQQQRELAGILESSEDLATAQKLLRNSSLAASDWKIENWAPYMVETARQIARKWK |
| WP_046244672.1 | Hymenobacter terrenus | MEPTPAILASLACNLDSDILTAAFPLLEDGQVEALEWSFDALFRTQELPEWFVELLQTYSRRQRLVGHGVFFSLLSSKWTPEQQQWLAQLRQVSLQFAFDHITEHF<br>GFFTGQNFHSGAPLPVPYTNTTLRIGQDRLRRISDACQCSVGLENLAFAYSLAEVRQHGEFLEKLIAPVNGFIILDLHNLYCQLHNFSVPYENLIGLYPLDCVREIHISG<br>GSWEDSGVLPHKKVRRDTHDDAVPAEVFHLLELTMPKCPNLKYVVLEQLGTGLRSDSSKAQFRRDFQKMASMVQQHRHGPHRQPAHEFRPLQLPVLGTAIEDEQ<br>LQTQQIQLSHILENASTFEAAHRQLLSSSLANSAWEIENWEPHMLETAVRIAQKWKK |
| WP_044218508.1 | Chitinophaga pinensis | MPKVLSAVACNLDANILAACLPLMEESRIEAIEWSFDALYKVKEVPDWFRELLTAFSDENRLIGHGVFFSLFSGKWLPEQENWLKHLSQTADAFQFDHITEHFGFMT<br>GKDFHQGAPLNIPYTAITLNIGRDRLKRIYEACKRPVGLENLAFSYSLDEVKRHGAFLEQLLEPVNGFIILDLHNLYCQLRNFDVTFEEMIALYPLDRVREIHISGGSWE<br>DSVATPSRSIRRDTHDDAVPTEVFQLLEMTIPRCPQLKYVVLEQLGNGLETETSRQYFYNDFLRMQEIIQQHNDQADDTSTDPFLPLIPFSTDAVVEDMMLYQQQLE<br>LSAIFESSASYGEAMQSLSTSSLANSDWRIEQWEPHMIETAVKIAQKWMK |
| WP_034713482.1 | Chryseobacteriu m soli | MGKPLLGLSMMAETEFVSAVLPVMQTNSIEVLEWSFDTFYHTEEPEWLSELLDFYAESNRLIGHGVYYSLFDARWTGRQEMWLEQLKKETQKRNYNHITEHFGFM<br>NTDNFHQGVPLPVSLNSKTLQVGKDRINRLQDAVDIPIGIENLAFSFSIDDVKEQGEFIHKLIEDIDGFIILDIHNLYCQACNFETDIWELIQLYPLEKVKEIHLSGGSWQK<br>SAYGRNPVRRDTHDDSIPEEIFAIIPQIFSQCTHLEYVIVERLGHTIKTQKEKQCFFNDFNKVKEIIDASDDRLDNKKVWHKKEINLSAPVEDLFLYEDQTKLTKLLFDAV<br>DETAIKRERFHYFEPEKWDLEMINTAQAIIKKWNPY |
| WP_062699011.1 | Chryseobacteriu m indologenes | MKKPLLGLSMMPEADFVSAILPLLQDQSVAVLEWSFDTVEHVLEPEWLRGLLDFYAENSRLLGHGVYYSLFDARWTERQEVWLEKLKKETQRRKYNHITEHFGFM<br>NTENYHQGVPLPVPLHSRILQIGKDRLAHLQAAVEIPVGVENLAFSFSADDVREQGIFLEKLIEDVEGFLILDLHNIYCQSCNFDIDIVDIVAMYPLEKVKEIHLSGGSW |

| | | |
|---|---|---|
| | | QESVYRKEPVRRDTHDDAIPEEVLAVLPQVLSRCPNLEYIIIERLGHTLNTTMEKEAFFNDFVRTKKIIEDSWFPAGNKTVWNRKECISSVPLEDHLLYHEQTKLTRLLYGNSDITSFKNENFSYFDVQKWDAEMIETARNIIKKWNPY |
| WP_051884865.1 | Chryseobacterium luteum | MSKPLLGLSMMPEADFVSAILPLLQNNSVDVLEWSFDTFYDTQEPEWLSGLLDFYAENGRLLGHGVYYSLFDARWTERQEGWIEKLKEEFRHRKYNHITEHFGFMNTENFHQGVPLPVPLHSKTLQIGKNRLRRLQDAVEIPVGVENLAFSFSINDVKEQGEFLDKLIEDIDGFLILDLHNIYCQSCNFEINMQEIIRLYPLEKVKEIHLSGGSWQESAYGEKTVRRDTHDDSIPDEILNILPEVLSRCHNLEYVIIERLGHTLNSEAKKQAFFDDFNRVKEIIEISDPFTGNKKVWNKKNGQYSEPVEDLLLHNEQTKLTKMLFEGDSIQSVKNKPFHYFKPGKWDEEMITAAQQIIKKWNPY |
| WP_083997424.1 | Chryseobacterium angstadtii | MKKPLLGLSMMPEADFVSAILPLLQDHSVEVLEWSFDTFFDEKEPEWLSGLLDFYSANNRLLGHGVYYSLFDARWTGRQEIWLEKLKEECKHRKYSHITEHFGFVNTENFHQGVPLPVPLHPQILQIGKDRLKRLQDAVEVPVGVENLAFSFSIEDVKEQGEFLYRLVEDIDGFLILDLHNIYCQSCNFEKDMQEIIRQYPLEKVKEIHLSGGSWQESAYGKKLIRRDTHDDRIPDEILLVLPEVLSQCPSPEYVIIERLGHTLNTEAERHIFLEDFKKVKDIIEAFDGYSDENKPGSRMTGKYSEPVEDFLLYDEQTKLTKLLFDENNVEFIKNQKFHYFKPENWDEEMINTAQQIIKKWNPY |
| WP_083996892.1 | Chryseobacterium sp. BLS98 | MKKPLLGLAMMPEADFVSAILSILQNNSVDVLEWSFDTFYDVEEPEWLSGLLDFYAENDRLLGHGVYYSLFDARWTSRQKDWLQKLKEEFRHRKYNHITEHFGFMNTENFHQGIPLPVPLHSKTLQIGKDRLKRLQDTVEVPVGVENLAFSFSMDDVKEQGEFLEKLIEDIDGFLILDLHNIYCQSCNFKIDMQELIRMYPLEKVKEIHLSGGSWQESSYGNKPVRRDTHDDDIPDEIIDVLPKVLSQCLNLEYVIIERLGHTLDTEGKKQLFFNDFNKVKEIIEVSDFPTGTTKPWNKRKRESSEPVEDLLLYEEQTKLTKILFEGNAAQSVKNEAFHYFKPENWDKEMIATAQQIIKKWNPY |
| KQM24483.1 | Chryseobacterium sp. Leaf201 | MMAEADFVSAILPLLESSGIDVLEWSFDTFYTVDEPQWLADLLNFYSDHKRLLGHGVYYSLFDAKWTERQDVWLQKLKAETDRRSYNHITEHFGFMNTENFHQGVPLPVPLHPELLKIGKDRLLRLQDVVEVPIGIENLAFSFSSDDVSEQGEFLYRLIEDIDGFVILDLHNIYCQASNFNMEIMELVNLYPLEKVKEIHLSGGSWENSIYTKNKRIRRDTHDDEIPQELFDILPDVISRCSHLEYIIIERLGHTIKTESDKQNFLRDFKKVKQIIDNSDPKENLQHQWKKKNRVLTDPMENLLLFDDQKNLNKFLFESQDPETIRNYAFHYFKMHNWDPEMIHTASQIVKKWNPY |
| WP_054508689.1 | Chryseobacterium sp. ERMR1:04 | MRKALLGLSMMAEPDFVTAILPLLENSEIEVLEWSFDTFYGVEEPQWLSNLLNFYAENNRLIGHGVYYSLLNAVWTDRHHEWLENLKTETQKRRYTHITEHFGFMNTENFHQGIPLPLPLNAVTLQTGKDRLQRLQNVLDVPVGIENLAFSFSKQDVEEQGEFLSRLIEDVDGFLILDLHNIYCQAHNFEIDMKELIRLYPLDQVKEIHLSGGSWQESIYSKDKYVRRDTHDDCIPEDIFQILPYVLKQCKNLEYVIIERLGNTIGAEKDKASFIKDFRRAKKIIDAIKLPLLTNHWIKKEKKIVEPFEDHLLFEDQKKLTKLLCESNTPEDIMTEKLHYFKSEDWNKEMINTAKNIIKKWNSY |
| WP_059069066.1 | Solirubrum puertoriconensis | MEAAPPLYASVACNLDADLLAAAYPLLEEGRVEALEWSFDALYWAEQVPEWFTNLLHTYAGAGRLVGHGVFFSLLSGKWSPEQQQWLAHLGQLAEQFRFDHITEHFGFFTGQNFHHGAPLPIPYTATALRIGQDRLRRIYDACCCPVGLENLAFAYTLDEVKRHGNFLEELLLPVNGFVILDLHNLYCQLHNFGVPYDELIALYPLERVREIHISGGSWHDSEQAPSQRIRRDTHDDAVPEEVFQLLELTMPRCPNLKFVVLEQLGHALHAEASRTQFREDFGRMESLVARHRAALRTLPTNDFRPQQTWPPSTAAEDEHLHRQQRQLSDILENAPSYAEAQRLLQASGLAHSDWRIEQWEPHMLETALHIARKWKT |
| KXH81096.1 | Chryseobacterium kwangjuense | MPEPEFVSAVLPILQDHSVDVLEWSFDTLYDLEEPEWLSGLLDFYAENNRLLGHGVYYSLFDARWTERQETWLEKLKKEVKHRNYNHITEHFGFMNTENFHQGVPLPVPLHSKTLQIGKDRLGRLQDAVEIPVGIENLAFSFSVNDVKEQGEFLDKLIEDIDGFLILDLHNIYCQSCNFETDMLEMIRMYPLEKVKEIHLSGGSWQESAYGKKPVRRDTHDDCIPDDILKILPEVLSLCPNLDYIIIERLGHTLKTESEQKIFFDDFESVKKIISKASYPSGQSKIWLKKKIGAYSEPVEDQLLYDEQTKLTQLLFNGTDIQSIKDQDFHYFKPENWDDEMIFSAQQIIKKWNPY |
| WP_162265596.1 | Legionella santicrucis | MTPPKMGLSLMPEQDYLDASLPLFESGQVEVIEWSFDMCWGKKIPQRCHDLLEKYSDDNCLLGHGVSFSILTAKRTKRQEIWLQYLQKEMSHLHYQNISEHFGFLTTDTFIQGPPLSMPFIPELVEVGINNLQEIKQIAQIPIGLENLGFAFSRQDVIDQGKFITALLDAVDGFLLLDIHNIYCHMLNFNMSFEEILETLPRHRLKEMHISGGSFRDICYQNKTIYCDSHDAAIPEDLFLLLEKNLPYLLPMESIIFERMGNTFSSPYEVSRFQDDFANLKSIVQKASDEMFK |
| WP_059013965.1 | Flavobacterium psychrophilum | MKKPQLGISLMTEPAFLEACFPLFAEGEIEVLEWSFDTIQTIEDEPLWLNELLSTYSRENKLLGHGVYYSLFDAYWGDKQDLWINKIKKETERYKYNHISEHFGFMSSENYHKGVPLPIPLSDSTLNIGIDRLKRIQNAVDLPVGIENLAFSFSKKDVFEQGVFIEKLIDSVNGFLVLDLHNIYCQSHNFEIDMMELINYYPLSKVKEIHISGGSWQDSIYKEKQIRRDTHDGNIPLEILSILPLVLSKCVNTDFVIFERLGNTFNNSYDVTEYQKDYKELKKIVQQIEYKSNQESWRKEYKISEKPLVNFELKKEQQKLSNFLLTEDYNEIKNKEFKYWDVNSWDIEMINTAIEISKKWNQ |
| WP_063962962.1 | Chryseobacterium kwangjuense | MKKPLLGLSMMPEPEFVSAVLPILQDHSVDVLEWSFDTLYDLEEPEWLSGLLDFYAENNRLLGHGVYYSLFDARWTERQETWLEKLKKEVKHRNYNHITEHFGFMNTENFHQGVPLPVPLHSKTLQIGKDRLGRLQDAVEIPVGIENLAFSFSVNDVKEQGEFLDKLIEDIDGFLILDLHNIYCQSCNFETDMLEMIRMYPLEKVKEIHLSGGSWQESAYGKKPVRRDTHDDCIPDDILKILPEVLSLCPNLDYIIIERLGHTLKTESEQKIFFDDFESVKKIISKASYPSGQSKIWLKKKIGAYSEPVEDQLLYDEQTKLTQLLFNGTDIQSIKDQDFHYFKPENWDDEMIFSAQQIIKKWNPY |
| WP_056291594.1 | Dyadobacter sp. Leaf189 | MPQIFSTIACNLDAHILQSSIPLFEMEKVEAIEWSFDALFKADVVPEWFTDLIGEFSDQGRLICHGVYFSLFSGKWSAEQQHWLDQLSKLCGQFHFDHITEHFGFMTGSDFHKGAPLGIPFNASTLAIGKDRLARIQNACNCPVGLENLAFSYSLDEVKLHGDFLGQLVESVNGFIILDLHNLYCQCHNFELDFTEFLSFYPLDRVREMHISGGSWDAQIPGRIVRRDTHDEAVPPEVFEYLRQAIPRCPDLKFVVMEQLGTALDTAEKQADFQKDFLQMDHIIKSVGETGRISANNFLPFSPTSPAGLPVEDLALYQQQMQLSNILEIAGDLKEAQRLLNASDLSDTDWHIEHWEPHMLQTAMNIAQKWKGGF |
| WP_062670360.1 | Chryseobacterium cucumeris | MRKPILGVSMMAEPDFVSAILPLLHNNSIEVMEWSFDTLYHTNEPDWLRDLLNFYAENNRLIGHGVYYSLFDARWTARQEEWLKKLKEEVSLRKYNHITEHFGFMNTENFHQGVPLPVSLHSKILQIGKDRLYRLQDAVNIPVGIENLAFSFSIEDVKEQGVFLDKLTEDTNGFLILDLHNIYCQSCNFELEMQQIIDLYPMDKVKEIHLSGGSWQDSVYRKKQVRRDTHDDVIPKDILSVLPSVMKKCPNLEYIIIERLGHTLKTEEKRNSFMEDFTKVRTIIETSDLKLKQQNGWCKEEIKLPAKPLEDLLLYEEQSRLTRLLYENANVENIKDLDFHYFKTENWDAEMILTAQNIIKKWNPY |
| WP_056076704.1 | Chryseobacterium sp. Leaf394 | MNNKPLLGTAMMAETEFVSAVLPLLENNEIDILEWSFDTFYHADEPKWLSGLLDFYSENNRLIGHGVYYSLFDAKWTDRQEHWIEKLKSETKKRNYNHITEHFGFMNTENFHQGIPLPVPLHSKILQLGKDRLLRLQDALEIPIGIENLAFSFSADDVKEQGEFLEKLIEDIDGFLILDLHNLYCQAVNFNIDILELVAMYPLEKVREIHLSGGSWENSSYTEEKRIRRDTHDDSIPREILEVLPKVLSVCHQLEYIIIERLGHTLTEKSDTTNFLNDFKKVRQIVNASEVNSFLKKNWKKKNQITGKPLEDLLLYEDQTKLTKLLFDSVNPEIIKNYHFNYFEIEKWDLEMIDTASKIIKKWNPY |
| WP_065397536.1 | Chryseobacterium arthrosphaerae | MKKPLLGLSMMPEADFVSAILPLLQSNSVEVLEWSFDTFYEAKEPEWLDELLNFYSENQRLIGHGVYYSLFDALWTERQENWLKKLKEEVRKRKYNHITEHFGFMNTENFHQGVPLPVPLHPKTLLIGKDRLYRLQDAVQLPVGIENLAFSFSIEDVKEQGVFLDNLIADINGFLILDLHNIYCQACNFEVEMEDIIRLYPLDKVREIHLSGGSWQESVYARKMIRRDTHDDLIPEEIFSVLPAVITQCNNLEYVIIERLGHTIKTDQEKRAFLDDFTRARRIIDDSEIKTGKKIRWGQKEMGFSNSPVEDDALYEEQSRLTKLLFDSSGAEMIKEQRFHYFKTENWDPDMILTAQNIIKKWNPY |
| WP_065394721.1 | Chryseobacterium artocarpi | MKKPLLGISMMAESEFVSAILPLLQSSSVEVLEWSFDTFYNAKEPVWLSDLLNFYSENNRLIGHGVYYSLFDAKWSDRQEVWLERLKQEVHKRQYNHLTEHFGFMNTENFHQGVPLPVSLHSQTLQIGKDRLHRLQDAVEIPVGIENLAFSFSVDDVKEQGLFLDKLIEEIDGFLILDLHNIYCQACNFDVDMLEIIQSYPLDKVKEIHLSGGSWQESVYGKKSIRRDTHDDDIPDEIWAVLPSVLLQCKNVQYVIIERLGHSIKTERDKQNFLSDFVKAQTIINVLDWSGRENRFWSKKEVNISKKPLEDVILYEEQTRLTKLLFDSIDPVLVKKQDFQYFNTETWCEEMIMTAQNILKKWNPY |
| WP_066690840.1 | Chryseobacterium contaminans | MGKPLLGLSMMADVEFLSTILPLLQSNSVEVLEWSFDTFYNAQEPDWLHDLLNFYSESNRLVGHGVYYSLFDALWTERQEVWLKKLREEFRKRKYNHITEHFGFMNAENFHQGVPLPVSLHAKTLQIGKDRLYRLQDAVEVPVGIENLAFSFSVDDVMEQGIFLDKLTEDTNGFLILDLHNIYCQSCNFEVEMSEIIQLYPLDKVKEIHLSGGSWQESVYRKKPVRRDTHDDAIPGSIFSVLPSVLSQCKNLEYIIIERLGDTIDTEDKKKSFLDDFIQVKTLIDSSGREIQNKNSWTKKEVSFPENALEDLLLHEEQTLLTRLLFDNTEATYIKSHEFHYFKTENWDPEMILTAQNIIKRWNPY |
| SEH31972.1 | Chryseobacterium culicis | MMAEADFVSALLPLLQSNSVDVLEWSFDTLYHNHEPVWLCDLLNFYADNERLIGHGVYYSLFDARWTERQEDWLKKLKEEVALRRYNHITEHFGFMNTENFHQGVPLPVSLHAKTLEIGKDRLYRLQDAVNIPVGIENLAFSFGIDDVKEQGAFLDKLIEDTDGFLILDLHNIYCQACNFGIEIQEIMDWYPLDKVKEIHLSGGSWQESVYGKRPVRRDTHDDSIPKELFSVLPSVVSQCQHLEYVIIERLGHTLKTDEEKNDFLTDFNRVKKIIEASDWKGEQKKNWKKKEIKLTEKPLEDRVLFEEQSRLTQLLFDHVDAAVIKDVDFHYFKTENWDLEMILTAQNMIRKWNPY |
| WP_068943005.1 | Chryseobacterium | MRRPLLGLSMMAEAEFISAIMPLLQSNSVEVLEWSFDTFYNMEEPSWLGDLLNFYAENNRLIGHGVYYSLFDAQWSERQEMWLSQLKQEFQKRKYNHITEHFGFMNTENFHQGVPLPVSLHSKTLEIGKDRLYRLQDAVQVPVGIENLAFSFSLDDVMEQGAFLDQLTEDVDGFLILDLHNIYCQSCNFEVEMQEIIKLYPLNKVREIHLSGGSWQESAYGKKPVRRDTHDDSIPEEILLILPSILSQCENLEYMIIERLGHTINTEEKRNRFLDDFDKVRALIDSSDLEIQNKNPWSKKEVQFPQKPLEDMILHEEQTLLTKLLFDSVEAVSIQNHEFHYFKTENWDPEMILTAQNIIKKWNPY |
| WP_245705439.1 | Chitinophaga filiformis | MPKVLSAVACNLDANVLAACLPLMEESRVEAIEWSFDALYKVKEVPSWFRELLTAFSQEKRLIGHGIFFSLFSGKWLPEQENWLKHLAQTAAEFQFDHITEHFGFMTGKDFHHGAPLNIPYTTTTLNIGRDRLKRIYEACGRPVGLENLAFSYSLDEVKRHGAFLEQLLEPVNGFIILDLHNLYCQLRNFDISFDEMIALYPLERVREIHISGGSWDDSNADPKRSIRRDTHDDAVPPEVFQLLEMTIARCPHLKYVVLEQLGNGLATEESRRCFYNDFLQMEEIVRRNNNENGTDASDPFIPLFPLSTGTVVEDMTLYQQQLELSSILESSASYTEAIYALTQSSLANSDWRIEQWEPYMIETAVKIAQKWKK |
| WP_090148480.1 | Dyadobacter soli | MSEIYASVACNLDAHILQASLPLFEQEKIEAIEWSFDTLFKFDEIPAWFTDLVSEFSNHNRLIGHGVYFSLFSGKWSPAQQQWLDKLKRLSAEFRFDHITEHFGFMTGEDFHKGAPIGIPRTARTLALGQDRLRRIQDACQCPVGLENLAFSYSVDEVKKHGEFLDQLVETVNGFIILDLHNVYCQVHNFDIAFDDIIKLYPLNKVREIHISGGSWQQTEAIPGKSVRRDTHDESVPEAVFEFLGKAIPECPYLKFVVLEQLGTALDTPSSQAAFQADFLRMDYIIKSLAHGYPQAAVYPFAPLPGLAPPVPLEDPDLHKQQVQLSRILETAGSVSEATAMLQSSDLRNTAWEVEKWAPYMLETAISIAQKWKDGFV |
| WP_090332217.1 | Dyadobacter koreensis | MPKILSSIACNLDTHILLAALPLFEQQKIEAIEWSFDTLFKYEEIPEWFTDLISEFSNQGRLVGHGVFFSLFSGKWLPEQDIWLKKLTAFSERFKFDHITEHFGFMTGSDFHKGAPIGIPYTKSFLDIGIDRLKRIQDACQRPVGLENLAFSYSLEEVGNHGDFLSKLVESVNGFIILDLHNLYCQIHNFRMDFEDIIVLYPLEKVRELHISGGSWDERKTATGEKIRRDTHDDRVPEEVFNLLSATIPKCPNLKFVVLEQMGTALTSEENQLAFQRDFLKMDAIVKSINYAFPQSLPNNFLPVNRAILSRFPLENLALFEQQQQLSYILETAENFGHAQQLLAKSALHQTEWKVENWEPYMLETAMAIAQKWRNGF |
| WP_218144591.1 | Hymenobacter actinosclerus | MASTPSSATTPASPVLAALACNLDADLLTAAFPLLEQGRVAALEWSFDALFAVEQVPEWFQELLRAYSAQGRLLGHGVYCSLLSGRWTHAQQQWLTQLRTLARQLPFDHVTEHFGFFTGANFHAGAPLPIPYTSATLRLGQDRLQRVQAACECPVGLENLAFAYSLDEVRRHGEFLEKLVEPVNGFLILDLHNVYCQLHNFRLPADELLALYPLHRVREIHLSGGSWAESGQEPGRQIRRDTHDDAVPAEVFQLLKWALPRCPNLKYVVLEQVGGALRTARSQAQFRVDFGRMEALVQEYQKGSARQPGNAFRPRLPLAPPGPVIEDELLHTQQRQLGHILETAASFEQARQRLQASSLAHTAWQIEHWEPAMLETAISIAQKWQ |
| ASS50882.1 | Candidatus Fluviicola riflensis | MIKQQHILASVACNLDQHLLQATLPLFIDEQVEAIEWSFDALYKVRNIPDWFVELLTTFGKEGRLIGHGVFFSLFSGKWSVEQERWLNHLKKVSGNFTFNHITEHFGFMTGEDFHKGAPIGIPYNSTTLGIGIDRLKRIQDTCQCPVGLENLAFSYSLEEVKRHGEFLNELIEPVNGFIILDLHNFYCQLHNFSLDAEELLELYPLDRVREIHISGGSWEQTRLEPQRTIRRDTHDDAVPEAVFELLKLVIPRCPNLKFVVLEQLGIALDSTEKQQQFQADFSRLKSIVSTFDSPNPVNDFLPKSTFVLSSSVAEDSFLASQQQELSDILENAINHENAVVLLAKSGLANSDWKVEEWSPYMLETAMRIAQKWKNGW |
| WP_245752765.1 | Chitinophaga arvensicola | MPEILSAAACNLDSNILAACLPLMEESRLEAIEWSFDALFNVKDIPSWFLELLTAYSHEKRLTGHGVFFSLFSGKWRQEQADWLKRLKKINSTFRFDHVTEHFGFMTGKDFHHGAPLNIPYTPTTLRIGQDRLKRIFEACECPVGLENLAFSYSLEEVKKHGAFLEQLLEPVNGFIILDLHNLYCQIHNFNIPFEEILALYPLHKVREIHISGGSWQPAPDNPVKKIRRDTHDDAVPEEVFALLEHTIARCPHLKYVVLEQLGNALYTANSKHLFYTDFGRMEKIVQANNKIHKAVSLQTFLPLSPIINGAVIEDETLNAQQLELSRILESAHSHHEAIYRLSQSSLAHSAWQVEEWEPDMIATAVSIAQKWKK |
| WP_090381944.1 | Dyadobacter sp. SG02 | MSEIYSSIACNLDAHILQASLPLFEQERVEAIEWSFDTLFRFDEIPAWFTDLVSEFSNGNRLIGHGVYFSLFSGKWSSGQQEWLDKLKKLSDEFLFDHITEHFGFMTGEDFHKGAPIGIPFTSQTLALGKDRLRRIQDACRCPVGLENLAFSYSMDEVKKHGDFLRELVESVNGFIILDLHNIYCQVHNFDIAFDDIIKLYPLDKVREIHISGGSWEHTLMNSGRPVRRDTHDDAVPEVVFEFLGSAIPKCPNLKYVVLEQLGTALDNESSQAAFQSDFLRMDSIIKSLTPKSPQIPADTFAPLPANANTTPLEDPRLHRQQVQLSRILETASDVNEARTMLLSSDLHNTAWNVENWAPYMLETAIAIARKWRNGFE |
| SFN39809.1 | Chryseobacterium oleae | MMPEAEFVSAILPLLQTQSVDVLEWSFDTFYDAEEPEWLSGLLDFYAENNRLLGHGVYYSLFDGRWANRQEIWLKKLKEEVKRRKYNHITEHFGFMNTENFHQGVPLPVPLLPKTLQIGKDRLSRLQDAVEIPVGVENLAFSFSMDDVKEQGEFLDKLVEDIDGFLILDLHNIYCQSCNFEVDMMEIINLYPLEKVKEIHLSGGSWQESAYGKKPVRRDTHDDRIPEEILNLLPEVLILCHPEYVIIERLGHTLNTEAEKQIFFDDFNRVKKIIELSDYPVDEEKIWNRQNTVHSKPVEDLLLHAEQTELTRMLFDGNTTESIKNRNFHYFKPESWDEEMITTAQQIIKKWNPY |
| WP_093832332.1 | Spirosoma endophyticum | MTPIRSTLACNLDTNLLSACLPLFEAGSVQAIEWSFDTLYQHRQSPDWFTELLTTFSDAGCLIGHGVFFSLFRGQWTPDQQAWLDQLKDLASTYQFNHITEHFGFMTGADFHKGAPLSIPLNASTLAIGQDRLRRIQDACGCPVGLENLAFAYSVDEVKRQSDFLNRLLEPINGFLILDLHNLYCQSQNFGMSGADLLDLYPLDRVREIHISGGSWSTSTIEPERIIRRDTHDDSVPAEVFAWLEQAINRCPNLRYVVLEQLGPALATEQERHFFQDDFRRMDQLVQAANQTLSVRETNTFLPANSSANPSVPVEDTFLFAQQRTLATILETAIDTRQAQTLLATSSLANSDWAIEQWEPAMLETALAIAQKWRNGFN |

| | | |
|---|---|---|
| WP_089733898.1 | Chryseobacteriu m jejuense | MGKPLLGLSMMAEAEFVSAVLPLLQSNSVEVLEWSFDTFYNVEEPDWLSELLDFYSENNRLIGHGVYYSLFDARWTDRQEIWLKKLQKEFRKRNYNHITEHFGFMNTENFHQGVPLPVSLHPKTLKIGKDRLYRLQDAVQVPVGIENLAFSFSLDDVMEQGVFLDKLTEDTDGFLILDLHNIYCQSFNFEVEMQEIIKLYPLDKVKEIHLSGGSWQESVYGKTLVRRDTHDDIIPEDILFVLPSILSQCKNLEYVIIERLGNTINTDEKKKNFLDDFTKVKNLIDSSDFGTENKDPWTKKEVRFSENPLEDLMLHEEQTLLTKLLFDNAGPESIKNHDFHYFKTKNWDPEMILTAQNIIKKWNPY |
| WP_255373497.1 | Chitinophaga sp. CF118 | MPKILSAVACNLDANILSACMPLMEESRVEAIEWSFDALYKVKEVPSWFRELLTAFSNEGRLIGHGVFFSLFSGKWLPEQESWLKHLEQTAAEFRFDHITEHFGFMTGKDFHHGAPLNIPYNTTTLNIGRDRLKRIYEACRRPVGLENLAFSYSLDEVKRHGAFLEQLLEPVNGFIILDLHNLYCQLRNFDIAFDEMIALYPLDRVREIHISGGSWDDSAADPNRSIRRDTHDDAVPPEVFRLLEMTITQCPHLKYVVLEQLGNGLITETSKRCFYNDFLQMEEIVRKNSNESDAGSPDPFLPLSPLSTGIAVEDMTLYQQQLELSSILESSASYAEAMNVLSQSSLANSDWRIEQWEPYMIETAVKIAQKWKK |
| WP_072362160.1 | Chitinophaga sancti | MPKILSAVACNLDANILAACLPLFEESRVEAIEWSFDALYKVKEVPGWFRELLSAFSDENRLIGHGVFFSLFSGKWLPEQEAWLSHLQQTATEFSFDHITEHFGFMTGEDFHHGAPLNIPYSSSTLNIGKDRLKRIYHACGRPVGLENLAFSYSLDEVKRHGTFLEQLLEPVNGFIILDLHNLFCQLKNFDLDFNTLISLYPLDKVREIHISGGSWDDSAAAPDRSIRRDTHDESVPPEVFQLLEMTIPQCPNLKYVVLEQLGNGLQSTASKQGFYNDFLQMQEIAQQHNYKEGNPFLPLESLPIITGAAIEDLRLYQQQLELSAILENAGTYAEAMQQLQQSSLAHSDWKIEEWEPYMIETAVKIARKWKK |
| WP_073063557.1 | Chryseobacteriu m oranimense | MNKPLLGLAMMPEADFVSAILPALQNNSVDVLEWSFDTFYDVEEPEWLSGLLDFYAENDRLLGHGVYYSLFDAQWTSRQKDWLKKLKEETKRRKYNHITEHFGFMNTENFHQGVPLPVPLHAKTLQIGKDRLKRLQDTVEVPVGVENLAFSFSMDDVKEQGEFLEKLVEGIDGFLILDLHNIYCQSCNFEIDMQEIIRMYPLERVKEIHLSGGSWQESAYGNKPVRRDTHDDNIPVEIIDVLPEVFSQCPNLEYIIIERLGHTLDTEVKKQLFFDDFNKVKEIIEVSDFPAGITKAWNKRKGESSEPVEDLLLYEEQTKLTKILFEGNTVQSVKNEAFHYFNPENWDKEMITTAQQIIKKWNPY |
| WP_217646801.1 | Pontibacter chinhatensis | MPGILATVACNLDADILSATYPLLEEGRVEAMEWSFDTLFQVEQLPDWFREMLLAYSREGRLIGHGVFYSLISGRWSREQAQWLKNLRQLSAQFHFDHITEHFGFFTGQNFHHGAPLSIPYTSTTLRIGRDRLLRIQDACQRPVGLENLAFSYSLDDVKQHGEFLEKLIAPANGFIILDLHNLYCQLHNFSVSYKELIRLYPLERVREIHISGGSWEEAPEAPGQKVRRDTHDDAVPAAVFGLLEQTMPLCPNLKYVVLEQLGNGLKTESSRAGFQRDFRQLEQLVQQANAVRTGHQPANLFLPLNALPGSAVEEDEALHQQQVLLSQILENAPSYQEAQRLLQTSSLAHSDWRIENWQPYMLETAIRIAQKWKK |
| WP_255373162.1 | Chitinophaga sp. YR627 | MPKVLSAVACNLDANILAACLPLMEESRIEAIEWSFDALYKVKEVPAWFQELLTAFSDENRLIGHGVFFSLFSGKWLPEQENWLKHLSQTANAFQFDHITEHFGFMTGKDFHHGAPLNIPYTATTLNIGRDRLKRIYEACKRPVGLENLAFSYSLDEVKRHGAFLEQLLEPVNGFIILDLHNLYCQLRNFDVTFEEMIALYPLDRVREIHISGGSWEDSVATPSRTIRRDTHDDAVPAEVFQLLEMTIPRCPQLKYVVMEQLGNGLETAESKQYFYNDFLRMQEIIQQHSDQTASPVTDHFLPLLSFSTDTIVEDMTLYQQQLELSAIFESAASYDEAIQALSASSLANSDWRIEQWEPHMIETAIKISQKWMK |
| OIN58769.1 | Arsenicibacter rosenii | MHILQAGLPLFASEQVAALEWSFDSLYAQTELPDWFDELLLTYGQANRLVGHGVFFSIFAGRWRPEQQTWLNQLADRANRYRFDHVTEHFGFMTGASFHAGAPMSVPLNTTTLRIGQDRLHRIQQAAGCAVGLENLALAYSLDDVKRQGDFLHQLLAPVDGLLIMDLHNLYCQSHNFGVDWRELLALYPLDRIREIHLSGGSWEDSISEPGRIIRRDTHDEAVPGLLFDWLPVAIDQCPNLKYVVLEQLGRGLVTDAARLQYQADFTRIANSVAAHPKASVQGPAPAFRPETPVELPAQPLEDETLAWQQQHLTTILETAGSAHEARQQLAASPLAHSDWGTERWQPAMLETALRIAQKWTNGFSGLQ |
| SEW45833.1 | Chitinophaga sp. YR573 | MEESRVEAIEWSFDALYKEKEVPSWFRELLTAFSNEKRLIGHGVFFSLFSGKWLPEQEAWLKQLEQTAAEFRFDHITEHFGFMTGKDFHHGAPLNIPYTTTTLNIGRDRLKRIYEACRRPVGLENLAFSYSLDEVKRHGTFLEQLLEPVNGFIILDLHNLYCQLRNFDIAFDEMIALYPLDRVQEIHISGGSWDDSAADPNRSIRRDTHDDSVPPEVFQLLEMTIAQCPHLKYVVLEQLGNGLVTEMSKRCFYNDFLQMEEIVRKNNNENDAGASDPFLPLFPLSTGTAVEDMMLYQQQLELSSILESSASYAEAMHVLSQSSLANSDWRIEQWEPYMIETAVKIAQKWKK |
| WP_073174028.1 | Chryseobacteriu m vrystaatense | MKKPLLGLSMMPEADFVSAIVPLLQSNSVEVLEWSFDTFYDAEEPEWLSGLLNFYAGNERLIGHGVYYSLFDARWNDRQENWLQKLKEECKCRKYNHITEHFGFMNTENFHQGVPLPVPLLPKTLQIGKDRLRRLQDAVEIPVGVENLAFSFSIDDVREQGHFLDMLVEDIDGFLILDLHNIYCQACNFQIDMQEIINLYPLGKVREIHLSGGSWQESAYGRRLIRRDTHDDSIPDEILKILPKVLSQCYNLEYIIIERLGNTLNNDTEKQIFFSDFNHVKEIIEVSVYPDIIRQDWKRTEKIYSEPVEDLLLHEEQTRLTKLLFDGNNVESVKNGKFLYFKPENWDEEMIMTAQQIIRKWNPY |
| WP_291042127.1 | Dyadobacter sp. 50-39 | MSEIYSSIACNLDAHILQAALPLFEQEKVEAIEWSFDTLFKFDEIPAWFTDLVGEFSNHDRLIGHGVYFSLFSGKWSREQQEWLDKLRKLSGEFRFAHITEHFGFMTGEDFHKGAPISIPFTSRTLALGKDRLERIQDACQCPVGLENLAFSYSIDEVKKHGDFLGQLVGSVNGFIILDLHNVYCQTHNFNIDFDDIMRLYPLDRVREIHISGGSWDDILAAPGRPVRRDTHDERVPEAVFDFLSKAIPKCPNIRFVVLEQMGTALETPERQAAFQSDFLRMDGIVRASASRSAHAPAYAFAPAGTLPNDMPLEDPDLYNQQVQLSRILETASDAYEARALLLASDLRNTAWQVENWAPYMLETAVSIARKWKDGFIWK |
| WP_073335882.1 | Chryseobacteriu m carnipullorum | MSKPFLGLSMMPEADFVSAILPLLQNNSVDVLEWSFDTFYDVNEPEWLSGLLDFYAANNRLLGHGVYYSLFDARWNERQEIWLKKLKKECANRKYNHITEHFGFMNTENFHQGVPLPVPMHPLTLQIGKDRLKRLQDAVEIPVGVENLAFSFSIDDVKEQGVFLDQLVEDIEGFLILDLHNIYCQSCNFGVSMQEIIKLYPLEKVKEIHLSGGSWQESAYAKKPVRRDTHDDRIPDEIVEMLPEVLSQCQNLEYIVIERLGHTLSAESDKLFFFADFNRVKEIISKSEDRSEDKKVWNRRIGVDSKPVEDMVLYDEQTKLTKMLFDESSVAKVKNEAFQYFKPENWDEEMIMTAQQIIKKWNPY |
| OJY27962.1 | Myxococcales bacterium 68-20 | MTLAPELVSRDHRGRMHPWIGLSLMLEDDFRLAAGPLFAEGVVDVLEWSFDTGWGRAIPEWADALLDHYGEAGRLLGHGVHYSAFSARWEERQARWLERLAGEVARRTYVHVSEHYGFMTAAPFMRGAPLPVPRGEASRSVGRDRLERMRAVLAAGRNDACPIGLENLALAWNRDEALAHGAFLGEVLDHAEDFIVLDVHNLHCQIENFDIDPDELLDSFPAARIRELHVSGGSFLPAWPHHPEETVRCDTHDGDVPAPVFDLLERALDRFPDVRAVILERLGGTLRTQADIDTFRRDYLRVKQIVDHSRDGEGARDGDG |
| WP_072923384.1 | Chryseobacteriu m sp. OV279 | MKKPLLGLAMMPEADFVSAVLPLLQTQSVDVLEWSFDTFYDAEELEWLSGLLDLYAENNRLLGHGVYYSLFDARWADRQEVWLKKLKEEVKRRRYNHITEHFGFMNTENFHQGVPLPVPLLPKTLQIGKDRLSRLQDAVEIPVGVENLAFSFSMDDVKEQGEFLDKLVEDIDGFLILDLHNIYCQSCNFEVDMREIINLYPLEKVKEIHLSGGSWQDSAYGKKPVRRDTHDDRIPEEILNLLPEVLIQCHPEYVIIERLGHTLHTEAEKQIFFDDFNRVKKIIELSDYPVGEEKIWNRKKTVHSQPVEDLLLYAEQTELTRLLFDGNTTESIKNRDFHYFKPESWDEEMITTAQQIIKKWNPY |
| WP_073852115.1 | Pontibacter flavimaris | MPGILATVACNLDADILSATYPLLEGGSVEAMEWSFDTLFQVEQLPDWFREMLQAYSQEGRLIGHGVFYSLISGRWTEKQAQWLKNLRQLSAAFHFDHITEHFGFFTGQNFHYGAPLSIPYTSTTLRIGRDRLLRIQDACQRPVGLENLAFSYSLDEVKQHGEFLEKLIEPANGFIILDLHNLYCQLHNFSVSYEELILLYPLERVREIHISGGSWEEAQEAPGHKVRRDTHDDAVPAEVFQLLKQTMPLCPNLKYVVLEQLGNGLKTESSRAGFQRDFRQMELLVQQVNAARTGLLPDNLFLPLSAFPDSFVEEDEALHQQQVLLSQILENAPSYQEAQRLLQTSSLAHSDWRIENWQPYMLETAIRIAQKWKK |
| SIO13075.1 | Chitinophaga niabensis | MQESRVEALEWSFDALYKVKDIPSWFIELLTAYSDEKRLIGHGVFFSLFSGKWLKEQENWLKQLKQTCTEFQFDHITEHFGFMTGKDFHHGAPLNIPYTSTTLNIGRDRLKRIYDVCRCPVGLENLAFSYSLDEVKRHGDFLEQLLSPVNGFIILDLHNLYCQIHNFNISFDDIIALYPLDRVREIHISGGSWEGSLVDPPRQIRRDTHDDAVPEEVFHLLENTIERCPNLKYIVLEQLGNGLSTDESKTTFYQDFLKMESIAGRFSSQGKQTFVPLSPWQTGYPVEDETLYQQQRELSSILESAASYETAVNALSVSSLAQSDWGIEHWEPYMIETAIKIAQKWKK |
| WP_074240080.1 | Chitinophaga niabensis | MPKILSAVACNLDNDILAACLPLMQESRVEALEWSFDALYKVKDIPSWFIELLTAYSDEKRLIGHGVFFSLFSGKWLKEQENWLKQLKQTCTEFQFDHITEHFGFMTGKDFHHGAPLNIPYTSTTLNIGRDRLKRIYDVCRCPVGLENLAFSYSLDEVKRHGDFLEQLLSPVNGFIILDLHNLYCQIHNFNISFDDIIALYPLDRVREIHISGGSWEGSLVDPPRQIRRDTHDDAVPEEVFHLLENTIERCPNLKYIVLEQLGNGLSTDESKTTFYQDFLKMESIAGRFSSQGKQTFVPLSPWQTGYPVEDETLYQQQRELSSILESAASYETAVNALSVSSLAQSDWGIEHWEPYMIETAIKIAQKWKK |
| WP_076552343.1 | Chryseobacteriu m ureilyticum | MKKPLLGISMMAESEFVSAILPLLQSSSIEVLEWSFDTFYNAKEPVWLSDLLNFYSENSRLIGHGVYYSLFDAKWNDRQKIWLEKLKKEVHKRQYNHLTEHFGFMNTENFHQGVPLPVSLHSQTLQIGKDRLHRLQDAVDISVGIENLAFSFSVDDVKEQGLFLDQLIEDIDGFLILDLHNIYCQACNFEVDMQEIIQSYPLDKVKEIHLSGGSWQESIYGKKSIRRDTHDDDIPDEIWAVLPSVLLQCKNVQYVIIERLGHSIKTERDKQNFLSDFIKAQTIINDLDWSGRENRFWSKKEVNISKKPLEDIVLYEEQTRLTKLLFDNVEPSLIKKENFQYFNTKIWNEEMIMTAQNILKKWNPY |
| OJV53697.1 | Bacteroidetes bacterium 43-16 | MEQMIEGIIPSIACNLDADMLSASLPLFEAGVVGGLEWSFDALHARNYQVPDWFHELLLAYSDAERLVGHGVFFSLFSGRFSEAQQAWLKRLSALTKVYQFDHITEHFGFMTGQDFHKGAPLSVPYTPSTLAIGRDRISRIADAAQCPVGLENLAFAYDLEEVKIQGRFLDELLTPVNGFIILDLHNLYCQLHNFNLDIATLLALYPLDKVREIHISGGSWESHPQAPFMKVRRDTHDDAVPAAVFELLRMALAQCPNLKFVVLEQLGIALSSPAQRVQFQQDFNTMKTIIEQYNRQQQWPRDIAAFRADQIQLAPAALSDEALFAQQQIISHILETASDHKDVIDKMQHTSLAGSDWHIENWHPAMVDTAIHIAQKWK |
| WP_170872041.1 | Chryseobacteriu m sp. RU33C | MLGVSMMAEADFVSAVLPLLQNNSIDVMEWSFDTLYHANEPDWLRDLLNFYAENNRLIGHGVYYSLFDARWTERQEEWLKKLKEEVNLRKYNHITEHFGFMNTENFHQGVPLPVSLHPKTLQIGKDRLYRLQEAVNIPVGIENLAFSFSIDDVKEQGVFLDKLTEDTNGFLILDLHNIYCQSCNFELEMQEIIDLYPLDKVKEIHLSGGSWQDSVYGKKQVRRDTHDDVIPRDILSVLPSVLTQCENLEYIIIERLGHTLKTDEERDHFLDDFSKVKTIIETSELNGKEKSNWSKMKVELSEKPLEDPVLYEEQSRLTQLLYDNADVETIKDLDFYYFKTENWDSEMILTAQNIIKKWNPY |
| WP_084180480.1 | Chryseobacteriu m joostei | MRKPLLGLSMMAETEFVSAILPLLQSNSVDVLEWSFDTLYHTKEPDWLGDLLNFFSENNRLIGHGVYYSLFDARWTERQNIWLNKLKKEVKKRKYNHITEHFGFMNTENFHQGVPLPVSLHSKTLQIGKDRIFRLQEALDIPVGIENLAFSFSIDDVREQGLFLDKLIEDTDGFLILDLHNVYCQSCNFEVEMREIINSYPLDKVKEIHLSGGSWQNSVYGEKLIRRDTHDDVIPEEILSVLPWVFSQCQNLEYVIIERLGHSIHLESEKDDFLKDFAKVKKLISESDWRDQEKNNWNKKVLKLSESPLEDTLLHKEQTMLTRLLFDSADVSLIKNQEFHYFKTDTWDTEMILTAQNIIKKWNPY |
| SIS50444.1 | Chryseobacteriu m shigense | MMPEADFVSAILPLLQNNSVDVLEWSFDTFYDVNEPEWLSGLLDFYAQNSRLLGHGVYYSLFDARWTARQEVWLEKLKKECSQRKYNHITEHFGFMNTENFHQGVPLPVPMHSKTLQIGKDRLKRLQDAVEVPVGVENLAFSFSMDDVKEQGVFLDQLVEDINGFLILDLHNIYCQSCNFEVGMQEIIQLYPLEKVKEIHLSGGSWQESAYAKKPIRRDTHDDRIPDEILDLLPEVLSQCQNLEYVIIERLGHTLSAESDKLSFFADFNRVKEMISNSEDRSEGKKVWKRRIEEDSKPLEDLVFYNEQTRLTKMLFDESDVAKVKDEGFQYFKPENWDEEMIITAQQIIKKWNPY |
| WP_076510010.1 | Chryseobacteriu m shigense | MRRPFLGLAMMPEADFVSAILPLLQNNSVDVLEWSFDTFYDVNEPEWLSGLLDFYAQNSRLLGHGVYYSLFDARWTARQEVWLEKLKKECSQRKYNHITEHFGFMNTENFHQGVPLPVPMHSKTLQIGKDRLKRLQDAVEVPVGVENLAFSFSMDDVKEQGVFLDQLVEDINGFLILDLHNIYCQSCNFEVGMQEIIQLYPLEKVKEIHLSGGSWQESAYAKKPIRRDTHDDRIPDEILDLLPEVLSQCQNLEYVIIERLGHTLSAESDKLSFFADFNRVKEMISNSEDRSEGKKVWKRRIEEDSKPLEDLVFYNEQTRLTKMLFDESDVAKVKDEGFQYFKPENWDEEMIITAQQIIKKWNPY |
| OMP75747.1 | Flexibacter sp. ATCC 35208 | MDANILAACLPLLQESRIEAIEWSFDALYKIKEVPDWFRELLTAFSDENRLIGHGVFFSLFSGKWLPEQEAWLSHLKQTSTAFNFDHITEHFGFMTGKDFHHGAPLNIPYSASTLAIGKDRLKRIYNACGRPVGLENLAFSYSLDEVKRHGDFLDQLLEPVNGFIILDLHNLYCQLHNFDLAFEELIALYPLDKVREIHISGGSWDDVGNRTIRRDTHDEGVPAEVFQLLEMTMPKCPHLKYVVLEQLGNGLVTEASRLGFYNDFLKMQEIVHKSSNEIEANLFQMQEIVPRSNHDTGGNLLQTQEIVHKSNRETEVNPFHPLLPLPIGPAVEDLHLYEQQMELSSILETAPSNAAAMQLLQHSSLAHSDWKIEQWDPYMIETAVKIARKWKK |
| WP_077414519.1 | unclassified Chryseobacteriu m | MKKPLLGISMMAEADFVSAVLPLLQNNTIDVLEWSFDTLYHANEPDWLRDLLNFYAENNRLIGHGVYYSLFDARWTERQEDWLKKLKEEVGLRKYNHITEHFGFMNTENFHQGVPLPVSLHPKTLQIGKDRLYRLQETIDIPVGIENLAFSFSIDDVKEQGIFLDKLTEDTNGFLILDLHNIYCQSCNFGVKMQEIIDLYPLDKVKEIHLSGGSWQESVYGKKQVRRDTHDDIIPEEIFSVLPSVLAKCQNLEYIITERLGHTLKTEEEKENFLNDFNKIKTIIEASDWKGRKISSWNKRQIQLPEKPVEDLALFEEQSRLTRLLFDNAGAAVIKDQDFHYFKTENWDPEMILTAQNIIKKWNPY |
| WP_235015951.1 | Chitinophaga ginsengisegetis | MPEILSAAACNLDANILAACLPLMEESRLEAIEWSFDALFKVQQIPSWFFELLTAYSNEQRLIGHGVFFSLFSGKWRQEQADWLKQLKKVREEFRFDHVTEHFGFMTGRDFHHGAPLNIPYTKTTLDIGRDRLKRIYEACECPVGLENLAFSYSLEEVKRHGGFLEQLLEPVNGFIILDLHNLYCQIHNFNIPFEEIIRHYPLDKVREIHISGGSWENAPNSPLKKIRRDTHDDAVPAEVFHLLENTIARCPHLKYVVLEQLGNALTTEESRTIFYNDFGKMENIVKKKNTPGISSYQSFLPSSPVITGKIIEDEILHHQQLELSAILESATSCGEAIQSLTRSSLAHTAWQIENWEPYMINTAIQIAQKWKK |
| WP_079241387.1 | Chryseobacteriu m | MRKPMLGVSMMAEADFVSAILPLLQNNSIKVMEWSFDTLYHTNEPDWLRDLLNFYAENDRLIGHGVYYSLFDARWTERQEEWLKKLTEEVSLRKYNHITEHFGFMNTENFHQGVPLPVSLHSKTLQIGKDRLCRLQDAVNIPVGVENLAFSFSIDDVKEQGVFLDKLTEDTNGFLILDLHNIYCQSCNFELGMQEIIDLYPLDKVKEIHLSGGSWQDSVYGKKQVRRDTHDDIIPQDILSVLPSVLTQCGNLEYVIIERLGHTMKTEQERDDFLNDFNQVKAIIETSDFNGKEKSSWNKMEVEHSEKPLEDLVLYEEQSRLTRFLYDNSNVETIKDLDFHYFKTEKWDSEMILTAKNIIKKWNPY |
| WP_078013439.1 | Hymenobacter sp. CRA2 | MHSEQAPIYSAVACNLDANILGAAFPLLEDGRIEALEWAFDTLFWAEQIPDWFTNLLDAYSQAGRLVGHGVFFSLLSGRWTPEQQQWLAHLRQVAARFGFDHITEHFGFFTGQNFHHGAPLPVPYSRTALRLGHDRLGRIAEACRCPVGLENLAFAYSLDEVKRHGTFLDELLAPVNGFVILDLHNLYCQLHNFAVPYDELLALQPLARVREIHISGGSWEDSALVPGRRIRRDTHDEAVPEEVFELLAWTLPRCPNLKYVVLEQLGSGLTTEQSRAQFRHEFERMAALVEQHRRARPATNAFRPPLPGLSSPAAEDAALHEQQRQLSHILETATSCEQAREQLQASALAHTDWRIEQWEPHMLETAVRIAQKWKE |
| WP_077921573.1 | Spirosoma sp. 209 | MTDIRSSVACNLDSHILRAALPLFADEKVGAIEWSFDALVNPRSMPDWFAELLRTYSGAGRLLGHGVFYSLFSGRWTGAQQQWLDQLKSLSRTYRFDHVTEHFGFMTGADFHKGAPMSISFTESTLAIGRDRLLRLQDACHCPVGLENLAFAYTVDEVDRHGAFLNQLIEPVNGFIILDLHNLYCQSRNFGVDPDGLLSLYPLYRVREIHLSG |

| | | |
|---|---|---|
| | | GSWAPSAVEPGRTIRRDTHDDAVPADVFRLLKQAIDLCPNLQYVVLEQLGAGLQTDGQQSQFQQDFLWMDHIVQAANQIRNPLINHSFLPRRFHVPDPSPLEDPLLYAQQLELSDILETSVSYAQAQQRLAGSGLARSDWAVETWQPAMLETALAIAQTWKNGFD |
| WP_080778923.1 | Chryseobacterium phocaeense | MKKPLLGLSMMPEPDFVSAILPILQDHSVDVLEWSFDTLYDLEESEWLSGLLDFYAENNRLLGHGVYYSLFDARWTERQEIWLEKLKKEVQHRNYNHITEHFGFMNTENFHQGVPLPVPLHPKTLQIGKDRLLRLQDAVEIPVGIENLAFSFSVNDVKEQGEFLDKLIEDIDGFLILDLHNIYCQSCNFETDMLEIIRMYPLEKVKEIHLSGGSWQESAYGKKPVRRDTHDERIPEDILNILPEVLSLCPNLDYIIIERLGHTLKTEAEQRIFFDDFERVKKIISKAGYPSGQNKIWRKKKIREHSEPVEDHLLYDEQTKLTQLLFNGTDTHSIKDQDFYYFKPENWDEEMIITAQKIIKKWNPY |
| WP_235021434.1 | Chitinophaga eiseniae | MPELISTVACNLDNNILAACLPLMEAARVEAIEWSFDALYAVENVPAWFEELLTAFSQENRLIGHGVFFSLFSGRWLPEQQQWLDHLKTVSRHYRFDHVTEHFGFMTGKDFHHGAPLNIPYTPVTLNIGRDRLKRMQDACECPVGLENLAFSYSLEEVKKHGDFLNELLEPVNGFIILDLHNLYCQAHNFDLSFDALKALYPLHRVREIHISGGSWHESEAAPGKTVRRDTHDDAVPEVVFNYLDKAIDLCPSLKYVVLEQLGTGLATEESRSAFGNDFLRMDQLLKNKNTNTAPLNTFLPQALRIPAQAREDETLYRQQLELSQILETAASFEATMKALQASSLAHSAWQIENWDPAMIETAAKIAQKWKRQTPDRRIT |
| SMC33782.1 | Chryseobacterium sp. YR221 | MMAEADFVSAILPLLQDNAVDVVEWSFDTFFAAEEPSWLKDLLNFYAGNGRLLGHGVYYSLFDARWTERQEMWLQKLKHEVQKRNYNHITEHFGFMNTENYHRGVPLPLSLHAKTLEIGKDRLYRLQDAVPIPIGVENLAFSFSVDDVREQGVFLDKLVDDIDGFLILDLHNIYCQSCNFDVSIQEIVSLYPLEKVKEIHLSGGSWQDSVYGKKMIRRDTHDDVIPEEIFSVLPSVLSQCHNLEYVIIERLGYTIHTEEEKQRFFNDFMRVKSILNDSDGMKRKKEKWMKREMLLPGPVEDSILYEEQILLTKLLFDHIDPTLIKNQNFHYFKTENWDPEMILTAQNIIKKWNPY |
| WP_081964927.1 | Legionella norrlandica | MDLPKLGLSLMPEQDYMDASRPLFESGDVDIIEWSFDMCWGRKIPQPCLDLLTKYSSQGCLLGHGVSFSILTAKRTKRQELWLQYLQNEVNQLNYQRISEHFGFLSTNTFIQGPPLSMPFIPEMLEVGIKNLQQIKQIAKIPIGLENLGFAFCKQDVIEQGKFLTALLEAVDGFLLLDIHNIYCHMINFDMSLDEILDSLPLDRAKEMHISGGSFRSIEQNNRNIYCDSHDAAIPQDLFLLLEKNLSRFSQIDSLIFERLGNTLSNHYEAAQFREDFMTLKSLIRR |
| WP_082215810.1 | Dyadobacter psychrophilus | MPEIYSSIACNLDAQILQASLPLFEQEKVEAIEWSFDTLFKFNEIPVWFTDLVSEFSDHSRLIGHGVYFSLFSGKWTSGQQEWLDKLQGLSLQFQFDHITEHFGFMTGEDFHKGAPISVPFTPQTLALGKDRLQRIQDACKCPVGLENLAFSYSMDEVKKHGDFLAQLVESVNGFIILDLHNIYCQVHNFNIDFDEIIKLYPLGRVREMHISGGSWDETLVGPGRQVRRDTHDEAVPEAVFELLQKAIPKCPNLKYVVMEQMGTALDTAERQAVFQSDFLRMDNIIKEIKTDFDPKITNTFGLLNRSAGTIPLEDQELHRQQLQLSHILETAGSVSEAQAMLRSSDLHDTAWEVDKWAPYMLQTAISIAQKWKNGFA |
| WP_082007927.1 | Cellulophaga baltica | MRKRIMGNKKPKLGIAIAPSLKFLEAALPLFAGEKIEIIEWSFDTLKDAADESAWLSLLLKEYGEKNRLLGHGVFYSLLDANWSSRQENWLKKVRQETLSHKYCQISEHFGLMSSANAHSGFPLPIQLSNPVLQIGIDRLKRLQATAQVDVGIENLALTANVADILEQGEFLFKLVNPVNGFVILDLHNIYCQSENFNIDMMTIIKSYPLSLVKELHISGGSWDTDSTLSRKIRRDTHDGRIPEVILEILPEVLKICPFLAFIIFEKIEDSFLTEKDGIDFRADFQKIREIIDATAFSVTPKEKKQVAAILGPPVIDVELLQAQVALRESIALGTYHTNTKWDKDMWKVATKLYEKWNN |
| WP_082474654.1 | Chryseobacterium sp. Leaf201 | MKRQLLLGVSMMAEADFVSAILPLLESSGIDVLEWSFDTFYTVDEPQWLADLLNFYSDHKRLLGHGVYYSLFDAKWTERQDVWLQKLKAETDRRSYNHITEHFGFMNTENFHQGVPLPVPLHPELLKIGKDRLLRLQDVVEVPIGIENLAFSFSSDDVSEQGEFLYRLIEDIDGFVILDLHNIYCQASNFNMEIMELVNLYPLEKVKEIHLSGGSWENSIYTKNKRIRRDTHDDEIPQELFDILPDVISRCSHLEYIIIERLGHTIKTESDKQNFLRDFKKVKQIIDNSDPKENLQHQWKKKNRVLTDPMENLLLFDDQKNLNKFLFESQDPETIRNYAFHYFKMHNWDPEMIHTASQIVKKWNPY |
| WP_081145449.1 | Niastella vici | MTKLLATAACNLDADMLNACYPLFLEEKVQAIEWSFDALFDHGALPQWFGELLQAYSQAKRLTGHGVFFSLFSGKWLPEQVQWLASLRDTIQRFPLEQVTEHFGFMTGKDFHEGAPLAIPYTATTLRIGRDRLRRIQDAAQCPVGLENLAFAYTPDEVKRHGDFLYKLIEPVNGFIILDLHNVYCQLKNFDLDANELLSLYPLHKVREVHISGGSWERVESEPGRQIRRDTHDDVVPAEVFLLLDKALDRCPECKYVVLEQLGNGLKSHTSRQQFYHDFVRMDNILQQKNWFRKGESQNDFMPLQLFDNGPAPEDELLFNQQRELSQILESGMSYSEAMQVLNRSSLANTPWDIEHWDPAMLETAMQIARKWKKQV |
| WP_084084025.1 | Chryseobacterium sp. YR221 | MGKPLLGLSMMAEADFVSAILPLLQDNAVDVVEWSFDTFFAAEEPSWLKDLLNFYAGNGRLLGHGVYYSLFDARWTERQEMWLQKLKHEVQKRNYNHITEHFGFMNTENYHRGVPLPLSLHAKTLEIGKDRLYRLQDAVPIPIGVENLAFSFSVDDVREQGVFLDKLVDDIDGFLILDLHNIYCQSCNFDVSIQEIVSLYPLEKVKEIHLSGGSWQDSVYGKKMIRRDTHDDVIPEEIFSVLPSVLSQCHNLEYVIIERLGYTIHTEEEKQRFFNDFMRVKSILNDSDGMKRKKEKWMKREMLLPGPVEDSILYEEQILLTKLLFDHIDPTLIKNQNFHYFKTENWDPEMILTAQNIIKKWNPY |
| ASE60851.1 | Chryseobacterium indologenes | MMRKPLLGLSMMPEAEFVSAALPLLQNNSVEVLEWSFDTLFNTNEPDWLRDLLNFYAENNRLIGHGVYYSLFDAQWTDRQEIWLSRLKDEVSKRNYNHITEHFGFMNTENFHQGVPLPVPLHPITLQIGQDRLQRLQDAVNIPVGIENLAFSFCLDDVKEQGVFLEKLIESSGGFLILDLHNMYCQSCNFDVDIQEIIKLYPLQKVKEIHLSGGSWQESVYGKKMIRRDTHDHIIPEEIFAILPFVLSHCENLEYIIIERLGPTIKTKEEEDDFFSDFMRVKTMIDSSAGHERQGDLWIKKEFELSKYPVEDPVLYEEQSRLTKLLFDGTHVDSIKNENFHYFNTAGWDPEMIFTAQEIIKKWNPY |
| OUJ69468.1 | Hymenobacter crusticola | MEQEPTMLASIACNLDADILSAAFPLLEEGQVEALEWSFDTLFWAEQVPSWFTELLTTYSNHNRLVGHGMFFSLLSGKWTLEQQQWLRQLHQLAAQFRFDHITEHFGFFTGQNFHYGAPLHVPYCEATLRIGQDRLRRIYEACQCPVGLENLAFSYSLEEVKQHGAFLEKLIKPVNGFLILDLHNVYCQLHNFSVAFEELVMLYPTDRVREIHISGGSWEGSVSAPGGRVRRDTHDEAVPEEVFQLLDATLPRCPNVKYVVLEQLGNGLKTEPSKIQFRQDFLRMQALVAKHRDHTSRPLTNTFLPLHPLPPSSIIEDEELYKQQLQLSHILETASSSDEAQRLLHASTLAKTAWKIEQWEPHMLQTAVDIAQKWKK |
| WP_088841642.1 | Hymenobacter gelipurpurascens | MRTVAPPDNANERAIYSSIACNLDANILTAAFPLLEQGKVEALEWSFDALFWAEQVPDWFTELLQAYSAQHRLVGHGVYFSLLSGRWTAEQQQWLQHLKELTRRFSFDHVTEHFGFFTGQNFHAGAPLPIPYGSNTLRIGQDRLRRMQEACGCPVGLENLAFAYSLEEVKRHGEFLEQLVEPVNGFLILDLHNVFCQLHNFSVPYEELMALYPLDRVREIHISGGSWEASGQVPGLQIRRDTHDEAVPEEVFQLLYNTMPRCPNLKYVVLEQLGTGLQTEDSRAHFQSDFHRMQALVNQHRSTTSIRAVQPFLPIHSAVAGPVAEDAQLHEQQQQLSHILETAPSYEDAQRQLQASALAYTAWKVEEWEPHMLETAISIAQKWKPQPR |
| WP_083729470.1 | Flexibacter sp. ATCC 35208 | MPKVLSAVACNLDANILAACLPLLQESRIEAIEWSFDALYKIKEVPDWFRELLTAFSDENRLIGHGVFFSLFSGKWLPEQEAWLSHLKQTSTAFNFDHITEHFGFMTGKDFHHGAPLNIPYSASTLAIGKDRLKRIYNACGRPVGLENLAFSYSLDEVKRHGDFLDQLLEPVNGFIILDLHNLYCQLHNFDLAFEELIALYPLDKVREIHISGGSWDDVGNRTIRRDTHDEGVPAEVFQLLEMTMPKCPHLKYVVLEQLGNGLVTEASRLGFYNDFLKMQEIVHKSSNEIEANLFQMQEIVPRSNHDTGGNLLQTQEIVHKSNRETEVNPFHPLLPLPIGPAVEDLHLYEQQMELSSILETAPSNAAAMQLLQHSSLAHSDWKIEQWDPYMIETAVKIARKWKK |
| WP_083421972.1 | Arsenicibacter rosenii | MAIYPALACNLDMHILQAGLPLFASEQVAALEWSFDSLYAQTELPDWFDELLLTYGQANRLVGHGVFFSIFAGRWRPEQQTWLNQLADRANRYRFDHVTEHFGFMTGASFHAGAPMSVPLNTTTLRIGQDRLHRIQQAAGCAVGLENLALAYSLDDVKRQGDFLHQLLAPVDGLLIMDLHNLYCQSHNFGVDWRELLALYPLDRIREIHLSGGSWEDSISEPGRIIRRDTHDEAVPGLLFDWLPVAIDQCPNLKYVVLEQLGRGLVTDAARLQYQADFTRIANSVAAHPKASVQGPAPAFRPETPVELPAQPLEDETLAWQQQHLTTILETAGSAHEARQQLAASPLAHSDWGTERWQPAMLETALRIAQKWTNGFSGLQ |
| WP_090024778.1 | Chryseobacterium oleae | MKKPLLGLAMMPEAEFVSAILPLLQTQSVDVLEWSFDTFYDAEEPEWLSGLLDFYAENNRLLGHGVYYSLFDGRWANRQEIWLKKLKEEVKRRKYNHITEHFGFMNTENFHQGVPLPVPLLPKTLQIGKDRLSRLQDAVEIPVGVENLAFSFSMDDVKEQGEFLDKLVEDIDGFLILDLHNIYCQSCNFEVDMMEIINLYPLEKVKEIHLSGGSWQESAYGKKPVRRDTHDDRIPEEILNLLPEVLILCHPEYVIIERLGHTLNTEAEKQIFFDDFNRVKKIIELSDYPVDEEKIWNRQNTVHSKPVEDLLLHAEQTELTRMLFDGNTTESIKNRNFHYFKPESWDEEMITTAQQIIKKWNPY |
| WP_097124551.1 | Spirosoma fluviale | MSRLYSSIACNLDTNSLQAALPLFESEKVQAIEWSFDTLFALGEMPDWFAELLRAYSQEGRLIGHGVFFSLFSGKWTSDQQQWLNQLKITSADFTFDHLTEHFGFMTGEDFHKGAPVSIPFTASTLAIGRDRLQRLQDAGSCPVGLENLAVAYSLDDVKRQGDFLAQLLEPINGFIILDLHNLYCQSYNFAIEITDLQALYPLDRVREIHISGGSWVPSAVNPTKQIRRDTHDDSVPAAVFHALERVMGQCPNLKYVVLEQLGTGLTTEASRRHFQEDFLRMDALVQAAHRADSPPMLNSFLPLNPPSIPEIAPENPLLYKQQTQLSAILETATDCAQAQLFLKASSLANSDWEIENWHPEMLETAIAIARKWKDGFVQ |
| OYV03992.1 | Verrucomicrobiales bacterium VVV1 | MWKGLIDMIRSGLSLMLEDEFLQAALPLFECGSVEALEWSFDVGWSLPSLPEWVTALLREYGEKGALVGHGVTFSALSGEWGDRAQNWSDQLREELNQRKYLHLSEHFGFMAGGNFHQGTPLPVPLTEKTLRLGQSRLKLLSDVAGAPVGLENLALALSPRDVADQGRFLGELLAPSDGFLVLDLHNIYCQMMNFDCTADSILASYPLERVREMHISGGSWMEAVTARGTPIRRDTHDEAVPETVFQMLDAVLPRCPGTQVVIFERLGHTLGGEGESETFRDDFLRMKSIVSSHAG |
| WP_099766995.1 | Chryseobacterium sp. 52 | MKKPLLGLSMMPEADFVSAILPILQNNAVEVLEWSFDTFYDLEEPEWLSGLLDFYAENNRLLGHGVYYSLFDARWTVRQDEWLKKLKKEVNQRKYNHITEHFGFMNTENFHQGVPLPVPLVSKTLQIGKDRLRRLQDAVEIPVGVENLAFSFSVDDVKEQGEFLNRLVEDIDGFLILDLHNIYCQSCNFEIDMQEIIRWYPLEKVKEIHLSGGSWQESAYGKKPVRRDTHDDRIPEEILNILPGVLSQCHHLEYTIIERLGHTLKTEEEKQTFLDDFKQVKEIIETSECFTGEEKKWNKTEEVYSEPVEDHLLYDEQTKLTKLLFDSNGVASVKEQEFHYFKPENWDEEMITTAQQIIRKWNPY |
| WP_101237823.1 | Chryseobacterium sp. PMSZPI | MKKTLLGLSMMAEADFVSAILPLLETHSVDVLEWSFDTFYDVEEPDWLSGLLNFYAENNGLIGHGVYYSLFDARWTDRQEAWLEKLKKEIDKRRYKHITEHFGFMNTENFHQGVPLPVSLNPQTLQIGKDRLYRLQDILQIPVGVENLAFSFCTDDVKEQGEFLDKLVEDIDGFLILDLHNIYCQSCNFGVDMENIIRLYPLEKVKEIHLSGGSWQESAYGKSPIRRDTHDDAIPKEILNILPIVFSKCENLEYVIIERLGNTIKTDKEKKSFFDDFVQVKTILESSSRKSRKKSSWAKKDLEFSEVPLEDLILYEEQTMLTKLLFDNTDVTSIKNYDFKYFKTEKWDPEMILTAQNIIKKWNPY |
| ATN04960.1 | Chryseobacterium indologenes | MMRKPLLGLSMMPEAEFVSAALPLLQNNSVEVLEWSFDTLFNTNEPDWLRDLLNFYAENNRLIGHGVYYSLFDAQWTERQEIWLSRLKDEVSKRNYNHITEHFGFMNTENFHQGVPLPVPLHPITLQIGQDRLQRLQDAVNIPVGIENLAFSFCLDDVKEQGVFLEKLIDNSGGFLILDLHNMYCQSCNFDVDIQEIIKLYPLQKVKEIHLSGGSWQESVYGKKMIRRDTHDHTIPEEIFAILPFVLSHCENLEYIIIERLGHTIKTKEEEGDFFSDFMRVKTIIDSSAGHERQGDLWIKKEFELSKYPVEDPVLYEEQSRLTKLLFDGTHVDSIKNENFHYFNTAGWDPEMIFTAQEIIKKWNPY |
| PWN62849.1 | Chryseobacterium viscerum | MPGLRNGIKMRKPLLGVSMMAEADFVSAILPLLQNNSIEVLEWSFDTLYHTHEPDWLCDLLNFYAENNRLIGHGVYYSLLDARWTARQEQWLEKLKEEVQQRKYNHITEHFGFMNTDNFHQGVPLPVSLHPKTLQIGKDRLNRLQEAVDIPVGIENLAFSLSVDDVKEQGMFLDKLTEDSNGFLILDLHNIYCQSCNFGVEMQEIINLYPLDKVKEIHLSGGSWQESVYGKMPVRRDTHDDVIPEDIFSVLPAVLTQCQSLEYVIIERLGHTLKTEEEKSNFLHDFNKVKTIIEASDWKNRRENNWTKKEIQLAEKPVEDLVLYEEQSRLTRLLFDNVGAAAIKDQDFHYFKTENWDPEMILTAQNIIKKWNPY |
| WP_099149092.1 | Flavilitoribacter nigricans | MPKIYSSIACNLDNHLLSAALPLFAEESVEAIEWSFDTLYKQRNIPDWFIELLTAFGQENRLVGHGVYFSLFSGKWTPEQEAWLQQLEKVSGHFQFDHISEHFGFMTGANFHAGAPIGIPFTRRTLQLGQDRLQRIFQACRCPVGLENLAFAYSLDEVKAHGQFLDELLEPVNGFIILDLHNLYCQLQNFTIPFEALLELYPLHRVREIHISGGSWEDSALHPRQKIRRDTHDDAVPDEVFQMLSAVVPRCPHLKFVVLEQLGIGLHTPASQEAFREDFRKLKRTVQSLSTPPPETVANDFLPKELCPLGPAPEDLSLHRQQQQLSRILETAPDLSAAREELQSSDLAGSAWKVEEWEPAMLETAMQIAQKWKNGFA |
| AUC75689.1 | Olleya sp. Bg11-27 | MWYGIMKINKPKLGLSIMPNPEFIAAALPLFESAQVEVIEWSFDTILDKKYQPEWLPLLLKEYGDNNRLLGHGVYYAPLDANWGANQDNWLQKAKLETLAYNYNHLSEHFGVMSSANAHHGFPLPFDLSNTILNIGIDRLKRLQNTVQLDIGIENLALASNVSDILKQGAFIEKLVAPINGFVILDLHNIYCQSINFDLDLLTIITSYPLHLVKEIHLSGGSWDHDHNLTKPIRRDTHDGRIPEVLLDILPEVLQRCPVLEFVIFERLGDTFQNKNDGLEFRTDFNKIQAIIDHTAFSSNARQWTLKKHDLGPPLIDLPLLNKQQELREHIRLETANNHAEWDTNMWITATKLYKKWNA |
| WP_104421012.1 | Neolewinella xylanilytica | MTYPSLACNLDHQLLSTALPLLEAGTVSGLEWSYDAVYQYETLPEWFAALLKAFAAEGRLIGHGIFYSVCSGAWSEDQRNYLDRLAHTVEAFPFDHVTEHFGFLTGANFHRGAPIAPPLTQSTLRIGQDRLHRLHDAARCPVGLENLAFAYSADDVFRQYAFLDQLVQPIDGLLILDLHNLYCQLHNFSLDLEELLRHIPLERVREMHLSGGSWDFHHDAPTGRVRRDTHDERVPDEVFHLLHEIAPRCSNLKFVVLEQLGSALRETEAQLGFQADFDRMTSVCTAMVQQPVNDRPAPCLPNTSPITHPPLQDRALANEQQELSTILETSSTLAEVQDRLSNSSLANSAWQMENWSPYMLETAWKIARKWR |
| WP_170061848.1 | Hymenobacter chitinivorans | MSTPAAAAEPGILATLACNLDADMLSASFPLLEQGRVEALEWSFDALFWAEQIPEWFTELLRAFAAQNRLVGHGVYFSLLSGRWTPEQQQWLSQLRKLAQQFRFDHITEHFGFFTGQNFHAGAPLPIPYTSTALRLGQDRLCRIQEACGCPVGLENLAFAYSLDEVRRHGEFLAQLVEPVNGFLILDLHNLFCQLHNFGVSYEELIQLYPLERVREIHISGGSWEESGQEPGRQVRRDTHDGAVPDEVFELLSRTLPRCPNLKFVVLEQLGNALQTEPSRAQFRRDFGRLEAMVHEHRAQAPAGGSPLFLPQRPLPTGPVAEDARLHEQQQQLTHILETAPSYDEARQRLAGSSLAHSDWQIEQWAPHMLETALRIAQKWK |
| MAR09898.1 | Blastopirellula sp. | MLEADFAAATQPLFATGDVEVLEWSFDVGWGRSVPPWAEQLVSFYSQHQQLLGHGVSFSPLSAEWQTRQTDWLRQLGQEVARRDYQHVSEHFGFMTAGDFHQSAPLPVPRTPATLAIGRDRLARLADTCPCSVGLENLAFAFGQRDVASQGDFLDALLEPVDGFLLLDLHNIYCQSCNFDVDPWSLIESYPLDRVRELHVSGGSWTEARDGQAVRRDTHDGSVPALLFELLPRVLDRCPQVQYVILERLGGTFTNAEAEFQADFRRLRHTVETWGEWATSSSANSKHQP |
| WP_105041844.1 | Rubritalea profundi | MPHVGLSLMPETQFLQASMPLFEAGDVGAVEWSFDTGWKNGVSEWLTSLIGTYSEAGRLTGHGVHFSLLSADWTQRQQGWLEKFKEEVTRHQYLHHTEHFGFMTAGDFHRSAPLPVPLLESTLKLGRERFAQLAAVAPCPMGLENLALALSVDDALEQHDFLEKIVSPSNGFLLLDLHNIYCQLCNFNLSPAELLDNYPFHLVREIHVSGGSWEPTASSKEPIRRDTHDEDVPAEVFSLLDYVLPRCPNLEVVFLERLGNTLHEHQFVSFRKDFHTLKSIVANG |
| WP_105702996.1 | Chryseobacterium sp. MYb7 | MDKPLLGISMMAETEFVSALLPLLQSNSVDILEWSFDTLYHSNEPGWLCDLLNFYAENDRLIGHGVYYSLFDARWTGRQEEWLKKLKEEVSLRKYNHITEHFGFMNTENFHQGVPLPVSLNPKTLEIGKDRLYRLQEAVNIPVGIENLAFSFSVDDVKEQGVFLDKLTEDTDGFLILDLHNIYCQSYNFEVEMQEIIDLYPLGKVKEIHLSGGSWQESAYGKKQVRRDTHDDVIPKEIFSVLPSVLTQCQNLEYIIIERLGHTLKTDEEKDNFLKDFNSVKGTIEASDWKREKKKNWNKKDVKIAEKPLEDLVLFEEQSRLTQLLFDNIDAAVIKDLDFHYFKTKNWDPEMILTAQNIIRKWNPY |

| | | |
|---|---|---|
| WP_103293743.1 | Chryseobacterium lactis | MGKPLVGLSMMAEAEFVSAILPLLQNEEVAVLEWSFDSLYQTDEPDWLCDLLNFFAKNNRLIGHGVYYSLLDAKWTGRQEIWLKKLKEEVCKRKYNHITEHFGFMN<br>TENFHQGIPLPVSLHPTTLKIGKDRLSRLQDAVEIPVGIENLAFSFSIDDVKEQGVFLDRLIEDIDGFLILDLHNIYCQACNFGIDMHEIIGLYPLEKVKEIHLSGGSWQES<br>VYGKRLIRRDTHDDVIPEEIFAVLPAVVSLCPNLEYIIIERLGHTLKSDEEKRSFFNDFMKVKSIIGSLEGCYLQKRDWVKKENNLLDAPVEDLLLYEEQSKLTKLLFDS<br>AEIETVKDQDFHYFNTKNWDPEMILTAQNIIKKWNPY |
| PQJ27362.1 | Rubritalea profundi | MPETQFLQASMPLFEAGDVGAVEWSFDTGWKNGVSEWLTSLIGTYSEAGRLTGHGVHFSLLSADWTQRQQGWLEKFKEEVTRHQYLHHTEHFGFMTAGDFHR<br>SAPLPVPLLESTLKLGRERFAQLAAVAPCPMGLENLALALSVDDALEQHDFLEKIVSPSNGFLLLDLHNIYCQLCNFNLSPAELLDNYPFHLVREIHVSGGSWEPTAS<br>SKEPIRRDTHDEDVPAEVFSLLDYVLPRCPNLEVVFLERLGNTLHEHQFVSFRKDFHTLKSIVANG |
| WP_106522985.1 | Taibaiella chishuiensis | MTPILSSIACNLDAGILSASLPLFAEERVEAIEWSFDTLFNYKEIPSWFLELLHTYSDAGRLVGHGVFFSLFSGRFLPEQKAWLEALRKMAGTFRFDHITEHFGFMTG<br>KDFHSGAPLNIPYSDVTLRIGRDRLMRIQDAAQCPVGLENLAFSYSADEVKRHGEFLEMLVKPVNGFIILDLHNLYCQLHNFSMPYEQLIGLYPLHRVREIHISGGSW<br>EDAASGQGRTIRRDTHDDAVPTVVFDLLRRTLPLCPNLKYVVMEQLGTGLQTEGSKQGFYNDFITMQEIVQAYRPQPQQAALNAFLPVQAVSIAGRPEEDEILYAQ<br>QLELSGILEQASSCREAQDTLATSGLAGTAWHTERWDADMLETAIRIAQKWKRQQA |
| WP_105682666.1 | Chryseobacterium culicis | MRKPMLGVSMMAEADFVSAILPLLQNNSIEVMEWSFDTLYHAHEPDWLHDLLNFYAENDRLIGHGVYYSLFDARWTERQEEWLKKLTEEVSLRKYNHITEHFGFM<br>NTENFHQGVPLPVSLHPQTLEIGQDRLYRLQEAVNIPVGIENLAFSFSIDDVKEQGVFLDKLTEGTNGFLILDLHNIYCQSSNFEVPIQEIINLYPLDKVKEIHLSGGSW<br>QDSVYGDKQVRRDTHDDVIPQDILSVLPSVLTQCQNLEYIIIERLGDTVKTEEERDNFLNDFNTVSTIIETSDWKVKGKSGWSKKEMKLSEKPLENLVLYEEQSRLTK<br>LLFDNVEVKAIKDQDFYYFKTENWDAEMILTAQNIIKKWNPY |
| PSK92621.1 | Taibaiella chishuiensis | MPTIPVLVVAWANRPAGIQGGPDIKHLKLFVDHEQMTPILSSIACNLDAGILSASLPLFAEERVEAIEWSFDTLFNYKEIPSWFLELLHTYSDAGRLVGHGVFFSLFSG<br>RFLPEQKAWLEALRKMAGTFRFDHITEHFGFMTGKDFHSGAPLNIPYSDVTLRIGRDRLMRIQDAAQCPVGLENLAFSYSADEVKRHGEFLEMLVKPVNGFIILDLH<br>NLYCQLHNFSMPYEQLIGLYPLHRVREIHISGGSWEDAASGQGRTIRRDTHDDAVPTVVFDLLRRTLPLCPNLKYVVMEQLGTGLQTEGSKQGFYNDFITMQEIVQ<br>AYRPQPQQAALNAFLPVQAVSIAGRPEEDEILYAQQLELSGILEQASSCREAQDTLATSGLAGTAWHTERWDADMLETAIRIAQKWKRQQA |
| PSL33846.1 | Dyadobacter jiangsuensis | MLLDQMAKKPPKIKRWTLVQPVEWADSYVSMSEIYSSIACNLDTHILQAALPLFEQEKVEAIEWSFDTLYRFDEVPAWFTDLVSEFSSHNRLVGHGVYFSLFSGKW<br>STGQQEWLDKLKKLSGEYNFDHITEHFGFMTGEDFHKGAPIGIPLTSRTLALGNDRLRRIQDACQCPVGLENLAFSYSIDEVRKHGTFLDQLVESVNGFIILDLHNLY<br>CQVHNFNIDFADILRLYPLNKVREIHISGGSWEDTMTPPHMPVRRDTHDESVPEAVFEYLGKVIPQCPNLKFVVLEQMGTALATPASQAAFQSDFLRMDRIVKSLAR<br>KSLPIPAHLFAPLAGRANPVPLEDPDLHKQQMQLSHILETAANVSEAKAMLRSSDLHNSAWEVENWAPYMLETAMAIAQKWKNGFV |
| WP_106930259.1 | Adhaeribacter arboris | MPEILPAIACNLDADILAAALPLFGESRIEAIEWSFDTLFRTAQIPSWFLELLKAFSNEKRLVGHGVFFSLFSGKWSPNQQNWLKQLRQLSSEFDFDHITEHFGFMTG<br>QNFHYGAPLPIPFSKTTLAIGQDRLARIYDACHCPVGLENLAFSYSLDEVKQHGEFLEQLITPINGFIILDLHNLYCQIYNFSIAYEDIINLYPLHRVREIHISGGTWQDSN<br>VAPARKIRRDTHDEAVPKEVFQLLDWTIDKCPNLKYVVLEQLGNALKTKESKKIFYQDFLKMEVIVQSKSNSLAPQPTNLFLPDLPNLPTTIIEDENLHQQQLQLSAILE<br>KAPTYTEAFELLFSSSLAHSDWKIESWEPYMLETARSIAQKWKK |
| WP_245901666.1 | Chitinophaga ginsengisoli | MPKVLSAVACNLDANILAACLPLMEESRIEAIEWSFDALYKVKEVPAWFRELLTTFSNEKRLIGHGVFFSLFSGKWLPEQENWLKHLEQTAAEFQFDHITEHFGFMT<br>GKDFHRGAPLNIPYTTATLNIGRDRLKRIYEACKRPVGLENLAFSYSLDEVKRHGAFLEQLLEPVNGFIILDLHNLYCQLQNFNISFDEMIALYPLDRVREIHISGGSWD<br>DSAAAPDRSIRRDTHDDTVPPEVFQLLEMTIARCPHLKYVVLEQLGNGLVTEESKRCFYDDFLQMEEIVRKNSDENDADSSDPFLPLSSLLTGTAVEDMTLYQQQL<br>ELSAILESSASYTEAMHALRQSSLANSDWRIEQWEPYMIETAVKIAQKWKK |
| WP_106915993.1 | Chryseobacterium aurantiacum | MGKPLLGLSMMAEAEFVSAVLPLLQSNSVDVLEWSFDTLYHTEEPDWLGDLLDFYSENNRLIGHGVYYSLFDARWTERQSIWLDKLKEEVKKRKYNHITEHFGFM<br>NTENFHQGVPLPVSLHPKTLQIGKDRLFRLQEALDIPVGIENLAFSFSVDDVREQGLFLDKLIEDTHGFLILDLHNIYCQSCNFEIEMQEIINSYPLDKVKEIHLSGGSW<br>QDSVYGKKHIRRDTHDDIIPEDILSVLPWALSQCHNLEYIIIERLGHSIQQESEKNDFLEDFAKVKKLVSESDWKNQQEVHWNKKEIKLPETPLEDMLLHEEQTLLTRL<br>LFDNADVSLIKNQEFHYFKTKNWDREMILTAQNIIKKWNPY |
| WP_109619450.1 | Chryseobacterium oncorhynchi | MGKPLLGLSMMAEAEFVSAILPLLQSNSVDILEWSFDTLYNTKEPDWLEDLLNFYSENNRLIGHGVYYSLFDARWTERQDVWLDKLKEEVKKRKYNHITEHFGFMN<br>TENFHQGIPLPVSLHHKTLEIGKDRLYRLQKAVDIPVGIENLAFSFSVDDVKEQGIFLDKLIEDTDGFLILDLHNIYCQSCNFEVEMREIINSYPLNKVKEIHLSGGSWQN<br>SVYGKKNIRRDTHDDVIPEEILSVLPWVFSQCQNLEYVIIERLGHSIHGESEKNDFLEDFVKVKELMSASDWKDQQKNHWNKNEIKLSENPLEDMLLYEEQTVLTRLL<br>FDSADIALIKSKEFHYFKTENWDPEMILTAQSIIKKWNPY |
| WP_110011639.1 | Chryseobacterium sp. AG844 | MRKPMLGVSMMAEADFLSAVLPLLQDNAIEVMEWSFDTLYHTNEPDWLRDLLNFYAENNRLIGHGVYYSLFDAIWTARQEEWLKKLKEEVSLRKYNHITEHFGFM<br>NTENFHQGVPLPVSLGPETLQIGKDRLYRLQETVDIPVGVENLAFSFSIDDVKEQGVFLDKLIEDTDGFLIFDLHNIYCQSCNFEVGLQEIINLYPLHKIKEIHLSGGSW<br>QESAYGKKQVRRDTHDDVIPEEILSVLPSVLTQCPNLEYIIIERLGHTLKTEEERNDFLDDFNKVRTIIETSDLKFKENKRWCKKEVKLSEKPLEDLLLYEEQSKLTRLL<br>YDNSDVKAIQDADFHYFKTQHWDAEMILTAQNIIKKWNPY |
| PWW16924.1 | Chryseobacterium sp. AG844 | MLGVSMMAEADFLSAVLPLLQDNAIEVMEWSFDTLYHTNEPDWLRDLLNFYAENNRLIGHGVYYSLFDAIWTARQEEWLKKLKEEVSLRKYNHITEHFGFMNTENF<br>HQGVPLPVSLGPETLQIGKDRLYRLQETVDIPVGVENLAFSFSIDDVKEQGVFLDKLIEDTDGFLIFDLHNIYCQSCNFEVGLQEIINLYPLHKIKEIHLSGGSWQESAY<br>GKKQVRRDTHDDVIPEEILSVLPSVLTQCPNLEYIIIERLGHTLKTEEERNDFLDDFNKVRTIIETSDLKFKENKRWCKKEVKLSEKPLEDLLLYEEQSKLTRLLYDNSD<br>VKAIQDADFHYFKTQHWDAEMILTAQNIIKKWNPY |
| WP_109709735.1 | Chryseobacterium phosphatilyticum | MEKPLLGISMMGEAEFLSAVLPLLQNNLVDVLEWSFDTFFNAKEPDWLDSLLNFYAENNRLIGHGVYYSLFDAKWTERQEIWLEKLKEEICKRKYNHITEHFGFMNT<br>ENFHQGVPLPVSLHPKTLQIGQDRLQRLQNVINLPVGIENLAFSFCLDDVREQGVFLQQLIKDVDGFLILDLHNIYCQACNFEIGMQEIIDLYPLDKVKEIHLSGGSWE<br>ESVYGRKMIRRDTHDHAIPEDIFSILPFVLSHCKNLEYVIIERLGSTIRTADEKKKFLDDFTRVKAMIDSSVWNERQKDSWTKKEYNLSKTPVEDVMFYEEQSRLTKIL<br>FDNTAVDALKNEKFHYFNTVNWEPEMIRTAQNIIKKWNPY |
| WP_233259989.1 | Chitinophaga sp. S165 | MPKILSAVACNLDTNILAACLPLMEESRVEAIEWSFDALYKVKEVPSWFRELLTAFSDEKRLIGHGVFFSLFSGKWLPEQENWLKQLEQTAAEFRFDHITEHFGFMT<br>GKDFHHGAPLNIPYTATTLNIGRDRLKRIYNACRVPVGLENLAFSYSLDEVKRHGAFLEQLLEPVNGFIILDLHNLYCQLRNFDIAFDEMIALYPLDRVREIHISGGSWE<br>DSAAAPERSIRRDTHDDAVPSEVFQLLEMTIGKCPQLKYVVMEQLGNGLVTDASRRAFYDDFLQMEEIVRKNSDESNTGSSDLFLPLSSFSTDIAVEDMALYQQQL<br>ELSSILESAVSYEDAMQVLTHSSLANSDWQIEQWEPYMIETAVKIAQKWKK |
| PTT33357.1 | Chryseobacterium sp. HMWF028 | MSKPLLGISMMAEAEFVSAVLPLLQDNSVDVLEWSFDTFYHAHEPDWLRDLLNFYAENERLIGHGVYYSLLDARWTERKEEWLKKLKEEVTLRKYNHITEHFGFMN<br>TENFHQGVPLPVSLHPKTLEIGKDRLYRLQEAVNIPVGIENLAFSFSIDDVKEQGVFLDKLIEDTDGFLILDLHNIYCQSCNFGVRLQEIIDLYSLDKVKEIHLSGGSWQ<br>DSVYGKKPVRRDTHDDVIPEEIFSVLSSVLTQCQNLEYVIIERLGHTLKTHEEKDNFLSDFNNVKAIIEASDWKKRGKKIRNKKEIKLSVKPLEDLILFEEQSRLTKLLFD<br>NADAKVIKDLDFHYFKTENWDLEMILTAQNIIKKWNPY |
| WP_106593918.1 | Dyadobacter jiangsuensis | MSEIYSSIACNLDTHILQAALPLFEQEKVEAIEWSFDTLYRFDEVPAWFTDLVSEFSSHNRLVGHGVYFSLFSGKWSTGQQEWLDKLKKLSGEYNFDHITEHFGFMT<br>GEDFHKGAPIGIPLTSRTLALGNDRLRRIQDACQCPVGLENLAFSYSIDEVRKHGTFLDQLVESVNGFIILDLHNLYCQVHNFNIDFADILRLYPLNKVREIHISGGSWE<br>DTMTPPHMPVRRDTHDESVPEAVFEYLGKVIPQCPNLKFVVLEQMGTALATPASQAAFQSDFLRMDRIVKSLARKSLPIPAHLFAPLAGRANPVPLEDPDLHKQQM<br>QLSHILETAANVSEAKAMLRSSDLHNSAWEVENWAPYMLETAMAIAQKWKNGFV |
| RPD50464.1 | Hymenobacter sediminis | MAEGRVEALEWSFDALFWAEQVPDWFSELLTTYSAANRLLGHGVYFSLLSGRWTPEQQQWLEQVRQVVAQYPLAHLTEHFGFFTGQNFHSGAPLPVPYTPATL<br>RLGQDRLSRIYEACRCPVGLENLAFAYSLDEVKRHGEFLGRLLEPVNGFLILDLHNLYCQIHNFSVSYEELIGLYPLERVREIHISGGSWENSELAPGRKIRRDTHDE<br>GVPEEVFQLLELTIPRCPRLKYVVLEQLGPSLKTEESRIRFRHDFGRMEEVVHRHRAGRLGYADAFFLPPQPLQTGPALEDAQLHAHQQQLSHILETAPSFAEAQR<br>QLHASALASSDWKLEQWAPYMLETAVSIAQKWK |
| WP_181416153.1 | Chryseobacterium sp. CBTAP 102 | MLGVSMMAEADFISAVLPLLQNNSIDVMEWSFDTLYYANEPDWLRDLLNFYAENNRLIGHGVYYSLFDARWTERQEEWLKKLKEEVNLRKYNHITEHFGFMNTEN<br>FHQGVPLPVSLHPKTLQIGKDRLYRLQEAVNIPVGIENLAFSFSIDDVKEQGVFLDKLTEGTNGFLILDLHNIYCQSCNFELEMQKIIDLYPLDKVKEIHLSGGSWQDSV<br>YGKKQVRRDTHDDVIPRDILFVLPSVLTQCENLEYIIIERLGNTLKTDEERDHFLDDFSKVKTIIETSELNGKEKSNWSKMKVELSEKPLEDPVLYEEQSRLTQLLYDN<br>ADVETIKDLDFYYFKTESWDAEMILTAQNIIKKWNPY |
| PZD74685.1 | Acaryochloris thomasi RCC1774 RCC1774 | MPQADFWQTAQPLLESSAVEVLEWSFDMGWGKVGLPTWVPPLLDQFSQQDRLLGHGVSYSLLSAQLDRSHWLACLEAERRQYHYQHISEHFGWLATETFRQSA<br>PLPMPLCPETLQLGRERLQRLAEVAQVPVGLENLAFAFSQQDVREQGIFLEKLLESVDGFLLLDVHNLYCQVCNFQISATELLALYPLHRVREIHISGGSWSKGTGG<br>QIRRDTHDQPVPEAVFELLTLALQQCLNVQFVIFERLGNTLGPASEQSQFRQDFARIQAIVQADREG |
| PZM82676.1 | Candidatus Melainabacteria bacterium | MPTKPKLGLSLMPTEDFEQATVPLFQNNQVEAIEWSFDFSWNGAHVGEWCNKILDEFSQKGALTGHGVNLSPLSARFSQRQQKWLDNAKEEFKARKYIHASEHF<br>GFSEAGPIAQGAPLAVPMNAESLRKGKEMLKRYADATQCPVGLENLAFAFSIEDVKRQGDFIDRLISSVDGFLLLDLHNIYCQIANFGVSELELLNSYPLAKVREIHLS<br>GGTWTHSISGKRAAVRRDTHDDAMPQEVFNLATLALKLCPNIEFVIMERLGYTMMELEEQQEFREDFETMEEILEFCYAN |
| WP_199474262.1 | Adhaeribacter pallidiroseus | MPEKVLAAVACNLDVDILAAALPLFAAGQVEAIEWSFDTLFRTPQMPDWFAELLQTYSQQNRLIGHGVYFSLFSGQWSVNQQNWLQHLRELSKLYTFDHITEHFGF<br>MTGQNFHHGAPLPIPYSETTLAIGQDRLARIQDACQCPVGLENLAFSYSLDEVKQHGQFLEQLITPVNGFLILDLHNLYCQLHNFQASYNEIINLYPLDRVREIHVSGG<br>SWESSAVEPDRKIRRDTHDEAVPEEVLELLAQIIEKCPNLKYVVLEQLGNALKSEKSKQLFYADFLKMEAILRQKNQKSRTNAGKTFLPAPFKLPEQPIEDQTLAAQQ<br>IYLSEILENAASAEEAIHLLQTSPLAHSDWQIETWEPYMIETAHNIARKWKK |
| WP_111952938.1 | Chryseobacterium lathyri | MKKPLFGLSMMPEADFVSAILPILQNNSVDVLEWSFDTFYDAQEPEWLSGLLDFYSANNRLLGHGVYYSLFDAKWTERQEGWLKKLKEEFRHRKYNHITEHFGFM<br>NTENFHQGVPLPVPLHPKTLQIGKDRLKRLQDAVEIPVGVENLAFSFSIDDVKEQGDFLDKLIEDIDGFLILDLHNIYCQSWNFEISMQEIIRLYPLEKVREIHLSGGSW<br>QDSAYGKKPVRRDTHDESIPEEILNILPEVLSQCHNLEYIIIERLGHTLNTDSEKDAFFDDFNRVKKIIQASDSLPGNKRVWHKKNEESSVPLEDLLLYDEQTELTKMLF<br>DGIQIHSVKKHDFHYFKPENWDEEMITTAQQIIKKWNPY |
| WP_114789236.1 | Niabella yanshanensis | MPEIFSSIACNLDNDILAACLPLFESEKVEAIEWAFDTIFSKEVIPAWFTELLQAYSSQGRLIGHGVFFSLFSGNWSKEQENWLLKLKALSKQFRFDHITEHFGFMTGS<br>DFHKGAPLSVPLTASTMRIGIDRLQRIQDACNCPVGLENLAFACSAEEVKRHGSFLLELIESVNGFIILDLHNFYCQSHNFNIPFDTLIKYYPLDKVREIHISGGSWEPV<br>SVQPARTIRRDTHDDAVPETVFGFLEKTLPRCPNLKFVVLEQLGNGLTTPESRLQFQKDFYRMRAIVQSAQKNKKDGSPDPFLPPMGQRVFSTPTEDPLLGLQQQ<br>ELSDIFENAESVNDAIRLLKRSSLAQSAWKIEYWDEAMLETAINIAQKWKNGFD |
| RBL88330.1 | Chitinophaga flava | MEASRVEAIEWAFDTLFNVTEVPSWFEELLTAYSQENRLIGHGVFFSLFSGKWLPEQESWLKHLRKVSNDFHFDHVTEHFGFMTGKDFHHGAPLNIPYTTATLDIG<br>RDRLKRICEACSCPVGLENLAFSYSPEEVKKHGAFLDQLLEPVNGFIILDLHNLYCQVHNFNLSFDDISMLYPLHRVREIHISGGSWENSIATPGRNIRRDTHDDAVP<br>EEVFAYLEKAIDLCPHLKYVVMEQLGTGLYTDASKIAFRNDFLRMDDIIKNKNKTIAVANSFLPETLIVPPEVKEDLSLYQQQLELSNILETALSYETAMTALSSSSLAHS<br>AWQIEQWDPAMIETAVNIAQKWKKKT |
| WP_123881242.1 | Chryseobacterium carnipullorum | MSKPLLGLSMMPEADFVSAILPLLQNNSVDVLEWSFDTFYDVNEPEWLSGLLDFYAANNRLLGHGVYYSLFDARWNERQEIWLKKLKKECANRKYNHITEHFGFM<br>NTENFHQGVPLPVPMHPLTLHIGKDRLKRLQDAVEIPVGVENLAFSFSIDDVKEQGVFLDQLVEDIEGFLILDLHNIYCQSCNFGVSMQEIIKLYPLEKVKEIHLSGGS<br>WQESAYAKKPVRRDTHDDRIPDEIVEMLPEVLSQCQNLEYIVIERLGHTLCAESDKLFFFADFNRVKEIISKSEDRSEDKKVWNRRIGVDSKPVEDMVLYDEQIKLTK<br>MLFDESSVAKVKNEAFQYFKPENWDEEMIMTAQQIIKKWNPY |
| WP_114822528.1 | Chryseobacterium sp. KLBC 52 | MRKPMLGVSMMAEPDFVSAILPLLQDNSIEVMEWSFDTLYHTNEPDWLRDLLNFYAENNRLIGHGVYYSLFDARWTERQEEWLKKLKEEVRLRKYNHITEHFGFM<br>NTENFHQGVPLPVSLHSKTLQIGKDRLYRLQDAVSIPVGIENLAFSFSIEDVKEQGVFLDKLTEDTHGFLILDLHNIYCQSCNFELEMHEIIDLYPLDKVKEIHLSGGSW<br>QDSVYGKKQMRRDTHDDVIPKDILSVLPSVMKKCSNLEYIIIERLGHTLKTEEERNNFLEDFNKVRTIIETSDLKLREQNGWCKEKLKFSAKPLEDLLLHEEQSKLTRL<br>LYDNANVETIKDLDLHYFKTENWDAEMILTAQNIIKKWNPY |
| REC44108.1 | Chryseobacterium pennipullorum | MKKPLLGLSMMAEAEFVSAILPLLQSNLVDVLEWSFDTFFHAEEPDWLNDLLDFYSDSRRLTGHGVYYSLFDAKWTERQEVWLDTLKKEVQTRTYNHITEHFGFM<br>NTENFHQGVPLPVSLHPVTLQIGKDRLFRLQDAVNIPVGIENLAFSFGIDDVKEQGVFLDKLTEDTNGFLILDLHNIYCQSCNFEIPMEEIINSYPLDKVREIHLSGGSW<br>QESAYGKKQIRRDTHDDAIPESLLDVLPSVLLKCKNLQYVIIERLGHTISTEKEKKRFTEDFQKVKKMLEAPGLLTENTTSWIKKGVVIPDVPLEDLALAEEQSRLTRLL<br>FEGADAAAFKKENFHFFNTRNWDSEMIRTAQNIIKKWNPY |
| WP_110984643.1 | Acaryochloris thomasi | MATTTTTRPSLGLSLMPQADFWQTAQPLLESSAVEVLEWSFDMGWGKVGLPTWVPPLLDQFSQQDRLLGHGVSYSLLSAQLDRSHWLACLEAERRQYHYQHIS<br>EHFGWLATETFRQSAPLPMPLCPETLQLGRERLQRLAEVAQVPVGLENLAFAFSQQDVREQGIFLEKLLESVDGFLLLDVHNLYCQVCNFQISATELLALYPLHRVR<br>EIHISGGSWSKGTGGQIRRDTHDQPVPEAVFELLTLALQQCLNVQFVIFERLGNTLGPASEQSQFRQDFARIQAIVQADREG |

| | | |
|---|---|---|
| WP_115833250.1 | Dyadobacter luteus | MREIYSSLACNLDANILLASIPLFEQENIEAIEWSFDALYKYDQIPPWFADLVSEYARNNRLVGHGIYFSIFSGKWLPEQQQWLNKLAVFCKEFRFDHISEHFGFMTGADFHKGAPISIPFTSKIVELGADRLARIQEICQCPVGLENLAFAYSFNEVKKQGEFLDCLIEKVNGFLILDLHNLYCQLHNFSISARELLKAYPLERVREIHISGGSWEQGLASDRKIRRDTHDDAVPEEVFALLNDTIPLCPNLKFVVLEQLGTGLTTDESRAQFQTDFNRMDSIVRLHRTNNADSLDNDFIPKVPLHINTTPVEDVLLYQQQQELSHILETASDLNHAQQLLLASSLNRSAWHIDQWTPEMLHTAISIAQKWKNGF |
| RDV15053.1 | Pontibacter diazotrophicus | MEEGRVEAIEWSFDTLYWAAQVPYWFTDLLLAYSQQSRLIGHGVFFSLLSGKWTKEQQHWLEQLKQLSSQYSFDHITEHFGFFTGQNFHSGAPLYVPYTSTTLRIGQDRLSRIYDACQCPVGLENLAFSYSLDEVKQHGEFLEKLVAPVNGFLILDLHNLYCQLHNFSITYEDLITLYPLNRVREIHISGGSWEASVLEPEKKVRRDTHDDAVPEEVFHLLELTVPKCSNLKYVVLEQLGNGLKTESSRNRFRQDFLKMEALVQKSRSSLSRQDDNLFLPVQPLQTGPVVEDDMLHDQQLQLSHILETAPSYEEARRLLFTSSLAGSDWKIENWAPHMLETAVSIAQKWKR |
| WP_115919746.1 | Chryseobacterium rhizosphaerae | MGRPLLGLSMMAEADFVSAILPLLQDNAVDVVEWSFDTFFAAEEPSWLKDLLNFYAENGRLLGHGVYYSLFDARWTERQEMWLQKLKHEVQKRNYNHITEHFGFMNTENYHQGVPLPLSLHPKTLEIGKDRLYRLQDAVPIPIGVENLAFSFSVDDVREQGVFLDKLVDDIDGFLILDLHNIYCQSCNFDVSIQEIVSLYPLEKVKEIHLSGGSWQDSVYGKKMIRRDTHDDVIPEEIFSVLPSVLSQCRNLEYVIIERLGHTIHTEEEKQRFFNDFVRVKSILNDSDGMKRKKEKWVKREMLLPEPVEDSILYEEQIRLTKLLFDHTDPTLIKNQNFHYFKTENWDPEMILTAQNIIKKWNPY |
| WP_115959376.1 | Chryseobacterium flavum | MERPLVGVSMMAEADFVSAVLPLLQDNAIDVLEWSFDTFFETEEPDWLCDLLNFYAGNDRLIGHGVYYSLFEAKWTDRQEIWLEKLKKEVCKRKYNHITEHFGFMNTENYHQGIPLPVPLHPKILEIGKDRLNRLQDAVQLPVGVENLAFSFSVEEVKEQGIFLDKLIEEMEGFLILDLHNIYCQACNFGVEMQEIIGSYPLHKVKEIHLSGGSWQNSVYRKKPVRRDTHDDAIPEELFAVLPSVLSQCSNLDYVIVERLGHTINTEDLKQNFLDDFIKVKTIVEKSGCQKQNTKDWFPKKNILAKPVEDAVLYEEQVRLTKLLFNHTDPIIIQHEDFHYFNTGNWDHEMILTAQNIIKKWNPY |
| WP_116014710.1 | Chryseobacterium elymi | MGKPLLGLSMMPEADFVSAILPILQNNSVDVLEWSFDTFYDAQEPEWLSGLLDFYSEHNRLLGHGVYYSLFDARWTERQEGWLKKLKEEFRHRKYNHITEHFGFMNTENFHQGVPLPVPLHFKTLQIGKDRLKRLQDAVEIPVGVENLAFSFSIDDVKEQGEFLDKLLEDIDGFLILDLHNIYCQSWNFDISMQEIIRLYPLEKVKEIHLSGGSWQESAYGKKPVRRDTHDGSIPEEILNVLPEVLPQCHNLEYIIIERLGHTLNTGSKKEAFFHDFNRVKEITQVSDSHTKNQKVWHKKNGECVEPVEDLLLYEEQTKLTRMLFDGTDIQSVKDHGFHYFKPENWDEEMITTAQQIIKKWNPY |
| WP_116105833.1 | Lewinella sp. IMCC34191 | MVYPSVACNLDRHLLATVLPLLEEGEVAGMEWSYDAIYRYESLPDWFGQLLTAFAREGRLVGHGIFYSVCAASWTEDQKNYLDQLSSVARTYPFDHVTEHFGFLTGKDFHRGAPLSPPLTAATLRIGRDRIRRLRDAANCPVGLENLAFAYRVEDVYRQYAFLDQLVEPVDGLLILDLHNLYCQLHNFSLDLEDLLRHIPLERVREMHVSGGSWESHSSAPTGRVRRDTHDERVPNEVFHLLREIGPRCPRLKFVVLEQLGTALHDLTAQDGFRSDFQRVAAICETIHSTVPADDRDKPLSADPIPEPPLQDRQLAAEQRELSDILETSSSYSEAHDRLARSLLANSEWHVESWAPHMVETAWKISRKWR |
| WP_273008157.1 | Chryseobacterium sp. | MRKPLLGLSMMAETEFVSAILPLLQSNSVDVLEWSFDTLYHTKEPDWLGDLLNFFSENNRLIGHGVYYSLFDARWTERQNIWLNKLKKEVKKRKYNHITEHFGFMNTDNFHQGVPLPVSLHSKTLQIGKDRIFRLQEALDIPVGIENLAFSFSIDDVREQGLFLDKLIEDTDGFLILDLHNVYCQSCNFEVEMQEIINSYPLDKVKEIHLSGGSWQDSVYGKKLIRRDTHDDVIPEEILSILPWVFSQCQNLEYVIIERLGHTIHLESEKDDFLKDFAKVEKLISESDWRNQKKINWNKKELKLSESPLEDTLLHKEQTMLTRLLFDSADVSLIKNQEFHYFKTDTWDPEMILTAQNIIKKWNPY |
| WP_115969000.1 | Chryseobacterium pennae | MGKPLLGLSMMAEAEFLSAVLPLLQSNSIEVLEWSFDTFYNTEEPDWLSDLLNFYSENNRLIGHGVYYSLFDARWTKRQELWLQKLKEEFHKRKYNHITEHFGFMNTENFHQGVPLPVSLHSKTLAIGKDRLYRLQDAVQVPVGIENLAFSFSLDDVLEQGVFLDKLTDDTDGFLILDLHNIYCQSCNFEVQMQDIIKLYPLDKVKEIHLSGGSWQESAYGKKWVRRDTHDDVIPEDILSVLPSVLSQCRNLEYIIIERLGHTINTEEKKKSFLDDFAKVKMLIDSSDWHAQNKENRNRKEIQFPEHPLEDRILHEEQTLLTKLLFDNVEPESVNHHTFQYFKTEKWDPEMIITARNIIKKWNPY |
| HAY78875.1 | Planctomycetaceae bacterium | MLEADFAAATQPLFATGDVEVLEWSFDVGWGRSVPPWAEQLVSFYSQHQQLLGHGVSFSPLSAEWQTRQTDWLRQLGQEVARRDYQHVSEHFGFMTAGDFHQSAPLPVPRTPATLAIGRDRLARLADTCPCSVGLENLAFAFGQRDVASQGDFLDALLEPVDGFLLLDLHNIYCQACNFDVDPWSLIESYPLDRVRELHVSGGSWTEARDGQAVRRDTHDGSVPALLFELLPRVLDRCPQVQYVILERLGGTFTNAEAEFQADFRRLRHTVETWGEWATSSSANSKHQP |
| RFM37115.1 | Chitinophaga silvisoli | MEESRVEAIEWSFDALYKVKQVPDWFRELLTAFSDENRLIGHGVFFSLFSGKWLPEQQAWLDHLKHTSTEFSFDHITEHFGFMTGKDFHHGAPLNIPYSAATLSIGRDRLKRIYNACGRPVGLENLAFSYSLDEVKRHGAFLDQLLAPVNGFIILDLHNLYCQLRNFDLAFDDLISLYPLDKVREIHISGGSWDDSAASPDRSIRRDTHDEAVPPEVFQLLEWTLPKCPHLKYVVLEQLGNGLVSQESKAAFYNDFLHMQEIVSQSNNKYSDNSFLPLLPTTTGPAVEDLALYQQQLELSAILETAASYDSAMHLLAQSSLANSDWRIEQWEPYMIETAVKIAGKWKK |
| WP_291873038.1 | Chryseobacterium sp. | MKKPLLGLAMMPEADFVSAILPLLQTQSVDVLEWSFDTFYDLEEPEWLSGLLDFYSDNNRLLGHGVYYSLFDARWMERQEIWLKKLKEEVKRRKYNHITEHFGFMNTENFHQGVPLPVPLLPKTLQVGKDRLSRLQDAVEIPVGVENLAFSFSMDDVKEQGEFLDKLVEDIDGFLILDLHNIYCQSCNFEVDMREIINLYPLEKVKEIHLSGGSWQESAYGKKPVRRDTHDDRIPEEILNLLPEVLIRCHPEYVIIERLGHTLHTEAEKQIFFDDFNRVKKIIELSDYPVDEEKIWNRKKTVHSKPVEDLLLHAEQTELTRLLFDGNTTESIKNSDFHYFKPESWDEEMITTAQQIIKKWNPY |
| HCN49489.1 | Chryseobacterium sp. | MEKPLLGLSMMAEAEFVSAILPLLQNNSVEVLEWSFDTFYNAEEPDWLNELLQFYSENNRLIGHGVYYSLFDARWTERQEIWLKKLKEEVRKRNYNHITEHFGFMNTENFHQGVPLPVSLHPKTLQIGRDRLYRLQDAVDVPVGIENLAFSFSIDDVKEQGMFLDQLVEDTNGFLILDLHNIYCQSYNFEVNIQDIISLYPLDRVREIHLSGGSWQESVYGKKPIRRDTHDDRIPKELLAILPSVLSQCQNLEYVIIERLGHTIKTEEQKKYFFDDFNKVKAIIEVSDTRSRKVNKWTGKEAVHTDHPEEDLALYEEQSRLTRLLFDQAGTEGIKEEKFHYFNTESWDPEMILTAQNIIKKWNPY |
| WP_276729926.1 | Chryseobacterium carnipullorum | MSKPLLGLSMMPEADFVSAILPLLQNNSVDVLEWSFDTFYDVIEPEWLSGLLDFYADNNRLLGHGVYYSLFDARWNERQGIWLKKLKKECANRKYNHITEHFGFMNTENFHQGVPLPVPMHPLTLQIGKDRLKRLQDAVEIPVGVENLAFSFSIDDVKEQGVFLDQLVEDIEGFLILDLHNIYCQSCNFGVSMQEIIKLYPLEKVKEIHLSGGSWQESAYAKKPVRRDTHDDRIPDEIVEMLPEVLSQCQNLEYIVIERLGHTLSAESDKLFFFADFNRVKEIISKSEDRSEDKKVWNRRIGVDSKPVEDMVLYDEQIKLTKMLFDESSVAKVKNEAFQYFKPENWDEEMIMTAQQIIKKWNPY |
| WP_118972670.1 | Taibaiella koreensis | MEYEPLRSTIACNLDAGILSASLPLFEEARVEAIEWAFDTLFRLETIPVWFEELLEAFSKEKRLIGHGVFFSLFSGRWQPEQQRWLDQLRTLCRKYHFDHITEHFGFMTGADFHHGAPLGIPFTAETLALGRDRLARIYDACQCPVGLENLAFSYSLEEVERHGAFLEELLQPLNGFIILDLHNLYCQLHNFSLDFDTLIKRYPLHRVREIHISGGSWEPSALFEGKSVRRDTHDDTVPEEVFVLLEKTIPLCPCLKYVVMEQLGTGLATEPSRQAFRNDFIRMQQIVSAATAQPAAFLPEAFLPPDILPAGPVVESATLHNQQLELSAILESSAGYEDARQRLAHSSLAGTGWQVEAWQPHMLETAVHIAQKWRKKNT |
| WP_119445210.1 | Hymenobacter oligotrophus | MQAAPAIYASVACNLDADLLAAAFPLLEEGRVEALEWSFDALYWAEQVPEWFTALLHTYAEAGRLVGHGVFFSLLSGKWSPEQQQWLAHLGQLAEQFRFDHVTEHFGFFTGQNFHHGAPLPIPYTRHALRIGQDRLRRIYEACRCPVGLENLAFAYTPDEVKRHGAFLEELLLPVNGFLILDLHNLYCQLHNFGLPYDELIALYPLERVREIHISGGSWHDSALAPGRRIRRDTHDDAVPAEVFQLLEWTMPRCPNLKFVVLEQLGHALHAEASRAQFRADFARMESLVAHHRATWRPQPANAFRPLQAWLPGPAAEDEHLHQQQRQLSDILENAGSYTEAQHLLQTSSLAHSDWCIEQWEPHMLETALHIARKWKK |
| AXY78851.1 | Paraflavitalea soli | MEESRIEALEWSFDTLYKVQEIPTWFQELLTAFSKEKRLIGHGVFFSLFSGKWLPEQEAWLKALQQTCSEFQFDHITEHFGFMTGKNFHQGAPLNIPYTPTTLAIGQDRLKRIYAACQCPVGLENLAFSYSLDEVKRHGRFLDKLLEPVNGFIILDLHNLWCQLHNFSIALEELIGLYPLDRVREIHISGGSWDHSQIEKGRTIRRDTHDDAVPAAVFQLLETVIPACPHLKYVVLEQLGNGLESIESKQQFYADFLEMESIIRRQSQPAYLTNLASFLPLSPIVMGPPVEDEGLYEQQAALSTILETSHSYEEARTLLGQSSLACSEWQVENWDPAMIETAMKIAQKWKAGFVGFIS |
| RKE81000.1 | Chryseobacterium sp. AG363 | MMAEPDFVSAILPLLHNNSIEVMEWSFDTLYHTNEPDWLRDLLNFYAENNRLIGHGVYYSLFDARWTARQEEWLKKLKEEVSLRKYNHITEHFGFMNTENFHQGVPLPVSLHSKILQIGKDRLYRLQDAVNIPVGIENLAFSFSIEDVKEQGIFLDKLTEDTNGFLILDLHNIYCQSCNFELEMQQIIDLYPLDKVKEIHLSGGSWQDSVYRKKQVRRDTHDDVIPKDILSVLPSVMKKCPNLEYIVIERLGHTLKTEEKRNSFMEDFTKVRTIIETSDLKLKQQNGWCKEEIKLPAKPLEDLLLYEEQSRLTRLLYENANVENIKDLDFHYFKTENWDAEMILTAQNIIKKWNPY |
| WP_121487547.1 | Chryseobacterium sp. 7 | MNKPLLGISMMAETDFVSAVLPLLQSNSVDVLEWSFDTLYHSHEPDWLRDLLNFYAENNRLIGHGVYYSLFDARWTGRQEEWLKKLKEEVSLRKYNHITEHFGFMNIENFHQGVPLPVSLNPKTLEIGKDRLYRLQEAVNIPVGIENLAFSFSVDDVKEQSVFLDKLTEDTNGFLILDLHNIYCQSYNFEVEMQEIIDLYPLEKVKEIHLSGGSWQESAYGKKQVRRDTHDDVIPKEIFSVLPSVLTQCQNLEYVIIERLGHTLKTEEEKDDFLNDFNQVKGIIEASDWKENEKKSWNKKEIKLSEKPLEDLVLFEEQSRLTRLLFDNVDAETIKDQDFHYFKTEKWDPEMILTAQNIIKKWNPY |
| GDY10763.1 | Planctomycetia bacterium | MTAPQVGLSLMLEPDFVAAALPLLQAGRVDVLEWSFDIGWSQAAMPEWADELLDHFASQDRLLGHGVKYSLLSADESPARDAWLERLSAECRRRHYRHVSEHFGFMAAGRFHEGAPLPVPLTEFTVRLGRERLQRLADAARCPVGLENLAFAFGPRDVQDQGRFLDELLTPLDGFLLLDLHNIDCQCCNFDVSGAELLATYPLHRVREIHVSGGSWSSVKVPSPKQLIRRDTHDAAVPREVFDLVRWAIPRCPNLEAVILEQLGGTLLEANAAEQFRTDFQTLRQVVSEVSNDRV |
| WP_119080626.1 | Chitinophaga alhagiae | MPRLLSTVACNLDPHLLRACLPLFESGEADAIEWSFDALYRLDALPDWFHGLINAFSEGNRLIGHGIFFSLFSGKWHPEQETWLRELSAVCRKYRFDHITEHFGFMTGQDFHHGAPLNIPYTDAALRIGRDRLLRIADACGRPVGIENLAFSWSVDEVRQHGDFLDQLLEPVNGFIILDLHNLYCQVANFNLPFETLAALYPLHRVREIHISGGSWELSEDQSRSIRRDTHDDSVPEEVFRYLENIIPRCPHLRYLVMEQLGSGLETEDSKNAFRADFRRMAAIARQWNLTKGDEILHPFSPPANPPQGAPLEDLALYQQQTELSSILENAPSFEAATRALQASSLAQSHWQVERWSPFMLETAMHIAQKWMKKN |
| WP_120230832.1 | Chryseobacterium sp. AG363 | MRKPILGVSMMAEPDFVSAILPLLHNNSIEVMEWSFDTLYHTNEPDWLRDLLNFYAENNRLIGHGVYYSLFDARWTARQEEWLKKLKEEVSLRKYNHITEHFGFMNTENFHQGVPLPVSLHSKILQIGKDRLYRLQDAVNIPVGIENLAFSFSIEDVKEQGIFLDKLTEDTNGFLILDLHNIYCQSCNFELEMQQIIDLYPLDKVKEIHLSGGSWQDSVYRKKQVRRDTHDDVIPKDILSVLPSVMKKCPNLEYIVIERLGHTLKTEEKRNSFMEDFTKVRTIIETSDLKLKQQNGWCKEEIKLPAKPLEDLLLYEEQSRLTRLLYENANVENIKDLDFHYFKTENWDAEMILTAQNIIKKWNPY |
| WP_123278455.1 | Chryseobacterium cucumeris | MRKPILGVSMMAEPDFVSAILPLLHNNSIEVMEWSFDTLYHTNEPDWLRDLLNFYAENNRLIGHGVYYSLFDARWTARQEEWLKKLKEEVSLRKYNHITEHFGFMNTENFHQGVPLPVSLHSKILQIGKDRLYRLQDAVNIPVGIENLAFSFSIEDVKEQGVFLDKLTEDTNGFLILDLHNIYCQSCNFGLEMQQIIDLYPLDKVKEIHLSGGSWQDSVYRKKQVRRDTHDDVIPKDILSVLPSVMKKCPNLEYIIIERLGHTLKTEEERNYFLADFNKVRTIIETSDLKLKQQNGWCKEEIKLPAKPLEDLLLYEEQSRLTRLLYENANVENIKDLDFHYFKTENWDAEMILTAQNIIKKWNPY |
| WP_123274716.1 | Chryseobacterium sp. G0240 | MRKPFLGISMMAEPDFLAAILPLLQDGSIDVLEWSFDTFYHAEEPDWLANLLNFYSQKDRLLGHGVYYSLFDARWTDRQEKWLEKLQQEVQQRRYAHITEHFGFMNTENFHQGVPLPVSLHPKTLEIGKDRLLRLQEVVDIPVGIENLAFSFSIDDVKEQGVFLEKLTADINGFLILDLHNIYCQACNFGVDMEELIRLYPLEKVREVHLSGGSWQTSSYGKRQIRRDTHDDSIPEEILTVLPSVLSQCKNLEYVIIERLGHTIQTEKEKKDFFDDFVRVKTIVGALAWKDYDKTCWTKKNIRLTGKPLEDEALYQEQTKLTKLLFDNVESGAFKEQDFQFFKTENWDPEMIRTAQAIIKKWNPY |
| WP_123319418.1 | Chryseobacterium sp. BIGb0232 | MGKPLLGLSMMAEAEFLSAVLPLLQSNSIEVLEWSFDTFYNTEEPDWLSDLLNFYSENNRLIGHGVYYSLFDARWTERQELWLQKLKEEFRKRKYNHITEHFGFMNTENFHQGVPLPVSLHSKTLAIGKDRLYRLQDAVQVPVGIENLAFSFSLDDVLEQGVFLDKLTDDTDGFLILDLHNIYCQSCNFKVEMQDIIKLYPLDKVKEIHLSGGSWQESAYGKKRVRRDSHDDVIPEDILSVLPSVLSQCRNLEYIIIERLGHTINTEEKKKSFLDDFAKVKMLIDSSDWQTQNKENRNRKEIQFPEHPLEDRILHEEQTLLTKLLFDNVEPESFNHHAFQYFKTEKWDPEMIITARNIIKKWNPY |
| AYY86288.1 | Chryseobacterium indologenes | MMRKPLLGLSMMPEAEFVSAALPLLQNNSVEVLEWSFDTLFNANEPDWLRDLLNFYAENNRLIGHGVYYSLFDAQWTERQEIWLSRLKDEVSKRNYNHITEHFGFMNTENFHQGVPLPVPLHPITLQIGQDRLQRLQDAVNIPVGIENLAFSFCLDDVKEQGVFLEKLIDNSGGFLILDLHNMYCQSCNFDVDIQEIIKLYPLQKVKEIHLSGGSWQESVYGKKMIRRDTHDHTIPEEIFAILPFVLSHCENLEYIIIERLGHTIKTKEEEGDFFSDFMRVKTIIDSSAGHERQGDLWIKKEFELSKYPVEDPVLYEEQSRLTKLLFDGTHVDSIKNENFHYFNTAGWDPEMIFTAQEIIKKWNPY |
| WP_123899531.1 | Chryseobacterium bernardetii | MGKPLLGLSMMAEEEFLSAVLPLLQGNSVEVLEWSFDTFYNSHEPDWLSDLLNFYAENDRLVGHGVYYSLFDARWTERQEIWLKKLKEEFRKRKYNHITEHFGFMNTENFHQGVPLPVSLHPEIVEIGKDRLYRLQDAVQVPVGIENLAFSFSLNDVMEQGIFLDRLTEGIEGFLILDLHNIYCQSCNFEVEMSEIIKLYPLDKVKEIHLSGGSWQESIYGKKLIRRDTHDDEIPEAILSVLPSVLLQCKNLEYVIIERLGNTINTREKKKRFLDDFLKVKALIDSSGLEIQNKNSWAKKELHFPESPLENIILHEEQTLLTKLLFDNVEAECIKNHKFHYFKTENWSPEMILTAQNIIKKWNPIKNDNPLVTTNQII |
| WP_123841941.1 | Chryseobacterium arthrosphaerae | MKKPLLGLSMMPEADFVSAILPLLQSNSVEVLEWSFDTFYEAKEPEWLNELLNFYSENQRLIGHGVYYSLFDALWTERQENWLKKLKEEVRKRKYSHITEHFGFMNTENFHQGVPLPVSLHPKTLLIGKDRLYRLQDAVELPVGIENLAFSFSIEDVKEQGVFLDNLIADINGFLILDLHNIYCQACNFEVEMEDIIHLYPLDKVREIHLSGGSWQESVYARKMIRRDTHDDLIPEEIFSVLPAVITQCNNLEYVIIERLGHTIKTDQEKRAFLDDFTRVRSIIDDSEIKTGKKIRWGQKETGFPNSPVEDDALYEEQSRLTKLLFDSSGAEMIKEQRFHYFKTENWDPDMILTAQNIIKKWNPY |
| WP_124634758.1 | Taibaiella sp. KBW10 | MDTPKILPTVACNLDADILSASLPLLEEGRVAAIEWSFDTLFKRNHILPEWFQELLETYSQANRLIGHGVFFSLFSGRLLPEQKTWLQHLETVSKQYTFDHITEHFGFMTGADFHKGAPMSIPYSKSTVAIGRDRLSRIQQACQCPVGLENLAFSYSLEEVKKHGDFLNELIEPLNGFIILDLHNFYCQLHNFNIPFDTLIALYPLDRVREIHISGGSWQQSIVNEQKQIRRDTHDGAVPQMVFELLQKALPLCPNLKYVVMEQLGIGLQTEADRLQFYRDFLQMEYLVMQYNEGHVYQPHNTFMPSGSLDTTRVVEDEYLYAEQQILSFILEHAKDHEDALHQMQQTALRQSSWDIEHWQPEMIDAALRIAQKWK |

| | | |
|---|---|---|
| WP_123939944.1 | Chryseobacteriu m sp. G0186 | MGKPLLGISMMAEAEFVSAILPLLQTNLVDVLEWSFDTMNTDEPDWLSNLLDFYAGNNRLIGHGVYYSLFDARWTERQRLWLEKLKNEVHRRNYNHITEHFGFMNTSNFHQGVPLPVSLHPKTLQIGKDRLYRLQEAVQIPVGIENLAFSFSIDDVMEQGVFLDKLTEDTNGFLILDLHNIYCQACNFEVEIEDIIKLYPLDKVREVHLSGGSWQESIYGKKMIRRDTHDDAIPEDILTVFSSVLSQCKNLQYVIVERLGHTIDTEEKKENFLNDFSKVKILIEASDQCIQQKKNWTKKEIRFPKAPLEDLVLYEEQVRLTKLLFDRAEVASIRNQEFYYFKTENWDQEMILTAQNIIKKWNPY |
| RTL44676.1 | Candidatus Melainabacteria bacterium | MKPAQLKPMLGVSLMPSDDFRKASRSLFESNQIQVIEWSFDFSWLGNEIEPWCTEIIDRFSSSNNLIGHGVTLSPLSANFSKRQENWLARVRDEFKTRNYVHASEHFGFSEAGPISHGAPIAVPMTAESLRTGTEMLKRYAEATQCPVGLENLAFAFSLDDVKRQGEFIDRLISAVDGFLVLDVHNLYCQVANFDVPITDLLNSYPLTKVREMHLAGGSWSESESRKRFAVRRDTHDDSVPQDVFNFAALGLKLCPNIEFVILERLGWTMLDEEAQAGFREDFETLAEILDATYSQSPLLNAEALFSNPDWRPVLRESFLPGSTEKQCLPEYQNRLLELLSSNKPSAEIFELLKSDPAFENYQNYVSNFDPDMVAVACELMRKWARRTEC |
| WP_123907925.1 | Chryseobacteriu m sp. G0162 | MGRPLLGLSMMAESEFISAVLPLLQSNSIEVLEWSFDTFYNTEEPVWLSELLNFYSENNRLIGHGVYYSLFDALWTERQELWLEKLKEEFHKRKYKHITEHFGFMNTENFHQGVPLPVSLHSKTLEIGKDRLYRLQDAVEVPVGIENLAFAFSLDDVTEQGVFLDKLTEDIDGFLILDLHNIYCQSCNFNVDMRDIIKLYPLNKVREIHLSGGSWQESVYSKTPVRRDTHDDAIPKEILSILSSVVFQCKNLEYIIIERLGHTINTEEKKKNFLDDFAQVRTLIDSSDWEGADKDLWNKKKVQFQKNPLEDMILHEEQTLLTKLLFDNAGVASIKNHEFHYFKTKDWDPEMILTAQNIIKKWNPY |
| WP_124531036.1 | Chryseobacteriu m sp. KBW03 | MRKPLLGVSMMAEADFVSAILPLLQNNSIEILEWSFDTLYYTHEPDWLCNLLNFYAENDRLIGHGVYYSLLDARWTARQEKWLEKLKEEVQQRKYNHITEHFGFMNTDNFHQGVPLPVSLHPKTLQIGKDRLYRLQEAVDIPVGIENLAFSFSVDDVKEQGVFLDKLTEDSNGFLILDLHNIYCQSCNFGVEMQEIINLYPLDKVKEIHLSGGSWQESVYGKMPVRRDTHDDVIPEDIFSVLPAVLTQCQNLEYVIIERLGHTLKTEEERSNFLYDFNKVKTIIEASDWKNRRENSWTKKEIQLTTEKPVEDLVLYEEQSRLTRLLFDNVGVAVIKDQDFHYFKTENWDPEMILTAQNIIKKWNPY |
| WP_123870988.1 | Chryseobacteriu m bernardetii | MGKPLLGLSMMAEAEFLSAVLPLLQGNSVEVLEWSFDTFYNSQEPNWLSDLLNFYAENDRLVGHGVYYSLFDARWTERQEIWLKKLKEEFRKRKYNHITEHFGFMNTENFHQGVPLPVSLHPEIVEIGKDRLYRLQDAVQVPVGIENLAFSFSLNDVMEQGIFLDRLTEDIEGFLILDLHNIYCQSCNFEVEMSEIIKLYPLDKVKEIHLSGGSWQESIYGKKLIRRDTHDDEIPEAILSVLPSVLLQCKNLEYVIIERLGNTINTREKKKRFLDDFLKVKALIDSSGLEIQNKNSWTKKELHFPESPLENIILHEEQTLLTKLLFDNVEAECIKNHKFHYFKTENWSPEMILTAQNIIKKWNPY |
| WP_123860487.1 | Chryseobacteriu m nakagawai | MGRPLLGLSMMAESEFISAVLPLLQSNSIEVLEWSFDTFYNTEEPVWLSELLNFYSENNRLVGHGVYYSLFDALWTERQELWLEKLKEEFHKRKYKHITEHFGFMNTENFHQGVPLPVSLHSKTLEIGKDRLYRLQDAVEVPVGIENLAFAFSLDDVTEQGVFLDKLTENIDGFLILDLHNIYCQSCNFNVDMQDIIKLYPLDKVREIHLSGGSWQESVYSKTPVRRDTHDDAIPKEILSILSSVVFQCKNLEYIIIERLGHTINTEEKKKSFLDDFEQVRTLIDSSGWERENEDLWNKKKVQFQKAPLEDMILHEEQTLLTKLLFDNAGVASIKNHEFHYFRTKDWDPEMILTAQNIIKKWNPY |
| RTZ46797.1 | Chryseobacteriu m arthrosphaerae | MKKPLLGLSMMPEADFVSAILPLLQSNSVEVLEWSFDTFYEEKEPEWLDELLNFYSENQRLIGHGVYYSLFDALWTERQENWLKKLKEEVRKRKYSHITEHFGFMNTENFHQGVPLPVPLHPKTLLIGKDRLYRLQDAVKLPVGIENLAFSFSIEDVKEQGVFLDNLIADINGFLILDLHNIYCQACNFEVEMEDIIRLYPLDKVREIHLSGGSWQESVYARKMIRRDTHDDLIPEEIFSVLPSVITQCNNLEYVIIERLGHTIKTDQEKRAFLDDFTRVRRIIDDSEIKTGKKIRWGQKETGFPNSPVEDDALYEEQSRLTKLLFDSSGAEMIKEQRFHYFKTENWDPDMILTAQNIIKKWNPY |
| WP_126650399.1 | Chryseobacteriu m aureum | MSKPLLGISMMAEADFVSAVLPLLQTDSVDVLEWSFDTLYHSYEPDWLHDLLNFYAENNRLIGHGVYYSLFDGRWTGRQEEWLKKLQEEVSLRKYNHITEHFGFMNTENFHQGVPLPVSLHPQTLEIGKDRLYRLQEAVNIPVGIENLAFSFSVDDVKEQGVFLDKLTENTDSFLILDLHNIYCQSCNFQVGLQELIELYPLDKVKEIHLSGGSWQESAYGKKQVRRDTHDDVIPDEIFSVIPYVLMKCQNLEYVIIERLGHTLKTQEEKDNFVMDFNEVKNIIEASAWKKKEKKSWNKKEVRLSEKPLEDLVLYEEQSRLTRLLFDNIDAATFKGVDFHYFKTENWDPEMIVTAQNIIKKWNPY |
| NSL90190.1 | Chitinophaga solisilvae | MEASKIEAIEWSFDALYKVKDIPDWFVELLTAFSNENRLIGHGIFFSLFSGRWRQEQEQWLQHLRKTSTHFRFDHITEHFGFMTGKDFHHGAPLNIPYTKSTLNIGVDRLKRIHDACRCPVGLENLAFSYSMDEVKRHGAFLDELLEPVNGFIILDLHNLYCQLHNFDITFGEMIILYPLHKVREIHISGGSWEASPAAPERMVRRDTHDDAVPEEVFQLLQKTIPLCPNLQYVVLEQLGTGLETDTSRTLFRQDFGRMQRIVHEQQPQTGSCPVNSFMPDNLLISAVPAADETLFQQQLELSAILESASSCEAAVHALQHSSLAHSDWLIEQWQPYMIETAVSIAQKWKKI |
| WP_129036615.1 | Chryseobacteriu m sp. CH21 | MSKPLLGISMMAEADFVSALLPLLQSNSVDVLEWSFDTLYHSNEPGWLRDLLNFYAENNRLIGHGVYYSLFDARWTSRQEEWLRKLKEEVALRKYNHITEHFGFMNTENFHQGVPLPVSLNPKTLEIGKDRLYRLQEVMNIPVGIENLAFSFSVDDVKEQGVFLDELTEDTNGFLILDLHNIYCQSCNFEIGLQEIIDLYPLDKVKEIHLSGGSWQESAYGKKQVRRDTHDDVIPDDIFAVLPSVLTQCHNLEYIIIERLGHTLKTDNEKDDFLNDFDKVKMIIAASDWKEKEKKNWNAKDIKLPEEPLEDLVLFEEQSKITQLLFENIDVAVIKDQDFHYFKTENWDAEMILTAQNIIKKWNPY |
| WP_126247406.1 | Chitinophaga rhizosphaerae | MPRLLSTVACNLDPQLLGACLPLFESGEADAIEWSFDALYRLHELPEWFHGLIGAFSEGDRLIGHGIFFSLFSGKWHPEQQAWLKELSSICDRYRFDHITEHFGFMTGQDFHHGAPLNIPYTAATLRIGQDRLQRIAEACGRPVGLENLAFSYSADEVRRHGDFLDNLLAPVNGFLILDLHNVYCQVENFDIDFDTLAAMYPLHRVREIHISGGSWEPSADGSQNIRRDTHDGAVPEKVFQYLERIIPQCPHLRYVVMEQLGAGLQTEASKSAFRADFRRMADIAGTWNNPASDESIHPFTPPASRLQSAPAEDPDLHRQQTELSSILENAASLEDALRSLETSSLAQSDWQVEQWSPYMVETAMRIARKWMKKD |
| WP_126693540.1 | Hymenobacter gummosus | METAPRICSAVACNLDADILAAVYPLLEEGRVEALEWAFDALYWAEQVPEWFEALLQAYAQAGRLIGHGVFFSLLSGRWTPEQQQWLQDLRQVASRYQFDHITEHFGFFTGQNFHHGAPLPIPYSAAALRIGQDRLLRIADACRCPVGLENLAFAYALDEVRRHGEFLDQLLQPVNGFIILDLHNLYCQLHNFEVPYEELIALYPLDRVREIHISGGSWEDSALTPARRIRRDTHDEAVPDEVFELLRLTLPRCPNLKYVVLEQLGPGLRTEASRLRFQADFGRMEALVQPQRAGLAAGPGNAFRPPLAPLTAPAAEDGQLHAQQRQLSDILETAGSYEEVQRRLHASALAGTDWRVESWAPHMLETAVNIARKWKQ |
| WP_129020704.1 | Edaphocola flava | MSRTILPTIACNLDADILSASLPLLEEGVVDALEWSFDALFRQDHVIPEWFHDLLTAYSNEGRLIGHGVFFSLFSGRLLPEQAAWLQRLKQLSRQYRFDHITEHFGFMTGSDFHKGAPMSVPYTSGTVAIGRDRLQRIQDAAQCPVGLENLAFAYSLDEVKRQGAFLEQLIEPVNGFIILDLHNVYCQLHNFNIPFEVLIQAYPLDRVREIHISGGSWETAVTDQSRSIRRDTHDGRVPESVFELLEHTLPLCPQVKYVVMEQLGTGLQTEAAKISFRKDFQQMQKITEQFLEHHSRMDAVPNNFLPEYSLPPGQVIEDEYLYGEQQILSYILEHAAGHAEAMEQLQHTALRNSVWGIEKWQPDMLETAIRIAQKWK |
| QBA23550.1 | Chryseobacteriu m indologenes | MRKPMLGVSMMAEADFLSAVLPLLQDNSIEVMEWSFDTLYHTNEPDWLRDLLNFYAENNRLIGHGVYYSLFDARWTARQEEWLNKLKEEVSLRKYNHITEHFGFMNTENFHQGVPLPVSLHSKTLQIGKDRLFRLQEAVNIPVGVENLAFSFSIDDVKEQGVFLDKLIDDTDGFLILDLHNIYCQSCNFEVGLQEIINQYPLDKIKEIHLSGGSWQESAYGKKQVRRDTHDDVIPEEILSVLSSVLTQCPNLEYIIIERLGHTLKTEEERNDFLDDFNKVRTIIETSDLKFKEKKRWCKKEVKLSEKPLEDLLLYEEQSELTRLLYDNADVKAIQDADFQYFKTQHWDAEMILTAQNIIKKWNPY |
| RYD58560.1 | Sphingobacterial es bacterium | MHEIFSSIACNLDNDMLSAALPLLQEERVEAIEWSFDALYDHRQIPPWFDELLHAFSNEGRLVGHGVFFSLFSGGWSVGQQAWLDHLKMLSRTFRFDHITEHFGFMTGRDFHSGAPLNIPYTNTNLQIGQSRLKKIYDACECPVGLENLAFSYTLDEVKRHGAFLEELLRPINGFIILDLHNLYCQLHNFSIPFDELIDLYPLDRVREIHISGGSWEGSEIEPGRQVRRDTHDDAVPEEVFYLLKRTIDKCPHLKFVVMEQLGNALKTTESRQQFYADFVRMDTVVKAKNHSAISASVNNFLPVLPLPEFPVNGDANLYRQQLELSDILENASSYSDALGKLQQSSLAKTEWDIERWKPAMLETAISIAQKWKKT |
| WP_129606978.1 | Spirosoma sordidisoli | MTDIRSSVACNLDSHILRAALPLFADEKVGAIEWSFDALVNPRSMPDWFAELLRTYSGAGRLLGHGVFYSLFSGRWTGAQQQWLDQLKSLSRTYRFDHVTEHFGFMTGADFHKGAPMSIPFTESTLAIGRDRLLRLQDACHCPVGLENLAFAYTVDEVDRHGAFLNQLIEPVNGFIILDLHNLYCQSRNFGVDPDGLLSLYPLYRVREIHLSGGSWAPSAVEPGRTIRRDTHDDAVPADVFRLLKQAIDLCPNLRYVVLEQLGAGLQTDGQQSQFQQDFLWMDHIVQAANQIRNPLINHSFLPRRFHVPDPSPLEDPLLYAQQLELSDILETSVSYAQAQQRLAGSGLARSDWAVETWQPAMLETALAIAQKWKNGFD |
| RYY55266.1 | Chitinophagacea e bacterium | MQKPFSSIACNLDTDILTASLPLLEQSKVEGIEWSFDTLFGSTVIPEWFRELLLTYSRAGRLVGHGVFFSLFTGRWLHGQQQWLDQLRKIATEFPFDHVTEHFGFMTGRDFHQGAPINIPFTGSTLAIGRDRLKRIQDAAGCAVGLENLAFAYSLDEVKRHGDFLQQLLLPVNGFIILDLHNLYCQLHNFNMDFEDLIELYPLDKVREIHISGGSFEDSILKPVRSIRRDTHDDRVPEAVFQLLEQTIPRCPNLRYVVLEQLGTALNSAGSKSGFREDFDRMDRIVKSLPATSNPTDIFLPHKNGLPFSIAEDEVLYQQQRQLSEILETSTDGTEALARLRKSALAGTAWDFENWDPYMLETAVGIARKWK |
| RYZ55261.1 | Sphingobacterial es bacterium | MKAGIAVNADADLLRAAFPLFEQEKVGVVEWAFDTLYDQRTLPPWFEELLGHYSSEGALLGHGVFFSLFSARWSPQQAAWLTALKALSRRFSFDHVTEHFGFMTGSDFHKGAPLNIPYNHTTLAIGQDRLARIQDACGCPVGLENLAFTWSADEAKKQGSFLEKLVTHVNGFIILDLHNLYCQLHNFKLGFEDMARFYPLHLVREIHISGGSWDSVAGRQIRRDTHDDQVPADVFDLLEMAIPLCPNLRFVILEQIGQGLVTGESQMQYQKDFLRLEMMLQHIPVDSGGRLDFSPLLPAFPSVPPEDILLAGQQSELCRMLESTESYEQLQHELKNSGICGTDWAIESWEPEMLETARRIAAKWKNGFFE |
| RZK63166.1 | Hymenobacter sp. | MSTPPAATAAPELRATLACNLDADILTAALPLLQEGRVAAIEWSFDTLYWAEQVPDWFTELLHAYGSQHRLIGHGVYFSLLAGRWSAEQAQWLGQLRQLARQFTFDHITEHFGFFTGQNFHSGAPLPMPYTSATLRLGQDRLRRIQDACQCPVGLENLAFAYSLREVEQHGEFLAKLVEPVNGFLILDLHNLYCQLHNFGASCESLLACYPLDRVREIHISGGSWEDSVLEPTRKIRRDTHDEAVPEEVFQLLALTLPQCPNLKYVVLEQLGHGLKTAVSHERFRQDFGRLEALVQQHQAGRRGAAGQPFLPQRPWQPGPVVVDEQLHAQQQQLSQILETATSVAEAQQRLQASALARTDWHIERWEPHMLETALRIAQKWQQ |
| VFA40327.1 | Chryseobacteriu m indologenes | MMRKPLLGLSMMPEAEFVSAALPLPQNNSVEVLEWSFDTLFNTNEPDWLRDLLNFYAENNRLIGHGVYYSLFDAQWTDRQEIWLSRLKDEVSKRNYNHITEHFGFMNTENFHQGVPLPVPLHPITLQIGQDRLQRLQDAVNIPVGIENLAFSFCLDDVKEQGVFLEKLIESSGGFLILDLHNMYCQSCNFDVDIQEIIKLYPLQKVKEIHLSGGSWQESVYGKKMIRRDTHDHIIPEEIFAILPFVLSHCENLEYIIIERLGPTIKTKEEEDDFFSDFMRVKTMIDSSAGHERQGDLWIKKEFELSKYPVEDPVLYEEQSRLTKLLFDGTHVDSIKNENFHYFNTAGWDPEMIFTAQEIIKKWNPY |
| WP_129056564.1 | unclassified Chryseobacteriu m | MKRPLLGLSMMAEAEFVSAVLPLLQSNSIEVLEWSFDTFYNAEEPDWLSNLLDFYAENNRLIGHGVYYSLFDARWTERQEIWLNNLKEEFHKRNYNHITEHFGFMNTENFHQGVPLPVSLHSKTLEIGKDRLYRLQDAVQVPVGIENLAFSFSLDDVMEQGVFLDKLTEDTNGFLILDLHNIYCQSCNFGVEMQEIIKLYPLHKVKEIHLSGGSWQESVYGKNPVRRDTHDDAIPEQILSVLSSVLSQCENLEYIIIERLGNTIDTDEKKKSFLNDFETVRRLIDLSAGSGIENNNFWTKKDVLLSENPLEDLLLHEEQTLLTKLLFDNAGPELVKDQKFHYFKTKNWDSEMILTAQNIIKKWNPY |
| RYD95878.1 | Sphingobacterial es bacterium | MGRRIEGIIPGIATNLDAAMLSASLPLLEAGKVGGLEWSFDALYARDYQVPDWFHELLLAYSEAQRLVGHGVFFSLFSGRFSAGQQAWLERLRALTAVYQFDHITEHFGFMTGRDFHKGAPLSVPYTANALAIGRDRISRIADAAQCPVGLENLAFAYDLDEVKIQGRFLDELLNPVNGFMILDLHNLYCQLHNFNMDITTLLDLYPLDKVREIHISGGSWEPHPGAHLQQVRRDTHDDAVPAAVFELLPIALRRCPNLKFVMLEQLGTALTTPVESTQFQEDFNTMQTIIEQYNREQSWPRNIAGFEPASLQLTPAALVDEQLFGHQQIISHILETAADYQDAMAKMQYTSLAGSDWHIEKWQPAMVDTAIHIAQKWK |
| RYF71102.1 | Cytophagaceae bacterium | MIRSTLACNLETNLILASLPLFEAERVEALEWSFDTLYAHEQLPDWFIDLLTTYSEAGRLIGHGVFFSLLSGRWTRDQENWLERLRALTRDYQFDHITEHFGFMTGADFHKGAPINIPCTPAALALGQDRLKRIQDACGCPVGLENLAFAYTLDDVKRQGDFLRQLIQPVNGFIILDLHNLYCQSCNFSIMHDDLLALYPLDCVREIHISGGSWSKSQTQPDVLIRRDTHDDAVPDEVFNWLETTIDQCPNLRYVVLEQLGSGLTSNAQQQMFQADFHRMDTIVQDRNQTTTATTMNSFLPKLPLIINATPLENHRLHNQQVELSAILESSTDVGQAQVRLAQSSLANSEWEIESWEPSMFETAVAIARKWKDGFVSRK |
| RZK32697.1 | Hymenobacter sp. | MSKEPSILAALACNLDADILAAAFPLLAEGRVEAIEWSFDTLYWTEQMPGWFAELLGAYSAQNRLLGHGVFFSLLSGKWTAEQQQWLGHLQQLTQQFSFAHITEHFGSFTGQNFHSGAPLPVPYTPATLRIGQDRLRRIYEACQCPVGLENLAFAYSLAEVQHHGEFLGKLLEPVNGFLILDLHNLYCQLHNFAVSEETLLALYPLDRVREIHISGGSWEESALAPGQKIRRDTHDAGVPEEVFHLLERSMPQCPHLKYVVLEQLGTGLKTEASRTQFRQDFRRMARLVKQCRKGWPTPPTQSFRPLHPHQPGPVAEDAQLHEQQTQLSHILESASSFAEAQRQLHNSALASTAWQLETWEPHMLETAIRIAQKWKG |
| TAL31200.1 | Alphaproteobacte ria bacterium | MPFAGLSIFGDARYLDAARPLFAAGAVDAIEWSVDSWSHEEAADEIDTLLDSYAKRGLLIGHGIHYPLLAVKADKLRANWLKQLKPVVQARKYAGVSVHFGFSTGWKLREGAPLPVPLCKEALDLGKKAMVALQKTVKVPVGIENLALAFSKQDVASQGVFIDEMLRAVDGYQLLDLHNIYCQSVNFGVPVLDLVKTYPLSRVLEIHVSGGSWSEHGKMKVRRDTHDGRVPQELFDALPAVLKLCPRARFVIFEKLPQSFKTKADTTGFRDDFMQLKALVPREKAAA |
| WP_131433417.1 | Chryseobacteriu m gleum | MRKPMLGVSMMAEADFVSAILPLLQNNSIDVLEWSFDTLYHPNEPDWLRDLLNFYAENGRLIGHGVYYSLFDSRWTERQDEWLKKLKEEVRVRNYNHITEHFGFMNTENFHQGVPLPVFLHPKTLQIGKDRLCRLQEAVDIPVGIENLAFSFSVDDVKEQGVFLDQLTEDTNGFLILDLHNIYCQSCNFGVEMKEIVDLYPLDKVKEIHLSGGSWQESDYGKKQVRRDTHDDVIPQDILSVLPSVMEQCQNLEYIIIERLGHTVKTEEEKVSFLNDFNTVRKIIEISDLKQAERSWTKKEMILPEKPLEDRMLYEEQSELTKLLFENADAAVIKNHDFHYFQTENWDPEMILTAQNIIKKWNPY |
| WP_136580099.1 | Niastella caeni | MTTLLATAACNLDSDILNACYPLFAEERVQAIEWSFDVLFNRNVLPPWFEELLHAYSQAKRLTGHGVFFSLFSGKWLPEQAQWLETLRATTKRFPLEQVTEHFGFMTGKDFHAGAPLTIPYSATTLRIGRDRLRRIQDAAQCPVGLENLAFAYTSDDVKRHGYFLHDLIEPVNGFIILDLHNVYCQLENFGIDANELLALYPLHKVREIHISGGSWEPVACDRDRRIRRDTHDDAVPAEVFSLLEMAIDRCAECKYVVLEQVGSGLKTPASRQQFYNDFLQMEKVLQQKNKSRHGNSQNNFLPLHVLNTNPPPEDEFLFQQQRELSRILESDIPYNEAVTVLRRSSLANSVWNIENWNPAMLETAMQIARKWKGE |
| TGE29932.1 | Hymenobacter metallicola | MEVGRVEALEWSFDTLFWAEQVPEWFTELLLTYSQQQRLVGHGVYFSLLSGRWTAEQQQWLQQLQELTKRFAFDHITEHFGFFTGQNFHSGAPLPIPYTRSALHLGQDRLQRIQQACGCPVGLENLAFAYSLDEVQRHGEFLDELVEPVNGFLILDVHNLFCQLHNFAVPYEELIRRYPLDRVREIHISGGSWDDSGPTPGRPVRRDTHDG |

| | | |
|---|---|---|
| | | AVPEEVFQLLEQTMPRCPNLRYVVLEQLGNALKTDAQHTQFRADFCRMEALVTRHRSRAAGRAGQLFLPLQPGAAGPVAEDESLYAQQQQLSRILETAPTFEEARRQLQASPLAHSAWKIEEWEPHMLETARRIAQKWQ |
| WP_135531367.1 | Hymenobacter wooponensis | MPNQPVKADTSEICASIACNLDANILTAAFPLLEEGRVEALEWAFDTLFWAEQMPEWFTELLQAYAAQHRLVGHGVYFSLLSGRWTAEQQQWLQHLRELSKQFAFDHVTEHFGFFTGQNFHSGAPLPIPYGSNTLRIGQDRLRRMQEACGCPVGLENLAFAYAPDEVKRHGEFLEKLVEPVNGFLILDLHNVFCQLHNFSLPYEELVTLYPLERVREIHISGGSWENSGQVPGRQIRRDTHDESVPEEVFQLLELTLPRCPNLKYVVLEQLGTGLRTELSQANFRADFHRMQELVNRHRSTTLRRTTNLFLSQQPLSSGPVAEDAQLHAQQQQLSHILENAPSFEAARQQLQTSSLAYSAWKIEEWEPHMLETAISIAQKWKD |
| WP_135435099.1 | Hymenobacter fodinae | MRAPAQSAPAPEPGIFSALACNLDANILTAAFPLLEEGRVEALEWSFDTLFWTKQVPAWFSDLLQAYAAQHRLVGHGVYFSLLSGRWTAEQQQWLRHLRELTKQFAFDHITEHFGFFTGQNFHSGAPLPIPYGANTLRLGQDRLRRMQEACACPVGLENLAFAYSLEEVKRHGEFLDKLVEPVNGFLILDLHNVFCQLHNFSVPYEELMDLYPLERVREIHISGGSWEDSGQVPGRQIRRDTHDEAVPEEVFQLLERTLPRCPNLKYVVLEQLGTGLQTELSQAQFRADFSRMQALINQHRSTSSHPAPEGFWPQQLLPLGPVAEDAELEAQQQQLSNILETAPSFEAARQQLHASSLANTAWNIEGWEPHMLETAISIAQKWKRKPV |
| QCE40223.1 | Psychroserpens sp. NJDZ02 | MRYGIMKNNKPKLGLSLMPNPEFIAAVLPLFEAAQVEVIEWSFDTILDKTYQPEWLPLVLKEYGDNNRLLGHGVYYSLLDANWSANQDNWLKKAKQETLTYNYTHISEHFGLMSATNAHNGFPLPFDLSDTILNIGIDRLKKLQNTVQLDVGIENLALASNASDILKQGEFLNKLVTPVNGFIILDLHNIYCQSENFDIDMLTIITSYPLDLVKEIHISGGSWDQDTSLTKPIRRDTHDGCIPEAILNILPEVLQRCPVLEFIIFERIGDAFKDQKDGIDFRADFNKIKAIIDHTNFSTTPRHWSLNKHNLDQPLIDLELVNQQNTLRQNIQLDQPNAHPEWDTDLWKTATKLYKKWNK |
| THV62431.1 | Chryseobacterium candidae | MKKTMLGVSMMAEADFVSAVLPLLQNNSIDVMEWSFDTLYHANEPDWLRDLLNFYAENNRLIGHGVYYSLFDARWTERQEEWLKKLKEEVNLRKYNHITEHFGFMNTENFHQGVPLPVSLHPKTLQIGKDRLYRLQEAVNIPVGIENLAFSFSIDDVKEQGVFLDKLTEDTNGFLILDLHNIYCQSCNFELEMQEIIDLYPLDKVKEIHLSGGSWQDSVYGKKQVRRDTHDDVIPRDILSVLPSVLTQCENLEYVIIERLGHTLKTDEERDHFLDDFSKVKTIIETSELNGKEKSNWSKMKVELSEKPLEDPVLYEEQSRLTQLLYDNADVETIKDLDFYYFKTESWDAEMILTAQNIIKKWNPY |
| QCR24856.1 | Pontibacter sp. SGAir0037 | MEEGRVEALEWSFDTLYWAEQVPDWFSELLHTYSGEERLVGHGVFFSLVSGKWTKEQQQWLEHLKQVCRQFHFDHITEHFGFFTGQNFHYGAPLNIPYTSSTLRIGQDRLMRISDACQRPVGLENLAFSYSLDEVKQHGHFLEKLVEPVNGFIILDLHNLYCQLHNFSVSFEELMQLYPLHRVREIHISGGSWESSVLDPAIRIRRDTHDDAVPAEVFQLLEYALPLCPELKYVVLEQLGNGLKLESSRIQFRQDFLKMEELVKRHTTTRKPAAKNLFLPSHPLQPGAAAEDKTLQAQQYTLSHILETATSYEEALQLLRSSSLANSEWQIESWAPHMLETAISIAQKWRK |
| WP_137903941.1 | Chryseobacterium sp. 2VB | MRKPMLGVSMMAEADFLSAVLPLLQDNAIEVMEWSFDTLYHTNEPDWLRDLLNFYAENNRLIGHGVYYSLFDAIWTARQEEWLKKLKEEVSLRKYNHITEHFGFMNTENFHQGVPLPVSLGPETLQIGKDRLYRLQETVDIPVGVENLAFSFSIDDVKEQGVFLDKLIEDTDGFLIFDLHNIYCQSCNFEVGLQEIINLYPLDKIKEIHLSGGSWQESAYGKKQVRRDTHDDVIPEEILSVLPSVLTQCPNLEYIIIERLGHTLKTEEERNDFLDDFNKVRTIIETSDLKFKENKRWCKKKVKLFEKPLEDLLLYEEQSKLTRLLYDHADVKAIKDLDFHYFKTQNWDAEMILTAQNIIKKWNPY |
| WP_131957160.1 | Dyadobacter psychrotolerans | MSTIFSSVACNLDANILLASLPLFESDQIEAIEWSFDALFKNDELPLWFNNLILEFSSHNRLVGHGVYFSLFSGKWMPEQHDWLKKLKVLAEKYRFDHITEHFGFMTGENFHKGAPMSIPLTAGTMSLGIDRLHRIQDACNCPVGLENLAFAYSIDEVKKQGDFIGKLVESINGFIILDLHNLYCQIHNFDVDYQNIIDLYPLGRVREIHISGGSWDKQLTRTGKPIRRDTHDDAVPEEVFRILSAVIPRCKNLKYVVLEQLGTALDSEQKRKAFRKDFLKMDAIVKESAFQFYNNPVNDFRPYQTIIPHPVPLEDLSFYNQQLQLSNILETAVDQENARHLLSKSSLRDSDWEVEKWESGMIQTAIAIAQKWKEGF |
| WP_138367034.1 | Dyadobacter luticola | MSEIYASVACNLDIPILQASLPLFEEGKIEGIEWSFDALFKWQEIPGWFTDLVHEFSENNRLVGHGVYFSLFSGKWSPEQQQWLEKLRKLSGDFHFDHVSEHFGFLTGDDFHKGAPIGIPFTNSTLTIGRDRLRRIQDACQCPVGLENLAFSYSLEEVKRHGEFLDALVEEVNGFIILDLHNLYCQSHNFNIPFDQMMRLYPLDRVREIHISGGSWDDQLSGFEKIVRRDTHDESVPEEVFEYLKLAIPQCKNLKYVVLEQLGAALETSDQQETFRNDFLKMQAIIQHPGTKSAQSVSDFVTKYFSPLSPIPPEDPHLYRQQKQLSDILENAADSFEAFKMLQNSTLANSDWRVENWEPVMLETAIAIARKWKNGF |
| TLX25343.1 | Chryseobacterium indologenes | MMRKPLLGLSMMPEAEFVSAALPLLQNNSVEVLEWSFDTLFNTNEPDWLRDLLNFYAENNRLIGHGVYYSLFDAQWTERQEIWLSRLKDEVSKRNYNHITEHFGFMNTENFHQGVPLPVPLHPITLQIGQDRLQRLQDAVNIPVGIENLAFSFCLDDVKEQGVFLEKLIDNSGGFLILDLHNMYCQSCNFDVDIQEIIKLYPLQKVKEIHLSGGSWQESVYGKKMIRRDTHDHTIPEEIFAILPFVLSHCENLEYIIIERLGHTIKTKEEEDDFFSDFMRVKTIIDSSAGHERQGDLWIKKEFELSKYPVEDPVLYEEQSRLTKLLFDGTHVDSIKNENFHYFNTAGWDPEMIFTAQEIIKKWNPY |
| TQM21133.1 | Chryseobacterium aquifrigidense | MLGVSMMAEADFVSAILPLLQNNSIEVMEWSFDTLYHTHEPEWLRDLLNFYAENSRLIGHGVYYSLFDARWTERQEEWLKQLKEGVSLRKYNHITEHFGFMNTENFHQGVPLPVSLHSKTLQIGKDRLYRLQETVNIPVGIENLAFSFSIEDVKEQGTFLDQLTEDTNGFLILDLHNIYCQSCNFEAGMQEIIDLYPLDKVKEIHLSGGSWQESVYGKKQVRRDTHDDIIPEEILSVFPSVMAQCPNLEYIIIERLGHTLKTEEERNNFLEDFNKVRTIIETSDLKLKEKSNWCKKEMKLPEKPLEDLLLYEEQSKLTRLIYDNAEVETVKDQDFHYFKTENWDSEMILTAQNIIKKWNPY |
| WP_138991310.1 | Larkinella sp. C7 | MANLHSALACNLDANILQASLPLLEAEQVDAIEWAFDTLYRTNKLPDWFDQLLRIYSDAGRLVGHGVFFSLFSGHWTADQQAWLKHLKILSSHYRFDHVTEHFGFMTGADFHTGAPLSVPLNSSTLSIGKDRLKRIQEAARCPVGLENLAFAYSLDDVKRQGDFLHQLIYPVNGFLILDLHNLYCQSHNFRLGYEQLLRLYPLDRVREIHLSGGSWEPATSEPGRTVRRDTHDEAVPTELFGWLEKAIDQCPNLKYVVLEQVGSGLRTEQARQGFARDFRQMKAIVKGKSNGLPSKLNTFLPGEPIVINELPVEDAELFAQQRQLATILESANDYQHARQLLASSNLAGSDWLAEKWQPGMLETAIAIAQKWRKGFS |
| TVT40773.1 | Hymenobacter setariae | MLAALACNLDADILGAALPLLEEGQVEALEWSFDALYWASEVPAWFVELLHAYSDQQRLLGHGVYFSLLSGRWTSEQQQWLNQLRELTQRYSFAHITEHFGAFTGQNFHAGAPLSVPYSPVSLRIGQDRLARIAEAARCPVGLENLAFAYSLEEVKRQGEFLGELLAPVNGFLILDLHNLYCQLHNFSIGYEELLECYALDRVREIHISGGSWENSQLLPTRKIRRDTHDEAVPVEVFNLLARTMPRCPNLKYVVLEQLGTSLKTEASHVQFRRDFVQMAALVQQHNHSPRSVPNAFLPRWPALPALAVQDDLLHEQQRQLSHILETAASLPEAQQRLRASALARSAWQIEHWEPYMLETALSIAQKWK |
| WP_138506478.1 | Spirosoma lacussanchae | MIDIRSSIACNLDSHILRAALPLFADEKVGAIEWSFDALINQRSMPDWFGELLRTYSEAGCLVGHGVFYSLFSGRWTGAQQQWLDQLRSLSRTYRFDHVTEHFGFMTGADFHKGAPMSIPFTESTLAIGRDRLVRLQDACRCPVGLENLAFAYTIDEVNRHGAFLSQLIEPVNGFIILDLHNLYCQSRNFGVEPDALLSLYPLYRVREIHLSGGSWAPSAVEPGRTIRRDTHDDAVPADVFRLLKQAIDLCPNLRYVVLEQLGAGLQTDGQQSQFQQDFLWMDHIVQTANQIRNSLINHSFLPRRFHVPDSSPLEDPLLYAQQLELSDILETSVSCAQAQQRLAGSGLARSNWAVETWQPAMLETALAIAQKWKNGFD |
| WP_142716532.1 | Chryseobacterium rhizoplanae | MKKPLLGISIMAEADFVSAVLPLLQNNAVDVLEWSFDTLYHTNEPDWLRDLLNFYAENERLIGHGVYYSLFDARWTERQEVWLQKLKEEVSLRKYNHITEHFGFMNTENFHQGVPLPVSLHPKTLQIGKDRLYRLQEAVDIPVGIENLAFSFSVDDVTEQGVFLDKLTEDTNGFLILDLHNIYCQSCNFGVEMQEIIDLYPLDKVKEIHLSGGSWQESVYGKKQVRRDTHDDVIPEEIFSVLPSVLAKCQNLEYSIIERLGHTLKTEEDKENFLNDFNKVKAIIEASDGKKKEKSSWNKWQIQLPEKPLEDLALFEEQSRLTRLLFDGAGAGVIKDQDFHYFKTENWDPEMILTAQNIIKKWNPY |
| WP_144334209.1 | Fluviicola chungangensis | MNQLLPKIRSAVSCNLDNQLLQATLPLFAAEKIEAIEWSFDTLYKVREIPIWFVELLHAFGNEGRLVGHGVYFSLFSGKWTAEQQRWLDHLRKVSSNFRFDHITEHFGFMTGEDFHKGAPISVPYKKETLRIGIDRLQRIREACACPVGLENLAFSYSLEEVQRHGEFLERLIEPVNGFIILDLHNVYCQLQNFELDFETLIRAYPLDRVREIHISGGSWDVSGIQPQRTIRRDTHDDAVPEAVFELLRKTLSVCPNLKFVVLEQLGIALHTTESQELFQQDFDHMQAIVNEFNGRNNLIPDNDFLLKQAVSLNEPVPESLELYRQQLELSSILEQAANYEEATRLLNNSTLAHSAWKTESWSPYMLETALRIAQKWRSGW |
| WP_142722334.1 | Chryseobacterium sp. ON_d1 | MKRPMLGLSMMAEADFVSAILPLMQNNSIDVLEWSFDTLYHTHEPDWLRDLLNFYAENSRLIGHGVYYSLFDAKWTERQEEWLQKLKEEVRLRKYNHITEHFGFMNTENFHQGVPLPVPLHSRILQIGKDRLCRLQDAVNIPVGIENLAFSFSMDDVKEQGIFLDRLTEDTNGFLILDLHNIYCQSCNFEVGMQEIINLYPLDKVKEIHLSGGSWQESVYGKKQVRRDTHDGVIPQDILSVLPSVLKQCQNLEYIIIERLGHTLKTEQEKVDFLNDFNTVKMIIETSGLKLRQGGTWSKKEMKLSEKPLEDLVLYEEQSRLTRLLFDKMGAEAIQGQDFRYFKTENWDSEMILTAQNIIKKWNPY |
| WP_142015289.1 | Chryseobacterium | MRKPMLGVSMMAEADFVSAILPLLQNNSIEVMEWSFDTLYHTHEPEWLRDLLNFYAENSRLIGHGVYYSLFDARWTERQEEWLKQLKEGVSLRKYNHITEHFGFMNTENFHQGVPLPVSLHSKTLQIGKDRLYRLQETVNIPVGIENLAFSFSIEDVKEQGTFLDQLTEDTNGFLILDLHNIYCQSCNFEAGMQEIIDLYPLDKVKEIHLSGGSWQESVYGKKQVRRDTHDDIIPEEILSVFPSVMAQCPNLEYIIIERLGHTLKTEEERNNFLEDFNKVRTIIETSDLKLKEKSNWCKKEMKLPEKPLEDLLLYEEQSKLTRLIYDNAEVETVKDQDFHYFKTENWDSEMILTAQNIIKKWNPY |
| WP_246129921.1 | Chitinophaga cymbidii | MNDRLFSTVACNLDANILSACLPLMEASKMEAIEWSFDALFRVKEVPSWFLELLTLYSNEKRLIGHGVFFSLFSAKWRQEQDEWLKQLKQTCSAFRFDHITEHFGFMTGRDFHHGAPLNIPYTPSTLNIGRDRLKRIQDACRCPVGLENLAFAYSLDEVKRHGDFLEQLLEPVNGFIILDLHNLYCQLHNFRLEFEDIIALYPLDKVREMHISGGSWDASQIDPDITVRRDTHDDAVPEEVFHLLDQTIGRCPHLKYVVLEQLGTGLSTPESRQSFYDDFLKMDNIIKTKKTSRSHPSFLPLQNDPPGEVVEDETLYRQQLELSHILETAGSCTDAMEALSRSSLANSDWQIEKWAPYMIETAVKIAQKWKK |
| WP_149388738.1 | Chryseobacterium panacisoli | MKKPLLGISMMADADFVSAVLPLLQSNAIDVLEWSFDTLYHTNEPDWLRDLLNFYAENNRLIGHGVYYSLFDARWTERQDNWLQKLKEEVSLRKYNHITEHFGFMNTENFHQGVPLPVSLHPKTLQIGKDRLYRLQETVGIPVGIENLAFSFSVDDVKEQGFFLDKLTEDTNGFLILDLHNIYCQSCNFGVKMQEIIDLYPLDKVKEIHLSGGSWQETVYGKKQVRRDTHDDVIPEEIFSVLPSVLAKCQNLEYIIIERLGHTLKTEEDKENFLNDFNKVKTIIEASDWKGRKKNSWNKGQIQLPEKPVEDLALFEEQSRLTRLLFDNVEAAVIKDQDFHYFKTENWDPEMILTAQNIIKKWNPY |
| WP_244620440.1 | Chitinophaga japonensis | MPEILPAVACNLDNNILAACLPLMEESRIAAIEWSFDALFKVKQVPPWFQELLTAFSNEQRLIGHGVFFSLFSAKWRQEQEDWLRHLQQTCTAFRFDHITEHFGFMTGKDFHHGAPLNIPYTPATLAIGRDRLKRIYEACQCPVGLENLAFSYSLEEVKRHGEFLEQLLVPVNGFIILDLHNLYCQLHNFDLALEDILALYPLDRVREIHISGGSWDSSRADPGRSVRRDTHDDAVPQEVFRLLEAVITRCPHLKYVVLEQLGNGLATEESRQCFYNDFGVMEIIIREKSPAAAGPSFQSFLPTSFSPPGPVLEDEMLYRQQLELSGILETAPSYAAAIRRLNSSSLAHSGWEIERWAPYMIETAMKIARKWKRK |
| KAA6432601.1 | Rufibacter glacialis | MQEGRVEGLEWSFDTLFQQEQVPSWFLALLETYGREKRLIGHGVFFSLFSGKWSEHQEAWLRHLTQMSAAYGFDHVTEHFGFMTGKNFHHGAPLPIPYTAATLAIGRDRLARIQEAAQCPVGLENLAFSYTLEEVKRHGAFLEELVEPVNGFLILDLHNLYCQLHNFSLPLEALLPWYPLHRVREIHISGGSWQDSAVLPQRKVRRDTHDSAVPEEVFDLLSATMPLCPQLKYVVLEQLGTGLKSPEDKALFYQDFCRMETLVKQHNASTFPSLTAPNLFLPLAPGKLPAPVEDEALYQQQLLLSSILEESPSLTQAHQRLLASSLANTAWAIEHWEPHLLETALHLAQKWKKPDLSA |
| WP_149072616.1 | Hymenobacter lutimineralis | MTKPPTIFAALALNLDADMLAAAFPLLADGAIGALEWSFDALSGANTPVWFTELLHAFSKEQRLIGHGVFFSLLSARWTTEQQQWLQRLNQLARQFSFDHVTEHFGFFTGQNFHSGAPLPVPYTPATLRLGQDRLCRISEACSCPAGLENLAFAYSLEEVTRHGEFLDKLLTPVNGFLILDLHNLYCQLHNFSISYEELIALYPLEKVREIHISGGSWADSSLLPDKKIRRDTHDDAVPDEVFSLLKATMPRCPHLKYVVLEQLGNGLQTEHSQTRFRQDFLRMQKLVAQQNQESKPTDYHFLPRQPVVSGPVEEDVRLYAQQQQLSHILETAHSYEEAHQRLLLSSLAQSEWQIEQWAPYMLETALGIAQKWQLKPS |
| KAA2242006.1 | Chitinophaga agrisoli | MEASKVEAIEWSFDALFKVKEIPSWFLELLTAFSDEQRLIGHGVFFSLFSGKWLQEQEDWLRQLEHTSTAFRFDHVTEHFGFMTGKDFHHGAPLNIPYTNATLTIGRDRLRRIQAACQCPVGLENLAFSYSLEEVKQHGAFLDELLAPVNGFIILDLHNLYCQLHNFEIPFDTLISLYPLDKVREIHISGGSWEDSLAAPPKKVRRDTHDDAVPDAVFELLENTIDLCPHLKYVVLEQLGNGLITAESKTQFQQDFLRMEAIVAEKNATRSTESIQSFLPAHPTIPAHIAEDATLYRQQLELSAILETAATYNDAVHALQTSSLAHSDWRIETWAPYMIETAMKIAQKWKK |
| WP_150032694.1 | Taibaiella lutea | MKTNIHAAIACNFDANILSAALPLFEQEKIEAIEWSFDALFKVKEIPGWFDELLLAFSKEQRLIGHGVFFSLFSGKWLPEQIAWLQHLQELCKQFQFDHITEHFGFMTGTDFHTGAPLGIPYTATTLAIGRDRLARIQEVCKCPVGLENLAFSYSLEEVKQHGHFLQELLNPVNGFIILDLHNLYCQLHNFNIPFEELIYLYPLELVREIHISGGSWESSAIFPEKQIRRDTHDDSVPEEVFELLEKALPLCPQLKYVVLEQLGIGLKPIESRDQFQKDFLKMESVITSFEKESIVKEFNSFLPGNPLPTGIPIEDEILYAQQLLLSDILENASNYNEALQRIQSSSLKQSAWNIESWQPDMLETAIRIAQKWKKGW |
| WP_149831712.1 | Chryseobacterium sediminis | MKKPLLGISMMPEADFVSAVLPLLQSNAVDVLEWSFDTLYNTNEPDWLRDLLNFYAENNRLIGHGVYYSLFDARWTERQEEWLQKLKEEVSLRRYNHITEHFGFMNTENFHQGVPLPVSLHPKTLRIGKDRLHRLQQAVDIPVGIENLAFSFSIDDVKEQGVFLDKLTEDTNGFLILDLHNIYCQSCNFGVGMQEIIDLYPLDKVKEIHLSGGSWQESVYGKKQIRRDTHDDIIPEEIFSVLPSVFAKCQNLEYIIIERLGHTLKTEDDKENFLNDFNNVKTIIEISDWKEREKSSWGKRQIQLPEKPVEDMALFEEQSRLTRLLFDGAGAIAIKDQDFHYFKTENWDPEMILTAQNIIKKWNPY |
| KAB1228757.1 | Chryseobacterium viscerum | MPGLRNGIKMRKPLLGVSMMAEAEFVSAILPLLQNNSIEVLEWSFDTLYHTHEPDWLCDLLNFYAKNDRLIGHGVYYSLLDARWTARQEQWLEKLKEEVQQRKYNHITEHFGFMNTDNFHQGVPLPVSLHPKTLQIGKDRLNRLQEAVDIPVGIENLAFSFSVDDVKEQGMFLDKLTEDSNGFLILDLHNIYCQSCNFGVEMQEIINLYPLDKVKEIHLSGGSWQESVYGKRPVRRDTHDDVIPEDIFSVLPAVLTQCQSLEYVIIERLGHTLKTEEEKSNFLHDFNKVKTIIEASDWKNRRENNWTKKEIQLAEKPVEDLVLYEEQSRLTRLLFDNVGAAAIKDQDFHYFKTENWDPEMILTAQNIIKKWNPY |
| WP_157814118.1 | Olleya sp. Bg11-27 | MKINKPKLGLSIMPNPEFIAAALPLFESAQVEVIEWSFDTILDKKYQPEWLPLLLKEYGDNNRLLGHGVYYAPLDANWGANQDNWLQKAKLETLAYNYNHLSEHFGVMSSANAHHGFPLPFDLSNTILNIGIDRLKRLQNTVQLDIGIENLALASNVSDILKQGAFIEKLVAPINGFVILDLHNIYCQSINFDLDLLTIITSYPLHLVKEIHLSGGSWDHDHNLTKPIRRDTHDGRIPEVLLDILPEVLQRCPVLEFVIFERLGDTFQNKNDGLEFRTDFNKIQAIIDHTAFSSNARQWTLKKHDLGPPLIDLPLLNKQQELREHIRLETANNHAEWDTNMWITATKLYKKWNA |

| | | |
|---|---|---|
| MXV51032.1 | Hufsiella arboris | MKSDPVTAIASTIACNLDSDILSAAIPLFEESRVEAIEWAFDSLDFSEVPEWFSDLIDAYSKEGRLIGHGVFFSIFSGRWSIQQQEWLDNLKKLSARFKFDHITEHFGFMTGANFHYGAPLAVPFNSKTLTIGRDRLNRIYDACQCPVGLENLAFSYSIDEVKKHGEFLETLVEPVNGFIILDLHNLYCQMHNFSISFDEIIKLYPVDRVREFHISGGSWENSSLSGKRIRRDTHDDAVPDEVYQLLDSALDHCKNVRYVTMEQLGSGLKTEESRSNFYNDFLKMQQIVRNKTSVPNDAENTFFPPKIVPPGDIIEDENLYIEQQQLSYILENSTSYDDAVSRIKSSVLAKSDWGIESWAPHMLETAISIAQKWKKK |
| WP_153391763.1 | Chryseobacterium | MKKPLLGLSMMPEADFVSAILPLLQDQSVAVLEWSFDTVEQMHEPEWLSGLLDFYAENNRLLGHGVYYSLFDARWTERQEVWLEKLKQEIQRRKYNHITEHFGFMNTENYHQGVPLPVPLHPKILQIGKDRLARLQAVVEIPVGVENLAFSFSADDVREQGIFLEKLIEEVDGFLILDLHNIYCQSCNFEIDILDIVTMYPLEKVKEIHLSGGSWQESVYGKKPIRRDTHDDAIPEEVLAVLPEVLSRCPNLEYVIIERLGHTISTNIEKESFFNDFVKTKKIIEGCLFPARKKTVWNRKEYISPEPLEDQLLYHEQTKLTRLLHENGDIAFFKNEDFSYFNVQKWDPEMIETARSIIKKWNPY |
| WP_149639431.1 | Dyadobacter sp. UC 10 | MSRIFSAIACNLDVHILQAAFPLFEMEMVEAIEWSFDSLFKVGTIPDWFIELVKEFSDHNRLIGHGVYFSLFSGKWSQDQQNWLDQLRKLSGEFRFDHVSEHFGFMTGADFHKGAPLGIPFNQSTLAIGQDRLARIQDACNCPVGLENLAFSYSYEEVRRHGDFLNQLVESVNGFIILDLHNLYCQCHNFNIHFEQMLSCYPLERVREIHISGGSWDNGTANRPVRRDTHDEAVPEVVFKYLEKALLQCPNLKFVVMEQLGTALDSGEKQYLFQQDFLQMSHIVKSFEENFPPVPVNSFLPPATFPLHSKPVEDLLLAKQQMQLSAILENASNLDHARGMLNASDLKNSDWHIDQWQPHMLKTAMDIARKWKDGF |
| WP_158978994.1 | Cellulophaga sp. L1A9 | MRKRIMGIKKPKLGIAIAPSLKFLEAALPLFAGEKIEIIEWSFDTLKDAADEPAWLSLLLKEYGEKNRLLGHGVFYSLLDANWSSRQENWLKKVRQETLSHKYCQISEHFGLMSSANAHSGFPLPIQLSNPVLQIGIDRLKRLQATAQVDVGIENLALTANVADILEQGEFLFKLVNPVNGFVILDLHNIYCQSENFNIDMMTIIKSYPLSLVKELHISGGSWDTDPTLSRKIRRDTHDGRIPEVILEILPEVLKICPFLTFIIFEKIEDSFLTEKDGIDFRADFQKIREIIDATVFSVTPKEKKQVAAILGPPVIDVELLQAQVALRESIALGTYQTNSEWDKDMWKVATKLYEKWNNY |
| WP_159477007.1 | Dyadobacter sp. 3J3 | MQKIFTSVACNLDTNILLAALPLFEQDKIQALEWSFDTLFKYEEIPDWFTDLVLEFSKNNRLIGHGVYFSLFSGKWLPSQQEWLEKLKIFTLKFRFDHITEHFGFMTGSDFHKGAPISIPFTPVTLALGKDRLNRIQDACQCPVGLENLAFSYSIEEVRKHGNFLDQLTDEINGFIILDLHNLYCQIHNFDITFKDIIHLYPVEKVREIHISGGSWDERLIGNGKQVRRDTHDQAVPQEVFELLINTIPLCPNLKYVVLEQLGTALGSAESRIMYQEDFLRMDAIVKGAEFNASGHNTFMPLYKTNLQATPLEDLNLYKQQQQLSQILETAVDLHQAQKLLSLSDLRNSAWNVENWEPYMLETAMSIAQKWKNGFD |
| QHT72239.1 | Rhodocytophaga rosea | MEESQVDAIEWSFDTLFNIWHTPGWFARLLTAFSKEERLIGHGVFFSLLSGKWSKEQYQWLSQLKQLSAEFHFDHITEHFGFMTGQSFHQGAPMCVPYSPATLAIGRDRLARIQEACSCPVGLENLAFAASLEQVKQQGEFLERLIEPVNGFIILDLHNLFCQMHNFFMDFEQLLSSYPLERVREIHISGGSWEKSGKQSERIIRRDTHDQAVPPEVFHLLEKTIPVCPNLKYVVLEQLGIALKTQESKAAFYQNFVQMKQIVEDNNQTIANSSENPFIPPSITLPETFVENQELYAQQRELSEILETSSSYEETNRLLKASSLAHTAWNIESWDPAMIETAIHIARKWKK |
| WP_202842763.1 | Chryseobacterium cucumeris | MRKPMLGVSMMAEADFVSAVLPLLQNNSIEVMEWSFDTLYHTNEPEWLRDLLNFYAENNRLIGHGVYYSLFDARWTARQEEWLKKLKEEVSLRKYNHITEHFGFMNTENFHQGVPLPVSLGPETLQIGKDRLYRLQEAVNIPVGVENLAFSFSIDDVKEQGVFLDKLIEDTDGFLILDLHNIYCQSCNFEVGLQEIINLYPLHKIKEIHLSGGSWQESAYGNKQVRRDTHDDVIPEEILSVLPSVLTQCPNLEYIIIERLGHTLKTEEERNDFLDDFNKVRTIIETSDLKFKENKRWCKKEVKLSEKPLEDLLLYEEQSKLTKLLYDHTDVKAIKDLDFHYFKTQNWDAEMILTAQNIIKKWNPY |
| NEQ25955.1 | Microcoleus sp. SIO2G3 | MFKFKRRTPTVGLSLMMESDFFQASQPLFEAGEVEVLEWSFDMGWGVDALPPWVEQLLQFYSQRDRLIGHGVSYSLLSAQPGERQIHWLKSLEEECLDYRYRHISEHFGWMAAGDFYQSAPLPLPLTPETLHLGREHLKRLSDTAKAPVGLENLAFAFGLQDVLQQGEFIERLLEPVDGFLVLDLHNLYCQVHNFQLSADQLLERYPLHRVRELHVSGGSWSVTSKGEQTRTIRRDTHNGTVPEAVFELLALALKRCPSVEAVIFERIGHTLHNEVDIEGFRQDFWRIKQVVKESV |
| WP_161990772.1 | Chryseobacterium viscerum | MRKPLLGVSMMAEAEFVSAILPLLQNNSIEVLEWSFDTLYHTHEPDWLCDLLNFYAKNDRLIGHGVYYSLLDARWTARQEQWLEKLKEEVQQRKYNHITEHFGFMNTDNFHQGVPLPVSLHPKTLQIGKDRLNRLQEAVDIPVGIENLAFSFSVDDVKEQGMFLDKLTEDSNGFLILDLHNIYCQSCNFGVEMQEIINLYPLDKVKEIHLSGGSWQESVYGKRPVRRDTHDDVIPEDIFSVLPAVLTQCQSLEYVIIERLGHTLKTEEEKSNFLHDFNKVKTIIEASDWKNRRENNWTKKEIQLAEKPVEDLVLYEEQSRLTRLLFDNVGAAAIKDQDFHYFKTENWDPEMILTAQNIIKKWNPY |
| WP_164720324.1 | Chryseobacterium gleum | MLGVSMMAEADFVSAILPLLQNNSIDVLEWSFDTLYHPNEPDWLRDLLNFYAENGRLIGHGVYYSLFDARWTERQDEWLKKLKEEVRLRNYNHITEHFGFMNTENFHQGVPLPVSLHPKTLQIGKDRLCRLQEAVDIPVGIENLAFSFSVDDVKEQGVFLDQLTEDTNGFLILDLHNIYCQSCNFGVEMKEIVDLYPLNKVKEIHLSGGSWQESAYGKKQVRRDTHDDVIPQDILSVLPSVMEQCQNLEYIIIERLGHTVKTEEEKVSFLNDFNTVRKIIEISDLKLAERNWMKKEMILPEKPLEDRVLYEEQSELTKLLFENADAAVIKNHDFHYFKTENWDSEMILTAQNIIKKWNPY |
| WP_165446360.1 | Chryseobacterium indologenes | MRKPLLGLSMMPEAEFVSAALPLPQNNSVEVLEWSFDTLFNTNEPDWLRDLLNFYAENNRLIGHGVYYSLFDAQWTDRQEIWLSRLKDEVSKRNYNHITEHFGFMNTENFHQGVPLPVPLHPITLQIGQDRLQRLQDAVNIPVGIENLAFSFCLDDVKEQGVFLEKLIESSGGFLILDLHNMYCQSCNFDVDIQEIIKLYPLQKVKEIHLSGGSWQESVYGKKMIRRDTHDHIIPEEIFAILPFVLSHCENLEYIIIERLGPTIKTKEEEDDFFSDFMRVKTMIDSSAGHERQGDLWIKKEFELSKYPVEDPVLYEEQSRLTKLLFDGTHVDSIKNENFHYFNTAGWDPEMIFTAQEIIKKWNPY |
| WP_162149646.1 | Chryseobacterium indologenes | MRKPLLGLSMMPEAEFVSAALPLLQNNSVEVLEWSFDTLFNTNEPDWLRDLLNFYAENNRLIGHGVYYSLFDAQWTDRQEIWLSRLKDEVSKRNYNHITEHFGFMNTENFHQGVPLPVPLHPITLQIGQDRLQRLQDAVNIPVGIENLAFSFCLDDVKEQGVFLEKLIDNSGGFLILDLHNMYCQSCNFDVDIQEIIKLYPLQKVKEIHLSGGSWQESVYGKKMIRRDTHDHTIPEEIFAILPFVLSHCENLEYIIIERLGHTIKTKEEEDDFFSDFMRVKTIIDSSAGHERQGDLWIKKEFELSKYPVEDPVLYEEQSRLTKLLFDGTHVDSIKNENFHYFNTAGWDPEMIFTAQEIIKKWNPY |
| WP_160139545.1 | Chryseobacterium sp. c4a | MGKPLLGLSMMAEAEFVSAVLPLLQSNSVEVFEWSFDTFYNAEEPGWLSDLLNFYAGNNRLIGHGVYYSLFDALWNERQEIWLGKLRDELRKRKYNHITEHFGFMNTENFHEGVPLPVSLHSKTLEIGKDRLHRLQDAVQVPVGIENLAFSFSLDDVMEQGLFLDKLTEDSDGFLILDLHNIYCQACNFKVGMEDIIKLYPLNKIKEIHLSGGSWQESIYGKKPIRRDTHDDIIPEEILSILPFVLSQCKNLEYVIIERLGHTIDTEEKRSDFLDDFRKVKMIIDSMDWKIFDRNSWIKKEIKISKKPLEDKGLYDEQTLLTKLLFDHVNVESIKNHEFHYFKTKDWDPEMILTAQNIIKKWNPY |
| WP_165748299.1 | Cellulophaga sp. Z1A5H | MGNKKPKLGIAIAPSLKFLEAALPLFAEEKIEIIEWSFDTLKDAADEPVWLSLLLKEYGEKNRLLGHGVFYALLDANWSSRQENWLKKVRQETLSHKYCQISEHFGLMSSANAHSGFPLPIPLSNPVLQIGIDRLKRLQATAQVDVGIENLALTANVADILEQGEFLFKLVNPVNGFIILDLHNIYCQSENFNIDMMTIIKSYPLSLVKELHISGGSWDTDPTLSRKIRRDTHEGRIPEVILEILPEVLKICPFLAFIIFEKIEDSFLTEKDGIDFRADFQKIREIIDATSFSVTPKEKKQVAIILGPPVIDVELLQAQVALRESIALDTYQTNSEWDKDMWKVATKLYEKWNN |
| NJK30207.1 | Acaryochloris sp. SU_5_25 | MPPLVGLSLMPQADFMAAAQPLFRAGEVEILEWSFDMGWGNVSLPPELPPLLDQFSQCHCLLGHGVSYSLLSAGIDRSHWLACLQAECQDYSYRHISEHFGWMATRHFSRSVPLPLPLLPATLKLGCDHIQRLAEVAQVPVGLENLAFAFGRQDVDQQGEFLDRLLEPVDGFLLLDLHNLYCQMHNFQLSASEILNLYPLGRVRELHLSGGSWSQSGTAKIRRDTHDHAVPEPVFDLLALALQECSHVEAIILERLGNTFTTDAEKIGFRQDFARIKEIVYSHRSSLSRKG |
| WP_167358395.1 | Chryseobacterium culicis | MNKPLLGISMMAEADFVSALLPLLQSNSVDVLEWSFDTLYHNHEPVWLCDLLNFYADNERLIGHGVYYSLFDARWTERQEDWLKKLKEEVALRRYNHITEHFGFMNTENFHQGVPLPVSLHAKTLEIGKDRLYRLQDAVNIPVGIENLAFSFGIDDVKEQGAFLDKLIEDTDGFLILDLHNIYCQACNFGIEIQEIMDWYPLDKVKEIHLSGGSWQESVYGKRPVRRDTHDDSIPKELFSVLPSVVSQCQHLEYVIIERLGHTLKTDEEKNDFLTDFNRVKKIIEASDWKGEQKKNWKKKEIKLTEKPLEDRVLFEEQSRLTQLLFDHVDAAVIKDVDFHYFKTENWDLEMILTAQNMIRKWNPY |
| WP_166692314.1 | Fibrivirga algicola | MAAIRSTLACNLETNLILAALPLFEAERVAALEWSFDALYAHEQLPDWFTELLTTYSEADRLVGHGVFFSLLSGRWTSEQQAWLIRLRALSREFRFDHITEHFGFMTGADFHKGAPINVPCTAVTLALGQDRLSRIQDACRCPVGLENLAFAYSLDDVKRQGDFLRQLIEPVNGFIILDLHNLYCQSKNFNLDLTALLDLYPLDRVREIHISGGSWAEVEIEPGRHIRRDTHDDAVPDDIFNWLSTAIDRCPNLGYVVLEQLGAGLTTGSERLAFQADFDRMDVIVQQKNKSIAHPDMQSFLPELPLLINPVPLENHRLYTHQLELSAILETAVDVQQAKRRLAESSLANSEWEVERWEPAMLETAVAIARKWKNGFVGKE |
| WP_165835292.1 | Chryseobacterium viscerum | MRKPLLGVSMMAEADFVSAILPLLQNNSIEVLEWSFDTLYHTHEPDWLCDLLNFYAENNRLIGHGVYYSLLDARWTARQEQWLEKLKEEVQQRKYNHITEHFGFMNTDNFHQGVPLPVSLHPKTLQIGKDRLNRLQEAVDIPVGIENLAFSLSVDDVKEQGMFLDKLTEDSNGFLILDLHNIYCQSCNFGVEMQEIINLYPLDKVKEIHLSGGSWQESVYGKMPVRRDTHDDVIPEDIFSVLPAVLTQCQSLEYVIIERLGHTLKTEEEKSNFLHDFNKVKTIIEASDWKNRRENNWTKKEIQLAEKPVEDLVLYEEQSRLTRLLFDNVGAAAIKDQDFHYFKTENWDPEMILTAQNIIKKWNPY |
| WP_165851835.1 | Chryseobacterium pennipullorum | MAIIAAPPVVWDKMKKPLLGLSMMAEAEFVSAILPLLQSNLVDVLEWSFDTFFHAEEPDWLNDLLDFYSDSRRLTGHGVYYSLFDAKWTERQEVWLDTLKKEVQTRTYNHITEHFGFMNTENFHQGVPLPVSLHPVTLQIGKDRLFRLQDAVNIPVGIENLAFSFGIDDVKEQGVFLDKLTEDTNGFLILDLHNIYCQSCNFEIPMEEIINSYPLDKVREIHLSGGSWQESAYGKKQIRRDTHDDAIPESLLDVLPSVLLKCKNLQYVIIERLGHTISTEKEKKRFTEDFQKVKKMLEAPGLLTENTTSWIKKGVVIPDVPLEDLALAEEQSRLTRLLFEGADAAAFKKENFHFFNTRNWDSEMIRTAQNIIKKWNPY |
| WP_167288130.1 | Paraflavitalea devenefica | MPKILSSVACNLEADILGACIPLWEEARIEAIEWSFDALFNVEEIPAWFRELLTAFSKEKRLIGHGIFFSLFSGKWLPEQQQWLNKLQQTCTEFQFDHITEHFGFMTGKDFHQGAPLNIPYTTSTLAIGRDRLQRIQEACQCPVGLENLAFSYSLDEVKRHGTFLDQLLEPVNGFIILDLHNLWCQLHNFSIPFEEVIDLYPLHRVREIHISGGSWDDSEVEAGRTIRRDTHDNAVPAAVFQLLEMTISRCPHLQYVVLEQLGTGLLTPESKQQFYQDFGRMESIVSEKNLSRTPHSTCSFLPQAPLQKGGVTEDLFLYQQQRELSAILESAPSYEDARRLLAQSSLAHSDWKIENWDPAMVETAIKIAQKWKQGWLT |
| WP_168679930.1 | Hymenobacter sp. BT18 | MTKPPTIFAALALNLDADMLAAAFPLLADGSVGALEWSFDALSGANTPVWFTELLHAFSKEQRLIGHGVFFSLLSARWTTEQQQWLQRLNQLARQFSFNHVTEHFGFFTGQNFHSGAPLPVPYTAATLRIGQDRLCRISEACSCPVGLENLAFAYSLEEVKRHGEFLDKLLTPVNGFLILDLHNLYCQLHNFSISYEELIALYPLEEVREIHISGGSWADSSLLPDKKIRRDTHDDAVPDEVFSLLKATMPRCPHLKYVVLEQLGNGLQTEHSQTRFRQDFLRMQKLVAQQNQESRPTDYHFLPRQPVVSGPVEEDVRLYAQQQQLSHILETAHSYEEAHQRLLLSSLAQSEWQIEQWAPYMLETALGIAQKWQLKPS |
| NJN38771.1 | Acaryochloridaceae cyanobacterium CSU_3_4 | MAWAKLPMPPLVGLSLMPQADFMAAAQPLFRAGEVEILEWSFDMGWGNVSLPPELPPLLDQFSQCHCLLGHGVSYSLLSAGIDRSHWLACLQAECQDYSYRHISEHFGWMATRHFSRSVPLPLPLLPATLKLGCDHIQRLAEVAQVPVGLENLAFAFGRQDVDQQGEFLDRLLEPVDGFLLLDLHNLYCQMHNFQLSASEILNLYPLGRVRELHLSGGSWSQSGTAKIRRDTHDHAVPEPVFDLLALALQECSHVEAIILERLGNTFTTDAEKIGFRQDFARIKEIVYSHRSSLSRKG |
| WP_167028800.1 | Chryseobacterium sp. Tr-659 | MGKPLLGLSMMAEADFVSAILPLMQNNAVDVLEWSFDTLYNVKEPDWLNDLLNFYAENNRLIGHGVYYSLFDARWTERQNVWLKKLKEEFLRRNYNHITEHFGFMNTENFHQGVPLPVSLHPKTLQIGKDRLYRLQEAVNVPVGIENLAFSFSSDDVKEQGEFLDQLIEDTDGFLILDLHNLYCQACNFEVEMQELINSYPLDKVREIHLSGGSWQKSVYGKRMVRRDTHDDLIPEEILSILPSVLAQCKNLEYVIIERLGNTIKTEAEKEGFFTDFNKVRAIIDASDLQVGKMGNWTKKEIKLPDEPLENMELFEEQSRLTKLLFDSVGPAILKGQQFHYFKTENWEPEMILTAQNIIKKWNPY |
| WP_172523132.1 | Chryseobacterium indologenes | MRKPLLGLSMMPEAEFVSAALPLLQNNSVEVLEWSFDTLFNTNEPDWLRDLLNFYAENNRLIGHGVYYSLFDAQWTERQEIWLSRLKDEVSKRNYNHITEHFGFMNTENFHQGVPLPVPLHPITLQIGQDRLQRLQDAVNIPVGIENLAFSFCLDDVKEQGVFLEKLIDNSGGFLILDLHNMYCQSCNFDVDIQEIIKLYPLQKVKEIHLSGGSWQESVYGKKMIRRDTHDHTIPEEIFAILPFVLSHCENLEYIIIERLGHTIKTKEEEGDFFSDFMRVKTIIDSSAGHERQGDLWIKKEFELSKYPVEDPVLYEEQSRLTKLLFDGTHVDSIKNENFHYFNTAGWDPEMIFTAQEIIKKWNPY |
| WP_168800700.1 | Psychroserpens sp. NJDZ02 | MKNNKPKLGLSLMPNPEFIAAVLPLFEAAQVEVIEWSFDTILDKTYQPEWLPLVLKEYGDNNRLLGHGVYYSLLDANWSANQDNWLKKAKQETLTYNYTHISEHFGLMSATNAHNGFPLPFDLSDTILNIGIDRLKKLQNTVQLDVGIENLALASNASDILKQGEFLNKLVTPVNGFIILDLHNIYCQSENFDIDMLTIITSYPLDLVKEIHISGGSWDQDTSLTKPIRRDTHDGCIPEAILNILPEVLQRCPVLEFIIFERIGDAFKDQKDGIDFRADFNKIKAIIDHTNFSTTPRHWSLNKHNLDQPLIDLELVNQQNTLRQNIQLDQPNAHPEWDTDLWKTATKLYKKWNK |
| QJB36205.1 | Chitinophaga oryzae | MEAAKVEAIEWSFDALYAAENVPAWFEELLAAFSKENRLIGHGVFFSLFSGRWLPEQQQWLDHLKTVSRHYRFDHVTEHFGFMTGKDFHHGAPLNIPYTPVTLGIGRDRLKRMQDACGCPVGLENLAFSYSLEEVKRHGDFLNELLEPVNGFVILDLHNLYCQAHNFDLSFEALKDLYPLHRVREIHISGGSWDESEATPGKTIRRDTHDDAVPATVFNYLDKAIDLCPSLKYVVLEQLGSGLASEESRSAFRNDFLRMDQLLNSKNRTTAPLNTFLPEAFQVPAQVPEDETLYRQQLELSQILETAASFDAAMKALQASSLAHSAWQIENWDPAMIETAAKIAQKWKRKDIVFKT |
| QJB42701.1 | Chitinophaga oryzae | MEAAKVEAIEWSFDALYAVENVPAWFEELLNVFSEENRLIGHGVFFSLFSGRWLPEQQQWLDHLKTVSRHYRFDHVTEHFGFMTGKDFHHGAPLNIPYTPVTLGIGRDRLKRMQDACGCPVGLENLAFSYSLEEVKRHGDFLNELLEPVNGFVILDLHNLYCQAHNFDLSFEALKDLYPLHRVREIHISGGSWDESEATPGKTIRRDTHDDAVPATVFNYLDKAIDLCPSLKYVVLEQLGSGLASEESRSAFRNDFLRMDQLLNNKNKSTVPLNSFLPEALQIPAQAQEDETLYRQQLELSQILETAASFGATMKALQASSLAHSDWQIESWDPAMIETAAKIAQKWKRKTTNP |
| WP_168237931.1 | Chryseobacterium gallinarum | MEKPFLGISMMAEADFLTAILPLLRDGSIDVLEWSFDTFYHAEEPDWLTRLLNFYSQNNRLLGHGVYYSLFDAKWTERQSMWLQKLKEEVQQRKYNHITEHFGFMNTENFHQGVPLPVSLHPKTLEIGKDRLCRLQEVLEIPVGIENLAFSFSIDDVKQQGIFLEKLIEDTDGFLILDLHNIYCQACNFGLGIEEIVRLYPLEKVREIHLSGGSWQTSPYGKRRIRRDTHDDTVPAEILAVLPSVLSQCENLEYVIIERLGHTIQTETEAKDFLDDFLKVKAIIGASEWNSRHKEIKIKKKPECAGRPVEDEGLYREQTILTKLLFDNVKADHFKDFDFQYFKTENWDPEMIRTAQNIIKKWNPY |

| | | |
|---|---|---|
| NNE28949.1 | Saprospiraceae bacterium | MIKAAISCNLDTHILQTALPLFRTAEVDAIEWSFDTLYKTKNIPDWFVELLRTFSDGKGLIGHGVFFSIFSGEWTEGQQRWLQHLEKVSKHFSFQHITEHFGFMTGADFHKGAPISVPMDNTSLSIGGDRLKRIANACSCPVGLENLAFTYSLEEVKKHGEFLSRLVSTVNGFIILDLHNLYCQMHNFQISFEEIIKLYPLELVREIHISGGSWQESDLAPEKKVRRDTHDQAVPIEVFNMLSKAIPLCPNLKFVVLEAIGTGLKTEKEKEDFQKDYQTMKALLTKSEPNPGIKNFIPDIPYELGEPMSDDEFALQQIELSNILETADNLTTAQASLSNSSLANTSWDIEHWSPHMLEAVIAIAQKWKMGFD |
| WP_171032588.1 | Chryseobacterium indologenes | MRKPLLGLSMMPEAEFVSAALPLLQNNSVEVLEWSFDTLFNTNEPDWLRDLLNFYAENNRLIGHGVYYSLFDAQWTERQEIWLSRLKDEVSKRNYNHITEHFGFMNTENFHQGVPLPVPLHPITLQIGQDRLQRLQDAVNIPVGIENLAFSFCLDDVKEQGVFLEKLIDNSGGFLILDLHNMYCQSCNFDVDIQEIIKLYPLQKVKEIHLSGGSWQESVYGKKMIRRDTHDHTIPEEIFAILPFVLSHCENLEYIIIERLGHTIKTKEEEDDFFSDFMRVKTIIDSSAGHERQGDLWIKKEFELSKYPVEDPVLYEEQSRLTKLLFDGTHVDSIKNENFHYFNTAGWDPEMIFTAQEIIKKWNPY |
| WP_168801929.1 | Chryseobacterium candidae | MLGVSMMAEADFVSAVLPLLQNNSIDVMEWSFDTLYHANEPDWLRDLLNFYAENNRLIGHGVYYSLFDARWTERQEEWLKKLKEEVNLRKYNHITEHFGFMNTENFHQGVPLPVSLHPKTLQIGKDRLYRLQEAVNIPVGIENLAFSFSIDDVKEQGVFLDKLTEDTNGFLILDLHNIYCQSCNFELEMQEIIDLYPLDKVKEIHLSGGSWQDSVYGKKQVRRDTHDDVIPRDILSVLPSVLTQCENLEYVIIERLGHTLKTDEERDHFLDDFSKVKTIIETSELNGKEKSNWSKMKVELSEKPLEDPVLYEEQSRLTQLLYDNADVETIKDLDFYYFKTESWDAEMILTAQNIIKKWNPY |
| NOT36767.1 | Saprospiraceae bacterium | MIYCGTALNLDHKLLSANLPLLANSKLEAIEWSFDALYKLEQVPDWFDQLVKLYSNQKRLIGHGVFFSIFSGKFSSEQKKWLEDLKILSNKYRFDHITEHFGFMTGEDFHKGAPLSIPMNLTALRIGQDRLKRIQDVSNCPIGIENLAFAFNVEDVKRQGEFLNDLLSPINGFMILDLHNLYCQAHNFKIDIDNLLQYYPLELVREIHISGGSWEDDIVHYNEKIRRDTHDDSVPSEVYICLESCLLKLDNLKYVILEQLSNALESESQQKKYQSDFYKILAIVKNNKVDEIRRNDFHPSFLDISTQEAFEDLHLYQQQAELSHILVDSKNLEEAKQKLSHSSLRNSDWNVELWQDNMLYTAMSIAQKWK |
| HIA55772.1 | Candidatus Melainabacteria bacterium | MPCKPKLGLSLMPTEDFEQATVSLFKNKQVQAIEWSFDFSWNGAVVSEWCDGILHEFSNRGALTGHGVNLSPLSARFSKRQEEWLANAKEEFKNRKYVHATEHFGFSEAGPIAQGAPLGVPMNAESLRKGKDMMKRYADATQCPVGLENLAFAFGLEDVKRQGDFIDQLISSVDGFLLLDVHNIYCQLANFGVSELELLNSYPLSKVREIHLSGGTWSGSISGKRAAVRRDTHDDAVPQEVFNLATLALKLCPNIEFVIMERLGYTMMELAEQQEFREDFETMEEILEFCYA |
| WP_169236210.1 | Chryseobacterium antibioticum | MSRPLLGLSMMPEADFVSAILPILQNNSVDVLEWSFDTFYDADEPEWLSGLLDFYAENNRLLGHGVYYSLFDARWTDRQDVWLKKLKEELRHRKYNHITEHFGFMNTENFHQGVPLPVPMHSRILQIGKDRLRRLQDVVDIPVGVENLAFSFSINDVKEQGEFLDKLVEDIDGFLILDLHNIYCQSCNFEVDMQEIIKLYPLEKVKEIHLSGGSWQESAYGKKPVRRDTHDDRIPDKIVEILPKVLSQCPNLEYVIIERLGHTLNVESEKQVFFADFNRVKEMIEMSDDLPGDRKVWNKKKGEYSKPVEDLMLYHEQTKLTKLLFDGEGVGSVKNQKFHYFKPENWDEEMIMTAQQIIKKWNPY |
| NQZ69047.1 | Lentisphaeria bacterium | MKPLIGLSLMHEPEFLQASLELFENDCVDCVEWSFDSLWNNFSSCPWLDELLIEYSSQGRLFGHGVHFSFLSAELSTEQMDYLDKLNNEFDSRNYVHSSEHFGFMARTALGKGAPLPVPLIPDVFQLAEQRLELLGKHTGTAVGLENLALAFNKEDALKHYSFINDLAANGFILLDLHNIYCQIENFGISLADVLERIDSSRVHEIHISGGSWEEAMGKRIRRDTHDDAVPDAVFDLLQAALPQLPNVKCVIFERLGNSFKNENDAIEFRSDFMKIRASVT |
| WP_171593135.1 | Hymenobacter | MSTPTQLAKADAPGIFSSIACNLDANILTAAFPLLEDERVEALEWSFDTLFWAEQVPEWFTALLQAYAAQNRLVGHGVYFSLLSGRWTAEQQQWLQHLKALAGQFAFDHITEHFGFFTGQNFHSGAPLPIPYTSTTLRLGQDRLHRMQEACRCPVGLENLAFAYSLDEVKRHGEFLEKLVEPVNGFLILDLHNLFCQLHNFSVSYEALLALYPLERVREIHISGGSWEASGQVPGRQIRRDTHDEAVPEEVFQLLEQTLPRCPNLKFVVLEQLGTGLKTESSQAQFRADFRRMQQLINHYRSTTHRQASQLFLPQQPYSAGPVAEDAQLHTQQQQLSHILETAPSYAEARQLLQTSALAHTDWKIEEWEPHMLETAISIAQKWKQ |
| NUO01438.1 | Saprospiraceae bacterium | MPRPFAALACNLDRNTLLTCLPLFEAEKVEAVEWSFDALFQFKEIPPWFTGLLQTFSRENRLVGHGVFFSLFSGKWLEGQQTWLDELKKLSKTYCFDHITEHFGFMTGADFHNGAPISLPFTTTTLGIGRDRLARIYDACECPVGLENLAFAYSIEEVKRHGEFLEQLLSPINGFIILDLHNLYCQLHNFDLEFADLIQLYPLDRVREIHISGGSWEDSPLRPGKPVRRDTHDDSVPQEVFQLLEQTIPLCPNLKFAVLEQLGTGLGTEASRVAFQQDFDAMSGILQQSRDDSGHPANAFFPQTRPDTSQPPIESNELEQQQQELSQILETATDYEAAKNRLATSRLAHSAWNIESWEPAMLETAMRISQKWKHGFLQTQR |
| NUQ53559.1 | Phycisphaerales bacterium | MESPLPVGISLMLEEDFLGATLPLFEEGAVDVLEWSFDVGWGRELPAWAQGLLQHFAVSGRLLGHGVSFSPLSARWEEPQRNWLHRAALEVAKLKYRHVSEHFGFMAAGSFTRGSPMPMPADAAVIRTGRDALSRLAEVTGLPVGLENLALALSSRDVDGQGDHLDALLEPVGGFVVLDLHNLWCQAVNFGRDPVGLLGRYPLSRGITLVLLKIDARVLLVSQSVSRSGASMQTLCEIDEPDQVASILLKASRNERSSIADRFQSLLSSSDRAMEKELESAGRQVFRTSSGDTVELASISTPVPLARINAGDAAARRERNEGAPNRLSQPIQPPVQGRPDRREGNPAPVRGYSVAGASRPLTGAEQLRRRLDAWRDQEERAS |
| MBA4075682.1 | Cyanobacteria bacterium PR.023 | MLQEDFLAAALPLFSNDEVEILEWSPDIGWATDIPEWADELLNFFSSTHSLYGHGVNYSPLSADWTAVDSGWMKNLERELEQRQFVHFSEHFGFSRLGELKQGAPLPVPHTAQALAAGQFKLAKLAHTTGGRIGLENLALAFSLSDVKEQGRFLDELLEPFDGFILLDLHNLYCQLENFGLSPEELLKTYPMHRVREIHISGGSWSQAINNKIRRDTHDNEVPDEVMALLPLVLAKCPNIEAVIFERLGGTMNTAEQQHRFRDDFRAIAKLIEQHWLEKNLADEALTAMEAIALVEPTKPAELKAGDFPDILRYQKKLMELLQSGQKPEEIIAKLSKDFDGEIADYASSIQPQMIAVGQELVAKWTR |
| MBA4064164.1 | Isosphaera sp. | MDRADPAARPRVGLSLMPEPDFLRAAAPLFAAGAVEVVEWSFDTGWPPAAVPGWIDELLRHYGDAGRLLGHGVSFSPLSAGTGDAQRRWLGRLTEEVARRNYRHVSEHFGFSAAGDFHRSAPLPVPLTGPALRLGRDRLRRLADAAGVPVGLENLAFAFGRRDVDEQGHFLDALLDPVDGFLLLDLHNVHCQAVNFAVPFDDLLDRYPLHRVRELHLSGGSWSDGVRRDTHDGPVPEELFAALPGVLRRCPRVEAVVLERLGGTLAEADDEPFRRDFLRLAGCVRGAG |
| WP_182412767.1 | Adhaeribacter radiodurans | MPQILSSIACNLDADILTAALPLFEESRVGAIEWSFDTLYGNSQIPPWFLELLKAYSNENKLIGHGVFFSLFSGKWSQNQHEWLNQLRQLSSEFHFDHITEHFGFMTGQNFHYGAPLPIPYSKTTLAIGQDRLARIYEASQCPVGLENLAFSYSLDEVKQHGEFLEQLVAPVNGFLILDLHNLYCQIHNFTISYEDIINLYPLDRVREIHISGGTWQDSTVEPERKIRRDTHDEAVPEEVFQLLKRTIDKCPNLKYVVLEQLGNALKTEASKKLFYSDFLKLEEIVRNKNHSLASHFTNSFLPSKSVTTGSILEDEKLYQQQIQLSSILEKASSYQAAIHLLNSSPLAHSDWKIEQWEPHMLETAISIAQKWKKNLD |
| WP_184174413.1 | Rhabdobacter roseus | MPKVLSSIACNLDPNLLLAALPLFEAERVEAVEWSFDALYKYPTLPGWFEELIGTFAENDRLIGHGVYFSLFAGKWSKQQDQWLRQLRSLAARYPFDHITEHFGYMTGADFHKGAPLGVPYTSTTLKLGIDRLRRIQEAGQCPVGLENLAFAYSLDEVKKHGEFLEKLVGAVNGFIILDLHNLYCQLHNFRLDFEEVIRLYPLERVREIHISGGSWEPATTDDKYPIRRDTHDDAVPEAVFALLETTLPMCPNLKYVVMEQLGTALDTDARRLAFQRDFIRMDTLVKRLNDKLPQPATNDFGPLPTGLPTTLPLEDLSLHQQQRELSHILENAKSYPEARDLLAASSLRHSAWNVEAWVPSMLETARQIAQKWKNGFE |
| MBA3855477.1 | Cyanobacteria bacterium PR.3.49 | MRIKKPKLGLSLMPTEDFQMATERLFAEERVKSLEWSFDFTWNGVVVDQWAQELLDKYSSAKALTGHGVNLSPLSARFSKRQQEWLKLAKEEFKTRNYVHASEHFGFSEAGPIKHGAPLSVPMDDASLTTGKEMLKRYADATGCPVGLENLAFAFSVNDVKRQGQFIDELISSVDGFLLLDLHNTYCQIANFGVSELELLNSYPLEKVREIHISGGSWSPSVSGKRAAVRRDTHDDGVPQEVFNLAALALKLCPNIEFVILERLGNTMFSLEAQSEFRDDFESMEQILELCYA |
| MBA2612290.1 | Bacteroidota bacterium | MNPWVGISLMPEADFIEAALPLFQSGEIDVIEWSFDTIRSGSKEPQWLSLLLSEYSKRGRLIGHGVRYSLLKGLWDGNQGKWLKQLAIEVKKYNYRHVTEHFGFMSSDDFHKGTPLPVPFNKTTLAIGQDRLKRLQNISQLPIGIENLAFAFSNKQIKEQGEFLEKLLEPVNGFLILDLHNIWCQSHNFKVDALKLIKSYPLHKVKEIHISGGSWATVKTSKNKVRRDTHDEAVPKEVFELLKEALPFCENVEYVIFERLGNTIKTEKDKLQFVKDFKQIKKIVSSQRKKAESPTEKIKRKKIDHKVPLKPLQDNILRKEQQFIIDTLFANKDVDKTIQLLQTKQITGWQPALWTDYMVETAMVLIKKWA |
| WP_172957645.1 | Chryseobacterium indologenes | MRKPLLGLSMMPEAEFVSAALPLLQNNSVEVLEWSFDTLFNANEPDWLRDLLNFYAENNRLIGHGVYYSLFDAQWTERQEIWLSRLKDEVSKRNYNHITEHFGFMNTENFHQGVPLPVPLHPITLQIGQDRLQRLQDAVNIPVGIENLAFSFCLDDVKEQGVFLEKLIDNSGGFLILDLHNMYCQSCNFDVDIQEIIKLYPLQKVKEIHLSGGSWQESVYGKKMIRRDTHDHTIPEEIFAILPFVLSHCENLEYIIIERLGHTIKTKEEEGDFFSDFMRVKTIIDSSAGHERQGDLWIKKEFELSKYPVEDPVLYEEQSRLTKLLFDGTHVDSIKNENFHYFNTAGWDPEMIFTAQEIIKKWNPY |
| WP_317171826.1 | Hymenobacter | MQATPTLCSALACNLDVDILSAALPLLAEGRVEALEWSFDALFWAGQIPDWFSELLRTYSASGRLLGHGVFFSLLSGRWSREQQQWLTQLRQLANQLPFAHITEHFGFFTGQNFHSGAPLPVPYTPVTLRLGQDRLSRLYEACQCPVGLENLAFAYALEEVKRHGEFLDKLLEPVNGFLILDLHNLYCQLHNFSISYEDLIDLYPLHRVREIHISGGSWEASALAPGQRVRRDTHDQGVPEEVFQLLQRTMPRCPHLRFVVLEQLGNSLRTEDSRARFRLDFGRMEALVQEHRAGLTASADQAFLPLRPPQPGPALEEPALHEQQQHLSRILETAPSVAEARQLLLASPLLASTDWNIEQWAPHMLETAVRIAQKWK |
| WP_184553106.1 | Chryseobacterium sediminis | MKKPLLGISMMPEADFVSAVLPLLQSNAVDVLEWSFDTLYHTNEPDWLRDLLNFYAENNRLIGHGVYYSLFDARWTERQEEWLQKLKEEVSLRKYNHITEHFGFMNTENFHQGVPLPVSLHPKTLRIGKDRLYRLQQAVNIPVGIENLAFSFSIDDIKEQGVFLDQLTEDTNGFLILDLHNIYCQSCNFGVEMQEIIDLYPLDKVKEIHLSGGSWQESVYGKQIRRDTHDDIIPEEIFSVLPSVFAKCQNLEYIIIERLGHTLKTEDDKENFLNDFNNVKTIIEISDWKERGKSGWNKQPIQLPEKPVEDMALFEEQSRLTRLLFDGAGANAIKDQDFHYFKTENWDSEMILTAQNIIKKWNPY |
| WP_185274038.1 | Adhaeribacter swui | MPEKILAAVACNLDVDILAAALPLFAAGKVEAIEWSFDTLFRTPQIPAWFSELLQTYGQAGRLIGHGVYFSLFSGQWSVNQQNWLQHLRELSGQYTFDHITEHFGFMTGQNFHHGAPLPIPYSASTLAIGQDRLARIQDACKCPVGLENLAFSYSLDEVKQHGQFLAQLITPVNGFIILDLHNLYCQLHNFRASYDEIVNLYPLDRVREIHISGGSWESSAIEPNRQIRRDTHDEAVPEEVFALLAQTIDKCPNLKFVVLEQLGNALKSEESKQLFYADFLKMEAILQRKNQKLRTNPAKDFLPLSLSIAEHPIEDKTLAAQQVHLSRILENASSAEEAIQRLQTSSLAHSDWQIETWEPYMIETAHNIARKWKM |
| WP_184160252.1 | Chryseobacterium shigense | MKKPLLGLAMMPEAEFVSAILPLLQNNDVDVLEWSFDTLYDVKEPEWLSGLLDFYAENNRLLGHGVYYSLFDARWTERQESWLKKLKKEFKHRKYNHITEHFGFMNTENFHQGVPLPVPLHSKTLQIGKDRLCRLQDAVEIPVGIENLAFSFSVDDVKEQGEFLAELIEDIDGFLILDLHNIYCQSCNFEIDMQDIIRMYPLEKVKEIHLSGGSWQESAYGKKPVRRDTHDDNIPEVILNVLPEVLAQCNNLEYIIIERLGHTLDTEAKKHVFFDDFYRVKAIIDISDFSAGSNKVWNKKNKGYSGPVEDLLLYDEQTKLTKMLFEGNSVQAVKNEKFHYFKPENWDEEMIATAQQIIKKWNPY |
| WP_187319759.1 | Hymenobacter citatus | MPAVPTAPAILSAIACNLDDQILTAALPLLAAGRVEAVEWSFDTLFNGQSLPNWFAELLTAYGEQSRLVGHGVYFSLLSGRWTPEQAQWLEQLRLLAGRFRFDHITEHFGSFTGQNFHTGAPLPVPYTATTLRLGQDRLLRVADACQCPVGLENLAFAYSLAEVQRHGEFLEELVAPVNGFLILDLHNLYCQLHNFNLSSDSLLALYPLHRVREIHLSGGSWEDSQVEQGRQIRRDTHDESVPDEVFELLERTLPQCPQLKYVVLEQLGTGLRTEASQHQFRQDFQRMAAIVEAGRGRVSGHQPNAFQPPASLPAGSVAYDEALHAQQRQLAQILETAASYEEAQQRLRASSLAHTSWQLEQWAPYMLETAISIAQKWKQ |
| WP_185133763.1 | Chryseobacterium indologenes | MRKPLLGLSMMPEAEFVSAALPLLQNNSVEVLEWSFDTLFNTNEPDWLRDLLNFYAENNRLIGHGVYYSLFDAQWTDRQEIWLSRLKDEVSKRNYNHITEHFGFMNTENFHQGVPLPVPLHPITLQIGQDRLQRLQDAVNIPVGIENLAFSFCLDDVKEQGVFLEKLIESSGGFLILDLHNMYCQSCNFDVDIQEIIKLYPLQKVKEIHLSGGSWQESVYGKKMIRRDTHDHIIPEEIFAILPFVLSHCENLEYIIIERLGPTIKTKEEEDDFFSDFMRVKTMIDSSAGHERQGDLWIKKEFELSKYPVEDPVLYEEQSRLTKLLFDGTHVDSIKNENFHYFNTAGWDPEMIFTAQEIIKKWNPY |
| WP_185246801.1 | Chryseobacterium | MKKPLLGLSMMAEPEFISAVLPLLQSNSIEVLEWSFDTFYNAQEPDWLSDLLHFYSENDRLIGHGVYYSLFDAIWNERQEVWLKKLKEEFHKRKYNHITEHFGFMNTENFHQGVPLPVSLHAKTLEIGKDRLYRLQDAVQVPVGIENLAFSFSLDDVTEQGIFLDKLTEDTDGFLILDLHNIYCQSCNFEVEIQDIIKLYPLDKVREIHLSGGSWQESVYSKSPVRRDTHDDAIPKEILSILPSVLSQCKYLEYIIVERLGHTINTEEKKIAFLDDFAQIRTLIDSSDRDQQYKAPWTKKQVEFQKNPLEDKVLHEEQTLLTKLLFNKEKPESIKNHEFQYFKTENWDPEMILTAQNIIKKWNPY |
| WP_185288143.1 | Chryseobacterium lactis | MQKPLLGLSMMAEAEFVSAILPLLQNNSVEVLEWSFDTFYNVEETDWLNELLQFYSENNRLIGHGVYYSLFDARWTERQEIWLKKLKEEVRKRNYNHITEHFGFMNTENFHQGVPLPVSLHPKTLQIGRDRLYRLHEAVDVPVGIENLAFSFSIDDVKEQGMFLDQLVEDTNGFLILDLHNIYCQSYNFEVNIQDIISLYPLDRVREIHLSGGSWQESVYGKKPIRRDTHDDRIPKELLAILPSVLSQCQNLEYVIIERLGHTIKTEEQKKYFFDDFNKVKAIIEVSDTRSRKVNKWTGKEAVHTDHPEEDLALYEEQSRLTRLLFDQAGTEGIKEEKFHYFNTESWDPEMILTAQNIIKKWNPY |
| MBC8152492.1 | Fibrella sp. | MSTIHSAIACNVDTNILLASLPLFDAEKIQAIEWAFDAVFRHDALPDWFDALLRTYADANRLVGHGVFFSLFSGRWTPDQQAWLTQLKTLAATYRFDHITEHFGFMTGADFHKGAPLSIPLTQSTLAIGQDRLRRIQDAGGCPVGLENLAFAYSLDEVKRQGDFLHRLLEPVNGFIILDLHNLYCQSHNFGLAGDELLALYPLEQVREMHISGGSWSASDSEPGRQIRRDTHDEAVPDAVFDWLEGAIDRCPNLRYVVLEQLGSGLTTEADQEQFRNDFLRMDRIVEGKNQVRAASQPNLFRNISGVLPSSMPLEDDDLHWQQRELATILETAADYRQARALLATSGLAHSAWEIERWEPAMLETAMAIAQKWQHGFD |
| WP_188686807.1 | Rufibacter glacialis | MPKVFASLACNLDVDILSASLPLLQEGRVEGLEWSFDTLFQQEQVPSWFLALLETYGREKRLIGHGVFFSLFSGKWSEHQEAWLRHLTQMSAAYGFDHVTEHFGFMTGKNFHHGAPLPIPYTAATLAIGRDRLARIQEAAQCPVGLENLAFSYTLEEVKRHGAFLEELVEPVNGFLILDLHNLYCQLHNFSLPLEALLPWYPLHRVREIHISGGSWQDSAVLPQRKVRRDTHDSAVPEEVFDLLSATMPLCPQLKYVVLEQLGTGLKSPEDKALFYQDFCRMETLVKQHNASTFPSLTAPNLFLPLAPGKLPAPVEDEALYQQQLLLSSILEESPSLTQAHQRLLASSLANTAWAIEHWEPHLLETALHLAQKWKKPDLSA |
| WP_192218895.1 | Microcoleus sp. FACHB-SPT15 | MFKFKRRTPMVGLSLMIESDFFQAAQPLFETGAVEVLEWSFDMSWGVKAKPPWVKELLQFYSQRDRLTGHGISYSLLSAQPGERQIHWLKSLQEECLDYRYRHISEHFGWMAAGDFYQSAPLPLPLTPETLQLGREHLKRLSDKANAPVGLENLAFAFGLQDVLQQGEFLARLLEPVDGFLVLDLHNLYCQVHNFQQSAEQLLERYPLQRVRELHVSGGSWSVTSKGQQTRTIRRDTHNGTVPEAVFELLALALKRCPSVEAVIFERIGHTLHTEADIEGFRQDFWRIKQVVKESV |
| MBC7862316.1 | Bacteroidia bacterium | LLQEYSANKRLIGHGVRYSMFDAKWTKRQSDWLSRLKKEVKKYQYNHITEHFGFMSNTDFHKGAPMPVPLNKSSLAIGIDRLKRMSAVAGVPVGIENLAFSFSAEDVKKQGLFLQKLVQPVNGFLILDLHNIYCQSKNFRIGMLKIIRAYPLKKVKEIHISGGSWQQSIYSTDLQKVRRDTHDGQVPEEIFKILPKVLKLCTNLEYIIFERMGNTMQDENEQLQFRKDFLRIKSVAGNVSSSASKRNGRNNWLKPRLSLDKKPITDSKLLSEQQHILEVMRTTENPYDALNKLNKEKLKDWQVLKWKPSMVETAIQLIRKWD |

| | | |
|---|---|---|
| MBC8083770.1 | Hymenobacter sp. | MEDEPILFSSIACNLDADILSAAFPLLAEGRVEAIEWSFDTLFWAEQIPDWFAELLSSYSSQNRLVGHGVYFSLLSGKWTKEQQQWLEHLQQLSAQFSFDHITEHFG<br>FFTGQNFHSGAPLPIPYTSTTLRVGQDRLRRIYEACQCPVGLENLAFAYSLDDVKQQGEFLEKLITPVNGFLILDLHNLYCQLHNFSISYEELIDLYPLDRVREIHISGG<br>SWEDSGLVPGKKIRRDTHDGAVPEQVFQLLEQTMPKCPHLKYVVLEQLGNGLQTQVSRIRFRNDFRTMTGLVGKWRNGLPRRSDQLFLPLQSFPAGPVAADAILH<br>EQQLQLSQILETAPSFEEARRRLYASSLAGTNWKIEDWEPHMLETAISIAQKWK |
| GGF13189.1 | Hymenobacter cavernae | MEEAPIPFAAIACNLDFDILAAAFPLLEAGQVEALEWSFDTLFRAGQVPDWFSELLSAYSQQGRLIGHGVFFSLLSGKWTPQQQQWLQHLQQLSAQFRFDHITEHF<br>GFFTGENFHYGAPLPIPYTQASLRIGQDRLSRIYEACQCPVGLENLAFAYTPDEVKRHGEFLNKLLEPVNGFVILDLHNLYCQLHNFAVSYEELIELYPLERVREIHIS<br>GGSWEESEQAPQGKVRRDTHDEAVPEAVFELLARTLPRCPNLKYVVLEQLGNGLKTESSRTQFRHDFLRMEALVQKHRASLLGHPEHLFRPPQPLQLGEVIEDQS<br>LHEQQRQLSRILETAASVEEAQRRLHASVLVGTDWQMEQWEPHMLETAIHLAQKWKQ |
| MBD1867410.1 | Cyanobacteria bacterium FACHB-471 | MFKRKQNLPWVGLSLMTDPHFFTATRPLFEQGEVEVLEWSFDAGWEDTIPSWASELIDFYSQRDRLLGHGVTYSLLSAEWSDRQMRWLKQFQAECSRHRYVHIS<br>EHFGWMSAGNFARSAPLPMPYTPDAVSLGRDRLQRLADAAKVPVGLENLAFAFGQKDVADQGKFITELLEPVDGFLLLDLHNLFCQIHNFQLSIPELLASYPLARVR<br>ELHLSGGSWSRTTPDSNLVRRDTHDGAVPEAVFDLLAIVLKQCTNVKAVILERLGHTLNSEPEIEQFRQDFNRIKQIVKRCG |
| WP_190774526.1 | unclassified Leptolyngbya | MPTPDFFQASQPLFEAGAVQVLEWSFDMGWEADLPDWVTHLLEFYSQRDRLLGHGVTYSLLSAQWEEAQDQWLKNLQEECKRYRYRHISEHFGWMAAGNFVQ<br>SAPLPLPLTADTLRLGCDRLNCLAAAAKVPIGLENLAFAFGQRDVEEQGTFLDQLLAHVQGFLVLDLHNLYCQIHNFQQSPEALLKRYPLHRVRELHISGGSWSKSA<br>SSDSMIRRDTHDGPVPEMVFELLAIALQHCSHVEAVIFERLGHTLSTETEQQQWRQDFQRIQQIVHQRRR |
| WP_190680896.1 | Leptolyngbya sp. FACHB-671 | MFKRKQNLPWVGLSLMADSHFLTATRPLFEQGEVEVLEWSFDASWEDTIPSWASELIDFYSQRDRLLGHGVTYSLLSAEWSDRQMHWLKQFQAECSRHRYVHIS<br>EHFGWMSAGNFARSAPLPMPYTPDAVSLGRDRLQRLADAAKVPVGLENLAFAFGQKDVADQGKFITELLEPVDGFLLLDLHNLFCQIHNFQLSIPELLASYPLARVR<br>ELHLSGGSWSRTTPDSNLVRRDTHDGAVPEAVFDLLAIVLKQCTNVKAVILERLGHTLNSEPEIEQFRQDFNRIKQIVKRCG |
| WP_190645960.1 | Oculatella sp. FACHB-28 | MFKRKQNLPWVGLSLMADSHFLTATRPLFEQGEVEVLEWSFDAGWEDTIPSWASELIDFYSQRDRLLGHGVTYSLLSAEWSNRQMHWLKQFQAECSRHRYVHIS<br>EHFGWMSAGNFARSAPLPMPYTPDAVSLGRDRLQRLADAAKVPVGLENLAFAFGQKDVADQGKFITELLEPVDGFLLLDLHNLFCQIHNFQLSIPELLASYPLARVR<br>ELHLSGGSWSRTTPDSNLVRRDTHDGAVPEAVFDLLAIVLKQCTNVKAVILERLGHTLNSEPEIEQFRQDFNRIKQIVKRCG |
| WP_191041552.1 | Spirosoma validum | MPKIYSSIACNVDTNILQATLPLFEAEKVQAIEWSFDTLYQHNQIPDWFAELLQAYGQENRLIGHGVFFSLFAGKWTTAQQEWLDRLKKLASEFTFNHITEHFGFMTG<br>EDFHKGAPISVPFISSTLALGQDRLKRIQNACNCPVGLENLAFAYSLDEVKQHGDFLNQLIEPINGFIILDLHNLYCQSQNFSVAVSDLLSLYPLDRVREIHISGGSWTS<br>SSTEPEKRIRRDTHDEAVPMEVFELLEQVIAQCPNLNFVVLEQLGSGLITEESRRLFQEDFLRMDDIITTYTNTIPPENLFLPLVPFDISSSPLQNHLLHTQQTELSTILE<br>TATNYQQAQHLLRSSSLANSAWQVENWNSTMLETALAIAQKWQHGFN |
| WP_197901599.1 | Rhodocytophaga rosea | MAKILSSVACNLDVHILSASIPLLEESQVDAIEWSFDTLFNIWHTPGWFARLLTAFSKEERLIGHGVFFSLLSGKWSKEQYQWLSQLKQLSAEFHFDHITEHFGFMTG<br>QSFHQGAPMCVPYSPATLAIGRDRLARIQEACSCPVGLENLAFAASLEQVKQQGEFLERLIEPVNGFIILDLHNLFCQMHNFFMDFEQLLSSYPLERVREIHISGGSW<br>EKSGKQSERIIRRDTHDQAVPPEVFHLLEKTIPVCPNLKYVVLEQLGIALKTQESKAAFYQNFVQMKQIVEDNNQTIANSSENPFIPPSITLPETFVENQELYAQQREL<br>SEILETSSSYEETNRLLKASSLAHTAWNIESWDPAMIETAIHIARKWKK |
| MBD2766342.1 | Hymenobacter montanus | MEAGEVEALEWSFDALMGEMPEWFSTLLSVYGEQGRLVGHGVYFSLLSGQWTPEQQQWLHDLGQLAARFSFDHVTEHFGFFTGESFHAGAPLPVPCSAATLRL<br>GRDRLRRIQDAGRCPVGLENLALAYSLADVQQQGAFLDQLLEPVNGFLILDLHNLYCQLHNFAIAPETLIDLYPLDRVREIHISGGSWEASARAPRGRVRRDTHDGA<br>VPEEVFQLLERTMPRCPHLKYVVLEQLGTALKTERQRAGFRRDFRRMARLVQQHDRATTRGSDQLFLPLHPSPGGPVVVDEKLHGQQRQLSRILEAAPTLDEAQR<br>LLHGSSLAHTEWHLENWPPYMLETARRIAQKWKR |
| WP_193545348.1 | Chryseobacteriu m culicis | MNKPLLGISMMAEADFVSALLPLLQSNSVDVLEWSFDTLYHNHEPVWLCDLLNFYAENERLIGHGVYYSLFDARWTERQEDWLKKLKEEVALRRYNHITEHFGFM<br>NTENFHQGVPLPVSLHAKTLEIGKDRLYRLQDAVNIPVGIENLAFSFGIDDVKEQGAFLDKLIEDTDGFLILDLHNIYCQACNFGIEIQEIMDWYPLDKVKEIHLSGGSW<br>QESVYGKRPVRRDTHDDSIPKELFSVLPSVVSQCQHLEYVIIERLGHTLKTDEEKENFLTDFNRVKKIIEASDWNGEQKKNWKKKEIKLTEKPLEDRVLFEEQSRLTQ<br>LLFDHVDAAVIKDVDFHYFKTENWDLEMILTAQNMIRKWNPY |
| WP_190796908.1 | Leptolyngbya sp. FACHB-541 | MFKRKQNLPWVGLSLMADSHFLTATRPLFEQGEVEVLEWSFDAGWEDTIPFWASELIDFYSQRDRLLGHGVTYSLLSAEWSDRQMHWLKQFQAECSRHRYVHIS<br>EHFGWMSAGNFARSAPLPMPYTPDAVSLGRDRLQRLADAAKVPVGLENLAFAFGQKDVADQGKFITELLEPVDGFLLLDLHNLFCQIHNFQLSIPELLASYPLARVR<br>ELHLSGGSWSRTTPDSNLVRRDTHDGAVPEAVFDLLAIALKQCTNVKAVILERLGHTLNSEPEIEQFRQDFNRIKQIVKRCG |
| MBE2206138.1 | Saprospiraceae bacterium | MSPSIHTGIACNLDSNIITAALPLLEQEHVEAIEWSFDALFNWNEVPAWFRDLLQVFADAGRLTGHGIFFSIFSARWSAKQDLWLEQLRQLNRELPFDHVSEHFGFM<br>SGRDFHHGAPLSVPLTASTLAIGQDRLLRIQDACQCPVGLENLAFAFSLEEVRQHEAFLEQLLEPVNGFIILDLHNLYCQMKNFDLGFDALVHSYPLHRVREIHISGG<br>SWEPSTIIPTRNIRRDTHDDAVPEEVFEMLAQTIALCPALKFIFLEQLGAGLHTVSQQSQFRSDYLRICEIVKKHAPAVAQGDAESFSPRPNLQPPSAHIQDAVLYEEQ<br>RRLSRILETASDCQDARQKLESSILKNSAWQTEAWPDYMLETALRIAQKWKDGF |
| MBI1268694.1 | Cryomorphaceae bacterium | MTVKPKLGLSLMSTPDFRQASAELFESGKVEVVEWSFDNAFNGAFIAPWCQNIVDRYSDANALLGHGVTLSPLSVRFSRRQQDWLLGAAEEFRRHNYIHASEHFG<br>FSEAGPVAHGAPLAVPMNSATLKSGKEMMKRFAEATRRPVGLENLAFAFCMEDVKRQGEFIDHLLSDVDGFLLLDLHNLYCQQLNFNIPAIELLESYPLARVREIHIS<br>GGSFSPSISGRRIFVRRDTHDEQVPQEVLELVVHALKRCPDIEYVIYERLGFTMEDEESQAEYREDFETLAQLLEMIYV |
| MBI1224055.1 | Bacteroidota bacterium | MPFASIACNLDRNTLLAALPLLESGNVEGLEWSFDALFKVREIPEWFQELVFTYGKASRLIGHGVYFSIFSGKWRPEQENWLSQLRQFSKTYQFDHVSEHFGFMTG<br>EDFHKGAPIGIPYNASTLSIGVDRLFRIQDACQCPVGLENLAFSYSLDEVKKHGDFLSKLVEPVNGFIILDLHNLFCQMHNFEVDFEELMALYPLERVREIHISGGSWE<br>ASNIVSNKKIRRDTHDSQVPETVFKMLESSLSRCPNLKFVVMEQLGNALGGDAEKAQFQADFLKMSSIIQHETPGNFTQQNDFSPNAARILGEPLENEFLAYQQNEL<br>SQILETSPNFETTRQLLANSSLANSDWKVENWEPTMLETAMVIAQKWKHGFV |
| MBJ6143402.1 | Hymenobacter sp. BT559 | MLAALACNLDADILSAALPLLEAGQVDALEWSFDALYWASEVPAWFTELLHAYSSEQRLLGHGVYFSLLSGRWTSEQQQWLCQLRELAQRYSFAHVTEHFGAFTG<br>QNFHAGAPLSVPYSPVALRIGQDRLARLAEAARCPVGLENLAFAYSLDEVKRQGDFLSALLAPVNGFLILDLHNLYCQLHNFSIDYEELLGCYELDRVREIHISGGSW<br>ENSQLLPTRRIRRDTHDEAVPTEVFELLARTLPRCPNLKYVVLEQLGTSLKTEASQVQFRRDFAQMAALVQQHNHAPRSVPSAFLPRWPALPALAVQDDLLHEQQ<br>RQLSHILETATSLPEAQQRLRASVLAHSAWQIEHWEPYMLETALSIAQKWK |
| MBK6512547.1 | Polyangiaceae bacterium | MGLAARGVRQGVSHPSGAPPRVGLSLMPADDWWAANEPLFFRGLIDAVEWSVDFGWGPDGVPEWLASLLERYGSQDRLYAHGVELSAMSAIWTPEQDAWLAE<br>LEATCARSRFRHLTEHYGFITAADFVRGTPLPLPPSAAALGLAERRLQTLRERAGVPVGIENLAFALSRRDVEAQASFVSQLLERTDAFLLLDVHNLYCQAHNFGIDP<br>LVLARSYPLHRAREIHVSGGSWSTPTSDPERRRFRRDSHDDRTPNEVFSLLSSVLELCPALEVVILERTDRSLFGRDEAERHREDFVRLRSLVHRRGQRGQAEPP<br>SGSRLPRVELARDDEVALEAFQGALVTTLASQADPHEVKAALQAAAPLAPYREHLDSYEPRALEIGAALVKQWGARAEPEDSMRAVVFRGPGSPLELRSLPRVAP<br>GPGQVLVRVVAVGLCGTDAHAYRGSFPVPTPIVLGHEIAGVVEALGSGVTDLEVGDRVGVSWVQAGCGACAACDRGAFQRCASPRTWIENGGGLSELTVAEASG<br>CTRLPDGLDPELAAPLFCAGHVAFSGLRKAAVRGAERVAVVGLGGLGHLALQIASALGHETLAVSSSLDKLEDAREGFGADHVVRVDDGDVGGALERAGGADIVL<br>ATTSNMSDVARSVRGLRLGGRVVLMGLGEGLLSIDPIELVQREASVIGSVQGSRAELEELLTLAAEGKVRPRVEVFPLLMAQRALARLLEGRVRYRAVVRVG |
| MBI3238826.1 | Flavobacteriia bacterium | MKNQNGILASVACNLDQQLLQATLPLFIQEEVEAIEWSFDTLYKVRNIPDWFVELLTAFGNEGRLIGHGVFFSLFSGKWSAEQEQWLHHLKKVSGNFTFDHITEHFG<br>FMTGEDFHKGAPIGIPYNANTLALGIDRLKRIQGTCQCPVGLENLAFSYSLEEVKRHGEFLEALIEPINGFIILDLHNLYCQLHNFSLDAAELLPLYPLHRVREIHISGGS<br>WEQTRLEPQRTIRRDTHDDAVPEAVFELLQLTIPRCPNLKFVVLEQLGTALDTNQNQLQFQSDFMRMKSIVSTFNTADSINNFLPSSPLILNPEVAEDTLLALQQQEL<br>SAILENATDYENAVALLRQSGLAHSDWNVEEWSPYMLETAMRIAQKWKHGW |
| MBK7839075.1 | Candidatus Obscuribacter sp. | MATEQLFAQDKVDALEFNFDQPLNGIVLAPWCQELLDKFSSASALIGHGVNLSLLSAQFTKRQQDWLTYTREEFKTRRYVHASEHFGFSEAGPIAQAAPLAVPMDA<br>QSLSVGKEMLKRYAEATQCPVGLENLAFAFSLDDIKKQGDFIDQLISSVDGFLLLDLHNIYCQVTNFDLDALELLHSYPLSKVRELHLSGGSWSPSLSGKRIAVRRDT<br>HDEAVPQEVFNLTALALKLCPNTQFVILERLGNTMLEPELQAEHRDDFDTIREILDYSYD |
| MBK8443732.1 | Sphingobacterial es bacterium | MPSIHVGLSLNLDGNLLSAALPLLQKGEVEAFEWSFDALYHRAATPLPGWFEALLDTFSKEHRLIGHGIVFSLFSGKWLPEQNLWLQQLSRIARRYNFAHVTEHFGF<br>MTGEDFHKGAPLGVPLTAATLRLGQDRLCRLQDACSCAVGLENLAFAYHIDEVKKHGDFLQQLVQPVNGFIILDLHNVYCQSVNFNIPFAELLSLYPLHLVREIHLSG<br>GSWAQVAHKTVRRDTHDAAVPAAVWDLLQTALPLCENVQWVILEQIGISLNTSEAQQLFQNDFVRMSQCVAQYCTLQNPAASPVKNFAPPLLSQIPSEIPEDDDLY<br>QQQMELSRILETAPDVEQARALLGNSVLAHSAWAVEKWDNAMLATAMQIAQKWKDGFERFFK |
| MBK9205492.1 | Candidatus Obscuribacter sp. | MATEQLFAQDKVDALEFNFDQPLNGIVLAPWCQELLDKFSSASALIGHGVNLSLLSAQFTKRQQEWLEYTREEFKSRRYVHASEHFGFSEAGPIAQAAPLAVPMDA<br>QSLYLGKEMLKRYADATQCPVGLENLAFAFSLEDVKRQGEFIDQLISSVDGFLLLDLHNIYCQVANFDLDALQLLHSYPLSKVRELHLSGGSWSPSLSGKRIAVRRD<br>THDDAVPQESLILLLLR |
| MBK9142105.1 | Candidatus Melainabacteria bacterium | MVGKIESRHVSRTSPLVGLSLMPTEDFRDFAWNLVDREMVDVIEWSFDMIWEDGEMTPSWCESLLDDFSARGRLLGHGVTFSPFSGGFSEHQKWWLEQLRVEC<br>DRRRYIQISEHFGFMEAGAFVESAPMPVPMTEGAISLGRRCLEALAQSSGRPVGLENLALAFCLEDVRWQGKFLEEVLSRVDGFLVLDLHNLYCQSVNFDVPMSEL<br>LGSYPLERVRELHISGGSWSARRSGTGPSIRRDTHDGPVPERVFEILPEALDRCPALEYVIFERLGGTIDETGGTEELIRDYERILETVDG |
| MBP7863127.1 | bacterium | MPEIKVGLHAMPTDDFRQAATELILSDQVDVLECTFDAHWQLAENKISDYQYPDWYQSLIQVYSDENSLLGHAILLSILSAKTSARQLCWLEKLERECIERNYLHVSE<br>HFGFFEADGFIDGAPLPVPYTKDAVMIGRAMLKKMAQSSNKDVGLENLALAFSKTEAQEQGAFLNDLLDKPNNFLLLDLHNIFCQIANFNLNAKDHLSLYPLHKVREI<br>HLSGGSYSKSWSGKRDAIRRDTHDGPVPAEVFDLLDLALTLCPNLEFVILERLGNTMDGEDDFAQFRADYKRMYDLARGK |
| MBK7746242.1 | Candidatus Obscuribacter sp. | MATEQLFAQDKVDALEFNFDQPLNGVVVAPWCQDLLNKFSSASALIGHGVNLSLLSAQFTKRQQDWLTYTREEFKTRRYVHASEHFGFSEAGPIAQAAPLAVPMD<br>AQSLHLGKEMLKRYADATQCPVGLENLAFAFSLEDVKRQGEFIDQLIASVDGFLLLDLHNIYCQVTNFDLDALELLHSYPLSKVRELHLSGGSWSPSLSGKRIAVRR<br>DTHDDAVPQEVFNLTALALKLCPNTQFVILERLGNTMLEPELQEEYRDDFDTIREILDYSYD |
| MBK9070561.1 | Myxococcales bacterium | MIDAAAARPNSRQIKVGLTLMPSKDWLVAAKPLLIGNYVEALEWSWDVETTQPPWALGLIEYFAGQGALYGHGVMYSAFSAEFEPRQQAALEKTATLRQLASAVHI<br>TEHVGFSTIKGMLRGAPLPSPPCRASVQTARDQLGRIADAVGRPVGVENLALAISMDDVALQGDMIEEILVASDAPFVLDLHNLYCQAINFSIPAEQLLRALPLHRVR<br>EIHVSGGRMHAVAGGATKRRDTHDQLVPEDVWTMLRQCLPQCPRVEVVFLERLAGTLQGQADELFLDFQRMKREVDTFLTASSDLHAGREFGDEARVTRQPLPH<br>PSEAHELAAYQQAFVGLLRKLPSVVTAHHASELRTALAAATPSFAGYAISINDDMLIIAHAAFSRWHKVF |
| MBK8564694.1 | Saprospiraceae bacterium | MLLPSIACNLDKNLLQTALPLFESGKVEGLEWSFDALFKRPNIPDWFQELLLAYANEGRLIGHGVYFSLFSGKWMPEQAEWLTQLRQFAKAYRFDHVTEHFGFMTG<br>EDFHKGAPLSIPFTNSTVAIGIDRLKRIQDACNCPVGLENLAFAYGLEEVKKHGEFLEKLIEPVNGFIILDLHNLFCQLHNFEVGFDEILALYPLQRVREIHISGGSWEDS<br>MVAPNRKVRRDTHDEPVPETVFDLLEKTIGRCPNLKFVVLEQLSNALNTEEEQLAFRQDFWKMNHIIVQHNAPIIAPTNDFMPMERLMLASQPAEDLLLAEQQAELS<br>NILETAANYEDGHQRLLNSTLENSAWQVENWNPAMLETAIAIARKWKQGF |
| MBK9447954.1 | Bacteroidota bacterium | MIHASLSFNVDSNLIQAALPLFAAEAVEGMEWSFDTLHNRPDIPEWFEELLKSYGNAGRLVGHGVFFSMFSGKWLPEQAAWLAELKSLTTRFRFDHISEHFGFFTG<br>ADFHKGAPLSVPYSKRTLALGQDRLLRIADACGCPVGLENLAFATDAHSARIHGEFLEALLAPANGFLILDLHNLYCQLHNFGLDSAELLAGIPLARVREIHISGGSWE<br>PSPFRPQPGVRRDTHDAGVPERVFELLATTMGQCPNLKFVVLEQMSHALADAESQSQYQADFLRMRDLVQLESAKRTSFTLLQNFLPPKIEFQSHPLEDETLHFE<br>QLLLSDILENSANFEEVKAKIQSSALASGEWSAVNWDPAMLETARQIARKWKDGF |
| MBK9619416.1 | Candidatus Obscuribacter sp. | MQTNKPKLGLSLMPTEDFRMATEQLFAQDRVDALEFNFDQPLNGVVIAPWCQDLLDKFSSASALIGHGVNLSLLSAQFTKRQQEWLEYTREEFKSRRYVHASEHF<br>GFSEAGPIAQAAPLAVPMDAQSLYLGKEMLKRYADATQCPVGLENLAFAFSLEDVKRQGEFIDQLISSVDGFLLLDLHNIYCQVANFDLDALELLHSYPLSKVRELHL<br>SGGSWSPSLSGKRIAVRRDTHDDAVPQEVFNLTALALKLCPNTQFVILERLGNTMLEPELQEEYRDDFDTIREILDYSYD |
| MBL9023440.1 | Myxococcales bacterium | MTPKVGLSMLLEDDLRLAQAPLFAEGIVEAIEWTIDCELGRATPPWVEPLLDHYAAAGALYGHGVHFSPLSAIFEPRQAAWLEHAKGALAKRRYQHVSEHFGFMTT<br>PGMTRGVPLPVPYTKGALELGRDRLARLRAATSGAPVGLENLALAMSRDDALVHGEFLDALLEPHDGFLLLDLHNLYCQAHNFDVPWGTLLGAMPLDRVREIHVS<br>GGSWYEVATPAGPQPFRRDTHDDAVPDEVFGLLELALPRCPGVEVVFLERLGGTLADASAGEAYREDYRRLVSVVSRFAKGAAAHG |
| MBL0015158.1 | Bacteroidota bacterium | MIHASLSFNVDSNLIQAALPLFAAEVVEGMEWSFDTLHNRPDIPEWFEELLKSYGDAGRLVGHGVFFSMFSGKWLPEQAAWLAELKSLTTRFRFDHISEHFGFFTG<br>ADFHKGAPLSVPYCKRTLALGQDRLLRIADACGCPVGLENLAFATDAHSARIHGEFLEALLAPANGFLILDLHNLYCQLHNFGLDSAELLAGIPLARVREIHISGGSWE<br>PSPFRPQPGVRRDTHDAGVPERVFELLATTMGQCPNLKFVVLEQMSHALADAESHSQYQADFLRMRDLVHLETERRSPNRILRDFLPSTIRLPKKPLEDTMLYSEQ<br>MLLSQILENSASYDAANAAIQSSALASGEWSAVNWDPAMLETARQIARKWKDGF |
| MBK9771754.1 | Candidatus Obscuribacter sp. | MQTNKPKLGLSLMPTEDFRMATEQLFAQDKVDALEFNFDQPLNGVVIAPWCQELLDKFSSASALIGHGVNLSLLSAQFTKRQQDWLTYTREEFKTRRYVHASEHF<br>GFSEAGPIAQAAPLAVPMDAQSLSVGKEMLKRYAEATQCPVGLENLAFAFSLDDIKKQGDFIDQLISSVDGFLLLDLHNIYCQVTNFDLDALELLHSYPLSKVRELHL<br>SGGSWSPSLSGKRIAVRRDTHDEAVPQEVFNLTALALKLCPNTQFVILERLGNTMLEPELQAEHRDDFDTIREILDYSYD |

| | | |
|---|---|---|
| WP_202092815.1 | Chryseobacterium endalhagicum | MKKPLLGLSMMPEPEFVSAILPVLQNNSVDVLEWSFDTLFDSEEPEWLSGLLDFYAENGRLLGHGVYYSLFDARWTERQEIWLEKLKKELQLRKYNHITEHFGFMNTENFHQGVPLPVPLHPKILQIGKDRLRRLQDAVEIPVGVENLAFSFSVDDVKEQGAFLSQLVEDIDGFLILDLHNIYCQSCNFEIDMREIINMYPLEKVKEIHLSGGSWQESAYGKKPVRRDTHDDRIPEAVFEILPEVLSQCENLEYAIIERLGHTLNTETDQQIFFDDFKRVKEIADAAGNSSGQENIWNKKEEGAHVKPVEDLLLFDEQTKLTQLLFEERSSQSVKDQDFHYFKPENWDEEMITTAQQIIKKWNPY |
| MBL7912957.1 | Bacteroidia bacterium | MLPWVGISLMPEADFIEAALPLFQNGEVDVIEWSFDTLRSNSKEPQWLSLLLNEYSKNGRLIGHGVRYSLLKAQWDSNQEKWLKQLSTAVKKYNYRHVTEHFGFMSSDDFHKGTPLPVPLNKTTLAIGQDRLKRLQNISQLPVGIENLAFALSTKQVKEQGEFLKKLLEPVNGFLILDLHNIYCQASNFKIDPIELVKSYPLDKVREIHVSGGSWADVSTSKNKVRRDTHDEAVPKEVFELLKSALPLCKNVEYIIFERLGNTIKTEKDKLQFAKDFKKIKKIVATQKKNENSPSERLTKKIIKHTVAKEPLESKVLRKQQQLIIETLLKHKDVAKATKVLTKNKALKDWDPALWKNYMVETAMVLIKKWN |
| WP_202271905.1 | Chryseobacterium sp. KMC2 | MGRPLLGLSMMAEADFVSAILPLLQDNAVDVVEWSFDTFFAAEEPAWLKDLLNFYAGNGRLLGHGVYYSLFDARWTERQEMWLQKLKHEVQKRNYNHITEHFGFMNTENYHQGVPLPLSLHPKTLEIGKDRLYRLQDAVSIPIGVENLAFSFSVDDVREQGVFLDKLVDDIDGFLILDLHNIYCQSCNFDVSIQEIVSLYPLEKVKEIHLSGGSWQDSVYGKKMIRRDTHDDVIPEEIFSVLPSVLSQCRNLEYVIIERLGHTIHTEEEKQRFFNDFVRVKSILNDSDGMKRKKEKWVKREMLLPEPVEDSILYEEQIRLTKLLFDHTDPTLIKNQNFHYFKTENWDPEMILTAQNIIKKWNPY |
| MBL7922055.1 | Bacteroidia bacterium | MNPWVGISLMPEADFIEAALPLFQNGEVDVIEWSFDTIPPKVKEPEWLKLLLKEYSQNGRLIGHGVHYSLLKAVWDSNQDKWLKQLATEVKKYNYRHVTEHFAFMTSDDFHKGTPLPVPFNKTTLAIGQDRLKRLQNVSQLPIGIENLAFAFSHAQIKEQGEFLKKLIEPVNGFLILDLHNIYCQSHNFKVDVVKLVNSYPLDKVKEIHISGGSWANVESSKNKVRRDTHDEAVPEEVFELLKEVLPLCKNVEYVIFERLGNTIKTQKDKLQFAKDFRQVKKIVLSQRKKEENLLEEITKTKIDHKVSLNPLEDENLRKQQQFIIETLLVNKDVAKTIQLLQNKKINSWQPELWTDYMVETGMVLVKKWN |
| MBL7782349.1 | Saprospiraceae bacterium | MLYPSIACNLDSDILLTALPLLESERVQGLEWSFDTLYQVRDIPAWFDELLLEYGRAGRLVGHGVFFSLFSGRWSPEQAQWLKRLKKQCDIFHFDHITEHFGFMTGADFHSGAPISLPFTASTLAIGHDRLRRIQEVCACPVGLENLAFAYSVEEVKRHGDFLNKLMEPVNGFIILDLHNLYCQAHNFDIPADELLPLYPLDRVREIHISGGSWENSSLLPDRPVRRDTHDDAVPQEVFQLLKNTIPLCPNLKFVVLEQMGTALHTEASRRQFRQDFDTMHTILEAYESPDNYSGNAFIPLPFPALGAPLEDADLHRQQRQLSEILEKAGSFEEAQRLLAVSDLAGTAWGTEHWAPYMLETATRIAQKWQKGFRQPE |
| MBL7682615.1 | Flavipsychrobacter sp. | MLQACYPLFEASKIEAIEWSFDALYKRRNIPDWFLQLLHAFANEGRLIGHGVFFSLFSGRWTYEQQAWLKHLSKVSSAFRFDHITEHFGFMTGADFHSGAPLSIPYTNTTLTIGKDRLARIYDACGKPVGLENLAFSYDIDEVKRHGDFLSKLIEPVNGFIILDLHNLYCQLQNFSVGVDEIINLYPLERVREIHISGGSWEDSIIDIGRKIRRDTHNDAVPQEVFALLEYAIDKCPNLKYVILEQVGGALHTQQSREDFAADFLKMSAIVKQAKGLPPETNDFLPKELIIPALVAEDEELHQQQTELSEIFETAAGYDDALMRLQQSSLANSDWQIENWAPYMLETAISIAQKWKK |
| MBL7707133.1 | Taibaiella sp. | MLSASLPLLEAGLVGGLEWSFDALYARDYQVPDWFHELLSAYSKEQRLVGHGVFFSLFSGRFSEEQQAWLERLRALRGMYQFDHITEHFGFMTGRDFHKGAPLSVPYTATTLAIGRDRISRIADAAQCPVGLENLAFAYDLEEVKIQGRFLDELLSPVNGFMILDLHNLYCQLHNFNLDSSTLLDLYPLEKVREIHISGGSWQPHPGAHLQQVRRDTHDDAVPAAVFELLKITLNRCPNLKFVLLEQLGIALTTPAQRAQYQDDFHTMRNIIEQYNREQQWPRNIDGFRPATLQLTPTPLVDEQLFGHQQIISHILETAADHKDVIDKMQYTSLAGSDWHIEKWQPAMVDTAIHIAQKWK |
| WP_204662631.1 | Dyadobacter sandarakinus | MSEIYSSVVCNLDMHMLRASLPLFEQEKVEAIEWSFDTLFRVEEVPAWFTNLVSEFSNHHRLIGHGVYFSLFSGKWAPEQHDWLIKLRQLSREFHFDHITEHFGFMTGEDFHKGAPISIPFTSTTLALGKDRLRRIQDACQCPVGLENLAFSYSLEEVRKHGDFLGRLVESVNGFIILDLHNVYCQVHNFNIPFDMLIELYPLDRVREIHISGGSWDETLAGPGKPVRRDTHDDYVPEVVFDFLNKAIPKCPNLKYVVLEQMGTALSTPADQLAFQSDFLRMDNIVKSWAPKMLQSPADTFAPIPGIASATPLEDPELHGQQMQLSRILETAGNAGEARSMLLSSSLHNTAWEVENWAPYMLETAISIAQKWKNGFA |
| MBN8684453.1 | Chitinophagales bacterium | MESERVEAIEWSFDALYSFEQIPEWFTDLLSEFSQAGRLIGHGVFFSLFSGKWLPEQSQWLESLRQLSQAHHFDHISEHFGFMTGADFHRGAPLSIPFTPGTLAIGQDRLKRISEACGRPVGLENLAFAYSLDGVRRHGAFLDELLEPINGFLIFDLHNFYCQLHNFEQDVDTLIRNYPLKRVREIHISGGSWEISEARPERPVRRDTHDHAVPESVFELLEMVMPRCPNLKYVVLEQLGPALNSEKERLDFQADFFKMEQAVQNFNRQQQGREIHDFLPMKYTLNAVPLEDEALYQQQIQLTDILENAGDLKSAQAQLIACGLADSDWQARYWAPEMLDTAINIARKWKNGF |
| MBN9380196.1 | Chitinophagaceae bacterium | MEGSRIDALEWSFDALYREEEIPDWFRDLLTAFANEHRLIGHGIFFSLFSGRWLAGQEEWLHQLRRVCQDYRFDHITEHFGFMTGKDFHHGAPLGVPYTPTTLRIGRDRLKRIYDACRCPVGLENLAFSTSAEEVKRHGVFLEQLLEPVNGFIILDLHNLYCQLHNFGIAFEDLIDLYPLHRVREIHISGGSWDNSKANPDRTVRRDTHDNAVPQEVFRLLENAIERCPHLKYVVLEQLGVGLQTAAARKTFYDDFLKMQSIVQEKSKTLDPGTVDTFLPGSVLLTEIKAEDETLHRQQLELSRILETAVSYEEVIHRLNNSSLVSSDWKIEAWEPCMVETAMKIAQKWMRGV |
| MBN9166157.1 | Myxococcales bacterium | MHPWIGLSLMLEDDFRLAAGPLFAEGVVDVLEWSFDTGWGRAIPEWADALLDHYGEAGRLLGHGVHYSAFSARWEERQARWLERLAGEVARRTYVHVSEHYGFMTAAPFMRGAPLPVPRGEASRSVGRDRLERMRAVLAAGRNDACPIGLENLALAWNRDEALAHGAFLGEVLDHAEDFIVLDVHNLHCQIENFDIDPDELLDSFPAARIRELHVSGGSFLPAWPHHPEETVRCDTHDGDVPAPVFDLLERALDRFPDVRAVILERLGGTLRTQADIDTFRRDYLRVKQIVDHSRDGEGARDGDG |
| WP_207426252.1 | Pedobacter sp. SYSU D00535 | MKTVYSSVACNLDRDILAATIPLLAESRVEAIEWSFDALYKIAEVPDWFQELLHTYSKENRLIGHGVFFSIFSGRFLPEQKEWLKQLRQTASHLKFDHITEHFGFMTGPNFHHGAPLNIPCNEKTLAIGRDRLARIQEACHCPIGLENLAFAFSLDDVQRQGAFLEKLVEPINGFIILDLHNLFCQLHNFQLDFDQLIQLYPLHRVREIHISGGSWESSEHLTNRQIRRDTHDDLVPEEVFQLLELAIPKCPELKYVVMEQVGAGLKTAESKTGFYADFLKMEAIVKRAVQLEGAAPVNRFLPEDILLEAQPVEDEALYQQQLELSEILETSSSYAETYSRVQHSSLSKSDWNIESWKPWMLETAVALANKWK |
| WP_208173464.1 | Hymenobacter negativus | MDETPPILASLACNLDSDILTAALPLLEAGQVEALEWSFDTLFRANELPEWFTELLHAYGNQQRLVGHGVFFSLLSSRWTPEQQHWLTQVRQVSATFAFDHITEHFGFFTGQNFHSGAPLPVPYTATALRIGQDRLRRIYDACQCPVGLENLAFAYSLEEVEQHGEFLEKLIAPVNGFIILDLHNLYCQLHNFSVSYEALIDLYPLDCVREIHISGGSWDNSAVLPHKKVRRDTHDDAVPEEVFRLLALTMPKCANLKFVVLEQLGTGLQSESSKAQFQRDFQRMANLVQQHRTRFLNHELANTFRPLQLPVLGSAIEDSQLHTQQVQLSDILENAATFEEAHQRLHASSLANSAWKIEAWEPHMLETAVRIAQKWKK |
| MBO3269797.1 | Hymenobacter defluvii | MPSILSAIACNLDDHLLAAALPLLEAGQVEAVEWSFDTLFDGRQLPNWFAELLTAYGEQGRLIGHGVYYSLLSGRWTPEQAQWLEQLRLLAARFHFDHITEHFGSFTGQNFHTGAPLPVPYTAATLRLGQDRLLRVADACQCPVGLENLAFAYSLEEVQRHGEFLEKLLAPVNGFLILDLHNLYCQLHNFNLSTDSLLALYSLHRVREIHISGGSWEYSQVEQGRQIRRDTHDEAVPDEVFALLERTIPQCPQLKYVVLEQLGTGLRTEASQHQFRQDFQRMAAIVEASRHRAPSHQPNAFQPPAALPTGPAAHDEALHAQQRQLSQILETAASYEEAQQRLRASSLAHTDWQLEQWAPHMLETAISIAQKWK |
| WP_207177520.1 | Cellulophaga sp. E16_2 | MRKRIMGNKKPKLGIAIAPSQKFLEAALPLFAEEKIEIIEWSFDTLKDAADEPAWLSLLLKEYGEKNRLLGHGVFYALLDANWSSRQENWLKKVRQETLSHKYCQISEHFGLMSSANAHSGFPLPIPLSNPVLQIGIDRLKRLQATAQVDVGIENLALTANVADILEQGEFLFKLVHPVNGFVILDLHNIYCQSENFNIDMMTIIKSYPLSLVKELHISGGSWDTDPTLSRKIRRDTHDGRIPEVILEVLPEVLKICPFLAFIIFEKIEDSFLTEKDGIDFRADFQKIREIIDATSFSVTPKEKKQVAIILGPPVIDVELLQAQVALRESIALDTYQTNSEWDKDMWKVATKLYEKWNN |
| WP_207331299.1 | Fibrella forsythiae | MIRSTIACNLETNLILASLPLFEAERVEALEWSFDTLYAHQQLPDWFVDLLTTYSEAGRLIGHGVFFSLLSGRWTHDQQNWLARLHALSRDYQFDHVTEHFGFMTGEDFHKGAPINIPCTPVTLALGQDRLKRIQDACGCPVGLENLAFAYTLDDVKRQGDFLRQLIQPVNGFIILDLHNLYCQSHNFSIMHDDLMALYPLDRVREIHISGGSWSGSETQPGLLIRRDTHDDAVPDDVFNWLSTTIDQCPNLKYVVLEQLGNGLTTEGQQQAFRADFHRMDTIVQEKNKPATTTTMNSFLPQLPLIINAMPLENHRLHNQQVELSAILESSTDVSQAQTRLAQSSLANSEWEVERWEPSMLETAVSIARKWKDGFVSKK |
| WP_207434928.1 | Sabulibacter ruber | MPNLLPSLACNLDLDLLAAALPLLEEARVDGLEWSFDALFQTRQVPAWFLDLLTAFGQEKRLVAHGVFFSLFSGKWSEGQRQWLKTLSEASQRYRFDHVTEHFGFMTGQNFHHGAPLAIPYTATTLAIGQDRLARIQEAAQCPVGLENLAFSYSVEDVKRHGEFLEKLIEPVNGFIILDLHNLYCQLHNFSVSFEELIPFYPLERVREIHISGGSWEDSLTTPAKKIRRDTHDNGVPEEVFNLLTQTLPLCPNLKYVVLEQLGNSLKTEQSKTQFYQDYLRMEAIVRDFNATHLNPIPEPERFLPPQATFSAELVEDEALYAQQVELSSILETAGSYEEAQQKLRTSTLAHSAWQVEQWEPAMLETAISLAQKWKKK |
| WP_291898394.1 | Chryseobacterium sp. | MKRPMLGLSIMAEADFVSAILPLMQNNSIDVLEWSFDTLYHTHEPDWLRDLLNFYAENGRLIGHGVYYSLFDAKWTERQEEWLQKLKEEVRLRKYNHITEHFGFMNTENFHQGVPLPVPLHSTILQIGKDRLCRLQDAVNIPVGIENLAFSFSMDDVKEQGIFLDRLTEDTNGFLILDLHNIYCQSCNFEVGMQEIINLYPLDKVKEIHLSGGSWQESVYGKKQVRRDTHDDVIPQDILSILPSVLKQCQNLEYIIIERLGHTLKTEQEKVDFLNDFNTVKMIIETSELKLREGSTWSKKEMKLSEKPLEDLVLYEEQSRLTKLLFDKAGSEVIQGQDFHYFKTEKWDPEMILTAQNIIEKWNPY |
| WP_291193020.1 | Dyadobacter sp. | MSEIYSAVACNLDTHILRAALPLFEQEKVEAIEWSFDTLYKFDEIPPWFTDLVSEFSKQNRLMGHGVYFSLFSGKWSSGQQEWLRKLKKLSAEFNFDHITEHFGFMTGEDFHKGAPIGIPLTPRTLALGQDRLQRIQDACQCPVGLENLAFSYSIDEVRKHGVFLDQLVESVNGFIILDLHNIYCQVHNFNIDFVDILKLYPLNKVREIHISGGSWEDTATPQGKPVRRDTHDESVPEIVFEFLGNAIPQCPNLKFVVLEQMGTALSTYASQAAFQSDFLRMDSIVKSLARKFDPIPERMFAPLAASLNAIPLEDADLYQQQIQLSYILETAGNVSEARAMLLASDLRNTAWEVENWAPYMLETAISIARKWKNGF |
| MBO9594034.1 | Niabella sp. | MAEIHSAIACNLDADILSASLPLFESGSVDAIEWSFDALHQRQELPEWFRALLTAFSRSNRLIGHGIFFSIFSGKWLPEQEQWLKSLKKYAADFHFDHITEHFGFMTGKDFHHGAPLSIPFSPVTLAIGIDRLQRIQEACRCPVGLENLAFSYSLDDVKRHGAFLEKLVEPVNGFIILDLHNLFCQTHNFNVDFEPLLACYPLDRVREIHISGGSWEPSVAVPSTIIRRDTHDEKVPDTVFDFLRTTIPRCPNLKFVVLEQLGTGLRSEASRVAFRDDFLQIQRIVKAASLRATSINTFQPPSFQLRTTPVEDSLLYRQQTALSRILEEAGSEAEARSQLEASVLAHSDWQIEHWNAAMLETAVHIAQKWKNGFTPDA |
| WP_209917743.1 | Chryseobacterium jejuense | MGRPLLGLSMMAETEFVSAILPLLQSNSVEVLEWSFDTFYDAEEPDWLSDLLDFYSENNRLIGHGVYYSLFDARWTKRQEIWLKKLKEEFRKRNYNHITEHFGFMNTENFHQGVPLPVSLHSKTLEIGKDRLYRLQDAVQVPVGIENLAFSFSLDDVMEQGIFLDKLTEDTNGFLILDLHNIYCQSCNFEVEMQEVIKLYPLDKVKEIHLSGGSWQESIYRKKPIRRDTHDDAVPEDILSVLPSVLSQCKNLEYIIIERLGNTIDAEEKKESFLDDFTKVKKLIDLSDVGTPNKDLWAKKEIRFSENPLEDMILHEEQTLLTKLLFDNAGPASIKSQEFHYFKTKNWDPEMILAAQNIIKKWNPY |
| WP_209784810.1 | Chryseobacterium sp. PvR013 | MLGVSMMAEADFVSAILPLLQNNSIEVMEWSFDTLYHAHEPDWLHDLLNFYAENDRLIGHGVYYSLFDARWTERQEEWLKKLTEEVSLRKYNHITEHFGFMNTENFHQGVPLPVSLHPKTLEIGQDRLYRLQEAVNIPVGIENLAFSFSIDDVKEQGVFLDKLTEGTKGFLILDLHNIYCQSCNFGVPIQEIINLYPLDKVKEIHLSGGSWQDSVYGDKQVRRDTHDDVIPQDILSVLPSVLTQCQNLEYIIIERLGDTVKTEEEKDNFLNDFNTVSKIIETSDWKVKGKNGWSKREMKLSEKPLENLVLYEEQSRLTKLLFDNVEVKAIKDQDFHYFKTENWDAEMILTAQNIIKKWNPY |
| MBP6746809.1 | bacterium | MLTEDFRQATGALFRSGKVEALEWSFDFGFNGAVEEPWGNDLLDKFSVAGSLTGHGVELSPLSATFSERQRQWLAQAREEFQKRKYLHASEHFGFSEAGPIEKGAPLSVPMNEQSLRLGKEMMKRFADATQCPVGLENLAFAFSLDDVKRQGDFIDQLISEVDGFLLLDLHNIFCQVANFNMPELELLNLYPLSKVRELHLSGGSWSRSVSGGRLAVRRDTHDDAVPQEVFNLTALALKLCPNVEFVILERLGYTMMEPEQQQEFRDDFDTIREILDHCYV |
| MBP1164381.1 | Chryseobacterium sp. PvR013 | MRKPMLGVSMMAEADFVSAILPLLQNNSIEVMEWSFDTLYHAHEPDWLHDLLNFYAENDRLIGHGVYYSLFDARWTERQEEWLKKLTEEVSLRKYNHITEHFGFMNTENFHQGVPLPVSLHPKTLEIGQDRLYRLQEAVNIPVGIENLAFSFSIDDVKEQGVFLDKLTEGTKGFLILDLHNIYCQSCNFGVPIQEIINLYPLDKVKEIHLSGGSWQDSVYGDKQVRRDTHDDVIPQDILSVLPSVLTQCQNLEYIIIERLGDTVKTEEEKDNFLNDFNTVSKIIETSDWKVKGKNGWSKREMKLSEKPLENLVLYEEQSRLTKLLFDNVEVKAIKDQDFHYFKTENWDAEMILTAQNIIKKWNPY |
| MBR9919930.1 | Bacteroidota bacterium | MPKVLASIACNLDSDMLLASIPLFASEKVQAIEWSFDTLYKHRNIPDWFVELLRTYSDSGRLIGHGVYFSLFSGKWSADQQAWLDRLSRLSANFNFDHITEHFGFMTGADFHKGAPLSLPFTSNVLNIGRDRLQRIAQVCQCPVGLENLAFAYTLEEVKRHGEFLNKLLEPVNGFIILDLHNLYCQLYNFEQDYEDIIQLYPLDRVREIHISGGSFEQSGIQDGKKIRRDTHDSAVPEAVFELLEKTLDRCPNLKYVVLEQLGIGLQTEDSKKQFQEDYFRMKKMLDTFNRRTTEFPNQDFNPPQKEPKTVPLEDLKLHKQQLELSAILESSESLADVNTKLRQSSLAKSPWQIENWAPEMLETAWCIARKWKAGFQ |
| MBS1663462.1 | Bacteroidota bacterium | MPRLLPAVACNLDADILAACLPLLEESRVESIEWSFDALFNIKQVPDWFLELLSAFSKEGRLIGHGVFFSLFSARWHSQQEAWLRQLQETSSLFRFDHITEHFGFMTGKDFHRGAPLNIPYTASTLAIGRDRLQRIQEACHCPVGLENLAFSYSLEEVKKHGEFLDRLLEPVNGFIILDLHNLYCQLQNFDLDFEELISLYPLDKVREIHISGGSWEDSDIAPGKRVRRDTHDDSVPGDVFQLLENTIDRCPHLKYVVLEQMGSALRTDDSRAGFRSDFSRMEAVVHRFNDDRIPELVDMSFLPPLHDISHPPVEDEQLYRQQLELSEILETAASYTEAAQRLSRSSLANSEWRVERWQPHMLETAVNIARKWARGWARQGS |
| MBS2001764.1 | Cyanobacteria bacterium SZAS LIN-5 | MPSDDFREASRSLFENNKVEVLEWSFDFSWLGNEIAPWCADIIDRFSKSNKLIGHGVTLSPLSARFSKRQEDWLARVRDEFKTRNYIHASEHFGFSEAGPISHGAPIAVPMTAESLHAGKEMLKRYAEATQCPVGLENLAFAFSMDDVKRQGEFIDRLISAVDGFLVLDVHNLYCQVANFDVPITELLNSYPLTKVREMHLSGGSWTESESRKRFAVRRDTHDDNVPQEVFNFAALGLTMCPNLEFVILERLGWTMLDDESQVGFREDFETVAEIVDATYSKEIQGADSFAQTGKSPSVEAQTESFTVDTSKESPSPSASTKSPSLTASTQAPALVSKSEPSLADYQDQLVELLSSNRTNSEILEILKSASQFRHYQDYISSFDPDMVAVACELMGKWARRADS |
| MBP9092271.1 | bacterium | MNYRVPKLGLSLMPTEDFRQATDELFQSGKVEALEWSFDFSFNGVIAEQSCNVLLDNFSSAGALTGHGVELSPLSAKFSKRQSEWLAHAREEFQARKYTHASEHFGFAEAGVIERGAPLAVPMNEQSLRVGKEMMKRFANATQCPVGLENLAFAFCLDDVKRQGDFIDQLISEVDGFLLLDLHNIFCQVANFNMSELELLNLYPLSKVRELHLSGGSWSRSISGARLAVRRDTHDDAVPQEVFNLTALALKLCPNVEFVILERLGYTMMEPEQQQEFRDDFDTIRELLDYCYD |

| | | |
|---|---|---|
| WP_215237124.1 | Dyadobacter helix | MSGILPAIACNLDINILRAVLPLLEEEKVEAIEWSFDSLYKYDEIPAWFESLMQEFGDQGRLVGHGVFFSLFAGKWLPQQQHWLDRLRILSKTYRFDHITEHFGFMTGEDFHKGAPISIPFTAQTLALGRDRLARIQDAASCPVGLENLAFAYSVEEVKKHGQFLDQLVQPVNGFIILDLHNLYCQTHNFNITHHELMALYPLEKVREIHISGGSWDESPFDHKEKIRRDTHDHAVPAEVFELLKIAVSRCPHLKFVVLEQLGTGLDTEESQIKFQEDFMTMRKIIEESNDGSRIPYSFIPLSVNSPDAVPAESFMLFDQQRVLSEILETSSDYEEVRMRLNTSILAHSDWGIESWKPEMLYTAMSIAQKWKNGFA |
| WP_215234353.1 | Dyadobacter linearis | MSRIFSAIACNLDTHILQASVPLFEMEKVEAIEWSFDSLFKVETIPEWFTDLVKEFSDHNRLIGHGVYFSLFSGKWSPEQQNWLSQLRKLSGEFHFDHISEHFGFMTGSDFHKGAPLGIPFNKSTLAIGRDRLARIQEACNCPVGLENLAFSYSYEEVKKHGDFLNQLVESINGFIILDLHNLYCQCHNFDIDFQEMLSCYPLDRVREIHISGGSWDNGSEERTVRRDTHDDSVPETVFEYLEKALPRCQNLKFVVMEQLGTALDTVEKQIAFQHDFVKMDGIVKSFDGDPGSNQTNSFLPFGKHPLDMQPVEDIALASQQMQLSAILETAVNLDHAREMLNASDLKHSDWHIEQWQPHMLKTAIDIAQKWKGGF |
| MBP9092304.1 | bacterium | MLQEDFLAATLPLFSNDEVEILEWSPDIGWAAPTMPEWANELLDFYSSANSLYGHGVNYSPMSAEWTAVDSEWIENLKRELEARKFVHFSEHFGFSRIGHLQQGAPLPVPFTTEAIRAGQFKLEKLAHTTGGRIGLENLALAFSKRDAMQQGRFLEDLLRPFDGFVLLDLHNLYCQMENFSLSAEELLGTYPLERVREIHISGGSWSESQAAQTKRKSKPTLAPASVTRIRRDTHDNEVPDEVLALLPLVLANCPNVEAVILERIGGTMPTAEQQARFRDDYRAIAKLIELHTLRQTVGDDAEAMAKTHITQVELSQGRGTLSRECSQSSDQTEILRYQKKLMELLQSEQSPERIIAELSSAFEGELAEYANLSQPQMLAVGQELVKKWTQ |
| MBT9556781.1 | Myxococcales bacterium | MALSWMREPAFLRAATPLLADGLIDALEWSFDTAWRDEPAPVRDALLGHYASAGRLLGHGVSYSLLSADPADDVRKARWLERWRGEQARHPYRHLSEHIGFMTAPGYQRGPPLPMPLTPEVVTLGRDRLNRLVDASNGPVGLENLAFAFSRREVLEQGELLERLLAPTGGFVVLDLHNLWCQAVNFGLDPAALLDSYPLARVHELHVSGGSWAETSHGPVRRDTHDGTVPFEVLELLQLALRRCPEVRFVVVERLGDSFESPDAAAELRADVMAVRGTLSTFDGVTSSLAKPLPLALWPDVHRDDALAVAQSALYATLDSGPTEDDAWSTSEAVSAIRSMIEHAEPRMLATGEALMRRWGRRNTG |
| WP_213279122.1 | Chryseobacteriu m indologenes | MKKPLLGISMMAEVDFVSAVLPLLQNNTIDVLEWSFDTLYHNNEPDWLRDLLNFYAENNRLIGHGVYYSLFDARWTERQEEWLQKLKEEVSLRRYNHITEHFGFMNTENFHQGVPLPVSLHPKTLQIGKDRLYRLQEAVDIPVGIENLAFSFSVDDVKEQGVFLDKLTEDTNGFLILDLHNIYCQSCNFGVEMQEIIDLYPLDKVKEIHLSGGSWQESVYGKKQVRRDTHDDVIPEEIFSVLPPVLAKCQNLEYSIIERLGHTLKTEGNKENFLNDFNKVKAIIEASDGKEKEKRSWNKRQIQLPEKPLEDLALFEEQSRLTRLLFDGAGAGIIKDQDFHYFKTENWDHEMILTAQNIIKKWNPY |
| WP_212662956.1 | Acaryochloris marina | MTPQLGLSLMPQPNFWQAAQPLFDADEIDVVEWSFDMGWGGVVLPDLLCSVLRQFSQGNGLLGHGVSYSLLSAQLDRRHWLACLQAECLDYSYRHISEHFGWMATRNFARSAPLPMPLLPETLKLGCDRIQKFADIAQVPIGLENLAFAFGLQDVRQQGEFLDQLLDPVDGFLLLDLHNLYCQMHNFQLSATEIFSLYPLERVRELHLSGGSWSHHGTATIRRDTHDQAVPEPVFELLKLALQQCPQVETVILERMGNTLNSKQEQLQFRRDFAHIRQIINHVSHAAEAGGE |
| WP_212651256.1 | Chryseobacteriu m arthrosphaerae | MKKPLLGLSMMPEADFVSAILPLLQSNSVEVLEWSFDTFYEAKEPEWLDELLNFYSENQRLIGHGVYYSLFDALWTERQENWLKKLKEEVRKRKYNHITEHFGFMNTENFHQGVPLPVPLHPKTLLIGKDRLYRLQDAVQLPVGIENLAFSFSIEDVKKQGVFLDNLIADINGFLILDLHNIYCQACNFEVEMEDIIRLYPLDKVREIHLSGGSWQESVYARKMIRRDTHDDLIPEEIFSVLPSVITQCNNLEYVIIERLGHTIKTDQEKRAFLDDFTRVRSIIDDSEIKTGKKIRWGQKETGFSNSPVEDDALYEEQSRLTKLLFNSSGAEMIKEQEFHYFKTENWDPDMILTAQNIIKKWNPY |
| MBU6341249.1 | Bacteroidota bacterium | MVFPAIACNLDPDLIQTALPLFETEKVAALEWSFDAVFSSEPLPNWFHELIQVYSKGERLIGHGVFFSLFSGKWLPEQSEWLQNLKRYCQIYQFDHITEHFGFMTGADFHKGAPLSIPYNQSTLAIGRDRLLRISQACNCPVGLENLAFSYSLDEVKRHGAFLNELLEPVNGFIILDLHNVFCQLHNFDLDAKDLRALYPLERVREIHISGGSWEFTRIQPEKPVRRDTHDDSVPKAVFDLLKDVLPCCPNLKYVVMEQLGTALHTQAQRDAYRADFYTMERILTQAQPKIPAKNNFVPPVLPPLGNPIEDLALFEQQRILSNILENALDYQGALNQSLLANSDWDIEHWNPAMLETAVAIARKWKNGW |
| MBT7958730.1 | Akkermansiacea e bacterium | MPHVGLSLMPETQFLQASMPLIEAGDVEALEWSFDTGWKNGVRDWLSNLINVYSEAGRLTGHGVHFSLLSAEWTDRQQGWLDQFREEVTRHQYLHHTEHFGFMTAGDFHRSAPLPVPLTNSTLKLGRERFSQLAEVAPCPLGIENLALALSIDDALQQHEFLEKIVTPANGFLLLDLHNIYCQLCNFNLSPAELLDNYPFHLVREIHVSGGSWAPTASSNSPIRRDTHDDDVPAEVFSLLEHVLPRCPDVEVVFLERLGNTLQEHQFEQFRKDFLTLKSIVANG |
| WP_216444824.1 | Chryseobacteriu m sp. PCH239 | MRKPILGVSMMAEPDFVSAILPLLHNNSIEVMEWSFDTLYHTNEPDWLRDLLNFYAENNRLIGHGVYYSLFDARWTARQEEWLKKLKEEVSLRKYNHITEHFGFMNTENFHQGVPLPVSLHSKILQIGKDRLYRLQDAVNIPVGIENLAFSFSIEDVKEQGVFLDKLTEDTNGFLILDLHNIYCQSCNFELEIQQIIDLYPLDKVKEIHLSGGSWQDSVYRKKQVRRDTHDDVIPKDILSVLPSVMKKCPNLEYIIIERLGHTLKTEEERNYFLADFNKVRTIIETSDLKLKQQNGWCKEEIKLPAKPLEDLLLYEEQSRLTRLLYENANVENIKHLDFHYFKTENWDAEMILTAQNIIKKWNPY |
| WP_218701771.1 | Olleya sp. HaHaR_3_96 | MKKNKPKLGLSIMPNPEFIAAALPLFESAQVEVIEWSFDTILDKKYQPEWLPLVLKEYGDNNRLLGHGVYYAPLDANWGPNQDNWLQKAKLETLAYNYNHLSEHFGVMSSANAHHGFPLPFDLSDTILKIGIDRLKRLQNTVQLDIGIENLALASNVSDILKQGAFIEKLVAPINGFVILDLHNIYCQSINFDLDLLTIITSYPLHLVKEIHLSGGSWDHAPNLTKPIRRDTHDGRIPEVLLDLLPEVLQRCPVLEFVIFERLGDTFQNENDGLEFRADFNKIQAIIDHTAFSSNARQWTLKKHDLGPPLIDLPLLNKQQELREHIRLETANNHPEWNTNMWITATKLYKKWNT |
| WP_218736096.1 | Cellulophaga sp. HaHa_2_1 | MGNKKPKLGIAIAPSLKFLEAALPLFAGEKIEIIEWSFDTLKDAADEPAWLSLLLKEYGEKNRLLGHGVFYSLLDANWSSRQENWLKKVRQETLSHKYCQISEHFGLMSSANAHSGFPLPIQLSNPVLQIGIDRLKRLQATAQVDVGIENLALTANVADILEQGEFLFKLVNPVNGFVILDLHNIYCQSENFNIDMMTIIKSYPLSLVKELHISGGSWDTDSTLSRKIRRDTHDGRIPEVILEILPEVLKICPFLTFIIFEKIEDSFLTEKDGIDFRADFQKIREIIDATAFSVTPKEKKQVAAILGPPVIDVELLQAQVTLRESIALGTYHTNSEWDKDMWKVARKLYEKWNNY |
| WP_220614797.1 | Hymenobacter sp. HSC-4F20 | MSSGLPTTPAEPAILATLACNLDVDILSAALPLLAEGRVEALEWSFDTLFWAEQLPDWFSELLGTYSAENRLLGHGVYFSLLSGRWTPEQQQWLEQVRQVVARYPLVHLTEHFGFFTGQNFHAGAPLPVPYTPATLRLGRDRLSRLYEACRCPVGLENLAFAYSLEEVKRHGEFLGRLLEPVSGFLILDLHNLYCQLHNFSVPYEELIGLYPLEQVREIHISGGSWEDSALTPGRKIRRDTHDEAVPEDVFQLLELTIPRCPHLKYVVLEQLGPSLRTEVSRAQFRHDFGRMEAIVQRLRTGRAGYADQLFLPLHPPPTGPAFEDGQLYAHQQHLSHILETAPTLAEAQRLLHASALASSAWKLEQWAPHMLETAVSIAQKWKR |
| MBW4543370.1 | Symplocastrum torsivum CPER-KK1 | MFKFKRRTPMVGLSLMMESDFFQATQPLFETGAVEVLEWSFDMGWGVKAKPPWVEELLQFYSQRDRLTGHGVSYSLLSAQPGERQIHWLKSLQQECLDYRYRHISEHFGWMAAGDFYQSAPLPLPLTPETLQLGREHLKRLSDKANAPVGLENLAFAFGLQDVLQQGEFLARLLEPVDGFLVLDLHNLYCQVHNFQQSAEQLLERYPLQRVRELHVSGGSWSVTSKGQQTRTIRRDTHNGTVPEAVFELLALALKRCPSVEVVIFERIGHTLHTEIDIEGFRQDFWRIKQVVKESV |
| MBX2922030.1 | Chitinophagacea e bacterium | MLQESRVEAIEWSFDALYQVQEIPGWFEELLHAYSNENRLLGHGVFFSLFSGRWSDHQAEWLNHLKQTCTKFHFDHITEHFGFMTGADFHHGAPLNIPYTPVTLSIGKDRLSRIYDACRCPVGLENLAFSYSLEEVKQHGEFLDKLIQDVNGFIILDLHNLFCQLHNFSVGYEEMIALYPLHKVREIHISGGSWDITDTGTQKTIRRDTHDDAVPSEVFGLLEKTIPKCTHLKYVVLEQLGNGLRTEESRQNFYNDFLRMEDIVQKNNKKIRDHENFSFLPVHPPVPSIIVEDEKLHFQQMELSHILETALSYEDAVKKLNASSLACSDWKIETWNSAMIETARKIARKWMKK |
| WP_218779874.1 | Hymenobacter crusticola | MLASIACNLDADILSAAFPLLEEGQVEALEWSFDTLFWAEQVPSWFTELLTTYSNHNRLVGHGMFFSLLSGKWTLEQQQWLRQLHQLAAQFRFDHITEHFGFFTGQNFHYGAPLHVPYCEATLRIGQDRLRRIYEACQCPVGLENLAFSYSLEEVKQHGAFLEKLIKPVNGFLILDLHNVYCQLHNFSVAFEELVMLYPTDRVREIHISGGSWEGSVSAPGGRVRRDTHDEAVPEEVFQLLDATLPRCPNVKYVVLEQLGNGLKTEPSKIQFRQDFLRMQALVAKHRDHTSRPLTNTFLPLHPLPPSSIIEDEELYKQQLQLSHILETASSSDEAQRLLHASTLAKTAWKIEQWEPHMLQTAVDIAQKWKK |
| MBW4573846.1 | Aphanothece sp. CMT-3BRIN-NPC111 | MFKSKRRTPMVGLSLMMESDFFRAAQPLFESGEVEVLEWSFDMSWGIAAKPPWVEELLEFYSQRDRLIGHGVSYSLLSAQQDERQTHWLKLLQQECLDYRYRHISEHFGWMAAGDFYQSAPLPMPLTPETLQLGRNNLKRFSDTANVPVGLENLAFAFGWQDVLQQGEFIDRLLEPVDGFLVLDLHNLYCQLHNFQQSAEQLLKRYPLHRVRELHVSGGSWSVTSKGQQTKKIRRDTHDDQVPEAVFELLAIALHSCPNVEAVIFERIGHTLHTEAEIVGFRQDFLRMKQVVKDSASCLMS |
| MBX2874589.1 | Saprospiraceae bacterium | MPEILSSIACNLDRNILTAALPLFQGAEVQAMEWAFDSLYQQPEIPLWFESLLHTFADAGRLVGHGVYFSIFAGKWSAKQEQWLFELEELARKYPFDHVTEHFGFMTGADFHTGAPLSVPYTPTTLAIGQDRLRRIQQAAQCPVGLENLAFAYTPEEVWRQGDFLDRLLTPVNGFLILDLHNLYCQLYNFQLTFENLIQAYPLQRVREIHISGGSWEATKNAPDQPIRRDTHDDRVPEAVFDLLSKALPILPNLKYVVLEQLGLGLDTAAQQEQFRSDFNRMRQIVASFNQTNTELVQHHFKAPKSLPASRPAQSLDLQKDQEVLSYILEESQSLAEAQSALLESRLAKSAWQVEHWSPKMLETAVQIAQKWKKGFR |
| MBW3130172.1 | Hymenobacter profundi | MPSILSAIACNLDDHILAAALPLLEAGQVEAVEWSFDTLFDGRQLPNWFAELLTAYGEQGRLIGHGVYYSLLSGRWTPEQAQWLEQLRLLATRFHFDHITEHFGSFTGQNFHTGAPLPVPYTAAILRLGQDRLLRIADACQCPVGLENLAFAYSLEEVQRHGEFLEKLLTPVNGFLILDLHNLYCQLHNFDLSTDSLLALYPLHRVREIHISGGSWEDSRVEQGRQIRRDTHDEAVPDEVFALLERIIPQCPQLKYVVLEQLGTGLRTEASQHQFRQDFQRMAAIVEASRHRAPSHQPNAFQPPAALPTGPAAHDEALHAQQRQLSQILETATSYEEAQQRLRASSLAHTDWQLEQWAPHMLETAISIAQKWK |
| MBX3213937.1 | Labilithrix sp. | MMSPWVGLSLMTEDDFRVAAAPLFAGGEVDMLEWSFDAGRRPAPAWAEALLDHYAGEGRLFGHGVHYSPFSARWEARQQSWLDALGRELSRRPYARVSEHYGFMTAGPYVRGAPLPVPRDRASLQIGRERLDRLRAVLASAGSCPLGLENLALAWSRDEALAHGGFLDDVLTGHDDFIVLDVHNLYCQAENFGLSPDELLATFPAARVGELHVSGGSVLRAWPGRRDEVRCDTHDDAVPARVFDLLERALARFPNVRAVVFERLGGTIRTVEDAEALRADFRQTKALAALARRPADARAGGPDAGASAGGGGAAPRVLAEEGDVSALAAFQSALLALLERENDESVIRAHLEIAPEIAAYRGVVRAFDGRALGIAARVVKRWARRRPAGEAAE |
| MBX3076151.1 | Candidatus Obscuribacterale s bacterium | MPSEDFQQATLELFEQDRVGVVEWSFDFALNGAEIDVWSGQILERFSDKNALLGHGVNLSPLSARFSERQEKWLAHARSEFELRKYVHASEHFGFSEAGPIAHGAPLAVPMTPESLRRGQEMLKRYADAVQCPVGLENLAFAFCMEDVKRQGEFIDQLISSVDGFLLLDAHNIYCQVENFGVSERELLVSYPLSKVREIHLSGGSWSFSETGARLAVRRDTHDDVVPQEVFNIATLALKLCPNIQYVILERLAYTMMEPEQQQGFREDFETIEEILELCYV |
| MBX2929408.1 | Saprospiraceae bacterium | MRKIHPGIACNLDAHILSAAVPLLEQAQVGAIEWSFDALFRLRDVPEWFQGLLQAFAEAGLLTGHGIYFSIFSAQWSKEQEAWLSHLRALSAEIKFDHISEHFGFMTGADFHNGAPMSVPLTNATLAIGADRLMRIQDACNCAVGIENLAFAFCEEEVKQHGDFLSRLLIPVNGFLILDLHNLYCQMKNFDVDYEALIGHYPLDRVKEIHISGGSWAPASGAPGGQVRRDTHDDAVPAEVFELLAHTIPRCPNLKFVLLEQIGSGLQTDLQKVRFRQDYLRMCAICESVEIERDAAAGGNNFLPAGEFHLPAPVEDAMLYEQQAALSQILENASGGADAARRLATSILAGTPWQTEHWPEYMLETAVQIAQKWKKGFVV |
| MBX3219529.1 | Labilithrix sp. | MTPWVGLSLMFEDDFRVAAAPLFAAGDVDVLEWSFDTGPRLVGAAGDRGPSWAEALLDHYAAEGRLFGHGVHYSPFSARWEERQQRWLEALGRELARRRYVHVSEHYGFMTAAPFVRGAPLPVPRDRASLDVGRERLDRMRAVLASAGGCRLGLENLALAWSRDEALAHGAFLDDVLTCDDDFIVLDVHNLYCQAENFGLSPDELFATFPAARVGELHVSGGSVLSAWPGGGGEVRCDTHDHAVPAPVFDLLDRALGRFPNVRAVVFERLGGTIRSAEQAEELRRDFRQVKAIATRPRRFGEEHASTDALRDSTSRPAAPAISAPPRLPSSSRGDVEALAVLQSALLDLLDRESDESIIRAHLEVAPAFAEYRDEVRAFDGRALGIAARIVKKWGRKATTHAS |
| MBX3262053.1 | Labilithrix sp. | MSKLLAVMTPWVGLSLMTEDDFRVAASPLFAGGEVDVLEWSFDAGRRPVPAWAEALLDHYAGEGRLFGHGVHYSPFSARWEPRQQSWLEALGRELSRRRYARVSEHYGFMTAAPYVRGAPLPVPRDRASLQVGRERLDRMRAVLSTGGTCPLGLENLALAWSRDEALAHGAFLDDVLTHDDDFIVLDVHNLYCQTENFGLSPDELLATFPAARVGELHVSGGSVLPAWPGGRDEVRCDTHDHAVPACVFDLLERALARFPNVRAVVFERLGGTIRTVEDAEALRSDFRRTRELAARARGVADAGAAPSLPDARRASAPERGGVVALAALPSGAASSARGDDDALAEFQSALLALFERESDEFVIRAHLEVAPEFAPYRAAVRAFDGRALGVAARVVKKWTRRAPATHDP |
| MBX7082900.1 | Nannocystaceae bacterium | MTVPIGLSLMLEDAWREAALPLFERGDVDALEWSFDQAWRRPLPSWAAALLDHYAAADRLWGHGVTMSPGSVDAGDRHEAWLQRLQHELPQRRLRGVSEHFGFMVAGAIDLGAPLPLPDGPGARAVLRRNLARLAAVAGVPVGIENLALALCRDDAWQQGPMLADVLAHVDGYLVLDLHNLWCQLVNFELDPLATLQRYPLQRARCIHVSGGSWWGEARFRRDTHDDEIPEPVLALLQQAVPLCPALEVVVIERLGETLVGHPDALQRDLERVRARLGAA |
| MBX9669130.1 | Candidatus Obscuribacterale s bacterium | MKQQVLAGLSLMPDDDFADSTYPLFEDGLVDWVEWTFDMGWSDRGVPGWLRELLDYFGERGRLSGHGVQYSALSARWTDFHDRWVARLDQEVRERKYAHISEHFGLARAGQYLFHAPLPVPYCDAAISVGRDNIAKIAAIARCPIGLENLALAFCPLDVAHQGEFLDKLLAPVVDGFLVLDLHNLYCQSVNFQVPFFELLKSYPLERVTEIHISGGSFSHSSAGNTTQPIRRDTHDGPVPQELFDFLPRVIESCPRVQVVILERLGNTLSNKSERSGFASEFRKLRKTLYRLQTKSTNIGSQSGFSHTTLQTLSRDRDLNHEEKTLLVYQDNLLNKLSEDRDATWVKMQLVDLHNNHSWLLGYIDELEVRMLEVGQELTKKWGVRV |
| MBX9696430.1 | Cyanobacteriota bacterium | MQQNKPRLGLSLMPTDDFHQASAELFERNLVKVVEWSFDLAWSGAIIEPWCQNIIDRFSVSNALIGHGVNLSVLSARFSKRQQEWLARTKDEFQRRTYIHATEHFGFSEAGPIAHGAPLAVPMNSESLKIGKEMMRRYADATQCPVGLENLAFAFGIEDVKRQGDFIDQLISEVDGFLLLDLHNIFCQVENFGFSAQELLDSYPLSKVREIHLSGGSWSESVSGKRAAVRRDSHDDGVLQDVFNFAALAMKLCPNIEFVILERLGNTMMDGPESQREFRDDFEAMEEILHYCYA |
| MBX9772085.1 | Candidatus Obscuribacterale s bacterium | MAPSLKTGFSLMPTDDFYSVLLNLLDSGKVDVVEWSFDMSWDTTTPDWCESILAEFGKRESLLGHGVRFSPLSGKRSARQERWLQLLEKDCRHHKYLHVSEHFGFFEAGSYTDGAPLPLPMNDGSLGAGRSSLQALRDVCKTQVGLENLAMAFSVNDVRQQGDFLEALLEPLDAFLILDLHNLFCQSVNFDIPAAELMNSYPLHRVKEIHISGGSWSSSNTGQRKHVRRDTHDGPVPDEVFGLVSVARAKCPQLEFAILERLGNTMDDPADQSQYMDDFHRLWSLMNE |
| MBX9567968.1 | Candidatus Obscuribacterale s bacterium | MVTPSKKEIFCGLALMDSPDFLAASLPLFDAGEVEVLEWSFDIGWENGQDEWASELVAFYSSENRLLGHGVNFSVLSGKWTAGQQAWIERFREECAQHSFMHVSEHFGFSRAGNVIQTSPLPVPYSKSALALGVERLKMMSSIAKVPIGLENLGFAFSIDCVKEQADFINQLLDKSDGFLLLDLHNAFCQIANFGVDAEEYFSLYPLQRVREIHISGGSWSADRDGNSIRRDTHDHEVPEEVFDLLMQVLWRCPNLKTVILERLGNTIRSEEEAARFRSDFLHLKKLIQVFCSESYTEVIPPVSTLRTNGPKKAAKG |
| MBX9947652.1 | Candidatus Obscuribacterale s bacterium | MPTKPKLGLSLMPTEDFEQATVSLFQNNQVHAIEWSFDFAWNGAVISEWCEGILDEFSDKGALTGHGVNLSPLSARFSKRQEAWLEFAKEEFRSRKYVHASEHFGFSEAGPIAQGAPLAVPMNSESLRIGKEMLKRYADATQCPVGLENLAFAFGLEDVKRQGDFIDQLISSVDGFLLLDVHNIYCQLANFGVSELELLNSYPLSKVREIHLSGGSWSSSISGKRAAVRRDTHDDAVPQEVFNLATLALKLCPDVEFVILERLGYTMMDSKSQEEFREDFETMEEILEYCYA |

| | | |
|---|---|---|
| WP_223401109.1 | Dyadobacter fermentans | MAEIYSSIACNLDTHILQAALPLFEQEKVEAIEWSFDTLYKFGEIPAWFTDLVSEFSSHRRLIGHGVYFSLFSGKWSPGQQEWLARLKKLSGEFNFDHITEHFGFMTGEDFHKGAPIGIPLTPRTLALGKDRLQRIQDACQCPVGLENLAFSYSIEEVRKHGTFLDQLVESVNGFIILDLHNIYCQAHNFDIDFADILGLYPLNKVREIHISGGSWDDTLTPTRKPVRRDTHDESVPEAVFDFLAEAIPHCPNLKFVVLEQMGTALETQASQAAFQSDFLRMDSIVKSLARKSLPIPEHLFAPLAGTAHPTPLEDPDLYKQQMQLSHILETAANVSDAHAMLRSSDLGNSAWEVENWAPYMLETAMAIAQKWKNGFV |
| WP_222165112.1 | Edaphocola aurantiacus | MSRTILPTIACNLDADILSASLPLLEEGVVGALEWSFDALFRQDHVIPEWFYDLLTAYSNEGRLIGHGVFFSLFSGRLLPEQAAWLQRLKQLTRQYRFDHITEHFGFMTGSDFHKGAPMSVPYTSGTVAIGRDRLQRIQDAAQCPVGLENLAFAYSLDEVKRQGAFLEQLIEPVNGFIILDLHNVYCQLHNFNIPFEVLIQAYPLDRVREIHISGGSWETAVTDQSRSIRRDTHDGRVPESVFELLERTLPLCPQVKYVVMEQLGAGLQTEAAKIGFRNDFQQMQKITEQFLEHHPRMAAVPNNFLPEQSLPPGQIIEDEYLYGEQQILSYILEHAAGHAEAMEQLQHTALRNSVWDIEKWQPDMLETAIRIARKWK |
| WP_223559707.1 | Chryseobacterium lathyri | MKKPLLGLSMMPEADFVSAILPILQNNSVDVLEWSFDTFYDAQEPEWLSGLLDFYSANNRLLGHGVYYSLFDAKWTERQEGWLEKLKKEFRHRKYNHITEHFGFMNTENFHQGVPLPVPLHPKTLQIGKDRLKRLQDAVEIPVGVENLAFSFSIDDVKEQGDFLDKLIEDIDGFLILDLHNIYCQSWNFEISMQEIIRLYPLEKVKEIHLSGGSWQESAYGKKPVRRDTHDESIPEEILNILPEVLSQCHNLEYIIIERLGHTLNTDSEKDAFFDDFNRVKKIIQASDSLPGNKRVWHKKNEEPSVPLEDLLLYDEQTELTKMLFDGTQIHSVKKHDFHYFKPENWDEEMITTAQQIIKKWNPY |
| WP_223598940.1 | Chryseobacterium sp. GVT01B | MKKTVLGISMMAEADFVSAVLPLLQNNAVDVLEWSFDTLYYINEPDWLRDLLNFYAENNRLIGHGVYYSLFDARWTERQEEWLQKLKEEVSLRKYNHITEHFGFMNTENFHQGVPLPVSLHPKTLQIGKDRLYRLQQAVDIPVGIENLAFSFSIDDVKEQGVFLDKLTEDTNGFLILDLHNIYCQSCNFEVEIEEIIDLYPLDKVKEIHLSGGSWQESVYGKKQVRRDTHDDVIPEEIFFILPSVLAKCQNLEYIIIERLGHTLKTEEDKENFLNDFNKVKTIIEASDWKEKEKRSWNKRQIQLTEKPLEDLALFEEQSRLTRLLFDGAGAGIIKDQDFHYFKTKNWDPEMILTAQNIIKKWNPY |
| MCA0235173.1 | Bacteroidota bacterium | MLYPSIACNLDTDILLAALPLLESERVQALEWSFDTLYQVRDIPAWFDELLLEYGRAGRLIGHGVFFSLFSGRWSPEQAQWLEHLKKQCSVFQFDHITEHFGFMTGADFHSGAPISLPFTASTLAIGHDRLRRIQEVCACPVGLENLAFAYSLEEVKRHGDFLNRLVEPVNGFIILDLHNLYCQAHNFDMPAEELLDLYPLERVREIHISGGSWENSPLLPERPVRRDTHDDAVPAEVFQLLRNTIPLCPNLKFVVLEQMGTALHTEANRRQFRQDFDAMHTILEAFDAPAQSTGNTFAPFSLPALGTPLEDADLYRQQRQLSEILETADSYEAARQMLENSDLAGTAWGTEHWAPYMLETAMRIAQKWQKGFQQPA |
| WP_223608183.1 | Chryseobacterium sp. OSA05B | MKKPLLGLSMMPEADFVSAILPVLQSHSVEVLEWSFDAFYDVAEPEWLSGLLDFYAGNNRLLGHGVYYSLFDAKWTDRQEIWLKKLKEEVSRRKYRHITEHFGFMNTENFHQGVPLPVPLLPKTLEIGKDRLSRLQDAIEIPVGVENLAFSFSIDDVKEQGEFLDKLVEDIDGFLILDLHNIYCQSCNFEVDMMEMIRLYPLGKVKEIHLSGGSWQESAYGKKPVRRDTHDDRIPEEILKILPEVLIQCHPEYVIIERLGHTLNTEVDKEFFFDDFNRVKEIIEFSDEPVGDEKIWNRKNRTYSKPVEDLLLHDEQTELTRLLFEGNGTEAIKNSDFHYFKPEDWDEEMIVTAQQIIKKWNPY |
| MCA9105848.1 | Planctomycetales bacterium | MSRSSAPWVGLSLMLGDAFHDAVRPLFDGDEVDVVEWTFDMGWGRPLPAWLEATLDEFSSRGRLLGHGVSYSVLDASLTDRQPRWLHHLQVELQRRRYHHLSEHFGLMGGGNFHLSSPLPIPLSDTAVEVGNSRLRDLARVARLPVGLENLALAFSLSDVQQQGEFLERLLAPVDGFLLLDLHNLYCQAMNFNVSLEELVDSYPLDRVRELHLSGGSWSQPLSHARPVRRDTHDDRVPEELFQILPTVLRRCPRVEAVILERMDGTIDSANLSDAANFRADYRRLRAITREAG |
| MCA9805219.1 | Cyanobacteria bacterium HKST-UBA02 | MPTEDFRQASAELFEQQKVQAVEWSFDFGFDGAQIEPWCLDILDRFSEGGSLIGHGVTLSMLSASFSERQEAWLASAREEFVNRSYVHASEHFGFSEAGPFADGAPLAVPMTSQSLRQGKEMIKRYADAVQCPVGLENLAFAFGMEDVKCQGDFIDQLISEVDGFLLLDLHNIYCQIENFGVEADTLLDSYPLSKVREIHISGGSWSHSLSGRRPAVRRDTHDDSVPQEVFNLAALALKLCPSVEFVILERLAFTMTESKSQRDFRDDFYTMQEILVVRA |
| WP_225718062.1 | Chryseobacterium rhizoplanae | MGKPLLGLSMMAEEEFLSAVLPLLQGNSVEVLEWSFDTFYNSQEPNWLSDLLNFYAENDRLVGHGVYYSLFDARWTERQEIWLKKLKEEFRKRKYNHITEHFGFMNTENFHQGVPLPVSLHPEIVEIGKDRLYRLQDAVQVPVGIENLAFSFSLNDVMEQGIFLDRLTEDIEGFLILDLHNIYCQSCNFEVEMSEIIKLYPLDKVKEIHLSGGSWQESIYGKKLIRRDTHDDEIPEAILSVLPSVLLQCKNLEYVIIERLGNTINTREKKKRFLDDFLKVKALIDSSGLEIQNKNSWTKKELHFPESPLENIILHEEQTLLTKLLFDNVEAECIKNHKFHYFKTENWSPEMILTAQNIIKKWNPY |
| MCB0707287.1 | Saprospiraceae bacterium | MSKIHSSIACNLDANMLLASLPLFASEKVQAIEWSFDTLYKHRHIPDWFVELLQAYSAENRLIGHGVYFSLFSGKWSPDQQAWLDHLSRLSTNFNFDHITEHFGFMTGEDFHRGAPLSLPFTTKTLLLGQDRLKRMQEACGCPVGLENLAFAYSLEEVEQHGAFLEALVEPVNGFIILDLHNLYCQLHNFDVSFDDIIKRYPLDRVREIHISGGSWEPSRQKPEKSIRRDTHDDAVPEAVFELLEQSLMLCPNLKYVVLEQLGIGLASEDSKQQFQADFYRMERLVEAYNLSLAPQSAISFLPVQAFESSSRPAVEDMQLYQEQQVLSRLLEDSTDLESFQAGLNNSILAHSDWRIETWAPEMLETAYQIAQKWKKGFA |
| MCB0644088.1 | Phaeodactylibacter sp. | HASIACNLDTHLLLTALPLFEAEQVQAIEWSFDTLFKIPEIPDWFLELLDVFGEADRLVGHGVFFSLFSGQWSPEQADWLAKLEALSQRFRFNHITEHFGFMTGADFHKGAPIPIPYTPTTLNIGRDRLQRIQQACQCPVGLENLAFAYCLDEVKQHGEFLAALVEPINGFIILDLHNAYCQAHNFELGLEEILAQYPLHLVREIHILGGSWQDSSILPERQIRRDTHDDAVPEAVFELLELALPRCPNLRFVVLEQIGTALYDPRQQAQFRADFLRMKTLVEHHNQERNTGPENDFQNPHPIQLAPTPAQNADLARQQQSLSDILETAENLAAAQSALAASPLAQSSWQVERWHPAMLETVLQIAQKWKEGWK |
| MCB1136801.1 | Chlamydiia bacterium | MRSQLEKTRPLAGLSLMTEADFLRAAFPLLASGEVEVLEWSFDTGWGANAPAPALREVIAAFSAKSRLIGHGVSFSLLSGEWTERHQTWVDALREEVEDLSYRHVSEHFGFLTAGNMHQGSPFPVPLTPSILELGQTRMCLLAETAKVPVGLENLSFAFTASDVARQGEFLDLLLEPVNGFLVLDLHNLYCQAVNFECELESLMLRYPLERVREMHISGGSWSETPSSTRPVRRDTHDEAVPEAIFEFLPTALEHCPGCEAVIFERIGNTLQHPTDVLQFQQDFRRLKHILESLYANQPSYARTVPSHTVGSSC |
| MCA9816168.1 | Cyanobacteria bacterium HKST-UBA01 | MTVKPKLGLSLMSTPDFHQASAELFESGKVEVVEWSFDNAFNGAFIEPWCQDIVDRYSDANALLGHGVTLSPLSVRFSQRQRDWLAGAAEEFRSHNYIHASEHFGFSEAGPVAHGAPLAVPMNSATLKSGKEMMKRFAEATRRPVGLENLAFAFCIEDVKRQGEFIDHLLSDVDGFLLLDLHNLYCQQLNFNIPAIELLESYPLERVREIHISGGSFSPSISGRRIFVRRDTHDEPVPQEVLELAVHALKRCAGAEFVIYERLGFTMEDEESQAEYREDFETLAQLLEMIYV |
| MCB9279281.1 | Lewinellaceae bacterium | MIHSAISCNLDADILEAALPLLEASEVEALEWSFDTLYKSGGMPPWFEALLDAFGREGRLVGHGVYFSLFSGRWSEDQAKWLAELGRLSDQFQFNHITEHFGFMTGENFHAGAPVSIPFTKATLAIGQDRLKRIQQVCRRPVGLENLAFAYSIEAVKVHGEFLEQLVAPVNGFIILDLHNLYCQLHNFGVDFAEMIRLYPLDRVREIHISGGSWEDATAQPGRRIRRDTHDDSVPDEVFRLLENTIPMCPNLKFAVLEQLGAGLKTEASKVAFRGDFRKMAAIIRRHNPTDNLSGINEFASKIEMPEGPPLEDEVLYGQQRELSYILENAADATTAENLLCKSSLAKTDWRIEDWDPAMLETAVRIARKWRRGFV |
| MCB9468714.1 | Candidatus Obscuribacterales bacterium | MTVKPKLGLSLMSTPDFHQASAELFESGKVEVVEWSFDNAFNGAFIEPWCQDIVDRYSDANALLGHGVTLSPLSVRFSQRQRDWLMGAAEEFRSHNYIHASEHFGFSEAGPVAHGAPLAVPMNSATLKSGKEMMKRFAEATRRPVGLENLAFAFCIEDVKRQGEFIDHLLSDVDGFLLLDLHNLYCQQLNFNIPAIELLESYPLERVREIHISGGSFSPSISGRRIFVRRDTHDEPVPQEVLELAVHALKRCPGAEYIIYERLGFTMEDEESQAEYRGDFETLAQLLEMIYV |
| WP_227769778.1 | Hymenobacter sp. 15J16-1T3B | MAESPSILATVACNLDADILSAAFPLLEEGRVEALEWAFDALYWTEQPPAWFADLLDSYSQAGRLIGHGVYFSLLSGRWTAEQQQWLRQLAQVAARFRFDHVTEHFGFFTGQNFHHGAPLPVPYSPAALRLGQDRLQRLQAACQCPVGLENLAFAYSLDEVQRHGEFLDQLLAPVNGFVILDLHNLYCQLHNFAVPYEALIELYPLERVREIHISGGSWEASSLVPGRTVRRDTHDEAVPDEVFALLARTLPRCPNLKYVVLEQLGTGLKTAASREQFRRDFRRMAALLEPQRTGRPAAPDSPFAPLPLAPAGPAAEDETLHEQQRQLSQILETAPSYDAARQQLLQSGLAGTDWRIEHWEPHMLETAVNIARKWKK |
| WP_227941441.1 | Chryseobacterium sp. X308 | MRKPMLGVSMMAEADFVSAVLPLLQNNSIEVMEWSFDTLYHTNEPDWLRDLLNFYAENNRLIGHGVYYSLFDARWTARQEEWLKKLKEEVSLRKYNHITEHFGFMNTENFHQGVPLPVSLHSKILQIGKDRLYRLQDAVNIPVGIENLAFSFSIEDVKEQGVFLDKLTEDTNGFLILDLHNIYCQSCNFEVGLQEIINLYPLHKIKEIHLSGGSWQESAYGKKQVRRDTHDDVIPEEILSVLPSVLTQCPNLEYIIIERLGHTLKTEEERNDFLADFNKVRTIIETSDLKFKEKNRWCKKEVKLSEKPLEDPLLYEEQSKLTRLLYDNSDVKAIQDLDFHYFKTQHWDAEMILTAQNIIKKWNPY |
| WP_226176240.1 | Hymenobacter lucidus | MSTQVPAAAVSEPGILSAIACNLDADILSASFPLLEQGRVEALEWSFDALFWAEQVPEWFTDLLRAFADQKRLVGHGVYFSLLSGRWTREQQQWLEQLRELAGRFAFDHITEHFGFFTGQNFHSGAPLPIPYTSAALRLGQDRLRRIQQAAGCPVGLENLAFAYSLDEVQRHGEFLDQLVEPVNGFLILDLHNLFCQLHNFAVPYEELIKLYPLERVREIHISGGSWEKSGQEPGRQVRRDTHDDSVPEEVFELLALTLPRCPNLKYVVLEQLGNALKTEASRTQFRADFGRMETLVGEFRRRAAPRASQLFLPPQPWLAGPVAEDAHLHEQQQQLSRILETAPSFEAARQQLHASSLAHSAWKIEEWAPHMLETALRIAQKWKQ |
| MCB9230954.1 | Bacteroidia bacterium | MIYSSLALNLDVPLLQAALPLLEAEKVEAIEWSFDTLFKVRDIPPWFVELLREFGQAGRLVGHGVFFSLFSGRWVPEQDHWLKRLREMSAEFQFNHVTEHFGYMTGVNFHQGAPMSVPLTPEIMALGQDRLQRIADAAQCPVGLENLAFANSLDEVRQQGEFLSRLLEPVNGFIILDLHNLWCQLHNFELDFADLIRLYPLDRVREIHISGGSWEDSGARPGKKIRRDTHDHAVPEEVFALLENTLPLVPNLKFVVLEQMGHSLADAAQQEIFRSDFLRMESLVQSSGQKEKAEIPGLFQPPRVSFPEIPLQNENLFLQQRELAHILETSTNLDQALAQLQASALAHSKWKIETWAPEMLETALHIAQKWKDGW |
| WP_228421676.1 | Chryseobacterium culicis | MLGVSMMAEADFVSAILPLLQNNSIEVMEWSFDTLYHAHEPDWLHDLLNFYAENDRLIGHGVYYSLFDARWTERQEEWLKKLTEEVSLRKYNHITEHFGFMNTENFHQGVPLPVSLHPQTLEIGQDRLYRLQEAVNIPVGIENLAFSFSIDDVKEQGVFLDKLTEGTNGFLILDLHNIYCQSSNFEVPIQEIINLYPLDKVKEIHLSGGSWQDSVYGDKQVRRDTHDDVIPQDILSVLPSVLTQCQNLEYIIIERLGDTVKTEEERDNFLNDFNTVSTIIETSDWKVKGKSGWSKKEMKLSEKPLENLVLYEEQSRLTKLLFDNVEVKAIKDQDFYYFKTENWDAEMILTAQNIIKKWNPY |
| MCC6723041.1 | Saprospiraceae bacterium | MLLPSIACNLDKNLLTTALPLFESGSVEGLEWSFDALFKQPNMPDWFQELLLTYANEGRLIGHGVYFSIFSGKWKPEQTDWLAQLRQFSKTYRFDHVTEHFGFMTGEDFHKGAPISIPYTSTTLGIGIDRLMRIQDACNCPVGLENLAFSYSLDEVKKHGEFLQKLIEPVNGFIILDLHNLFCQLHNFEVGFEEMLKLYPLQRVREIHISGGSWEDSMVVPNRKIRRDTHDEPVPASVFELLEKTIPHCPNLKFVVLEQLSSALNTGEERTAFQADFLKMNDIVAQHSGIEQLTTNAFLPLTTLYFNSQPVEDEALAEQQAILSQILETANSYEDGRNMLRNSSLAATAWNVENWDPAMLETAIAIARKWKHGF |
| MCC6283020.1 | Saprospiraceae bacterium | MKTLSSIACNLDADILLAALPLLEAEKVQAIEWSFDTLYKFREVPPWFTELLQEFGRAGRLIGHGVFFSLFSGKWSAAQQDWLNQLTQLSKTFRFDHITEHFGFMTGADFHAGAPISLPFTPVTLAIGTDRLSRIFDACQCPVGLENLAFSYSLDEVKKHGDFLFRLIEPINGFIILDLHNLYCQSHNFGLEPDALLQLYPLDRVREIHISGGSWEDSPLLPSRPIRRDTHDDSVPAAVFQMLEYALGQCPNVKFVVMEQLGAGLKTETSRRQFQADFETMDAILRQQESSTQPTNAFLPAGRIELGQPLEDELLHRQQRQLSDILEQASGCTDAQRLLSASELAHTAWDVEKWEPYMLETALRIAQKWKKGFS |
| CAH0184169.1 | Chryseobacterium sp. Bi04 | MMAEADFVSAILPLLQNNAVDVIEWSFDTFFEAEEPSWLKDLLDFYAGNDRLLGHGVYYSLFDARWTERQEMWLQKLKQEVQKRNYNHITEHFGFMNTENYHQGVPLPLSLHPKTLEIGKDRLYRLQDAVDIPIGVENLAFSFSVNDVREQGVFLDKLIDDIDGFLILDLHNIYCQSCNFEVSMQDIVGLYPLEKVKEIHLSGGSWQDSVYGKKMIRRDTHDDVIPEEIFAVLPSVLSQCHNLEYIIIERLGHTIHTEEEKQCFFNDFMRVKKILNDSGGMKRKNEKWVKRERLLSEPVENSILYEEQIRLTKLLFDHADPTLIKNQNFHYFKTKNWDLEMILTTQNIIKKWNPY |
| MCC6672286.1 | Planctomycetota bacterium | MDVRVGLAWMPEPEFTAAVLPLLAAGEVGALEWSFDTLPDGHDEDPALTGLLDAFAAQERLYLHGVTFNTFGGRLDARSRRWLERLRSACARRRVRHVSEHFGFLRAGGWRDGPPLPVPTDACALDLGRERLRLLRDAACCPVGIENLAFAFARDEARRQGAFLDDLLAPVDGFLLLDLHNLHCQAANFGMDPEELLASYPLHRVREIHVSGGSWREGSHGPVRRDTHDGTVPSAVLDLLAIALPHCPRLEVVILERLGASLPDRAAQLAFASEYRGLAARIAEARVH |
| MCC6978279.1 | Candidatus Melainabacteria bacterium | MFEEERVESLEWSFDFTWNGVVVDPWALELIDKYSEANALTGHGVNLSPLSARFSKRQEEWLARAREEFKTRKYVHASEHFGFSEAGPIKHGAPLSVPMDAASLKVGKEMLKRYADATGCPVGLENLAFAFSINDVKKQGHFIDELISSVNGFLLLDLHNIFCQIANFGISELELLNSYPLDKVREIHISGGSWSPSISGKRAAVRRDTHDDGVPQEVFNLAALALKLCPNIEFVIFERLGNTMFSLEAQSEFRDDFESMEQILEFCYA |
| MCC7453008.1 | Crocinitomix sp. | MKTSQSTIYSTVACNLDQHLLQATLPLFASEKIEAIEWSFDTLFKVKNIPDWFVELLKTYSQANRLIGHGVYFSIFSGKWSEDQTNWLRHLKKTANHFKFDHVTEHFGFMTGSDFHKGAPISVPYNQSTLALGRDRLVRIYDACQCPVGLENIAFAYTLDEVKQQGEFLTKLIEPINGFIILDLHNLYCQLQNFSLSFDTLIKAYPLDRVREIHISGGSWEPSDVHPTRNIRRDTHDNAVPDEVFELLKLSLNRCPNLKFVVLEQLGTALYTSEQQIQFQHDFDRMQNIIHQYLKKNTPQISNTFLPEIKSSLSLEIPQNEELAQQQAELSFILETAKNHSDAHEKLRQSSLASTDWKIENWSAYMLETALKIAQKWKQGW |
| WP_229047851.1 | Chryseobacterium arthrosphaerae | MKKPLLGLSMMPEADFVSAILPLLQSNSVEVLEWSFDIFYEAKEPEWLDELLNFYSENQRLIGHGVYYSLFDALWTERQENWLKKLKEEVRKRKYSHITEHFGFMNTENFHQGVPLPVPLHPKTLLIGKDRLYRLQDAVELPVGIENLAFSFSIEDVKEQGVFLDNLIADINGFLILDLHNIYCQACNFEVEMEDIIRLYPLDKVREIHLSGGSWQESVYARKMIRRDTHDNLIPEEIFSVLPSVITQCNNLEYVIIERLGHTIKTDQEKRAFLDDFTRVRRIIDDSEIKTGKKIRWGQKETGLSNSPVEDDALHEEQSRLTKLLFNSSGAEMIKEQRFHYFKTENWDPDMILTAQNIIKKWNPY |
| WP_229755264.1 | Hymenobacter cavernae | MMEEAPIPFAAIACNLDFDILAAAFPLLEAGQVEALEWSFDTLFRAGQVPDWFSELLSAYSQQGRLIGHGVFFSLLSGKWTPQQQQWLQHLQQLSAQFRFDHITEHFGFFTGENFHYGAPLPIPYTQASLRIGQDRLSRIYEACQCPVGLENLAFAYTPDEVKRHGEFLNKLLEPVNGFVILDLHNLYCQLHNFAVSYEELIELYPLERVREIHISGGSWEESEQAPQGKVRRDTHDEAVPEAVFELLARTLPRCPNLKYVVLEQLGNGLKTESSRTQFRHDFLRMEALVQKHRASLLGHPEHLFRPPQPLQLGEVIEDQSLHEQQRQLSRILETAASVEEAQRRLHASVLVGTDWQMEQWEPHMLETAIHLAQKWKQ |
| MCC6953627.1 | Deltaproteobacteria bacterium | MVRCAIAAMPDERWLGAVMPLLEQGEVEALEWSFDCDWNRGTVPEWTHQLIRHYSEGARLFGHGISLSLLSTELAHHEQWFEKFERVCRDLSFVHVSEHFGFTRTGNAHHEAPLPCPYDPIIATLGASRLQRMKSICGSPVGVENLALALSPDDVRMHSRFIEEIVTPADGFVLLDLHNLYCQSVNFGLGMMELVQSYPLERVCELHLSGGSWNEVNVYGVSSSIRRDTHDAAVPDSLWEVLPEVIRKTPNLRVVTLEQLPDGLAHAPAAEAYREDYRKLRRIVEGGRDAGA |

| | | |
|---|---|---|
| MCC7537066.1 | Deltaproteobacteria bacterium | MDAALPVGVSLMLEEDFLGATLPLFEEGAVDVLEWSFDVGWGRELPAWAQGLLQHFAVSGRLLGHGVSFSPLSARWEEPQRNWLHRAALEVAKLKYRHVSEHFGFMAAGSFTRGSPMPMPVDAAVIRTGRDALSRLAEVTGLPVGLENLALALSSRDVDGQGDHLDALLEPVGGFVVLDLHNLWCQAVNFGRDPVGLLGRYPLGRVRELHVSGGSWSQPSSAEEPFRRDTHDAAIPDEVHALLPEALASCPHVEAVI |
| WP_230035574.1 | Chryseobacterium sp. Bi04 | MEKPLLGLSMMAEADFVSAILPLLQNNAVDVIEWSFDTFFEAEEPSWLKDLLDFYAGNDRLLGHGVYYSLFDARWTERQEMWLQKLKQEVQKRNYNHITEHFGFMNTENYHQGVPLPLSLHPKTLEIGKDRLYRLQDAVDIPIGVENLAFSFSVNDVREQGVFLDKLIDDIDGFLILDLHNIYCQSCNFEVSMQDIVGLYPLEKVKEIHLSGGSWQDSVYGKKMIRRDTHDDVIPEEIFAVLPSVLSQCHNLEYIIIERLGHTIHTEEEKQCFFNDFMRVKKILNDSGGMKRKNEKWVKRERLLSEPVENSILYEEQIRLTKLLFDHADPTLIKNQNFHYFKTKNWDLEMILTTQNIIKKWNPY |
| GJM31625.1 | Saprospiraceae bacterium | MSKLHASIACNLDANILLAALPLFEAEKVQAIEWSFDALFKTREVPPWFTALLSTYSKNNRLVGHGVFFSLFSGKWSPAQQNWLDQLKRLSAVYQFDHITEHFGFMTGANFHQGAPISVPFNPVLLAIGQDRLKRIQEACQAPVGLENLAFAYSIDEVKRHGEFLDQLIAPINGFIILDLHNLYCQIHNFEIDFEDIIQLYPLEQVREIHISGGSWEDSPLIPGKKIRRDTHDDAVPQAVFDLLERTIPRCPNLKYIVLEQIGTGLKTEASKRQFQQDFFKMEQIVQNKQRSTTKDQHTFLPSQPFHLGKPLEDVILHAQQLELSDILETATDYQHAQQCLQASSLANSDWKIETWAPAMVETVLRIAQKWK |
| WP_232831792.1 | Taibaiella helva | MDKIRSTIACNLDAGILSASLPLLEEARVEAIEWSFDTLFKLETVPQWFEALLEAFSKEGHLIGHGVFFSLFSGRWLPEQQAWLDHLRRLCRQYHFDHITEHFGFMTGADFHHGAPLSIPFTAETLAIGRDRLARIYEACGCPVGLENLAFSYSLDEVKRHGAFLEALLEPLNGFIILDLHNLYCQLHNFSLDFDTLIRLYPLHRVREIHISGGSWEPSDLFGGQPVRRDTHDDAVPETVFTLLEKTIPMCPLLKYVVMEQLGAGLETEASRQAFRNDFLRMDGIVSAAASDPQAFRPEAFLPQVPVIPGPVTESMALHTQQLELSAILEASAGYEEAYHRLQHSSLAATGWQVETWQPHMLETAIHIAQKWRKKST |
| MCE3228656.1 | Bacteroidota bacterium | MKPLVGLSLMHEQEFLNAAIPLFLKNEVDVLEWSFDTILVEKYKPEWVHQLLKEYSINNRLIGHGVRYSLFDAKWTKRQESWIRKLKAEVKRYNYNHITEHFGFMSHSDFHKGAPLPVPFNKKSLAIGTDRLKRLQHAAELPVGVENLAFAFSKHEINKQGEFLDSLVESVNGFLILDLHNLYCQSQNFKTAIIELVKSYPLKKVKEIHISGGSWQNSVYTKDIKKVRRDTHDEKVPAEIFKVLPEVMKLCPNVEYVIFERLGDTLHNEKEAEDFRKDFRRLRKIIAGINSSKKTQTQNQKPSAAKGLPTARPITNNQKLNPPLFDQPLYNEQQFILDVLRKSQRPSEAFSLLKQKQLPDWNIASWNTSMLETAIGLIRKWDL |
| MCD6068446.1 | Bacteroidota bacterium | MKPLVGLALMHEQEFLRAALSLFAADEVELLEWSFDTILYEKYKPEWLHQLLREYSSKGRLIGHGVRYSLLDAKWTRRQSGWLNKLRKEVNRYPYQHITEHFGFMTSSDFHKGAPLPVPFNKHSLRIGIDRIKRLKDAIGSLPLGIENLAFSFSAADVKEQGAFLAELIRPVDGFLILDLHNVYCQAENFKIDILEIIRSYPLDKVREIHISGGSWKNSIYSKEIKKVRRDTHDERVPEQLFKILPDVLSLCPNIAYVIFERLGNSLQEEEEQEQFRKDFRRIKSIVHRSSPDKKASTVSERSIPYSKAPLNDLSFYKEQRFLSKTILDAGDPVEALLRLKKEELKYWDVDQWNSSMLETAMSIAKKWDL |
| WP_232720674.1 | Chryseobacterium gleum | MRKPMLGVSMMAEADFVSAILPLLQNNSIDVLEWSFDTLYHPNEPDWLCDLLNFYAENGRLIGHGVYYSLFDARWTERQDEWLKKLKEEVRLRNYNHITEHFGFMNTENFHQGVPLPVSLHPKTLQIGKDRLYRLQEAVDIPVGIENLAFSFSVDDVKEQGVFLDQLTEDTNGFLILDLHNIYCQSCNFGLEMKEIVDLYPLDKVKEIHLSGGSWQESVYGKKQVRRDTHDDAIPQNIVSVLPSVMEQCQNLEYIIIERLGHTVKTEEEKVSFLNDFNTVRKIIEISDLKKAERSWMKKEMIFPEKPLEDWMLYEEQSELTKLLFENADAAVIKNHDFHYFKTENWDSEMILTAQNIIKKWNPY |
| WP_233332259.1 | Chryseobacterium gleum | MRKPMLGVSMMAEADFVSAILPLLQNKSIDVLEWSFDTLYHPNEPDWLRDLLNFYGENDRLIGHGVYYSLFDARWTERQDEWLKKLKEEVRVRNYNHITEHFGFMNTENFHQGVPLPVSLHPKTLQIGKDRLYRLQEAMDIPVGIENLAFSFSVDDVKEQGVFLDQLTEDTNGFLILDLHNIYCQSWNFGVEMKEIVDLYPLDKVKEIHLSGGSWQESAYGKKQVRRDTHDDVIPQDILSVLPSVMEQCQNLEYIIIERLGHTVKTEIEKVSFLNDFNTVRKIIEISDLKLAERSWMKKEMILPEKPLEDWMLYEEQSELTKLLFENADAAVIKNHDFHYFKTENWDSEMILTAQNIIKKWNPY |
| WP_233877771.1 | Dyadobacter sp. CY323 | MPRMRNGMIYSSIACNLDAHILMASLPLFEQEKVDAIEWSFDTLFKFNEIPSWFSDLIGEFSKNDRLVGHGVYFSLFSGKWLPEQQQWLDKLKKLSDQFHFDHISEHFGFLTGADFHKGAPIGIPLTTSTLAIGSDRLKRIQDACRCPVGLENLAFSYSLEEVKKHGDFLNQLVESVNGFIILDLHNLYCQSHNFSIPFNEIIRLYPLDRVREIHISGGSWDDLSNTHGKIVRRDTHDESVPEQVFDYLKRTVPKCANLKYVVLEQLGTALDSESRRVAFQNDFLRMDGIVKKESSAHINSITNNFLPAIGVIPDKVPPEDLPLYNQQTELSYILENARDLTDARMRLESSALSNSEWNVEQWEPAMLDTAIAIAQKWKNGF |
| WP_233635913.1 | Hymenobacter setariae | MGTPVAPAQPTVLAALACNLDADILGAALPLLEEGQVEALEWSFDALYWASEVPAWFVELLHAYSDQQRLLGHGVYFSLLSGRWTSEQQQWLNQLRELTQRYSFAHITEHFGAFTGQNFHAGAPLSVPYSPVSLRIGQDRLARIAEAARCPVGLENLAFAYSLEEVKRQGEFLGELLAPVNGFLILDLHNLYCQLHNFSIGYEELLECYALDRVREIHISGGSWENSQLLPTRKIRRDTHDEAVPVEVFNLLARTMPRCPNLKYVVLEQLGTSLKTEASHVQFRRDFVQMAALVQQHNHSPRSVPNAFLPRWPALPALAVQDDLLHEQQRQLSHILETAASLPEAQQRLRASALARSAWQIEHWEPYMLETALSIAQKWK |
| WP_233789821.1 | Dyadobacter sp. CY343 | MSRIFSAIVCNLDTHILQASVPLFEMEKVEAIEWSFDSLFKVDTIPEWFTDLVKEFSDHNRLIGHGVYFSLFSGKWSPEQQNWLSQLRKLSGEFHFDHISEHFGFMTGSDFHKGAPLGIPFNKSTLAIGRDRLARIQEACNCPVGLENPAFSYSYEEVKKHGDFLNQLVESINGFIILDLHNLYCQCHNFDIDFEQMLACYPLERVREIHISGGSWEMAIEERLIRRDTHDDAVPETVFAYLKKALPRCPNLKFVVMEQLGTALDSVEKQQAFQQDFFQMDKMVKSFEENSNSIQTKHFLPLTALTLQSQPVEDLLLAAQQMQLSSILETAVNLDHARKMLISSNLNNSDWHIEQWQPHMLKTAIDIAQKWKDGF |
| MCE9579119.1 | Deltaproteobacteria bacterium | MTRTPPRVGLNFLPEGDFRAAALPLFEAGLVDALERDIDYAWGSHPRGERRDEPAWALEILDAYAADDALYGHGVWMSVMSGRWTSRQERWIAQLTDECRARRYRHVSEHFGWLSAGPFWRNTMLPAPYTPATVALGVDRLRRLADAVGCPVGLENLGMGLGPVDAQEQGAFLGEILDAVDGFLLLDVHNVWTQAHNARLDPLALLRTYPLARVRELHVSGGDWYEPAPGARPIRLDSHDGDVPGDVWPLVAEALACCPALEVVFLERRGQTLGREPDRAAYRADFLALRAAVEAAFDVGEERCA |
| WP_233525516.1 | Chitinophaga silvisoli | MPEILSAVACNLDANILAACLPLMEESRVEAIEWSFDALYKVKQVPDWFRELLTAFSDENRLIGHGVFFSLFSGKWLPEQQAWLDHLKHTSTEFSFDHITEHFGFMTGKDFHHGAPLNIPYSAATLSIGRDRLKRIYNACGRPVGLENLAFSYSLDEVKRHGAFLDQLLAPVNGFIILDLHNLYCQLRNFDLAFDDLISLYPLDKVREIHISGGSWDDSAASPDRSIRRDTHDEAVPPEVFQLLEWTLPKCPHLKYVVLEQLGNGLVSQESKAAFYNDFLHMQEIVSQSNNKYSDNSFLPLLPTTTGPAVEDLALYQQQLELSAILETAASYDSAMHLLAQSSLANSDWRIEQWEPYMIETAVKIAGKWKK |
| WP_233823285.1 | Dyadobacter sp. CY312 | MTKIYTSVACNLDANILLASLPLFEQEKIEAIEWSFDALFKLEQIPPWFADLVSEFGKNNRLIGHGVYFSLFSGKWLPEQQQWLDSLRAFSREFHFDHISEHFGFMTGADFHKGAPISIPYTSSTLALGTDRLMRIQDAANCPVGLENLAFAYTLDEVRRQGAFLSELTNKVNGFIILDLHNLYCQIHNFNVPFEKLLHSYPLEKVREIHISGGSWEQNHAAPEDKIRRDTHDDEVPETVFELLSAAIPKCPNLKYVVLEQLGTALSDEQSRSRFQQDFLRMEQIVKNSKTASDGRIVNNFLPHTSLQSGHVPYEDLQLYREQNELSQILETATDLAHAQKLLASSVLNGTAWQTETWQPHMLETAIAIAQKWKNGFELK |
| WP_234615733.1 | Dyadobacter fanqingshengii | MSEIYSSIACNLDAHILQASLPLFEQEKVEAIEWSFDTLFKVSEIPAWFTDLVKEFSNHNRLIGHGVYFSLFSGKWSPGQQEWLDKLQKLASQFHFDHITEHFGFMTGEDFHKGAPISIPFTAQTLALGKDRLQRIQDACKCPVGLENLAFSYSLDEVKKHGDFLAQLVESVNGFVILDLHNIYCQVHNFNIDFDDIIKLYPLERVREMHISGGSWDETQVGPGRQVRRDTHDESVPEAVFEFLHNAIPKCPNLKYVVMEQMGTALDTLEKQAIFQSDFLRMDSIIKRLGFDYDPKIENTFTPLNTSASNIPLEDHELYKQQLQLSHILETAGSISEAQAMLRSSSLNNTAWEVDKWAPYMLQTAISIAQKWKNGFA |
| WP_235109853.1 | Acaryochloris sp. 'Moss Beach' | MTPQIGLSLMPQPDFWQAAQPLFDADEVDVVEWSFDMGWGGVVLPDLLCSVLRQFSQRNCLLGHGVSYSLLSAQLDRRHWLACLQAECRDYSYRHISEHFGWMATQDFARSAPLPMPLLPETLKLGCDRIQQFADIAQVPVGLENLAFAFGLQDVRQQGEFLDQLLEPVDGFLLLDLHNLYCQMHNFQLSATEIFSLYPLERVRELHLSGGSWSHHGTATIRRDTHDQAVPEPVFELLKLALQQCPQAEAIILERMGNTLNSEQEQLQFRQDFAHIRQITDHVSHVTEAGGK |
| WP_233850042.1 | Dyadobacter sp. CY326 | MPEIYASIACNLDAHILQASLPLFEEEKVEAIEWSFDALFNFNEIPSWFTELVREFSDHNRLIGHGVYFSLFSGKWSPEQQEWLHRLRKVSNEFHFDHITEHFGFMTGADFHKGAPIGIPFTTQTLALGKDRLRRIQDACRCPVGLENLAFSYSIEEVKKHGDFLNQLVESVNGFIILDLHNLYCQVHNFDIAFDEIVKLYPLDKVREIHISGGSWDKNPLRGNKKVRRDTHDASVPEAVFQYLRKAMRFCPNLKFVVMEQMGTALHTVERQQMFRNDFLRMSSIIESSNLDQQHPTNTFAPVSNFNINEPPLEDLFLHSQQVQLSQILENASNVDEARVMLQSSDLANTAWKVENWAPYMLETAISIAHKWKNGFS |
| WP_235149479.1 | Dyadobacter sp. CY345 | MPQVYSSIACNLDTNILSATLPLFEQEKVEAIEWSFDTLFKYEEIPEWFSDLISEFGNEGRLVGHGVYFSLFSGKWLPGQDAWLKKLNTFSRKFRFDHITEHFGFMTGSDFHKGAPIGIPYTKTFLDIGIDRLKRIQEACQCPVGLENLAFSYSLEEVKNHGEFLSKLVESVNGFIILDLHNLYCQIHNFDIDFQDILALYPLEKVREIHISGGSWDERKTSRGEKIRRDTHDDTVPKEVFDLLKATIPKCPNLKFVVLEQMGTALYSTESQLAFQQDFLKMNAIVKHKEYAFPEKLFNNFLPIDTVSLAPSPLENLTLFEQQLELSSILETAKNFDHARQLLSRSALHHSEWKVENWEPYMLETALAIAQKWQKGFL |
| WP_234656798.1 | Dyadobacter chenwenxiniae | MSGIYSSIACNLDAHILQAALPLFEQEKVEAIEWSFDTLFKFSEIPAWFIDLVSEFSRHDRLIGHGVYFSLFSGKWSSEQQEWLHKLQKLSREFQFDHITEHFGFMTGEDFHKGAPISIPFTSRTLALGKDRLQRIQDACKCPVGLENLAFSYSLDEVKKHGEFLDQLVESVNGFIILDLHNIYCQVHNFNIDFDYIIKLYPLDRVREMHISGGSWDETLADSGRQVRRDTHDERVPEAVFEFLGKAIPRCENLKYVVMEQMGTALDTLESQAAFQSDFFRMDSIIKEARTCFDPKIKNSFAPTNRSADIIPLEDHDLYKQQLQLSHILETAGSVSEAQALLRASSLNDTAWGVDSWAPYMLQTAISIARKWKNGFA |
| WP_234608367.1 | Dyadobacter chenwenxiniae | MSEIYSSIACNLDAHILQAALPLFEQEKVEAIEWSFDTLFKFSEIPAWFIDLVSEFSRHDRLIGHGVYFSLFSGKWSSEQQEWLHKLQKLSREFQFDHITEHFGFMTGEDFHKGAPISIPFTSRTLALGKDRLQRIQDACKCPVGLENLAFSYSLDEVKKHGEFLDQLVESVNGFIILDLHNIYCQVHNFNIDFDDIIKLYPLDRVREMHISGGSWDETLADSGRQVRRDTHDERVPEAVFEFLGKAIPKCVNLKYVVMEQMGTALDTLESQAAFQSDFFRMDSIIKETGTGFDPKIKNSFAPTNRSADIIPLEDHDLYKQQLQLSHILETAVSVSEAQALLRASSLNDTAWGVESWAPYMLQTAISIARKWKNGFA |
| WP_235139901.1 | Dyadobacter fanqingshengii | MPEIYSSIACNLDAHILQASLPLFEQEKVEAIEWSFDTLFKVSEIPAWFTDLVSEFSDHKRLIGHGVYFSLFSGKWSPGQQEWLDKLQKLASQFHFDHITEHFGFMTGEDFHKGAPISIPFTAQTLALGKDRLQRIQDACKCPVGLENLAFSYSLDEVKKHGDFLAQLVESVNGFIILDLHNIYCQVHNFNIDFDDIIKLYPLERVREMHISGGSWDETQVGPGRQVRRDTHDESVPEAVFEFFHNSIPKCPNLKYVVMEQMGTALDTLEKQAVFQSDFLRMDSIIKGLRFDFDPKIENTFTPLNTSASNIPLEDHELYKQQLELSHILETAGSISEAQAMLRSSSLNNTAWEVDKWAPYMLQTAISIAQKWKNGFA |
| WP_235938302.1 | Chitinophaga solisilvae | MPEIRSAVACNLDANILAACLPLMEASKIEAIEWSFDALYKVKDIPDWFVELLTAFSNENRLIGHGIFFSLFSGRWRQEQEQWLQHLRKTSTHFRFDHITEHFGFMTGKDFHHGAPLNIPYTKSTLNIGVDRLKRIHDACRCPVGLENLAFSYSMDEVKRHGAFLDELLEPVNGFIILDLHNLYCQLHNFDITFGEMIILYPLHKVREIHISGGSWEASPAAPERMVRRDTHDDAVPEEVFQLLQKTIPLCPNLQYVVLEQLGTGLDTDTSRTLFRQDFGRMQRIVHEQQPQTGSCFINSFMPDNLLISAVPAADETLFQQQLELSAILESASSCEAAVHALQHSSLAHSDWLIEQWQPYMIETAVSIAQKWKKI |
| WP_236858879.1 | Chryseobacterium sp. MEBOG06 | MTKPLLGLSMMPETDFVSAVLPLLESSSVDVLEWSFDTFYDAKEPAWLNELLNFYSENNRLIGHGVYYSLFDAIWTERQEIWLKKLKEEVHKRNYNHITEHFGFMNTENFHQGVPLPISLHPKTLQIGKDRLYRLQDTVNIPVGIENLAFSFCIDDVKEQGVFLDQLVEDTNGFLILDLHNIYCQACNFDVGMQEIISLYPLHKVKEIHLSGGSWQESAYGKKQIRRDTHDDLIPEEIVSILPSVLAQCPDLEYIIIERLGSTIKTDSQKKDFFNDFKKVEKITEASDHQICKKDNWLKKTVEFPEKPIEDLILYEEQTRLTKLLFDSKGSGIIKEESFHYFKTENWDPEMIRTASNIIKNWNPY |
| WP_235296475.1 | Portibacter marinus | MRNGVGKELYSGISCNLDSNILSACFPLFKGGEVEVIEWSFDTLFSHNNLPDWFDHLLTVFGEEGRLIGHGVYFSLFNGTFTKDQQEWLNGLRVTSDRYEFDHISEHFGFMTGQDFHKGAPLNVPLSDNSLAIARDRISRMSYACGKPVGLENLAFCYDKEGLIRQGSFINQILEEVNGFLILDLHNLYCQCMNFDIEFSELILKYPLDRVREIHISGGSWEIAQSEDRKIRRDTHDDAVPREVFEYLQEVIPKLPNLKFVILEQLGTGLDTIVKQARFRDDFETMKSIVDHKQLEKPLATNDFKSKNLKLSKHPISDETLAKQQQELTFILENSVDFRSVKKRLQNSSLASSPWNIESWEDSMLETAFAIANKWKSGF |
| WP_237144733.1 | Pontibacter pamirensis | MDRKELHPVGKEPEILSTIACNLDADILSAAFPLLEEGRVEAIEWSFDTLYWAAQVPYWFTELLLAYSQQNRLVGHGVFFSLLSGKWTKEQQHWLEQLKQLASQYSFDQITEHFGFFTGQNFHSGAPLHVPYTSATLRIGQDRLSRIYEACQCPVGLENLAFSYSLDEVKQHGEFLDKLVAPVNGFLILDLHNLYCQLHNFVISYEDLITLYPLERVREIHISGGSWEASVLEPEKKIRRDTHDDAVPEEVFQLLALTIPKCPNLRYVVLEQLGNGLKTESSRIRFRQDFLKMEALVQKSRSSLSRDVDNLFLPVQPLQKGPAVEDDMLHNQQLQLSHILETAPSYEEARRLLLTSSLAGSDWKIESWAPHMLETAVSIAQKWKKQS |
| WP_236850266.1 | Chryseobacterium sp. MEBOG07 | MRKPLLGVSMMAEADFVSAILPLLQNNSIEVLEWSFDTLYHTHEPDWLCDLLNFYAENNRLIGHGVYYSLLDARWTERQEEWLGKLKEEVLQRKYNHITEHFGFMNTDNFHQGVPLPVSLHPKTLQIGKDRLYRLQEAVDIPVGIENLAFSFSVDDVKEQGMFLDKLTEDSNGFLILDLHNIYCQSCNFGVEMQEIINLYPLDKVKEIHLSGGSWQESVYGKMPVRRDTHDDVIPEDIFSVLPTVLTQCQNLEYVIIERLGHTLKTEEEKSNFLHDFNKVRTIIEASDWKNRRENSWTKKEIQLTEKPAEDLVLYEEQSRLTRLLFDNVGAAVIKDQGFHYFNTENWDPEMILTAQNIIKKWNPY |
| WP_236387059.1 | Chitinophaga filiformis | MLAACLPLMEESRIEAIEWSFDALYKVKEVPAWFRELLTAFSNEKRLIGHGVFFSLFSGKWLPEQESWLKHLEQTTDEFHFDHITEHFGFMTGKDFHHGAPLNIPYTTATLNIGRDRLKRIHEACGRPVGLENLAFSYSLDEVKRHGAFLEQLLEPVNGFIILDLHNLYCQLRNFDISFDEMIALYPLDRVREIHISGGSWDDSAAVPNRRIRRDTHDDAVPPEVFQLLEMTMARCPHLKYVVLEQLGNGLETATSKQCFYNDFLQMEEIVRKNSDDNDADTPDPFLPLSPLSTDTAIEDMILYQQQLELSAILESSDSYAQAMDALRQSSLANSDWKIEQWEPYMIETAVKIARKWKK |
| WP_234645261.1 | Dyadobacter sp. CY356 | MSTIFSSVACNLDTNILLASLPLFEREQIEAIEWSFDTLFKNDEVPDWFSDLILEFSSQNRLIGHGVYFSLFSGKWMSEQQDWLKKLKILGERFQFDHITEHFGFMTGANFHKGAPISIPLTASTMSLGVDRLHRIQDACNCPVGLENLAFSFSLDEVKRQGNFLDKLVESINGFIILDLHNLYCQIHNFDVDYDYIIDLYPLERVREIHISGGSWDIQLTRTGKKVRRDTHDDAVPEEVFQILSAVIPRCTNLKFVVLEQLGTALDSEVKRTRFRSDFERMDAIVKQSGLALSNSPVNNFKPHHKITPHLIPLEDLSLYNQQLQLSTILETAVDQQHARHLLLNSDLRDSVWEVEKWDPCMLQTAIAIAQKWKDGF |

| | | |
|---|---|---|
| MCH2081132.1 | Saprospiraceae bacterium | MTKIHSTLACNLDKNILMAALPLLAAEKVEAIEWSFDALFKTRNIPNWFVDLLRTYSQGQALIGHGIFFSLFSGRWSPEQQQWLNHLKRITKEFNFEHITEHFGFMTGKNFHQGAPISVPYNERTLALGQDRLKRMYEACQRPVGLENLAIAYSTAEVQQHGEFLEALLSPINGFIILDLHNFYCHLHNFEENFDRLIQSYPVHMVREIHISGGSWQNSEWFDDKKVRRDTHDDGVPEIVFELLEKVIPLCPNLKYVVLEQLGTGLKTDGQKTRFQKDFLRMDEIVQPSIPHKKTQNDFLPKQKIKFDEIVEDEVFHQQQMQLSAILENAVDAASAEKLLMNSALAKTDWKVEEWKTHMLDAVIQIAQQWKKGFQSPAN |
| WP_238387972.1 | Hymenobacter sediminis | MQPEPAILATLACNLDADILSAALPLLAEGRVEALEWSFDALFWAEQVPDWFSELLTTYSAANRLLGHGVYFSLLSGRWTPEQQQWLEQVRQVVAQYPLAHLTEHFGFFTGQNFHSGAPLPVPYTPATLRLGQDRLSRIYEACRCPVGLENLAFAYSLDEVKRHGEFLGRLLEPVNGFLILDLHNLYCQIHNFSVSYEELIGLYPLERVREIHISGGSWENSELAPGRKIRRDTHDEGVPEEVFQLLELTIPRCPRLKYVVLEQLGPSLKTEESRIRFRHDFGRMEEVVHRHRAGRLGYADAFFLPPQPLQTGPALEDAQLHAHQQQLSHILETAPSFAEAQRQLHASALASSDWKLEQWAPYMLETAVSIAQKWK |
| MCG8329440.1 | Chitinophagales bacterium | MYASIACNLDTNILLAALPLFETEKVEAIEWSFDALFNKAQVPAWFYELLQAYAKEKRLIGHGIFFSLFSGRWREEQEAWLAQLQEVARDFQFQHITEHFGFMTGENFHAGAPLSIPITEKTLRIGQDRLKRIYQACECPVGLENLAFSYSLEDVKAQGIFLDQLLEPINGFIILDLHNLYCQLYNFEISFEELIALYPLQRVREIHISGGSWEDSVLTPQKRVRRDTHDEAVPDEVYKLLEKTIPLCPNLKFVTLEQIGTGLENEPKRERYRQDFLKMEKIVASYTKSTPVTNDFLPISPLQVSTSPEEDQILHKQQVELSYILENAPDYSSAQQQLEASSLAQSDWKVENWAPHMLETALKIAQKWA |
| WP_239808446.1 | Flavihumibacter fluvii | MQKIYSSLASNLDIQLLSAAYPLLAGGKVDAIEWSFDTLYKVDHIPMWFEDLLAAYSAQNRLIGHGVFFSLFSGRWSDAQQNWLAALRQTSQRYRFDHITEHFGFITGKDFHQGAPMSIPYTPATLAIGQDRLKRIQDACSCPVGLENLAFSYSLDEVKEHGEFLQRLIEPVNGFIILDLHNLYCQINNFDIGFEEIISLYPLDKVREIHISGGSWDEVSEPFPKNVRRDTHDSAVPEPVFDLLKMTIEKCPALKYVVLEQLGSALSTPESQAVFQQDFLILEEIVQEKNKGRWDFFNDAFLPSPLVIPDDIIEDEALYQQQKQLAEILEHCTTAAEAQQQMRQTTLANSAWDIEHWQPYMLETAIRIAQKWKDGFE |
| WP_240737107.1 | Hymenobacter metallicola | MAEQLQPVGASAGTPDILSAIACNLDADILSASFPLLEVGRVEALEWSFDTLFWAEQVPEWFTELLLTYSQQQRLVGHGVYFSLLSGRWTAEQQQWLQQLQELTKRFAFDHITEHFGFFTGQNFHSGAPLPIPYTRSALHLGQDRLQRIQQACGCPVGLENLAFAYSLDEVQRHGEFLDELVEPVNGFLILDVHNLFCQLHNFAVPYEELIRRYPLDRVREIHISGGSWDDSGPTPGRPVRRDTHDGAVPEEVFQLLEQTMPRCPNLRYVVLEQLGNALKTDAQHTQFRADFCRMEALVTRHRSRAAGRAGQLFLPLQPGAAGPVAEDESLYAQQQQLSRILETAPTFEEARRQLQASPLAHSAWKIEEWEPHMLETARRIAQKWQ |
| WP_241286337.1 | Chryseobacterium arthrosphaerae | MKKPLLGLSMMPEADFVSAILPLLQSNSVEVLEWSFDTFYDAKEPEWLDELLNFYSENQRLIGHGVYYSLFDALWTERQENWLKKLKEEVRKRKYNHITEHFGFMNTENFHQGVPLPVPLHPKTLLIGKDRLYRLQDAVELPVGIENLAFSFSIEDVKEQGVFLDNLIADINGFLILDLHNIYCQARNFEVEMEDIIRLYPLDKVREIHLSGGSWQESVYAGKMIRRDTHDDLIPEEIFSVLPSVIAQCNNLEYVIIERLGHTIKTDQEKRAFVDDFTRVRRIIDDSEIKTGKKIRWSQKETGFSNSPVEDDALYEEQSRLTKLLFDSSGAEMIKEQEFHYFKTENWDPDMILTAQNIIKKWNPY |
| WP_241315041.1 | Chryseobacterium arthrosphaerae | MKKPLLGISMMPEADFVSAILPLLQSNSVEVLEWSFDTFYEAKEPEWLDELLNFYSENQRLIGHGVYYSLFDALWTERQENWLKKLKEEVRKRKYSHITEHFGFMNTENFHQGVPLPVPLHPKTLLIGKDRLYRLQDAVELPVGIENLAFSFSIEDVKEQGVFLDNLIADINGFLILDLHNIYCQACNFEVEMEDIIRLYPLDKVREIHLSGGSWQESVYARKMIRRDTHDDLIPEEIFSVLPSVITQCNNLEYVIIERLGHTIKTDQEKRAFFDDFTRVRRIIDDSEIKTGKKIRWGQKETGFSNSPVEDDALHEEQSRLTKLLFDSSGAEMIKEQEFHYFKTENWDPDMILTAQNIIKKWNPY |
| WP_242696435.1 | Longitalea luteola | MTKLLATAACNLDIDMLNACYPLFEEERVQAIEWSFDALFDRGVLPPWFEGLLQVYSNAKRLIGHGVFFSLFSGKWLPEQAQWLETLRTTSRRFPLEQVTEHFGFMTGKDFHEGAPLAIPYSATTLRIGRDRLKRIQEAALCPVGLENLAFAYSMDEVKRHGDFLYELIQPVNGFIILDLHNVYCQLENFQISAEDLLSLYPLDKVREIHISGGSWEPVESEPDRRIRRDTHDDAVPAEVFSLLDMAIDRCPECKYVVLEQLGAGLQTAASRQQFYQDFLQMENMLDAKNKHRNGEAQNDFLPFHIVSNEPPVEDAQLYQQQRELSQILESDMSYNEAITALNQSSLAHSEWNMEQWHPAMLETAMQIARKWKKKV |
| WP_243900110.1 | Hymenobacter defluvii | MAAVPAVPSILSAIACNLDDHLLAAALPLLEAGQVEAVEWSFDTLFDGRQLPNWFAELLTAYGEQGRLIGHGVYYSLLSGRWTPEQAQWLEQLRLLAARFHFDHITEHFGSFTGQNFHTGAPLPVPYTAATLRLGQDRLLRVADACQCPVGLENLAFAYSLEEVQRHGEFLEKLLAPVNGFLILDLHNLYCQLHNFNLSTDSLLALYSLHRVREIHISGGSWEYSQVEQGRQIRRDTHDEAVPDEVFALLERTIPQCPQLKYVVLEQLGTGLRTEASQHQFRQDFQRMAAIVEASRHRAPSHQPNAFQPPAALPTGPAAHDEALHAQQRQLSQILETAASYEEAQQRLRASSLAHTDWQLEQWAPHMLETAISIAQKWK |
| WP_240773366.1 | Pontibacter sp. SGAir0037 | MAWDKVVQSYLAIMKAPEVFASIACNLDTDILSAAFPLLEEGRVEALEWSFDTLYWAEQVPDWFSELLHTYSGEERLVGHGVFFSLVSGKWTKEQQQWLEHLKQVCRQFHFDHITEHFGFFTGQNFHYGAPLNIPYTSSTLRIGQDRLMRISDACQRPVGLENLAFSYSLDEVKQHGHFLEKLVEPVNGFIILDLHNLYCQLHNFSVSFEELMQLYPLHRVREIHISGGSWESSVLDPAIRIRRDTHDDAVPAEVFQLLEYALPLCPELKYVVLEQLGNGLKLESSRIQFRQDFLKMEELVKRHTTTRKPAAKNLFLPSHPLQPGAAAEDKTLQAQQYTLSHILETATSYEEALQLLRSSSLANSEWQIESWAPHMLETAISIAQKWRK |
| MCI5081785.1 | Saprospiraceae bacterium | MKEIHATLACNLDQHVLLSALPLLAAEKVDAIEWSFDALFRTPNIPDWFVELLRTFSHARRLVGHGIFFSLFSGRWTPEQQQWLTHLRKVSHEFKFDHITEHFGFMTGENFHQGAPISVPMNERTLAIGHDRLKRMYQACERPVGLENLAIAYSLGDVERQGGFLDRLLEPVNGFMILDVHNIYCQAHNFRLDPMKIIEAYPLDRVREIHISGGSWEESEYFEGDKIRRDTHNDAVPETVFEILAKAIPLCPNLKYVVLEQIGEGLQSPHQKKRFQQDFLRLREIIQSSAQSPTAQNDFLPKGEILVGQAKEDEHLHQQQMQLSAILENAGDLASAKHLLHNSSLADTDWEIERWSDHMLDTVIQIAQKWKAGFRPPAEAGRPKQ |
| WP_293564951.1 | Phaeodactylibacter sp. | MEHSGLQRPRATLACNLDTNLLRAAHPLLAAGQVEALEWSFDALFQFPEIPDWFTALIQAYSEEGRLVGHGVFFSLFSGRWSAEQQQWLDRLKALSEVYRFDHVTEHFGFMTGADFHKGAPLGIPFTPQTLQLGRDRLKRISAAAGCPVGLENLAFAYTLDEVKRHGDFLEQLLEPINGFIILDLHNLYCQLHNFRQSAEELLRLMPLHRVREIHLSGGSWEPAATAGNRQIRRDTHDESVPQAVFDLLEKALPKCPNLKYVVLEQLGGALETPEQQQAFQADFRTMHRILKGHNHMTGKPAPLNDFQAPAFTLSATPRSDETLHQQQQALSEILENAENLRDARTQLSASPLADTAWKVEHWEDDMLETARRIAQKWKGGWG |
| WP_244674893.1 | Hymenobacter cellulosilyticus | MSTQGQPAGAAEPGILSAIACNLDADILSASFPLLEEGRVEALEWSFDTLFWAEQIPDWFTELLQAFAAQHRLVGHGVYFSLLSGRWTPEQQQWLQQLRALAGQFTFDHITEHFGFFTGQNFHSGAPLPIPYTSSALHLGQDRLRRIQDACQCPVGLENLAFAYSLDEVQRHGEFLDKLVEPVNGFLILDLHNLFCQLHNFAVPYEELIRLYPLDRVREIHISGGSWEESGQVPGRQVRRDTHDDAVPDEVFELLKLTMPRCPNLKFVVLEQLGNALQTEPSRAQFRHDFGRMEALVAEYRSRAVGRISQLFLPLQPWVAGPVAEDARLYEQQLQLSRILETAPSFEQAHRQLQASDLAHSAWKIEEWEPHMLETALRIAQKWQ |
| WP_245125988.1 | Hymenobacter volaticus | MEKQPVILSSIACNLDGDILSAAFPLLEEGKVEALEWSFDTLYWAEEVPAWFTELLSAYSEQKRLIGHGVFFSLLSSKWTKEQQQWLQKLQQLSSEFSFDHITEHFGFFTGQNFHYGAPLPIPYTTTTLRIGQDRLSRIYEACQCPVGLENLAFAYSLDEVKQHGEFLDKLIEPVNGFIILDLHNLYCQLHNFSVPYDQLLPLYPLNRVREIHISGGSWEASDRGPNQKVRRDTHDDSVPEEVFQLLELTLAKCPNLKYVVLEQLGNGLKTESSRVQFRHNFLRMESIVKRNANHLCRHSNELFLPAQRIPLGPAVENEQLHEQQVQLSQILETASSLEMARQLLASSSLAYTDWKIESWEPYMLETAIHIAQKWKR |
| WP_245957479.1 | Chitinophaga flava | MPEILSTVACNLDNDILAACLPLMEASRVEAIEWAFDTLFNVTEVPSWFEELLTAYSQENRLIGHGVFFSLFSGKWLPEQESWLKHLRKVSNDFHFDHVTEHFGFMTGKDFHHGAPLNIPYTTATLDIGRDRLKRICEACSCPVGLENLAFSYSPEEVKKHGAFLDQLLEPVNGFIILDLHNLYCQVHNFNLSFDDISMLYPLHRVREIHISGGSWENSIATPGRNIRRDTHDDAVPEEVFAYLEKAIDLCPHLKYVVMEQLGTGLYTDASKIAFRNDFLRMDDIIKNKNKTIAVANSFLPETLIVPPEVKEDLSLYQQQLELSNILETALSYETAMTALSSSSLAHSAWQIEQWDPAMIETAVNIAQKWKKKT |
| WP_241614085.1 | Chryseobacterium sp. SSA4.19 | MKPYLGLSMMAETDFVSAILPLLQNSEIEVLEWSFDTFFNAEEPNWLSNLIDFYSDNNRLIGHGVYYSLFDAKWTSRQQQWLDSLKLETQKRKYSHITEHFGFMNTENFHQGIPLPVPLCSKTLQIGKDRLSRLQNVLEIPVGLENLAFSFSADDIKQQGDFLDQLIRDIDGFLILDLHNIYCQAHNFRMDIKEIINSYPLNKVKEVHLSGGSWQESVYGKISMVRRDTHDDCIPKELFSILPFVLEKCHHLNYIIIERLGNSIKTKKEIQNFLHDFRRVRKIMDDSSAESVIEKWVKPEFLQEEPLEDLLLYQEQALLTKMLFEKENQETIRNYPFTYFNTASWDLEMIKTAENIIKKWNPY |
| WP_245096096.1 | Hymenobacter aerilatus | MSAPAILSAIACNLDDQTLTAALPLLEAGQVEAVEWSFDTLFDGRQLPDWFVELLTAYGEQGRLVGHGVYFSLLSGRWTPEQAQWLEQLRLLAGRFRFDHITEHFGAFTGQNFHTGAPLPVPYTAATLRLGQDRLLRVADACQCPVGLENLAFAYSLEEVQRHGEFLEQLVAPVNGFLLLDLHNLYCQLHNFNLLSDSLLALYPLHRVREIHISGGSWEDSRVEQGRQIRRDTHDEAVPDEVFSLLKRTIPKCPQLKYVVLEQLGTGLRTDASQDRFRQDFQRMAAIVEASRCHAHSHQPNAFQPPASLPAGPAAHDETIHTQQRQLAQILETAASYEEAQQRLRASSLAHTDWQLEQWAPHMLETAVRIAQKWKQ |
| WP_245999635.1 | Paraflavitalea soli | MSKILSTVACNLDNDILGACLPLMEESRIEALEWSFDTLYKVQEIPTWFQELLTAFSKEKRLIGHGVFFSLFSGKWLPEQEAWLKALQQTCSEFQFDHITEHFGFMTGKNFHQGAPLNIPYTPTTLAIGQDRLKRIYAACQCPVGLENLAFSYSLDEVKRHGRFLDKLLEPVNGFIILDLHNLWCQLHNFSIALEELIGLYPLDRVREIHISGGSWDHSQIEKGRTIRRDTHDDAVPAAVFQLLETVIPACPHLKYVVLEQLGNGLESIESKQQFYADFLEMESIIRRQSQPAYLTNLASFLPLSPIVMGPPVEDEGLYEQQAALSTILETSHSYEEARTLLGQSSLACSEWQVENWDPAMIETAMKIAQKWKAGFVGFIS |
| WP_246000679.1 | Pontibacter diazotrophicus | MNGKELHPVGKEPEILSTIACNLDADILSAAFPLLEEGRVEAIEWSFDTLYWAAQVPYWFTDLLLAYSQQSRLIGHGVFFSLLSGKWTKEQQHWLEQLKQLSSQYSFDHITEHFGFFTGQNFHSGAPLYVPYTSTTLRIGQDRLSRIYDACQCPVGLENLAFSYSLDEVKQHGEFLEKLVAPVNGFLILDLHNLYCQLHNFSITYEDLITLYPLNRVREIHISGGSWEASVLEPEKKVRRDTHDDAVPEEVFHLLELTVPKCSNLKYVVLEQLGNGLKTESSRNRFRQDFLKMEALVQKSRSSLSRQDDNLFLPVQPLQTGPVVEDDMLHDQQLQLSHILETAPSYEEARRLLFTSSLAGSDWKIENWAPHMLETAVSIAQKWKR |
| WP_246197742.1 | Chitinophaga agrisoli | MPKILSAVACNLDVDILGACLPLMEASKVEAIEWSFDALFKVKEIPSWFLELLTAFSDEQRLIGHGVFFSLFSGKWLQEQEDWLRQLEHTSTAFRFDHVTEHFGFMTGKDFHHGAPLNIPYTNATLTIGRDRLRRIQAACQCPVGLENLAFSYSLEEVKQHGAFLDELLAPVNGFIILDLHNLYCQLHNFEIPFDTLISLYPLDKVREIHISGGSWEDSLAAPPKKVRRDTHDDAVPDAVFELLENTIDLCPHLKYVVLEQLGNGLITAESKTQFQQDFLRMEAIVAEKNATRSTESIQSFLPAHPTIPAHIAEDATLYRQQLELSAILETAATYNDAVHALQTSSLAHSDWRIETWAPYMIETAMKIAQKWKK |
| WP_247814364.1 | Chitinophaga filiformis | MPEVLSAIACNLDANILAACLPLMEESRVEAIEWSFDALYKVKEVPSWFRELLTAFGNEQRLIGHGVFFSLFSGKWLPEQERWLRHLEQTATEFRFDHITEHFGFMTGKDFHHGAPLNIPYTAATLNIGRDRLKRIYHACRRPVGLENLAFSYSLDEVKRHGAFLEQLLEPVNGFIILDLHNLYCQLRNFDISFDEMIALYPLERVREIHISGGSWDDSAVDPSRRVRRDTHDDAVPDEVFHLLEMTMPRCPHLKYVVLEQLGNGLSTAISRQCFYNDFLQMEEIVRKHNHDRDTGSSDPFLPISPLSIDAVVEDMTLYQQQLELSAILESSGSYAEAMYALRQSSLAGSDWRIEQWEPYMIETAVKIAQKWKK |
| WP_246269322.1 | Chitinophaga oryzae | MSKLVSTVACNLDNNILAACLPLMEAAKVEAIEWSFDALYAAENVPAWFEELLAAFSKENRLIGHGVFFSLFSGRWLPEQQQWLDHLKTVSRHYRFDHVTEHFGFMTGKDFHHGAPLNIPYTPVTLGIGRDRLKRMQDACGCPVGLENLAFSYSLEEVKRHGDFLNELLEPVNGFVILDLHNLYCQAHNFDLSFEALKDLYPLHRVREIHISGGSWDESEATPGKTIRRDTHDDAVPATVFNYLDKAIDLCPSLKYVVLEQLGSGLASEESRSAFRNDFLRMDQLLNSKNRTTAPLNTFLPEAFQVPAQVPEDETLYRQQLELSQILETAASFDAAMKALQASSLAHSAWQIENWDPAMIETAAKIAQKWKRKDIVFKT |
| WP_250225925.1 | Chryseobacterium gallinarum | MEKPFLGISMMAEADFLTAILPLLRDGSIDVLEWSFDTFYHAEEPDWLTRLLNFYSQNNRLLGHGVYYSLFDAKWTERQSMWLQKLKEEVQQRKYNHITEHFGFMNTENFHQGVPLPVSLHPKTLEIGKDRLCRLQEVLEIPVGIENLAFSFSIDDVKQQGIFLEKLIEDTDGFLILDLHNIYCQACNFGLGIEEIVRLYPLEKVREIHLSGGSWQTSPYGKRRIRRDTHDDTVPAEILTVLPSVLSQCENLEYVIIERLGHTIQTETEAKDFLDDFLKVKAIIGASEWNSRHKEIKIKKKPECAGRPVEDEGLYREQTILTKLLFDNVKADHFKDFDFRYFKTENWDPEMIRTAQNIIKKWNPY |
| MCP5090420.1 | Gammaproteobacteria bacterium | MIEEDFLEAARPLFDTDTVEVLEWSFDIGWQQPSVSRRVEEIIDEYDRRGRLLGHGVTYSPMSAEFSEHQRQWLRHLERECRRRQYIHVSEHFGFMAAGGFAEAAPLPVPRTRATLGVGQDNFKRLKDAAQVPIGLEDLAFAFCERDVIEQGRFLDELLEPVDGFLVLDLHNVYCQSRNYGIPFDELLQTYPLDRVREIHVSGGSWSTLWHSAESRRVRRDTHDNSVPEEIFRGVAAMLQRCDGVVAVILERIGGSLGSELALREFRRDYERLRKIVDDFRNR |
| WP_255814981.1 | Chryseobacterium sp. MA9 | MKKPLLGISMMAEADFVSAVLPLLQNNAIDVLEWSFDTLYHTNEPDWLRDLLNFYAENNRLIGHGVYYSLFDARWTERQEEWLQKLNEEVSLRKYNHITEHFGFMNTENFHQGVPLPVSLHSKTLQIGKDRLYRLQEAVDIPIGIENLAFSFSIDDVKEQGVFLDRLTEDTNGFLILDLHNIYCQSCNFEVEMREIIDLYPLDKVKEIHLSGGSWQESVYGKKQVRRDTHDDVIPETIFSVLPSVLEQCQNLEYIIIERLGHTLKTEEDKENFLNDFNKVKTVIEASDWKRREKGSWSKQKIQIPKKPVEDLALFEEQSRLTRLLFDNAGAVVIKDQDFHYFKTDNWDPEMILTAQNIIKKWNPY |
| MCP4807725.1 | Pseudomonadota bacterium | MSERHIGVGLSVMPEPAWRDAAVPLLSDGQVDLLEWSFDTGWGRDLPAWLDPLLDFASENGVLWGHGVSYSLLTAGFPERQEAWLDNLRRELDRRSYVGISEHFGFCHAEGFRRAPPLPVPWCEATVRIGQQRLARLAEVAGVPVGLENLAFAFGRRDMLDQGPFLDELLSAVDGYLVLDVHNLWCQVQNFGVEAGSLLSRYPLDRVRVAHVSGGSWSIAGSSPWRRDTHDGPVPDEVFALLPTVINICPSLEAVVLERLGTSLEGQGQAFRDEFLRLRQVVG |
| WP_246601894.1 | Hymenobacter profundi | MAAVPAVPSILSAIACNLDDHILAAALPLLEAGQVEAVEWSFDTLFDGRQLPNWFAELLTAYGEQGRLIGHGVYYSLLSGRWTPEQAQWLEQLRLLATRFHFDHITEHFGSFTGQNFHTGAPLPVPYTAAILRLGQDRLLRIADACQCPVGLENLAFAYSLEEVQRHGEFLEKLLTPVNGFLILDLHNLYCQLHNFDLSTDSLLALYPLHRVREIHISGGSWEDSRVEQGRQIRRDTHDEAVPDEVFALLERIIPQCPQLKYVVLEQLGTGLRTEASQHQFRQDFQRMAAIVEASRHRAPSHQPNAFQPPAALPTGPAAHDEALHAQQRQLSQILETATSYEEAQQRLRASSLAHTDWQLEQWAPHMLETAISIAQKWK |
| WP_250254317.1 | Chryseobacterium sp. Marseille-Q3244 | MGRPLLGLSMMAESEFISAVLPLLQSNSIEVLEWSFDTFYNTEEPVWLSELLNFYSENNRLIGHGVYYSLFDALWTERQELWLEKLKEEFHKRKYKHITEHFGFMNTENFHQGVPLPVCLHSKTLEIGKDRLYRLQDAVEVPVGIENLAFAFSLDDVTEQGVFLDKLTEDIDGFLILDLHNIYCQSCNFNVDMRDIIKLYPLNKVREIHLSGGSWQESVYSKTPVRRDTHDDAIPKEILSILSSVVFQCKNLEYIIIERLGHTINTEEKKKSFLDDFAQVRTLIDSSDWEGVNKDLWNKKKVQFQKTPLEDMILNEEQTLLTKLLFDNAGVASIKNHEFHYFKTKDWDPEMILTAQNIIKKWNPY |

| | | |
|---|---|---|
| WP_257200057.1 | Chryseobacterium sp. WG14 | MEKPLLGLSMMAEADFVSAILPLLQNNAVDVIEWSFDTFFEAEEPSWLKDLLDFYAGNDRLLGHGVYYSLFDARWTERQEMWLQKLKQEVQKRNYNHITEHFGFMNTENYHQGVPLPISLHPKTLEVGKDRLYRLQDAVDIPIGVENLAFSFSVNDVREQGVFLDKLIDDINGFLILDLHNIYCQSCNFEVSMQDIVGLYPLEKVKEIHLSGGSWQDSVYGKKMIRRDTHDDVIPEEIFAVLPSVLSQCHNLEYIIIERLGHTIHTEEEKQCFFNDFMRIKRIISYSDTMNGKAEKWVKRERLLSEPLEDLIIYEEQIRLTKLLFDHADPTLIKNQNFHYFKTKNWDPEMILTAQNIIKKWNPY |
| WP_255373044.1 | Chitinophaga sp. YR573 | MPKILSAVACNLDANILSACLPLMEESRVEAIEWSFDALYKEKEVPSWFRELLTAFSNEKRLIGHGVFFSLFSGKWLPEQEAWLKQLEQTAAEFRFDHITEHFGFMTGKDFHHGAPLNIPYTTTTLNIGRDRLKRIYEACRRPVGLENLAFSYSLDEVKRHGTFLEQLLEPVNGFIILDLHNLYCQLRNFDIAFDEMIALYPLDRVQEIHISGGSWDDSAADPNRSIRRDTHDDSVPPEVFQLLEMTIAQCPHLKYVVLEQLGNGLVTEMSKRCFYNDFLQMEEIVRKNNNENDAGASDPFLPLFPLSTGTAVEDMMLYQQQLELSSILESSASYAEAMHVLSQSSLANSDWRIEQWEPYMIETAVKIAQKWKK |
| WP_257883251.1 | Hymenobacter sp. DG01 | MQANPAICSALACNLDVDILSAALPLLAEGRVEAVEWSFDTLFWTEQMPDWFSELLRTYSAGNRLLGHGVFFSLLSGRLSPEQQQWLSQLRKVAAELPFAHITEHFGFFTGQNFHSGAPLPVPYTPATLRLGQDRLSRLYEACRCPVGLENLAFAYSLDEVKRHGEFLDKLLEPVNGFLILDLHNLYCQLHNFSVSYEELIRLYPLHRVREIHISGGSWEPSALAPEQKVRRDTHDQGVPEEVFQLLQRTMPQCPRLRFVVLEQLGNSLRTEENRARFRLDFGRMETLVQEHRASLTVSAAEQAFLPLQPPRRGPVLEDPALHEQQQHLSRILETAPSVAEARQLLHTCPQLSATAWKIEQWEPYMLETAVGIAQKWKK |
| WP_258539328.1 | Chitinophaga oryzae | MSKLVSTVACNLDNNILAACLPLMEAAKVEAIEWSFDALYAVENVPAWFEELLNVFSEENRLIGHGVFFSLFSGRWLPEQQQWLDHLKTVSRHYRFDHVTEHFGFMTGKDFHHGAPLNIPYTPVTLGIGRDRLKRMQDACGCPVGLENLAFSYSLEEVKRHGDFLNELLEPVNGFVILDLHNLYCQAHNFDLSFEALKDLYPLHRVREIHISGGSWDESEATPGKTIRRDTHDDAVPATVFNYLDKAIDLCPSLKYVVLEQLGSGLASEESRSAFRNDFLRMDQLLNNKNKSTVPLNSFLPEALQIPAQAQEDETLYRQQLELSQILETAASFGATMKALQASSLAHSDWQIESWDPAMIETAAKIAQKWKRKTTNP |
| WP_259830879.1 | Chryseobacterium pyrolae | MSRPLLGLSMMPEADFVSAILPILQNNSVDVLEWSFDTFYDADEPDWLSGLLDFYAENNRLLGHGVYYSLFDARWTDRQEVWLKKLKEELRHRKYNHITEHFGFMNTENFHQGVPLPVPMHSRILQIGKDRLRRLQDVVDIPIGVENLAFSFSIDDVKEQGEFLDKLVEDIDGFLILDLHNIYCQSCNFEVDMQEIIKLYPLEKVKEIHLSGGSWQESAYGKKPVRRDTHDDRIPGKIVEILPEVFSQCPNLEYVIIERLGHTLNVESEKQTFFADFNRVKEMIEMSDDLPGNRQVWNKKKGEYSKPVEDLLLYDEQTKLTKLLFDEESTGSVKNQKFHYFKPESWDEEMIMTAQQIIKKWNPY |
| WP_257188871.1 | Chryseobacterium sp. WG23 | MEKPLLGLSMMAEADFVSAILPLLQNNAVDVIEWSFDTFFEAEEPSWLKDLLDFYAGNDRLLGHGVYYSLFDARWTERQKMWLQKLKQEVQKRNYNHITEHFGFMNTENYHQGVPLPLSLHPKTLEIGKDRLYRLQDAVDIPIGVENLAFSFSVNDVREQGVFLDKLIDDIDGFLILDLHNIYCQSCNFEVSMQDIVGLYPLEKVKEIHLSGGSWQDSVYGKKMIRRDTHDDVIPEEIFALLPSVLSQCHNLEYVIIERLGHTIHTEEEKQCFFNDFMRVKKILNDSGGMKRKNEKWVKKERLLSEPVEDLIIYEDQIRLTKLLFDHADPTLIKNQNFHYFKTKNWDPEMILTAQNIIKKWNPY |
| MCR9102186.1 | bacterium | MEHADLQRPRATLACNLDTNLLRAAHPLLAAGEVEALEWSFDALFQFPEIPDWFTALIQAYSEEGRLVGHGVFFSLFSGRWSAEQQQWLDRLKALAEVYRFDHVTEHFGFMTGADFHKGAPLGIPFTPQTLQLGQDRLQRISAAAGCPVGLENLAFAYTLDEVKRHGDFLEQLLEPINGFIILDLHNLYCQLHNFRQSAEELLRLMPLHRVREIHLSGGSWEPAAGESKRLIRRDTHDEAVPQAVFDLLEKALPKCPNLKYVVLEQLGGALETPRQQRAFQADFRTMHGILKRQNHSGKTSAPANDFQPPAFTLMATPRSDEALHQQQQALSEILENAANLRTARAQLSSSPLAETAWNVEDWEDDMLETARRIAQKWKGGWG |
| MCR9285899.1 | Bacteroidota bacterium | MSKILSAIACNLDQHILQTALPLFAEEKVQAIEWAFDTLFNHKNIPDWFVELLESFGKEGRLIGHGVFFSLFSGKWKPEQDEWLLRLKRMSNHFHFNHVTEHFGFMTGENFHKGAPISIPYTPTTLSIGHDRLKRIAEAAQCPVGLENLAFAYSMEEVKKHGDFLNALIEPINGFIILDLHNLYCQMHNFEVSFEEIIELYPLDRVREIHISGGSWEDSDLQPSKKIRRDTHDDSVPKEVFQILEKSIPKCPNLKFVVLEQLGLGLKTEESKINFRNDFLKMDKIISGVNSKNENIRQKSFLPSTNFQLKAPLESIQLHHQQLELSQILENALHYENAQHLLQDSSLAQSDWQIENWSPHMVETVIKIAQKWKAGFN |
| WP_258049066.1 | Hymenobacter sp. NBH84 | MKLSKALLGAVLVGITAQTTACTKGDEPAPKGEKVAKGSEKKNARLLPRVWPGLSNAPETMAAVPALPSILSAIACNLDDHLLAAALPLLEAGQVEAVEWSFDTLFDGRQLPNWFAELLTAYGEQGRLIGHGVYFSLLSGRWTPEQAQWLEQLRLLAARFHFDHITEHFGSFTGQNFHTGAPLPVPYTAAILRLGQDRLLRIADACQCPVGLENLAFAYSLEEVQRHGEFLEKLLAPVNGFLILDLHNLYCQLHNFDLSTDSLLALYPLHRVRGNPHFGRQLGRFAGGAG |
| WP_259835626.1 | Chryseobacterium herbae | MKKPLLGLSMMPEADFVSAILPILQTHSVEVLEWSFDTFYDVEEPEWLSGLLDFYAGNNRLLGHGVYYSLFDARWTDRQETWLKKLKEEVSRRKYNHITEHFGFMNTENFHQGVPLPVPLLPKTLEIGKDRLIRLQDAVEIPVGVENLAFSFSLDDVKEQGEFLDKLVEDIDGFLILDLHNIYCQSCNFEVDMKEIIKLYPLEKVKEIHLSGGSWQESAYGKKPVRRDTHDDRIPEEILQMLPEVLIQCHPEYVIIERLGHTLNTEAEKEFFFDDFNRVKEIIEFSDHPVDHEKIWNRKNSTYAQPIEDLLFHDEQTELTRLLFDGNSTEAIKNRNFHYFKPENWDEEMIITAQQIIKKWNPY |
| WP_261511996.1 | Chryseobacterium paludis | MKKPLLGLSMMAEADFVSAILPLFENNQIEILEWSFDTFYDVEEPEWLKGLLDFYSQNNRLIGHGVYYSLFDAKWTERQEEWLNNLKTETEKRHYNHITEHFGFMNTENFHQGIPLPVSLNSKTLQIGKDRLLRLQDVVNVPIGIENLAFSFSIEDVKQQGQFLDQLITDIDGFLILDLHNIYCQAYNFDVQLEEIISAYPLEKVKEIHLSGGSWNQSIYTKNKLIRRDTHDDAIPEEILNILPFVLPKCENLQYIIIERLGCTIKTKDEKEAFFDDFRRVREIIDSIEYHENNYPAIWEKKNVNLVSPLEDLQLYEDQTRLTQLLFDDKPELIKSQTFHYFKTDDWDIEMINTAKSIIKKWNPY |
| WP_260545184.1 | Chryseobacterium oranimense | MKKPLLGLAMMPEADFVSAILPVLQNNSVDVLEWSFDTFYDVEEPEWLSGLLDFYAENDRLLGHGVYYSLFDARWTSRQKDWLQKLKEEFKHRKYNHITEHFGFMNTENFHQGVPLPVPLHSKTLQIGKDRLKRLQDTVEVPMGVENLAFSFSIDDVKEQGEFLGKLIEDIDGFLILDLHNIYCQSCNFEIDMQEIIRMYPLEKVKEIHLSGGSWQESAYGNKPVRRDTHDDYIPEEIINVLPEVFSQCLNIEYVIIERLGHTLDTEVKKQLFFDDFKKVKAIETSDFPAGITKEWNRKKREYSEPVEDMLLYEEQTKLTKMLFEGNTVQSVKNEPFHYFKPENWDKEMITTAQQIIKKWNPY |
| MCU0533392.1 | Hydrococcus sp. Prado102 | MFRQKQTLPWVGLSLMADPSFFQATRPLFEAGDVDVLEWSFDVGWGLESLPSWVEQLLEFYSQRDRLLGHGVSYSLLSAQWSDRQAQWLKLLKSECQRYRYRQISKHFGWMAAGNFERSAPLPMPLTRDTLRIGCMNLRRLTEVTSSPIGLENLAFAFGLPDVLNQGNFLEQLLQSVNGFLVLDLHNVFCQIHNFDRAADEIINSYPLHRVREIHISGGSWSQSTRGDRGLMRRDTHDGDVPEEVFQLLALVLKKCPTTESIILERLGKTLDDEAKMARFRQDFYRIKEIVKNSLCKQKN |
| MCU0328123.1 | Chitinophagales bacterium | MYSTLAINPYPHLLTTALPLLEAGKVEALEWSFDTLPFPQIPEWFQELLQTYSQKDRLIGHGVFFGLMKGNFSQNQQEWLNQLKGISKKIKFHHISEHFGYFSGRDFHSGAPLPVPMNKITLDIGIDRLQRIQEASECPVGLENLAFAGNIEEVKYHGEFLDKLLSPVQGFIVLDLHNIYCQMVNFDLRLEEILASYPLNKVREIHISGGSWDTVSSLKKIRRDTHDDSVPTEVMTLLKKTIPLCPCIEFITLEQMDYALITDQMQQNFRNDFEEIQKIVFSTSLPKSDLIITNKENSDIHLYATPIESEELNRAQELFSTLLEDSDNLNQLKINLFKVQNTLSILEVNQWQDDMLETAMRIAKKWR |
| WP_264518140.1 | Chryseobacterium viscerum | MKKPLLGISMMAEADFVSAVLPLLQNNAIDVLEWSFDTLYHTNEPDWLRDLLNFYAENNRLIGHGVYYSLFDARWTERQGKWLQKLKEEVSLRKYNHITEHFGFMNTENFHQGVPLPVSLHLKTLQIGKDRLYRLQEAVDIPVGIENLAFSFSIDDVKEQGVFLDKLTEDTDGFLILDLHNIYCQSCNFEVEMQEIIDLYPLDKVKEIHLSGGSWQESVYGKKQVRRDTHDDVIPETIFSVLPSVLEQCQNLEYIIIERLGHTLKTEEDKENFLNDFNKVKTMIEASGGKEREKSSWSKQEIQFSKKPVEDPALFEEQSRLTRLLFDNVGTAVIKDQDFHYFKTDNWDPEMILTAQNIIKKWNPY |
| WP_265130928.1 | Chryseobacterium oranimense | MNKPLLGLAMMPEADFVSAILPVLQNNSVDVLEWSFDTFYDVEEPEWLSGLLDFYAENDRLLGHGVYYSLFDAQWTSRQKDWLKKLKEETKRRKYNHITEHFGFMNTENFHQGVPLPVPLHAKTLQIGKDRLKRLQDTVEVPVGVENLAFSFSMDDVKEQGEFLEKLVEDIDGFLILDLHNIYCQSCNFEIDMQEIIRMYPLERVKEIHLSGGSWQESAYGNKPVRRDTHDDNIPVEITDVLPEVFSQCPNLEYVIIERLGHTLNTEVKKQLFFDDFNKVKEIIEVSDFPAGIKKAWNKRKGESSEPVEDLLLYEEQTKLTKILFEGNTVQSVKNETFHYFNPENWDKEMITTAQQIIKKWNPY |
| WP_273635928.1 | Chryseobacterium rhizosphaerae | MGKPLLGLSMMAEADFVSAILPLLQDNAVDVVEWSFDTFFAAEEPSWLKDLLNFYAGNGRLLGHGVYYSLFDARWTERQEMWLQKLKHEVQKRNYNHITEHFGFMNTENYHQGVPLPLSLHPKTLEIGKDRLYRLQDAVPIPIGVENLAFSFSVDDVREQGVFLDKLVDDIDGFLILDLHNIYCQSCNFDVSIQEIVSLYPLEKVKEIHLSGGSWQDSVYGKKMIRRDTHDDVIPEEIFSVLPSVLSQCHNLEYVIIERLGHTIHTEEEKQRFFNDFMRVKSILNDSDGMKRKKEKWVKREMLLPEPVEDSILYEEQIRLTKLLFDHIDPTLIKNQNFHYFKTENWDPEMILTAQNIIKKWNPY |
| WP_276161883.1 | Chryseobacterium arthrosphaerae | MKKPLLGLSMMPEADFVSAILPLLQNNSVEVLEWSFDTFYEAKEPDWLDELLNFYSENQRLIGHGVYYSLFDALWTERQENWLKKLKEEVRKRKYSHITEHFGFMNTENFHQGVPLPVPLHPKTLLIGKDRLYRLQDAVELPVGIENLAFSFSIEDVKEQGVFLDNLIADINGFLILDLHNIYCQACNFEVEMEDIIRLYPLDKVREIHLSGGSWQESVYARKMIRRDTHDDLIPEEIFSVLPAVITQCNNLEYVIIERLGHTIKTDQEKRAFFDDFTRVRRIIDDSEIKTGKKIRWGQKETGFSNSPVEDDALHEEQSRLTKLLFDSSGAEMIKEQEFHYFKTENWDPDMILTAQNIIKKWNPY |
| WP_267406217.1 | unclassified Chryseobacterium | MKPHLGLAMMAEADFVSAILPLLQSNEIDVLEWSFDTSYNADEPDWLNELLDFYSENNRLIGHGVYYSLFDAKWTDRQEDWLKKLKSETQKRHYNHITEHFGFMNTENFHQGIPLPISLHQKTLQIGKDRLSRLQDVLNVPVGIENLAFSFSIDDVKEQGEFLEKLIEDIDGFLILDLHNIYCQAHNFDVDIQDLINSYPLDKVKEIHLSGGSWQKSVYGKEKLIRRDTHDDAIPEELFNVLLYVFSKCNNVEYVIIERLGNTITTGIEKENFFDDFKKVREIMNVSDFQLNSKKWNKPDLSLTEPLEDLSLYKEQVLLTKMLFENEKPELIKEYPFSYFNTENWDLEMLNTARNIIKKWNPY |
| WP_269427484.1 | Pedobacter punctiformis | MNKPYLGISIMPEHDFLQAVLPLLQDGDIETIEWSFDTIKFEKYKPKWLNRLLQEYADNNRLIGHGVYYSLFQAKWSKEQANWLKKLKKEVEKYNYRHISEHFGFMSDHVDHHKGCPFPVPLNKTTLAIGIDRLQRLQDAAQLPVGVENLAFSFSKYDVKRQGEFITKLIEPINGFVILDLHNIYCQSENFGIEMMDLIDCYPLDSVKEIHISGGSWQESVYQGVIKRIRRDTHDGNVPKKIFDVLPAVLQKCKSVEFVIFEKMGNSFQKPGDFVDYRNDFNKLKNIIENTTLKFEAKEWGNKIIPPKNPIEDGLLFTEQNLLSRTLLESSNYKELLSKISLKNWEIDKWDPRMIDTAIALSKKWS |
| WP_276349236.1 | Daejeonella sp. JGW-45 | MPAIIPSIACNLDADILSASLPLFSEHRVGGIEWSFDALYGIEQVPEWFTELLTAYSNERRLVGHGVYFSIFSGRWSEDQQNWLNELKLLSQKFKFDHISEHFGFMTGKSFHHGAPLSIPYTGSTLRLGRDRLSRIYDACRCPVGLENLAFSYSLDEVKRHGEFLEKLVEPLNGFIILDLHNLYCQVHNFSADFEDLIKLYPLDRVREIHISGGSWEQSVLTGKQVRRDTHDESVPDEVFDLLRKTLSRCPNLKYVVLEQLGDALKTSGSRKAFYSDFIEMESIIAAGSKSRPDTQTNLFSPDYNLVSDQVAEDHDLYLQQRELSGILETSGSYEQAVDRLKLSSLHNSEWDTQNWQPYMLETAMNIAQKWKSTDD |
| WP_276484500.1 | Paraflavitalea pollutisoli | MLEACLPLLEEARVEALEWSFDVLYQTAETPAWFHDLLTAFSKEERLIGHGVFFSLFSGRWLPEQDVWLQHLRTCCATFHFDHITEHFGFMTGTDFHQGAPLNIPYTASTLAIGQDRLKRIYEACACPVGLENLAFAYSIDDVQRQGSFLDELLHPINGFIILDLHNLWCQLHNFSIPYEQLIHLYPLDRVREIHISGGSWDDSLLETGRRIRRDTHDDAVPAAVFDLFRQTIPRCPHLKYVVMEQLGNGLTSHKSKQQFYADFVTMASIVDTANQQREAGATHAFLPAEQLYLGQPLEDETLYQQQRELANILESAGSFDAARHLLLTSSLAGTAWHVEQWDAAMIETAVKIAQKWKEGF |
| WP_277235126.1 | Hymenobacter sp. YC55 | MHPDSAMEKQPTILSSIACNLDANILSAAFPLLEEGKVEALEWSFDTLYWAEQVPDWFTELLSAYSEQKRLIGHGVFFSLLSSRWTKEQQQWLQKLQQLSLEFNFDHITEHFGFFTGQNFHYGAPLPIPYTDITLRIGQDRLKRIYEACQCPVGLENLAFAYSLDEVKQHGEFLEKLIEPVNGFVILDLHNLYCQLHNFSVSYDELIALYPMDRVREIHISGGSWEDSALGPDQKVRRDTHDDSVPKEVFQLLERTICRCPNLKYVVLEQLGNGLKTESSRTLFRHDFLEMERIVKLNASSLSTRPNELFLPPQLLLTGPAVEDEPLYEQQLQLSQILETAPSLERARQLLAASSLAQSDWKIESWEPYMLETAIHIAQKWKK |
| WP_276964884.1 | Chryseobacterium sp. | MLGVSMMAEADFVSAVLPLLQNNSIDVMEWSFDTLYHANEPDWLRDLLNFYAENNRLIGHGVYYSLFDARWTERQEEWLKKLKEEVNLRKYNHITEHFGFMNTENFHQGVPLPVSLHPKTLQIGKDRLYRLQEAVNIPVGIENLAFSFSIDDVKEQGVFLDKLTEDTNGFLILDLHNIYCQSCNFELEMKEIIDLYPLDKVKEIHLSGGSWQDSVYGKKQVRRDTHDDVIPRDILSVLPSVLTQCENLEYVIIERLGHTLKTDEERDHFLDDFSKVKTIIETSELNGKEKSNWSKMKVELSEKPLEDPVLYEEQSRLTQLLYDNADVKAIKDLDFYYFKTENWDSEMILTAQNIIKKWNPY |
| WP_264691061.1 | Hymenobacter sp. YIM 151858-1 | MYATIACNLDADLLATAFPLLQQGRVEALEWSFDTLYGAGQMPEWFTALLRTYAEAGRLVGHGVFFSLLAGKWSPEQQQWLAHLGQLAGQFRFDHITEHFGFFTGQNFHHGAPLPIPYTAAALRIGQDRLRRIHDACRCPVGLENLAFAYTLEEVKRHGAFLEELLLPVNGFLILDLHNLYCQLHNFGVAYDALIALYPLERVREIHISGGSWHDSEQAPGRRIRRDTHDGAVPDEVFRLLALTIPRCPNLKFVVLEQLGHALHTESSRTQFRQDFGRLEHLVAQHRAALRPEPTNDFRPRQTALPAAPAAEDEHLHEQQRQLSDILENAPSYAEAERQLRASALAHSDWRIEQWQPHMLETALHIARKWKP |
| WP_277694591.1 | Chryseobacterium sp. WX | MRKPILGVSMMAEPDFVSAILPLLHNNSIEVMEWSFDTLYHTNEPDWLRDLLNFYAENNRLIGHGVYYSLFDARWTARQEEWLKKLKEEVSLRKYNHITEHFGFMNTENFHQGVPLPVSLHSKILQIGKDRLYRLQDAVNIPVGIENLAFSFSIEDVKEQGVFLDKLTEDTNGFLILDLHNIYCQSCNFELEMQQIIDLYPLDKVKEIHLSGGSWQDSVYRKKQVRRDTHDDVIPKDILSVLPSVMKKCPNLEYIIIERLGHTLKTEEERNYFLADFNKVRTIIETSDLKLKQQNGWCKEEIKLPAKPLEDLLLYEEQSRLTRLLYENANVENIKHLDFHYFKTENWDAEMILTAQNIIKKWNPY |
| WP_276878162.1 | Chryseobacterium joostei | MRKPLLGLSMMAETEFVSAILPLLQSNSVDVLEWSFDTLYHTKELDWLGDLLNFFSENNRLIGHGVYYSLFDARWTERQNIWLNKLKKEVKKRKYNHITEHFGFMNTENFHQGVPLPVSLHSKTLQIGKDRIFRLQEALDIPVGIENLAFSFSIDDVREQGLFLDKVIEDTDGFLILDLHNVYCQSCNFEVEMQEIINSYPLDKVKEIHLSGGSWQNSVYGKKLIRRDTHDDVIPEEILSVLPWIFSQCQNLEYVIIERLGHSILLESEKDAFIEDFAKVEKLISESDWRNQKKINWNKKELKLSESPLEDMLLHEEQTMLTKLLFDSADVSLIKNQEFHYFKTDTWDTEMILTAQNIIKKWNPY |
| WP_029449562.1 | Cellulophaga baltica | MGNKKPKLGIAIAPSLKFLEAALPLFAGEKIEIIEWSFDTLKDAADEPAWLSLLLKEYGEKNRLLGHGVFYSLLDANWSSRQENWLKKVRQETLSHKYCQISEHFGLMSSANAHSGFPLPIQLSNPVLQIGIDRLKRLQATAQVDVGIENLALTANVADILEQGEFLFKLVNPVNGFVILDLHNIYCQSENFNIDMMTIIKSYPLSLVKELHISGGSWDTDPTLSRKIRRDTHDGRIPEVIFEILPEVLKICPFLAFIIFEKIEDSFLTEKDGIDFRADFQKIREIIDATAFSVTPKEKKQVAAILGPPVIDVELLQAQVALRESIALGTYHTNTKWDKDMWKVATKLYEKWNNY |

| | | |
|---|---|---|
| WP_168737805.1 | Chitinophaga eiseniae | MEAARVEAIEWSFDAVEQAPSWFEELLTVYSQENRLIGHGIFFSLFSGKWLPEQQQWLSHLHKISRRYHFDHVTEHFGFMTGKDFHHGAPLNIPYTPVTLNIGRDR<br>LKRIQGACECPVGLENLAFSYSLEEVKKHGDFLERLLEPVNGFIILDLHNLYCQAHNFGLSFEALKELYPLHLVREIHISGGSWEDSDAMQGRIVRRDTHDNAVPEAV<br>FIYLEKAIDLCPSLKYVVMEQLGTALTTTDSQATFRSDFLRMDQIIRNKNRMTASLNTFMPLPLSIQPRPLEDIPLYQQQLELSDILETAGSFDTLMEALQASSLAHSD<br>WRIERWDAAMIETAASIAQKWKKK |
| WP_169229115.1 | Chitinophaga fulva | MEEARVEAIEWSFDALYAVEEVPAWFGELLSVFSNENRLIGHGVFFSLFSGRWLPEQQQWLNDLKAVSRRYQFDHVTEHFGFMTGRDFHHGAPLNIPYTPVTLEI<br>GRDRLKRMQDACSCPVGLENLAFSYSLEEVKKHGDFLNELLEPVNGFMILDLHNLYCQAHNFNLTFEALNSLYPLHRVREIHISGGSWEVSGATPGKTVRRDTHDD<br>AVPEEVFAYLDKAIDFCPSLKYVVLEQLGSGLDCDSSRAAFGHDFLRMDELIKYKNKTAAPLNNFLPEALQIPAQAREDESLYQQQLELSQILETATSFDALMKALQA<br>SSLAHSAWQIESWDPAMIETAAKIAQKWRRKDPIF |
| WP_157303704.1 | Chitinophaga oryziterrae | MEESRVEAIEWSFDALYKVKEMPSWFRELLTAFSNEKRLIGHGVFFSLFSGKWLPEQESWLRHLERTAAEFRFDHITEHFGFMTGKDFHHGAPLNIPYTTTTLNIGR<br>DRLKRIYEACRRPVGLENLAFSYSLDEVKRHGTFLEQLLEPVNGFIILDLHNLYCQLRNFDISFDEMIALYPLDRVREIHISGGSWEDSAVAPNRSIRRDTHDDAVPPE<br>VFQLLEMTIAKCPHLKYVVLEQLGNGLVTETSKRCFYNDFLQMEDIVRKNSNESNTPDPFLPLFPLSTGPVVEDMTLYQQQLELSSILESSASYAEAMHVLTQSSLA<br>NSDWRIEQWEPYMIETAVKIAQKWKK |
| WP_168764034.1 | Chitinophaga polysaccharea | MEAARVEAIEWAFDILYTAEQVPSWFEELLTAYSRENRLIGHGIFFSLFSGKWLPEQQQWLNHLRKTSRYYHFDHVTEHFGFMTGKDFHHGAPLNIPYTPVTLNIGR<br>DRLRRMQDACECPIGLENLAFSYSLEEVKRHGDFLEKLLEPVNGFIILDLHNLYCQAYNFGLSFEALKGLYPMHLVREIHISGGSWEDSETMPGRIVRRDTHDDAVP<br>EAVFIYLEKAIDFCPSLKYVVMEQLGTGLTTAGRQAAFRNDFHRMDQLIRNKNRTAAPVNTFIPAPLSLPTLAIEDNFLYQQQLELSAILETAGSFDTLMEALQGSSLA<br>HSDWQIESWDPAMIETAARIAQKWKKK |
| WP_188091977.1 | Chitinophaga qingshengii | MEAASVEAIEWSFDTLYAVEEVPAWFEELLTAYSQENRLIGHGVFFSIFSGRWLPEQQQWLHHLKKISQQYRFDHVTEHFGFMTGKDFHQGAPLNIPYTAVTLDIG<br>RDRLKRMQDTCRCPVGLENLAFAYSLEEVKRHGDFLNELLEPVNGFIILDLHNFYCQAHNFELSFDELKSLYPLHRVREIHISGGSWDHSEAMPGKTVRRDTHDNA<br>VPDTVFSYLDKAIDLCPSLKYVVLEQLGTGLTTVDSQVAFRNDFLRMNQLIKNNNNTSLPENSFLPQPLQLPAQVREDESLYQQQLELSHILEAAVSFDAVMCALQA<br>SSLAHSDWQIERWDPAMIETAAKIAQKWKHGWT |
| WP_236015555.1 | Chitinophaga solisilvae | MPEIRSAVACNLDANILAACLPLMEASKIEAIEWSFDALYKVKDIPDWFVELLTAFSNENRLIGHGIFFSLFSGRWRQEQEQWLQHLRKTSTHFRFDHITEHFGFMTG<br>KDFHHGAPLNIPYTKSTLNIGVDRLKRIHDACRCPVGLENLAFSYSMDEVKRHGAFLDELLEPVNGFIILDLHNLYCQLHNFDITFGEMIILYPLHKVREIHISGGSWEA<br>SPAAPERMVRRDTHDDAVPEEVFQLLQKTIPLCPNLQYVVLEQLGTGLETDTSRTLFRQDFGRMQRIVHEQQPQTGSCPVNSFMPDNLLISAVPAADETLFQQQLE<br>LSAILESASSCEAAVHALQHSSLAHSDWLIEQWQPYMIETAVSIAQKWKKI |
| WP_169190781.1 | Chitinophaga sp. Ak27 | MEAARVEAIEWAFDTLYTAEQVPSWFEELLTAYSRENRLIGHGIFFSLFSGKWLPEQQQWLNHLRKTSRYYHFDHVTEHFGFMTGKDFHHGAPLNIPYTPVTLNIG<br>RDRLRRMQDACECPIGLENLAFSYSLEEVKRHGDFLEKLLEPVNGFIILDLHNLYCQAYNFGLSFEALKGLYPMHLVREIHISGGSWEDSETMPGRIVRRDTHDDAV<br>PEAVFIYLEKAIDFCPSLKYVVMEQLGTGLTTAGRQAAFRNDFHRMDQLIRNKNRTAAPVNTFIPAPLSLPTLAIEDNFLYQQQLELSAILETAGSFDTLMEALQGSSL<br>AHSDWQIESWDPAMIETAARIAQKWKKK |
| WP_168873219.1 | Chitinophaga varians | MDAARVEAIEWSFDALYAVTKVPVWFEELLTVFSNENRLIGHGVFFSLFSGRWLPEQQQWLNDLRAVSRRYQFDHVTEHFGFMTGRDFHHGAPLNIPYTPTILEIG<br>RDRLKRMQEACSCPVGLENLAFSYSLEEVKKQGDFLDKLLGPVNGFMILDLHNLYCQAHNFNLSFEALISLYPLHRVREIHISGGSWEVSGAAPGKTVRRDTHDDA<br>VPEEVFGYLDNAIDLCPSLKYVVLEQLGAGLDSDSSRAAFGHDFLRMDDLIKQKNETIAPLNSFLPEALQIPAQPPEDESLYRQQLELSQILETAVSFDGLMQALKAS<br>SLAHSAWQIESWDPAMIETAAQIAQKWKKR |
| WP_188100661.1 | Chitinophaga varians | MEAAQIEAIEWSFDALYAVEKVPSWFEELLTVFSDENRLIGHGVFFSLFSGRWLPEQQQWLDHLKAVSRRYRFDHVTEHFGFMTGRDFHHGAPLNIPYTPVTLEIG<br>RDRLKRMQDACGCPVGLENLAFSYSLEEVKKHGDFLNELLEPVNGFIILDLHNLYCQTHNFNLTFEELNKLYPLHRVREIHISGGSWEISGATPGKTVRRDTHDDAV<br>PEEVFAYLDKAIDLCPSLKYVVLEQLGTGLDTEDSRIAFGSDFLRMDELIKNKNKTAAPLNNFLSEALPVAAQAREDESLYQQQLELSQILETATSFEALMKALHASSL<br>AHSAWQIESWDPAMIETAAKIAQKWKRKDVIL |
| WP_207335620.1 | Fibrella aquatilis | MAAALPLFEAGEVGAIEWAFDSLYRYDELPGWFADLLTTYGEAGRLVGHGVFFSLFSGQWTPEQAAWLTSLRALSQRFRFDHVTEHFGFMTGANFHQGAPLGVP<br>CTPQTLALGRDRLQRIQDACGCPVGLENLAFAYAPDEVKQHGEFLRQLIEPINGFIILDLHNLYCQLMNFDISAPDLLALYPLDRVREIHISGGSWHDSAADPTRAIRR<br>DTHDDAVPDAVFNLLTYVMDQCPNLRFVVLEQLGTALDTDESRQQFRADFRRMDAIVKAMDTDGNGLVNSFLPPKQASVSNPPLTDAALHAQQRTLATILETATDY<br>QQAQTRLASSELAHTAWQPERWQPAMLETAIAIAQKWRRGIEGKL |
| WP_085413297.1 | Fibrella sp. ES10-3-2-2 | MAAIRSTLACNLETNLILAALPLFEAERVAALEWSFDALYAHEQLPDWFTELLTTYSEADRLVGHGVFFSLLSGRWTSEQQAWLIRLRALSREFRFDHITEHFGFMT<br>GADFHKGAPVNVPCTAVTLALGQDRLSRIQDACRCPVGLENLAFAYSLDDVKRQGDFLRQLIEPVNGFIILDLHNLYCQSKNFNLDLTALLDLYPLDRVREIHISGGS<br>WAEVEIEPGRHIRRDTHDDAVPDDIFNWLSTAIDRCPNLGYVVLEQLGAGLTTGSERLAFQADFDRMDVIVQQKNKSIAHPDMQSFLPELPLLINPVPLENHRLYTH<br>QLELSAILETAVDVQQAKRRLAESSLANSEWEVERWEPAMLETAVAIARKWKNGFVGKE |
| WP_214460436.1 | Flavihumibacter fluvii | MWFEDLLAAYSAQNRLIGHGVFFSLFSGRWSDAQQNWLAALRQTSQRYRFDHITEHFGFITGKDFHQGAPMSIPYTPATLAIGQDRLKRIQDACSCPVGLENLAFS<br>YSLDEVKEHGEFLQRLIEPVNGFIILDLHNLYCQINNFDIGFEEIISLYPLDKVREIHISGGSWDEVSEPFPKNVRRDTHDSAVPEPVFDLLKMTIEKCPALKYVVLEQLG<br>SALSTPESQAVFQQDFLILEEIVQEKNKGRWDFFNDAFLPSPLVIPDDIIEDEALYQQQKQLAEILEHCTTAAEAQQQMRQTTLANSAWDIEHWQPYMLETAIRIAQK<br>WKDGFE |
| WP_246398786.1 | Hymenobacter | MSAALPLLAEGRVEALEWSFDALFWAGQIPDWFSELLRTYSASGRLLGHGVFFSLLSGRWSREQQQWLTQLRQLANQLPFAHITEHFGFFTGQNFHSGAPLPVPY<br>TPVTLRLGQDRLSRLYEACQCPVGLENLAFAYALEEVKRHGEFLDKLLEPVNGFLILDLHNLYCQLHNFSISYEDLIDLYPLHRVREIHISGGSWEASALAPGQRVRR<br>DTHDQGVPEEVFQLLQRTMPRCPHLRFVVLEQLGNSLRTEDSRARFRLDFGRMEALVQEHRAGLTASADQAFLPLRPPQPGPALEEPALHEQQQHLSRILETAPS<br>VAEARQLLLASPLLASTDWNIEQWAPHMLETAVRIAQKWK |
| WP_223847018.1 | Hymenobacter montanus | MSAAFPLLEAGEVEALEWSFDALMGEMPEWFSTLLSVYGEQGRLVGHGVYFSLLSGQWTPEQQQWLHDLGQLAARFSFDHVTEHFGFFTGESFHAGAPLPVPC<br>SAATLRLGRDRLRRIQDAGRCPVGLENLALAYSLADVQQQGAFLDQLLEPVNGFLILDLHNLYCQLHNFAIAPETLIDLYPLDRVREIHISGGSWEASARAPRGRVRR<br>DTHDGAVPEEVFQLLERTMPRCPHLKYVVLEQLGTALKTERQRAGFRRDFRRMARLVQQHDRATTRGSDQLFLPLHPSPGGPVVVDEKLHGQQRQLSRILEAAP<br>TLDEAQRLLHGSSLAHTEWHLENWPPYMLETARRIAQKWKR |
| WP_262904639.1 | Hymenobacter lucidus | MSAIACNLDADILSASFPLLEQGRVEALEWSFDALFWAEQVPEWFTDLLRAFADQKRLVGHGVYFSLLSGRWTREQQQWLEQLRELAGRFAFDHITEHFGFFTGQ<br>NFHSGAPLPIPYTSAALRLGQDRLRRIQQAAGCPVGLENLAFAYSLDEVQRHGEFLDQLVEPVNGFLILDLHNLFCQLHNFAVPYEELIKLYPLERVREIHISGGSWE<br>KSGQEPGRQVRRDTHDDSVPEEVFELLALTLPRCPNLKYVVLEQLGNALKTEASRTQFRADFGRMETLVGEFRRRAAPRASQLFLPPQPWLAGPVAEDAHLHEQ<br>QQQLSRILETAPSFEAARQQLHASSLAHSAWKIEEWAPHMLETALRIAQKWKQ |
| WP_208419998.1 | Paraflavitalea devenefica | MEADILGACIPLWEEARIEAIEWSFDALFNVEEIPAWFRELLTAFSKEKRLIGHGIFFSLFSGKWLPEQQQWLNKLQQTCTEFQFDHITEHFGFMTGKDFHQGAPLNI<br>PYTTSTLAIGRDRLQRIQEACQCPVGLENLAFSYSLDEVKRHGTFLDQLLEPVNGFIILDLHNLWCQLHNFSIPFEEVIDLYPLHRVREIHISGGSWDDSEVEAGRTIR<br>RDTHDNAVPAAVFQLLEMTISRCPHLQYVVLEQLGTGLLTPESKQQFYQDFGRMESIVSEKNLSRTPHSTCSFLPQAPLQKGGVTEDLFLYQQQRELSAILESAPSY<br>EDARRLLAQSSLAHSDWKIENWDPAMVETAIKIAQKWKQGWLT |
| WP_237487918.1 | Hufsiella arboris | MIDAYSKEGRLIGHGVFFSIFSGRWSIQQQEWLDNLKKLSARFKFDHITEHFGFMTGANFHYGAPLAVPFNSKTLTIGRDRLNRIYDACQCPVGLENLAFSYSIDEVK<br>KHGEFLETLVEPVNGFIILDLHNLYCQMHNFSISFDEIIKLYPVDRVREFHISGGSWENSSLSGKRIRRDTHDDAVPDEVYQLLDSALDHCKNVRYVTMEQLGSGLKT<br>EESRSNFYNDFLKMQQIVRNKTSVPNDAENTFFPPKIVPPGDIIEDENLYIEQQQLSYILENSTSYDDAVSRIKSSVLAKSDWGIESWAPHMLETAISIAQKWKKK |

### **Table S3. Calculated $[M+H]^+$ masses for peptides characterized in this study.**

$[M+H]^+$ masses shown are monoisotopic (in Da) unless otherwise noted.

| Peptide | Peptide sequence | Modification | Expected $[M+H]^+$ |
|---|---|---|---|
| DybA | GSSHHHHHHKISKPVLQAVAVAVALSTITACADMD<br>IKPEKDKKAKTRTLDPCPACGMG | N/A | 6137.12 |
| DybA + DybH | GSSHHHHHHKISKPVLQAVAVAVALSTITACADMD<br>IKPEKDKKAKTRTLDPCPACGMG | -4.03 | 6133.09 |
| DybA + DybH (LysC digested fragment) | TRTLDPCPACGMG | -4.03 | 1317.54 |
| ChrA | GSSHHHHHHSQMKIPALVMASLLAVSVSGQTTKP<br>VKKGTKSVKKVKKMDSPKTVKAEPTKVVKRDTILK<br>HGGGCPACGMG | N/A | 8442.54 |
| ChrA + ChrH | GSSHHHHHHSQMKIPALVMASLLAVSVSGQTTKP<br>VKKGTKSVKKVKKMDSPKTVKAEPTKVVKRDTILK<br>HGGGCPACGMG | 9.98 | 8452.52 |
| ChsA | MHHHHHHKLSKSLLSAIMIGIAVQTTVVSCSKDEQ<br>VKPKKADQANKQSESKPVDNPDSCPACGMG | N/A | 7013.45 |
| ChsA + ChsH | MHHHHHHKLSKSLLSAIMIGIAVQTTVVSCSKDEQ<br>VKPKKADQANKQSESKPVDNPDSCPACGMG | -4.03 | 7009.42 |

| | | | |
|---|---|---|---|
| HymA | MHHHHHHKLSQAILGAVLVGLTAQTTGCIKKSDPT<br>PKEEQGKSGKKSPEIPYNCPGCGLG | N/A | 6351.22 |
| HymA + HymH | MHHHHHHKLSQAILGAVLVGLTAQTTGCIKKSDPT<br>PKEEQGKSGKKSPEIPYNCPGCGLG | -4.03 | 6347.18 |
| ArlA | GSSHHHHHHNQNLREKATQLLDVASNTQGMKSS<br>SPLPCPACGMMMALPESSPQLVTPATR | N/A | 6453.07 |
| ArlA + ArlH | GSSHHHHHHNQNLREKATQLLDVASNTQGMKSS<br>SPLPCPACGMMMALPESSPQLVTPATR | -4.03 | 6449.04 |
| MelA1 (LysC digested fragment) | LFPQPAPCPACGMG | N/A | 1388.62 |
| MelA1 + MelH + MelHc (LysC digested fragment) | LFPQPAPCPACGMG | -4.03 | 1384.58 |
| DybA His-embed + DybH | MKISKPVLQAVAVAVALSTITACADMHHHHHHDIK<br>PEKDKKAKTRTLDPCPACGMG | -4.03 | 6033.04 |
| lipo-DybAH | CADMHHHHHHDIKPEKDKKAKTRTLDPCPACGM<br>G | -4.03<br>(+800.73) | 4641.47 |
| lipo-DybAH (observed N-terminal LysC fragment) | CADMHHHHHHDIKPEK | 800.73 | 2772.60 |
| NedA | MKTTSTLMALAGALVLAGCQVEPAQSSSPEVSVQ<br>QKAGEGKCGAASQAGKAAEGKCGEGKCGSKHH<br>HHHH | N/A | 7008.90 (ave.) |
| NedA + NedBC | MKTTSTLMALAGALVLAGCQVEPAQSSSPEVSVQ<br>QKAGEGKCGAASQAGKAAEGKCGEGKCGSKHH<br>HHHH | -12.10 | 6996.80 (ave.) |
| lipo-NedABC | CQVEPAQSSSPEVSVQQKAGEGKCGAASQAGKA<br>AEGKCGEGKCGSKHHHHHH | -12.10<br>(+801.34) | 6067.00 (ave.) |
| lipo-NedABC (N-terminal LysC fragment) | CQVEPAQSSSPEVSVQQK | 800.73 | 2731.74 |

**Table S4. Assignment of proton and carbon chemical shifts of LysC-digested DybAH in 90% $D_2O$ and 10% $H_2O$.** Chemical shifts were referenced to $D_2O$ at 4.790 ppm. Br: broad.

| Residue number | Residue | NH<br>CO | alpha | beta | gamma | delta | other |
|---|---|---|---|---|---|---|---|
| 1 | T | -<br>168.2 | 3.90 (C:58.8) | 4.17 (C:66.7) | 1.32 (C:18.8) | | |
| 2 | R | 8.84<br>173.2 | 4.47 (C:53.6) | 1.79, 1.85<br>(C:28.3) | 1.65 (C: 24.3) | 3.22 (C:40.7)<br>ζC=O:157.0 | Side chain NH: 7.20 |
| 3 | T | 8.39<br>171.4 | 4.29 (C:59.5) | 4.15 (C:66.8) | 1.20 (C:18.9) | | |
| 4 | L | 8.38<br>173.9 | 4.36 (C:52.4) | 1.60 (C:39.8) | 1.59 (C:24.2) | 0.86 (C: 20.7),<br>0.92 (C: 22.0) | |
| 5 | D | 8.46<br>170.5 | 4.93 (C:48.5) | 2.79, 2.88<br>(C:36.0) | γC=O<br>174.7 | | |
| 6 | P<br>(trans) | -<br>173.6 | 4.33 (C:61.1) | 1.92, 2.28<br>(C:29.6) | 2.01 (C:29.2) | 3.76, 3.76<br>(C: 47.9) | |
| 7 | C | 8.58<br>N/A | 4.92 (C:53.5) | 2.98 (C:33.3) | | | |
| 8 | P<br>(trans) | -<br>174.4 | 4.43 (C: 62.5) | 2.04, 2.37<br>(C:29.0) | 2.04 (C:24.5) | 3.47, 3.79<br>(C: 47.7) | |
| 9 | A | 8.52<br>172.4 | 4.72 (C:48.2) | 1.34 (C:14.9) | | | |
| 10 | C | 7.90 | 5.02 (br) | 5.80 (br) | | | |

| | | | | | | |
|---|---|---|---|---|---|---|
| | | 157.0 | (C:51.7) | | | |
| 11 | G | -<br>172.0 | 4.42, 4.15 (C:45.5) | | | |
| 12 | M | -<br>170.6 | 4.97 (C:53.5) | 2.43 (C:26.2) | 2.59 (C:30.0) | 2.09 (C: 14.3) |
| 13 | G | 8.42<br>174.4 | 3.93, 3.88 (C:42.4) | | | |

*N/A: not observed.*
*-: No such proton exists.*

**Table S5. Assignment of proton and carbon chemical shifts of LysC-digested, IAA-alkylated DybAH in DMSO-d6.** Chemical shifts were referenced to DMSO-d6 at 2.50 ppm ($^1$H) and 39.5 ppm ($^{13}$C).

| Residue number | Residue | NH<br>CO | alpha | beta | gamma | delta | other |
|---|---|---|---|---|---|---|---|
| 1 | T | 8.09<br>($-NH_3^+$)<br>166.6 | 3.57 (C:58.0) | 3.79 (C: 65.9) | 1.14 (C:19.6) | | Side chain OH: 5.55 |
| 2 | R | 8.66<br>170.6 | 4.49 (C:52.0) | 1.69, 1.53<br>(C:29.1) | 1.53 (C:24.5) | 3.11 (C:40.3) | Side chain NH: 7.44<br>$C\zeta$: 156.7 |
| 3 | T | 8.01<br>169.0 | 4.26 (C:57.7) | 3.97 (C:66.2) | 1.00 (C:19.1) | | Side chain OH: 5.03 |
| 4 | L | 7.84<br>169.2 | 4.31 (C:50.7) | 1.42 (C:40.7) | 1.60 (C:23.8) | 0.82 (C: 21.3)<br>0.86 (C:23.0) | |
| 5 | D | 8.35<br>171.8 | 4.75 (C:47.5) | 2.37, 2.73<br>(C:35.5) | | | |
| 6 | P | -<br>N/A | 4.12 (C:60.2) | 1.86, 1.81<br>(C: 24.2) | 2.04, 1.74<br>(C:29.1) | 3.66, 3.56<br>(C:46.6) | |
| 7 | C | 8.33<br>N/A | 4.82 (C: 51.8) | 2.61, 2.96<br>(C:33.7) | | | |
| 8 | P | -<br>170.7 | 4.24 (C:61.8) | 1.87, 1.82<br>(C:24.2) | 2.18, 1.80<br>(C:28.5) | 3.65, 3.32<br>(C:46.5) | |
| 9 | A | 7.99<br>170.9 | 4.62 (C:46.3) | 1.22 (C:15.5) | | | |
| 10 | C | 7.34<br>N/A | 5.20 (C:57.3) | 5.92 (C:64.5) | | 3.39, 3.16<br>(C:32.8)<br>(IAA: $\alpha$H) | 170.2<br>(IAA:O=C-$NH_2$) |
| 11 | G | -<br>171.0 | 4.27, 4.04<br>(C:44.9) | | | | |
| 12 | M | -<br>167.8 | 4.60 (C:52.9) | 2.37, 2.28<br>(C:26.9) | 2.38<br>(C:29.9) | 2.02 (C:15.2) | |
| 13 | G | 8.42<br>170.9 | 3.82, 3.69<br>(C: 40.8) | | | | |

*N/A: not observed.*
*-: No such proton exists.*

**Table S6. Protein sequences of predicted MNIO-modified RiPP-lipoproteins.**
Sequences are grouped by cluster as shown in the SSN in Fig. 5c.

| Cluster | Annotation | Sequence |
|---|---|---|
| I | Chryseobasin-like | MKVSKSVLQAVAVAVTVTALATACTDSTIGPNGEKTTKTKTVDPCPACGMG |
| I | Chryseobasin-like | MKISKSVLQAVAVAVTVTALATACTDSSVKPNGEKTSKTKTMDSCPACGMG |
| I | Chryseobasin-like | MKISKSVLQAVAVAIVVTTTISACANDDVSPEKEKATKSQKLDPCPACGMG |
| I | Chryseobasin-like | MKLSKPLLQAIAVAVTVTTISSCTKDKVIDPKAPQGEQQREPYNCPACGMG |
| I | Chryseobasin-like | MKLSKSLLQAIAVAVTVTTISSCGKGTVVDPKAPNGEQQKKVPYNCPGCGLG |
| I | Chryseobasin-like | MKLPKAVLGAVLVGLAVQTTSCIKKDEVKPNGEKTGQKGGKTIDSCPACGMG |
| I | Chryseobasin-like | MKISKSVLQSVAVAVALTTLVSTTACVDNIIKPNSEKKLRQKVDPCPGCGMG |
| I | Chryseobasin-like | MKLPQALLGAILLGVVAQATTGCKKDAPSPKQEEGSGKGKPVPINCPACGMG |
| I | Chryseobasin-like | MKLSQAILGAVLVGLTAQTTGCIKKSDPTPKEEQGKSGKKSPEIPYNCPGCGLG |
| I | Chryseobasin-like | MKLSQAVLGAVLVGLTVQTTGCIKKEDPTPKEEQGKPSKKNPKVPDSCPGCGMG |
| I | Chryseobasin-like | MKISKPLLQAMALAVAVTTVGSSCKDGLVKPKKEEAKKEQQRKLPDNCPACGMG |
| I | Chryseobasin-like | MKLSQAVLGAVLVGLTVQTTSCIKKDDPTPKEEQGKPSKKSPKVPDSCPGCGLG |
| I | Chryseobasin-like | MKIPQAVLGALLVGLAAQTTGCIQKNEPVVKQEQAKPKAPAPEPYSDPCPACGMG |
| I | Chryseobasin-like | MKLSPALLQAITLGVAVTALSSSCHQNEVNSGRTKEIKAKKEKPVRDNCPVCGMG |
| I | Chryseobasin-like | MKISQALLGAMLVGLAAPTTSCTKKGDPTPKQEEGKQKKGESTKTPDSCPACGMG |
| I | Chryseobasin-like | MKLPKTLLSAIVVGIAVQTSACQKDDLPKPTAENGGKDGKEVIKNPANCPACGMG |
| I | Chryseobasin-like | MKLSKTLLSALLLGITVQTTTSCEKKELPQPNSGKEESSKEQGKEAPVNCPGCGMG |
| I | Chryseobasin-like | MKIPQAVLGAVLVGLAVQTTGCSSKNDPKPKQTSEQEAKKSGEASKEPINCPACGLG |
| I | Chryseobasin-like | MKISKKLIGAIVVGIAIQVSTTSCNKKEIKPQKAPNAENGQPVNHDPVLDCPACGMG |
| I | Chryseobasin-like | MNLPKALLSAILLGITVHTTTSCTDKNELPGPKTEKEKTSKTPEKNDPGNCPGCGMG |
| I | Chryseobasin-like | MKLSKALLAAMLTGITLQGVQSCTKDKDEPSKEQKEKEKEGENKPKPHPYSCPACGMG |
| I | Chryseobasin-like | MKLPKSLLSALVIGIAVQAVPACTKTKESPSQKAKKDAKEKETEKTSIPFNCPACGMG |
| I | Chryseobasin-like | MKLSKSLLSAIMIGIAVQTTVVSCSKDEQVKPKKADQANKQSESKPVDNPDSCPACGMG |
| I | Chryseobasin-like | MKLSKTLLAALLTGITLQGLESCTKDKDGPSKEEKEKEEEGKGKPKPPTVPYNCPACGQG |
| I | Chryseobasin-like | MKVSKSLLQAIALGVALGSSATSCMLLDNSDEVKPDTEKAAEDDRCPTDDGDKPWYNCPACGMG |
| I | Chryseobasin-like | MKLSKSLLGAILVGVTVQTAVTSCNKKSNETFKPTSEAQANPESQANPTNQNPNSVPSECPACGMG |
| I | Chryseobasin-like | MKIPKTVLQAVAAAVVITTVTACSVSDVNPEGEKVTKNKIVDSCPGCGMG |
| I | Chryseobasin-like | MKLSKALLQTIAVAVAVTTLSSCDKGKVIDPKKDDTKQQIPYDCPGCGLG |
| I | Chryseobasin-like | MKISKSVLQAVAVAVALSTITACADIDIKPGKEKKAKTRTMDPCPGCGMG |
| I | Chryseobasin-like | MKVPASVLKAMAAAVIVASVTACTGEDIGPKNGKKKEEKTLASCPACGMG |
| I | Chryseobasin-like | MKISKPVLQAVAIAVALSTITACADSDIKPEKEKKTNTKTMGSCPACGMG |
| I | Chryseobasin-like | MKISKSVIQAVAAAVIVSSVVSCTSGLINPEKEKGSKTKNVDPCPGCGMG |
| I | Chryseobasin-like | MKIQKNTLQAIAVAVAVTAVTAACAVDNVGPKEKQAKTKVIDNCPACGLG |
| I | Chryseobasin-like | MKISKSVLQAVAVAVTVAALTTACTDNAVGPNGEKTSKTKTLDNCPACGMG |
| II | Bufferin-like | MKRTSIKIGLTLAATAAALFATGCTTLCGESQEALVKCSNVNACKGSSECATPSNSCKGRNSCKSSGW VYMTKSECLANGGSVRD |
| II | Bufferin-like | MNYKRSLLVGAAIAGLSLSACTTTNTSNTGSKVEGECHGINACKGQGACGGKGSSCAGTNACKGKG WLKTAENECVSKGGKFVKG |
| II | Bufferin-like | MKKSGLFFTAAVGGLMFAATCSPQAEKPKAEAVKGECHGINSCKGQGACGGKDHSCAGQNACKGQ GWLKMTEEECNAQKGTFKPLSMSM |
| II | Bufferin-like | MGLKTKGTGIALAVSASMLIAGCASNQQQADASSHGSADGMVQCHGVNSCKGTSSCKAEGSSCAGK NACKGKGWLPMSQSECESHGGRMGGFNMN |
| II | Bufferin-like | MKTASKVTGFTLASAAAALLLAGCSTGGDSAKSASMSKDKKMADVKCAGINSCKGTSACATATSACK GLNSCKGQGWVKASATDCADKGGKVLGQA |
| II | Bufferin-like | MSAFALSAAALFLGACDKTKSGEGSQAPAEQPPAASSQAKDAKVKCFGVNECAGQSVCAVNKPELGI EHACAGENECKGKGWIKLSSAECDDLGGEIL |
| II | Bufferin-like | MTNTNKQLMSAFALSAAALFLGACDKSSGGTTDPGDAAGGDANADEVVKCFGINECKGESACNVNK PELGIEHSCAGENDCKGKGWIKVPRSECDAKSGEVL |
| II | Bufferin-like | MTKNSKQFMSAFALSAAALFLGGCKKTTTEVTSEPTATPGPAVTADEQVKCFGINSCTGESACAVNKP DLGIEHACAGENACEGKGWIKVSRSDCEEKSGEVLGTL |
| II | Bufferin-like | MTRNNKQFVSAFALSAAALFLGGCKKTEPTAEPTAEPTATAPASNADEDVKCFGINSCSGESVCAVNK PDLGIEHACAGENACEGKGWIKVSRSECETQSGEVLGTL |
| II | Bufferin-like | MSIETSKARRGAMLALGAAALILAGCAATGAGTNTAANSTVGACVGVNSCKGTSDCKTAGNSCKGQN ACKTATNACKGQGSCKSAANSCKGQNECKGQGHLALTGADCDAKGGHLV |
| II | Bufferin-like | MGLKTKKTGIALAVTASMLIAGCASNQEQASAKGHGHGHGHSHGADNMGQCHGANSCKGKGSCGA AIADSCKGTDSCKAKKDNSCKGKAACKTKKDNSCKGKSGCKAKAEAHSCAGMNSCKGKGWLPMSQ SDCDAKGGRFKGAK |
| II | Bufferin-like | MKALTGAAMAMMVAGLVGCNSTENNSTSAQAAAAETDLVHCYDVNVCGGHNDCKTASNACSGQAS CKGTGFVAMPSKACSDVGGNQKDAWVGSVAKADLVHCHDVNVCGGHNDCKTASNACAGHASCKG TGFVSMPAKACADIGGKVKS |
| II | Bufferin-like | MKKFTGAAMAVMVAGLVGCNSTEADTSAASVANMNSTTDLVHCYDVNVCGGHNDCKTADNACSGQ ASCKGTGFVAMPSKACGDVGGAIKDKWVGTVAQTDLVHCNDVNVCGGHNDCKTADNACAGQASCK GTGFVSMPAKSCGDIGGSVDS |

| | | |
|---|---|---|
| II | Bufferin-like | MNKNTKNVMSGLAVAFAAAAIAGCSATANTNTTAAAANTTDLVHCAGVNVCKGHNDCGGASNACAGQGSCKGSGFVVMPSKACGDVGGKVQDDWRGEVSKAELVQCTGVNVCKGHNDCSGADNSCAGQASCKGTGFVKMGAKACGDIGGKS |
| II | Bufferin-like | MKKFTGAAMAVMVAGLVGCNATETQTSSSNTVANNTATTDLVHCYDANVCGGHNDCKTAENACAGQASCKGTGFVAMPAKACADVGGKVKDKWVGTVTKTDLVHCNDVNVCGGHNDCKTANNACAGHASCKGTGFVSMPAKACGDIGGSVDS |
| II | Bufferin-like | MKALTGAAMAVMVAGLVGCNSTDNTSSNTANANTSGTSTDLVHCYDVNVCGGHNDCKTASNACSGQASCKGTGFVAMPAKACADVGGKVKDKWVGSVAKSDLVKCNDVNICGGHNDCKTANNACGGHASCKGTGFVKMPAKACQDIGGKVDS |
| II | Bufferin-like | MKQTKLNLSGVAIAMTAASLISGCVGPSKSTHSSTAAAGKTDLAHCYGVNICGGHNDCKTANNSCKGQASCKGQGFVAMPTKACNDVGGKVKDDWRGQVSKADLVKCYDVNLCKGHNDCKTANNACAGQASCKGQGFVLTPQKSCEDIGGKVG |
| II | Bufferin-like | MITNMKTKLSGAAIAMAAAGLAGCMGANAANSSTAAPTDTVELGHCYGINSCKGHNDCKTANNACKGMGSCKGQGFVTMPTKACTDAGGKVRDDWRDQIAKAELTQCFGVNVCKGHNDCKTADNACAGHGSCKGTGFVLTTAKSCADIGGTTK |
| II | Bufferin-like | MSQHQRLLRGAGLAVAAAGLAACGHTQTTAQTSSAAAEKTDLIHCYGVNVCKGHNDCKTASNACAGHGSCKAQGFVAMPTKSCEDAGGKASREWVGEIARADLIHCYGVNICKGHNDCKTKENACAGHATCKGTGFIEAPAKSCKDIGGTVGK |
| II | Bufferin-like | MKALTGAAMAMMVAGLVGCNSTSESGATSGASAANAGAAKTDLVHCYDVNVCGGHNDCKTASNACAGQASCKGTGFVGMPSKACADVGGKQKDDWVGSIAKADLVHCHDVNICGGHNDCKTASNACAAQASCKGTGFVNMPAKACKDIGGKVGS |
| II | Bufferin-like | MKALTGAAMAMMVAGLVGCNSTSDSGATSNASSANAGGAKTDLVHCYDVNVCGGHNDCKTASNACAGHASCKGTGFVGMPSKACADVGGKRKDDWVGSIAKADLVHCHDVNICGGHNDCKTASNACAAQASCKGTGFVNMPAKACKDIGGKVGS |
| II | Bufferin-like | MKALTGAAMAVMVAGLVGCNATDSSQSSTASATASSGGAKTDLVHCYDVNICGGHNDCKTASNACAGQASCKGSGFVGMPAKACADVGGKQKDSWVGSVAKADLVHCHDVNICGGHNDCKTANNACAAQASCKGTGFVNMPAKACKDIGGKVGS |
| II | Bufferin-like | MKALTGAAMAVMVAGLVGCNATDSSQSSSAGATASSGGAKTDLVHCYDVNICGGHNDCKTASNACAGQASCKGSGFVGMPAKACADVGGKQKDDWVGSVAKADLVHCHDVNICGGHNDCKTANNACAAQASCKGTGFVNMPAKACKDIGGKVGS |
| II | Bufferin-like | MKALTGAAMAMMVAGLVGCNATDSSQSSNASTAASSKGATTDLVHCYDVNICGGHNDCKTANNACAGHASCKGSGFVGMPAKACADVGGKQKDSWVGSIAKADLVHCHDVNICGGHNDCKTANNACAAQASCKGTGFVNMPAKACKDIGGKVGS |
| II | Bufferin-like | MNTKAQATLTGAALALAMAGLTGCNSTDKSETTQAAMAKGSSDLVHCYGVNACKGHNDCKTATNACGGHASCKGTGFVAMPAKSCGDVGGKVQDDWVGSIAKTDLVHCYSVNVCKGHNDCKTANNACGGHASCKGTGFVAAPAKACADIGGKVGA |
| II | Bufferin-like | MKKVSQKITGAAIAVAAATLMGCTTTHSGNSASTATAQGGTTDLVHCYSVNQCKGHNDCKTAENACAGHAECKGHGFVAMPSKACDDIGGKIKDSYRGTIAESELVHCYGVNQCKGHNDCKTAENACAGHAECKGHGFVAMPAKSCGDVGGKEGA |
| II | Bufferin-like | MNTKAQATLTGAALALTMAGLTGCASTEKSETTQAAMAKGSTDLVHCYGVNVCKGHNDCKTATNACGGHASCKGTGFVAMPSKSCADVGGKIQDDWVGSIAKTDLVHCYSVNVCKGHNDCKTANNACGGHASCKGTGFVSAPAKSCADIGGKVGA |
| II | Bufferin-like | MNSNLKSSFSGAAIAVAAAGLAGCMGSTSASTSSAPMASGNTVELGHCYGVNSCQGHNDCKTADNDCKGQSACKGQGFVTMPLKACTDAGGNVRDDWRDQIAKADLTQCFGINVCKGHNDCKTADNACAGQGSCKGTGFVMTTEKSCDDIGGTTK |
| II | Bufferin-like | MKPNTKSALQGTAVALAVAGMMGCAQTSSQSGSSSYAAKNTAELGHCYGVNKCSGHNDCKTADNACKGQASCKGHGFVAMPTKSCGDIGGTVKDDWRGKVTTADMAHCYNVNVCGGHNDCKTANNACKGQASCKGQGFVKMSPKACNDVGGKVGK |
| II | Bufferin-like | MKALTGAAMAMMVAGLVGCNSTSTSNSSAGSATASAAASASSTDLVHCYDVNVCGGHNDCKTASNACAGQASCKGTGFVAMPKKACGDVGGKTKDAWVGEIAKAALVHCHDVNICGGHNDCKTASNACAGQASCKGTGFVSMPAKSCGDIGGKVKS |
| II | Bufferin-like | MKRTSKTALALAAAAAALFATGCSTMCGDSHEALVKCAGSNACKGSSECATPDNSCKARNSCKATGWSYMTKGDCSAAGGSVVD |
| III | Oxazolin-like | MNVKTLAALVGTLSLSALATGCASTKSAEAQPAAAEKGAEGSGAATTEGQAAEGAQTQEGAAPAATPEKGSEMSCGSGSCGGGKK |
| III | Oxazolin-like | MNVKHIAATLAILGSASFAVTGCKKNAEGTEVPAEGEKKADGSCGANKDGSCGANKDGSCSAKKEGEGDVPAAGDAPAEATPPPAN |
| III | Oxazolin-like | MNVKALAAVVGTLSLTALATGCASSKSAEATPASTDTSTQSPGSQAANPEQGAAGDTTAAPSAPKGEEGKCGEGKCGEGKCGEGKCGGKK |
| III | Oxazolin-like | MNVKALAAVVGTLSLSALATGCASSKSAEAAPASGDTSAPSPGSQAAAPEQGAAGDTTAAPSAPKGQEGTCGEGKCGEGKCGEGKCGGKKE |
| III | Oxazolin-like | MQKRNLSLALSVGAILVAGSTLSACNHTPVKSDDGASTAAKPADGKCGAKKDGSCGAKKDGSCGANKDGSCGAKKDATCGAKKDGSCGAKQ |
| III | Oxazolin-like | MNVKTLAAIVGTLSLGSLAAGCATSSGSAQPGTEKGAQGSCSATGGEASCSGKATQKGGEASCGEGTCSGAPKAATPEKGTHASCGESGCGGGR |
| III | Oxazolin-like | MANLKDIVATLAILGAGSLALTGCKKEAETANPDEATEATEATDAEATDAEASCAGEGEADAEASCAGEGEADAEASCAGEGEGEADAEAATEAV |
| III | Oxazolin-like | MANLKDIVATLAILSAGTLTLTGCKKEGGETTNPDEVTADDAAGEGEGEGEGEASCSGEAEGEASCSGEDAAAEGDAEGDAEGEAADAEGEASCSGAA |

| | | |
|---|---|---|
| III | Oxazolin-like | MNVKTLAALVGTLSLGSLAAGCATTHRSAEPGTEKGTQGAGSTRGGEASCGEGTCSGKATQKGGEASCGEGTCGGDQKAAAPEKGSHASCGESGCGGGR |
| III | Oxazolin-like | MANLKDIVATLAILSAGSLALTGCGDKKDTTNPTDAAGEGGDVAAPAEGEGSCKGEDGEGSCKGDADGEGSCKGDAAEGEGSCKGDAAEGEGSCKGDAAEGEEAAAE |
| III | Oxazolin-like | MNVKALAAVVGTLSLGALATGCASNKAAEGSKVHAASPGAPEASESSAAPAAEGAAPAAAPEKGAEGSCGAGSCGAGSCSGKK |
| III | Oxazolin-like | MNAKALAAIVGTLSLASLATGCASTKAAEGQPAAAEKGAEGSCGAAKTEENKPAEGTDAAAPAATPEKGAEHSCGAGGCNGQKK |
| III | Oxazolin-like | MNVKALAAIVGTLSLGALATGCASNKAAEGSTEGAATTESSAKGTEASCKGGTEAKGTEASCKGATEAKGTNASCGEGTCANKK |
| III | Oxazolin-like | MNVKTLTALVGTLSLGALATGCATAKPAAAPSEQGSATVNSAAATPEKAAEGTCSGKASPTTEKAAEHGCGAGGCGASGCGGKK |
| III | Oxazolin-like | MNVKTMAAIVGTLSLGSLATGCATAKPAAATSEAKGTEASCKGGAEATQEKGGEAKCSGKAASTSEKGSEHGCGKDGCGAAKPQ |
| III | Oxazolin-like | MNLKTLTALVGSLSLGTLATGCATAKTAAAPAEQESTSAHAASAVPAESSTAPAAEGTATPSTEKSSEHSCGSGGCGANGCGGKK |
| IV | Oxazolins | MEKKSKSILWSSVAAGAMLGAASIFASCSTESKDVTTEVQTPAKDTTGEHTCGTEHTCGKDSAEAKCGEGKAADAKCGEHKTTEGKCGEGKCGADKKK |
| IV | Oxazolins | MKTTSSLIALAGFLALAGCQGQSNGSAESTPAASVSAPKAEEGKCGEGKCGASHAGEKAGEGKCGEGKCGASHAGEKAGEGKCGEGKCGTSHAGEKAGEGKCGEGKCGSH |
| IV | Oxazolins | MSNLNRKITGLAAATLLATGLSACKEEKMQAQPLAQGYQNADTAQPAVAQPNQAVVSDTKTAEGKCGEGKCGEAGCSATMGDKSASASCGASTANADKSASAACGASTDKNANASCGGSK |
| IV | Oxazolins | MQNTISQSSQNKKHLAKNLALSSLLAVSVVGCAESQMGAKPMEQGYQNNQNPPQASQENMSQASMSQENMSQASTTVNEQQSNLFKAPEAKCSEGMCGASHKDSSGGCGASHKDASGGCGASH |
| IV | Oxazolins | MKTTSSLIALAGLLALAGCQDKNQQPAPTAAAASAAASKAEEGKCGEGKCGAASHAESKAEEGKCGEGKCGAASHAHSKAEEGKCGEGKCGAAAKAGSKAEEGKCGEGKCGAGSKADAKSAEGKCGEGKCGSK |
| IV | Oxazolins | MTKTSSSLIALAGLLALAGCQGQSNSEPVQATSAASSSTSKAEEGKCGEGKCGAASQAAGKSAEGKCGEGKCGAASQAAGKSAEGKCGEGKCGAASQAADKSAEGKCGEGKCGAASQAAGKSAEGKCGEGKCGSK |
| IV | Oxazolins | MSNLNRKITGLAVATLLATGLTACKEEKVEAQPMEQGYQNAEPAQPATAEPNTATATKTAEGKCGEGKCGEAGCSATMTDKSAEGKCGEGKCGEGKCGAEAKAGDKSANAGCGANTATATADKSATASCGASTDKNANASCGGNK |
| IV | Oxazolins | MKTTSTLMALAGALVLAGCQVEPAQSSSPEVSVQQKAGEGKCGAASQAGKAAEGKCGEGKCGSK |
| IV | Oxazolins | MKISSSLIALAGLLALAGCQQEAVSGKPAVEAASHADKSAEGKCGEGKCGASHSKNAKSAEGKCGEGKCGSK |
| V | BON-domain containing protein | MKKYYLNGLIGFCLAILVGCQSLTHENFFRVGPSDETITTSVQSAFSNNPDLSAQRIHVETHKGTVVLSGYVKTIRQSDTAGDIAGKISGVKSVQNELIVRK |
| V | BON-domain containing protein | MKNYYLKGIIGFFFVLLVGCQSLTNENFFRIGHSDASITASVNSAFASHQDLSTQRIHVETQKGTVVLSGYVKTIRQSDTAGDIAGKIPGVKSVQNGLIVRK |
| V | BON-domain containing protein | MRKHVQIVLLALIGLILVGCSFTNGRPVFTPRLSDDAITTSVHQAFVNNRILSDAPVHVETHQGNVVLSGYVKTIRQSDTAADVAAKVPGVKFVQNNIIVRK |
| V | BON-domain containing protein | MMRRYIQVVLVGALFVLIASCQMISSPSIFNPRLSDQAITNSVYQAFADSKILSDVPVHVETHQGNVLLSGYVKTIRQSDTAQDVATKVPGVRGVQNNIIVRK |
| V | BON-domain containing protein | MRRYCCIALISISLIGMAGCQMLTSGNLFAPRFSDDAITSSVNQAMMNNINLAGIPIHVETHQGNVMLSGYVKTIRQSDTAGDVASKVPGVKSVQNNLIVRKW |
| V | BON-domain containing protein | MRRYGRMVLISLGFFAMTGCQVLTGGNVFSPRLSDDTITTSVSQAMMNNQNLINVPIHVETHQGNVILSGYVKTIRQSDTAGEVASKVHGVKTVQNNLIVRKW |
| V | BON-domain containing protein | MMRKYCRIALMCISFMAMSGCQMLSGGNIFAPRLSDEAITSSVKQAMMNNNNLMNVPIQVETHQGNVMLSGYVKTIRQSDTAGDVASKVPGVKSVQNNLIVRKW |
| V | BON-domain containing protein | MKRISERIPVVIMLTLVMLLAGGCAGNGETRSTGQYVDDAALTSKVKTALFRDDDVSGFQVDVDSFKGRVQLSGFVDSEEQKLRAEQVARGIEGVQEVTNNLEVK |
| V | BON-domain containing protein | MLKYLLSALISVFLVLVSGCQTHTTAGSGLFSPTYPSGMTLAQSVQEALIQSNDPVIAQVRVETNQNTVILSGYVKKIRQSDIAEQIARNVQGVQMVQNNIIVRP |
| V | BON-domain containing protein | MARENTAALLLAAIMASTVVACSSTPTQQSTGQAIDDGVVTAKVKAKLIEDPVTKAHQINVETFKGAVQLSGFVETDQARTRALQLARDTDGVKSVKDALEIRKGG |
| V | BON-domain containing protein | MRILLTACALMLLSGCTALMVGGAAAGGYQLGKDERSASQVTRDGATTASIKSRLIADKLVSAFNVNVDTYENRVTLRGTVGSYAARARAAAIANEVDAVVAVDNQLKVVNSR |
| V | BON-domain containing protein | MNCIRRVFLVLITVTTTIAISGCATAVMTGAAQGGYDPDDGRSARQAPADAEISAAVTRALVHHPQIPAMPITVRTERGRVTLSGKVPDRDTARRAARLAAAVPGVESVRNLLRVEQR |
| V | BON-domain containing protein | MRFPQLSFAASAGVLAAIGATTLLTACAWGDSSTASGARMLDDSLLSYQVKAALDQDTALSTRQIRIRSNPDGKVTLTGWVDTPEMARRAGEDVKRFVDPAKLDNQLRVLSRMQVLGGGPYIPAGLPPEAPASAPAAR |
| V | BON-domain containing protein | MRFPQLPFAAPAGALAAIGAAVLLTACAWGDSPNSTGARMTDDSLLAYQVKAALDQDTALNPRQIRIHTTPEGKVTLTGWVDTPEMARRAGDDVKRFVDPAKFDNQLRVLSRSQVLGGGPLIPAGLPPEAPASAPAAR |

| | | |
|---|---|---|
| VI | LPS translocon maturation chaperone LptM | MMRLPLKLAMLATVALLSACGQQGPLYFPKDAPAQNEPVNPETSQATQASTQP |
| VI | LPS translocon maturation chaperone LptM | MMRLTLKLAALATVALISACGQQGPLYFPEDAPAQNEQVNPETPQDTQESTQP |
| VI | LPS translocon maturation chaperone LptM | MRTILNASLLMAIVTLSGCGQRGPLYFPEDAPATQGSPADNSQQDTAPDSTKTQ |
| VI | LPS translocon maturation chaperone LptM | MRMTLKLAVLSAAILLSACGQQGPLYFPEDAPAQNTAPAQTETAHDTQAQDPAQQ |
| VI | LPS translocon maturation chaperone LptM | MRTTLKLAVLSATVLLSACGQQGPLYFPEDAPAQNAAPAQTEIANDTQAQDPAQQ |
| VI | LPS translocon maturation chaperone LptM | MMRTTLKLFVLAAVTVLSACGQQGPLYFPEDAPAQNSAPATSDTTQDTQPQAASQP |
| VI | LPS translocon maturation chaperone LptM | MMRTTLKLVVLAAITLLSACGQQGPLYFPEDAPAQNSAPAETGNTQDTQAQGTDQP |
| VI | LPS translocon maturation chaperone LptM | MLHSKLSSQTVILIVKSTLALQFTALACTAALLGGCGQQGPLYLPKQPAKPASANTQGKPGVAPPSAPV PSTPPASQQSPVN |
| VII | Glycine zipper two transmembrane domain-containing protein | MRKSVLLAACFTTLSLLLGGCASSLTGDSYSRDEARRVQTVRMGTIESLRPVKIEGTKTPIGGAAGAVI GGVGGSAIGGGRGSIVTAVIGAVAGGLLGSATEEGLTRTQGVEITVREDDGSTRAYVQAVQENEIFRIG DRVRIMTVDGTSRVTR |
| VII | Glycine zipper two transmembrane domain-containing protein | MRKSVLLIAAFSTMTVLLGGCTSNLTGDSYSRDEARRVQTVRMGTIVALRPVQIEGTKTPIGAATGAIV GGVGGSAIGGGRGSIVTAVIGAVAGGLLGSAAESGLTKTQGVEITVREDDGSTRAYVQQVEPNQVFRT GERVRIMTVDGTSRVTQ |
| VII | Glycine zipper two transmembrane domain-containing protein | MRKSVLLVACFTTLSLLLGGCASSLTGDSYSRDEARRVQTVRMGTIESLRPVKIEGTKTPIGGAAGAVI GGVGGSAIGGGRGSIVTAVIGAVAGGLLGSATEEGLTRTQGVEITVREDDGSMRAYVQAVQENEIFRI GDRVRIMTVDGTSRVTR |
| VII | Glycine zipper two transmembrane domain-containing protein | MRKSVLLVACFTTLSLLLGGCASSLTGDSYSRDEARRVQTVRMGTIESLRPVKIEGTKTPIGGAAGAVI GGVGGSAIGGGRGSIVTAVIGAVAGGLLGSATEEGLTRTQGVEITVREDDGSTRAYVQAVQENEIFRIG DRVRIMTVDGTSRVTR |
| VII | Glycine zipper two transmembrane domain-containing protein | MRKSVLLVACFTTMSLLLGGCASSLTGDSYSRDEARRVQSVRMGTIESLRPVKIEGTKTPIGGAAGAVI GGVGGSAIGGGRGSIVTAVIGAVAGGLLGSATEEGLTRTQGVEITVREDDGSMRAYVQAVQENEIFRI GDRVRIMTVDGTSRVTR |
| VII | Glycine zipper two transmembrane domain-containing protein | MRKSVLLVASFSTMAMLLTGCQSNLSGDSYSRDEARRVQTVRMGTIESLRPVKIEGTKTPIGGLAGAA VGGVGGSAIGGGKGSIVAAVIGAVAGGLIGSATEEGLTRTQGVEITVREDDGSMRAYVQQVQENEVFR VGERVRIMSVNGTSRVAH |
| VII | Glycine zipper two transmembrane domain-containing protein | MRKSVLLVASFSTMAMLLTGCQSSLTGDSYSRDEARRVQTIRMGTIEALRPVKIEGTKTPIGGAAGAVV GGVGGSAIGGGRGSIVAAVIGAVAGGLIGSATEEGLTRTQGVEITVREDDGSMRAYVQQVQENEVFR VGERVRIATVDGTSRVSH |
| VII | Glycine zipper two transmembrane domain-containing protein | MRKSVLLVASFSTMAMLLTGCQSSLTGDSYSRDEARRVQTIRMGTIEALRPVKIEGTKTPIGSIAGAAV GGVGASSIGQGNGSIVAGIIGAVAGGLIGSATEEGLTRTQGVEITVREDDGSMRAYVQQVQENEVFRV GERVRIASVDGTSRVSH |
| VII | Glycine zipper two transmembrane domain-containing protein | MRKSVLLVASFTAMATLLGGCASNLTGDSYSRDEARRVQTVRMGTIESLRPVKIEGTKTPIGGAAGAVI GGVGGSAIGGGRGSIVTAVIGAVAGGLLGSATEEGLTRTQGVEITVREDDGSMRAYVQQVQENEVFRI GDRVRIMTVNGTSRVTR |
| VIII | TssQ family T6SS-associated lipoprotein | MLVSALAAAVLAAGCAQTPPPPVGLLDVTSRPAERALQGGIRAYEDGQYPEAEKQLNLALTTGLVSPR DRAAAHKHLAFIYCTSSRTSDCEAAFRAARQADPAFALNKSEQGHPAWGPVYKRVQP |
| VIII | TssQ family T6SS-associated lipoprotein | MRWPALALVVLSISGCALPPAPSPGLMDVAERPAEKALLGGMRAYDDGQYVQAEAQLKQALATGLAA PRDRAAARKYLAFIYCTSQRVPACEAGFRAARRDDPAFALTRAEAGHPLWGPVWLKSRD |
| VIII | TssQ family T6SS-associated lipoprotein | MNRLTAISALFALACAGCATPPAAPPAGGLSDLLERPAERALYEGMRAYDDGQYPAAEKSLRRALGE GLASPRDRANAYKLIAFIACTSERLGECEAAFMAARTVYPAFVLSRSEAGHPLWGPVYRRVAP |

| | | |
|---|---|---|
| VIII | TssQ family T6SS-associated lipoprotein | MRTLLGTTLASLVLATLAGCVTQPKQAPIGLLDVVSRPAEKALQAGLRAYDDAQYGEAEKQLNNALKVGLVSPRDQAEAHKVLAFIYCTSRRPVECEAEFRAAKASDPAFALSKSEQGHPLWGPIYKKIP |
| VIII | TssQ family T6SS-associated lipoprotein | MTKTLLLAMSATALVLIAGCVQPPPAPPVGLLDVTSRPAERALLAGIRAYEDGQYPESEKQLNAALQSGLVSPKDAAAAQKHLAFIYCTSNRTPQCEGAFREARKADPAFALSRSEQGHPSWGPVYKRMSP |
| VIII | TssQ family T6SS-associated lipoprotein | MRLPPVATLVSTFAGMLLAAGCAQTPPAPVGLLDVTSRPAERALQGGIRAYEDGQYPEAEKQLNLALTTGLVSPRDRAAAHKQLAFIYCTSNRTSDCEAAFRAARQADPAFALNKSEQGHPVWGPVYKRVQP |
| IX | Hypothetical protein | MKLVTSVLMGLLLLGANGCAELKEVGTVIGHTTRDATKAIGHASRDAVNSIKEDMSDEE |
| IX | Hypothetical protein | MLFNKILLGAGLAVVMSGCAELKEAGTAIGHSTRDTTTAIGHASRDAVNSVKEDLSSED |
| IX | Hypothetical protein | MKRVLLVAGLLIVVAGCAELKQAGTEVGHATRDVTTAIGHATRDTTKAIGHASRDAVNSVKDDLSED |
| IX | Hypothetical protein | MLKLGKMLILSVLLGASGCAELKQAGTDIGHATRDATTAIGHATRDATKAIGHASRDAVNSVNDDLNSDG |
| IX | Hypothetical protein | MTQCQKLSFSLFTTLLLLGCAELKQAGKEVGHATRDATTAIGHATRDTTKAIGHASRDAVTNISNDLSKE |
| IX | Hypothetical protein | MLKLGKMFVLSALVLLGASGCAELKQAGTEIGHATRDAATTIGHATRDTTKAIGHASRDAVNSVKDDLSDGN |
| IX | Hypothetical protein | MQTLTKSMVLTGMMLFTISGCAELKQAGSDIGHATKDATTAIGHAARDTTKAIGHASRDAVNSVKDDLSSED |
| IX | Hypothetical protein | MSKFTQALWLTTGILLGVTGCAELKEAGRTVGHATRDATTAIGHATRDTTKAIGHASRDAVNSVKDDLSGDSSD |
| X | Likely uncharacterized RiPP precursor | MLSKKSIYETLTALGAGAMISACGGSQTPVNATEVPAAQDATPAAEDPATPADVTPDETPATAPPSEPGAAAAPVDPKPAETAQAPATPATAKPAPKPAAKKPAAKKPDASAACGAGTCA |
| X | Likely uncharacterized RiPP precursor | MNAKSIYETLAALGASAMIAGCAGTQEPVKAAEEPQAEAPAEAAPAEKTEEAAGTEAAGATNAAAAPAEAPKEAAAPAATTTPAPAAAPAPAATPAPAKAAAPAPKPAAKKAAAPKKAGAEAACGEGSCA |
| X | Likely uncharacterized RiPP precursor | MLSKKSIYETLAALSAGAMVVSGCGGAQAPVNAAEVPAEHETAPAAPTEATSGDDAAGATTPATPAEPSSTTEAPTPAPPTDVQSAATAPQVAPTATPAPAAPAPPSPTTPAKPTAKKPAGKKPGHAGCGAGTCG |
| X | Likely uncharacterized RiPP precursor | MLSKKSIYETLAALSAGALLMGGCGGAQAPVNATEVPAEQEATPAAPAAPAEPAPAETTSGDAAAGANVPAAPAAPDSAAPPSDAQSAATSPQVAPTATPAATAPTPPTPTAAPKPAAKKAAAKKPGHAGCGQGTCA |
| X | Likely uncharacterized RiPP precursor | MLSKKSIYETLAALSAGAMVMSGCAGAQAPVNASEVPAEQTAPAPAEPAPAPGPTEAASGDAAAGANVPAAPAAPAPPSDAQSAATSPQVAPTATPAATAAPPAPPTAAPATAAPAKPAAKKTGAAAKKPGHAGCGQGTCG |
| X | Likely uncharacterized RiPP precursor | MLSKKSIYETLAALGTGAMVMSGCGSAQAPVNAAEVPAAQETPPAAPAEATSGDAAAGEGAPAAPAPAAPAAAAPAPAAPAAPAAPASADTPTPPADAPSALTAPQVAPTATPAPAAPGPAPTSAPAKPAAKKPAAKKPAHAGCGQGTCG |
| Other | - | MKKILFLFLLAGLFYACKKDSIGTKPILSFKSYSIDSVISSTQQMVLTMNVEDGDGDIEDSIWIGPVFKSNGPNADTFYSVKKMGDIGANKGNKVKAEVQILLRSIEFKLVQNTGVDSIHFVVFVRDNAGHFSDTISTPKIPYNY |
| Other | - | MKKFLFLFLVVALAASCGKDSYGPVPILSFEGYSVPSIDSNTTTFEAIFRVKDGDGDIDSSIFYTIHYYIPSTLEEQANARMPNIGQNTGKSVNAQVKVTLEAIDFVRWVEHTGTRPDSLWMEVFVQDRAGHISDTIQTTKIPIYKRQ |
| Other | - | MKKILYLLLAVGLLYACKKDNVGTKPLLFFKSYQPDSITPDTRQFVLTMRVEDGDGDIEDSIGVAMLKDSEQAVNKDTIWQFYKMPKIGQNRGNSIKADVIMPFEEIDFAAAYNPAPGDSAHYIVFLRDNSGNISDTVPTPKFPFRRNR |
| Other | - | MKKILFLILAAGLFYACKKDKIGTKPLLSFKSYSEDSITPTTKTFVLTMRVEDGDGDIEDSIAVAMFKDSQQATSHDTIWTFYKMPNIGQNRGNKVKADVIMPFQEIDFAAAYNPVPNDSAHYIVFLRDNAGNFSDTIPTPKFPFRRNR |
| Other | - | MKKFLLLFLFAAFVVLSCNKDNYGPIPTISFESWSLPSIDSNTAVVNAFFRVKDGDGDIDSAIFYRIHYFIANPATDTARFGMKHMAGIGENTGKSVNAQVKLPLEKIDFVRWIEHTEDRPDSMWMEVFIQDRAGNISDTITTKKIPIFKKRAP |
| Other | - | MKKILMFAAVCSVLAACKKESSGTKPSLSFASYDPSVYFASTAAGVPQNGYNFNVTFNIADADGDINDTLGIRAHYKSKDINNTLPDTTAWIYAQMPDIGANKGHSVKGQVTIALQSIDIIGFNPTSTNDSIWFSAYVRDAAGHFSDTVLTAKTAIIKE |
| Other | - | MKGKLTVLSVLLSAGLAAGCQGMNQQETASDSKLQQELAGAMDKQDFRLYYTTGRRPVVPGFEQFEFKALEARCGVKAMPGSGDTLRSEADKAARAEAYQYAKAYNLKIYDACLNRL |
| Other | - | MKDKITAVGALLSIGLTMGCQGMHKQEDADVDDKLQQALTEAMDKQDFRLFYTTGRRPVVPGFEQFAFKDLEARCGVKVMPGTGDTLRSEADKAARSEAYQYAKSYNLKIYDACLNKL |
| Other | - | MKVLMGFCLLCCLTTGCANDYVQRQTDSPVNDNTTEVSEAVALALKDKDYRLFATTGRRPIFPGLEKMSFSTLKAKCGVKYLSGAGDVLKSEQDKQQRFERYEYAKAYNIEMYAKCKAIK |
| Other | - | MKGFIGIALFCCLAVGCVNDYAQRQTDSSDPVRDNTTKVSQAVASALRDKDYRLFATTGRRPVFPGLEKLAFSMLKAKCGVKYLSGSGDVLKSEQDKQQRLERYEYAKAYNIEMYAKCKALK |
| Other | - | MKTYIGLMLSAWLLAGCADGYGEHQNESTTDMTLQKDVSDAIARAVENKDFRLYATTGRRPVFPGLEQLDFDELKVSCGLKYLPSTGDVIKSEQDKQQRLKQYEFAKSYNLKIYAKCKKNAKA |

| | | |
|---|---|---|
| Other | - | MKWIQILACLGLMIGCAHEQPQGSKQSNMNKRSDDLNVDVAIEKALENKDYRLLLTQGRRPVSPGLEHIPIEELKDRCGTKFLTGMGDVIKSTAEKKARVAKYNFAKAYNLKMYAICQKATDKYNKV |
| Other | - | MYFKLSGSKLLTAAALVIAFTVSSCGLIPSQVESSINAVNVDESSLAIKVYDAVAYFTQAKPVLGSSQFTAEHLAATYYFSSAPHQSLFKGDPNKYAAQYGRKLLTNRSQNYKFLVGINSMLWLRYAT |
| Other | - | MNKSYFLFPFLLLLSCCGSVQVADPVYKADGRTAIKGYDPVSYFTENKPVAGEEKFQTSWNGAQWKFSSRKNLDAFRKNPENYAPQYGGYCAYAMRDGETYEIDPNAWKIVNGKLYLNYNEKVNGFWSRDIPGNIKKADEQWSKLPKKSQAP |
| Other | - | MNKSYFLPIVFLLLLDCGSRQLVEPVFKPDGKTAISGYDPVSYFTESKPKEGNPKFSFRWKGADWRFSSQKNLESFKKSPENFAPQYGGYCAYAMRDGEAYETDPKAWKIVSGKLYLNYNEKVHGFWERDVPGNITKADNQWKVLPRKESNP |
| Other | - | MNKLYFLFPSLLLFIAACSARQAVDPIFKADGLVAIRGYDPVAYFSENKAVAGNEKFQTTWNGAKWKFSSNKNMEAFKKKPESFAPQYGGYCAYAMRDGETYEIDPNAWKIVDGKLYLNYNEKVNGFWSRDIPGNIKKANDQWIKLPKKKEIP |
| Other | - | MAAGMLIVLAAITAACTTTPRNPNPPDFRYQPRYSVNPACSHGFRPTNALSCSY |
| Other | - | MSKFRAFMAAGMLIVLAAITAACTTTPRNPNPPDFRYQPRYSVNPACSHGFRPTNALSCSY |
| Other | - | MPREETHMSKFRAFMAAGMLIVLAAITAACTTTPRNPNPPDFRYQPRYSVNPACSHGFRPTNALSCSY |
| Other | - | MLIVLAAITAACTTTPRNPNPPDFRYQPRYSVNPACSHGFRPTNALSCSY |
| Other | - | MHRICVLAVTALILVAGCSGGNVVVAGAGPDDMLKLRAGPGLGFTVIMGLPDGTALTQQDCVTELGQRWCRVTLADAPGVTGYVSADYLSPR |
| Other | - | MRTHFAALLVLLIATGCTAGSGAVVKGAGPNDLLKLREGPGLEHEIIIGLPDGTRLTRQNCVTTDGKVWCRVFLTDKPSVSGYVSADYLAHR |
| Other | - | MRRSLGALAGAMLMAACSGSPADVFADRYEVHGVESGDMLKMRGGPGTGFNIITGLPNGTVVRVQECTQTGGTRWCEVVMDRPGGLRGYVSFAYLRAV |
| Other | - | MRGPVLGALAGAMLMTACGSQPADIFQTRHEVYGVEAPDMLKMRAGPGTGFNIITGLPNGTVLWVHDCTQTGGTRWCEVSMDRAQGLRGYVSNAYLRKL |
| Other | - | MRAAFLLVPLLLLAACKDEPDFDTRYDKAAKEIEARAKAMDADIAEAEKAAASKGLQAETKPSNPPASSGE |
| Other | - | MRALLIIPLLLLAACQDEPDFDTRYDKAAKEIDARAKAMDADIAEAEKAAAAASDLPDAAPPSTAAASSGE |
| Other | - | MRAAVLLVPLLLLSACKDEQRFEDRYDKAAKEIEARAKAMDADIAKSDEAAAAASDLSERPKPSNAPTSSGE |
| Other | - | MRAIVPILPLLLLAACQDEPDFDTRYDKAVEAIDARAKAMDADIAEAERAAATASDLPDESPSSNAAASSGE |
| Other | - | MTRIKRLVRPDTQALNWKAAIPVLGLAATLLSACASTAPAPEEIKERAVVDFSTCAKPVWPAESLKNENTGAVTLAFLIGTDGKVKDSSVKKSSGFVPLDEAARVGIEKCSFKPATVNGKPVEDWMLMQYVWMLK |
| Other | - | MSLSRAALAGLALAAGLTGCGKSSQPPAAPTVPPTELAALKTPPPEYAPELACAGVGGTSVLRVVIGIEGKPTDVSVTQSSGQPVLDEAAMKRVREWQFKPATRNGQAVPQTIQVPVAFKPPVPQPDECFAIEERARRGG |
| Other | - | MSMSRAALAGLTLAASLTSCGKPSQPPAAPRVAPTELAALKTPPPEYAPELACAGVGGTSVLRVVIGTEGKPTDVSVTQSSGQPVLDEAALKRVREWQFKPATRNGQAVPQTIQVPVAFKPPVPRPDECFAIEERARRGG |
| Other | - | MRPMYAVAILFLLAGCSSVPAEKTRAIPAERLLGYQQPVSGGGRLEVHRDYGYLGGGCYVAFLIDRQVAARIGVGEEASFQVPAGEHVVGIGIDTQDDTLCGKGLLNRELRTRIAADGNARFRIVSEASSGFDIRAE |
| Other | - | MRLISAVLPLLLLAGCSSFRADPEDVRPVPADRLLAFQEAREGGGQIVVNRDLGMMGGGCYVAIEVDRQVAARIGVGEVASFQVPAGTRVVGITLDKQDDTLCSKGRLLRELAVPVKVGESHAFRIVSQNKGGFDILPEQPRP |
| Other | - | MRSILAVLPLVLLAGCSSFRPDAEHITQVPDDRLLAYQEAREGGGQLVVNRDMGLMGGGCYVAIEVDRKLAARIGMAEVASFNVPAGTRVLGLTIDPLDDTLCGMGRLHKEVAVKVAPGSVQYFHIVSENRGGFDIRPDAQPPKLQ |
| Other | - | MNIFRTKLCVSFGLLTTSVLFSGCNAIAPVKASPATAATPAKASDVTASNTPVKSIEEIANEAMQSKGQLSREQIDALIRANAKCKPNDPRYNS |
| Other | - | MHTSPQLTVIVSILLATSLITGCTETLPSKATEPQSVQDIAKEAMNSKGKLSKEQIDALIRANAACRPDDK |
| Other | - | MNKAHPHAATAASLFLTASLFAACSEAPPTKEADSKSVQEIAKEAMTSKGKLSKEQINALIKANAACRPGDQH |
| Other | - | MKLVKGILFALSVVAMSACGTQAEMSAAQEGEAALSQQESRQEVGTLAALDCSVSIQCSNGTTRSCSGSSGACSASASGSGSVTCNGVTSSCALTLCSCRADGCCNNTCAADPDCGLSNCPQGAACSSNTQCGSNGRCVSGQCMCLIEP |
| Other | - | MNQERVMKLVKGILFALSVVAMSACGTQAEMSAAQEGEAALSQQESRQEVGTLAALDCSVSIQCSNGTTRSCSGSSGACSASASGSGSVTCNGVTSSCALTLCSCRADGCCNNTCAADPDCGLSNCPQGAACSSNTQCGSNGRCVSGQCMCLIEP |
| Other | - | MKKFTLTIIMGVFGLTACNSFANMPIKDNTTQAQVQTPITPKAKGVWIDVRTPEEYQAGHLTDSINVPVDDITKRIFAIEPNKNSPINLYCRSGRRAEVARTQLLKLGYTNVTNHGGYEDLKQQGYR |
| Other | - | MKKLTLAIAISVLGLTACNSFANTSSKQNNAKSQTQTQSQIKPKAKGVWIDVRSPEEYQAGHLSNSVNIPVGDIASKISAIEPNKDNPINLYCRSGRRAEAARTELLKLGYTNVTNHGGYEDLKQQGYR |
| Other | - | MKYIAPLLAALALSACSSRPVTSGISVEQHKNSFSDQLRVDNLNLAKKLAITDVKTRQTNQLTDVVVTLSSNYKKSQYLQYQFNWFDKDGFIIKGNHSPWQALTLFGFAKTQLPGLAPSPDAVTFSLAVREVSTKSQEFKD |

| | | |
|---|---|---|
| Other | - | MKYLVPLLAVSLLTACAGRPTTSGIAVENASAQYQQQLKVNNPQLGKRLVISDVKTRQTNQLTDVVVTLSSQYKKSQYLQYQFTWFDGDGFVIKGNHSPWQALTLFGFANIQLPGLAPTQEAVTFSLAVREVSTQAQEFKE |
| Other | - | MKTFTFTKNIFFAALTSLVMLLTSMGCSDNEDKTATATSEHNPFDHPHDFPVTDIQKHKFEHEFAKQCVQNELRNSVNKELDKERYTEPCMCIATFMMKDLTAQEAKKFITEHENPRSLQIKFDSAAYHCLQQKAQAIKGPQLFGKR |
| Other | - | MKTPTFTKNFFFAALTSMVIALAVTGCSDDEQKTAAAKPAAANVVHNPFDHSHDGEVTDIQKHKFEHDFASQCVDRELKNSTNKEFDKVRYATPCMCIAKFLMKDLTAEEAEKFIGEHKNAQSLVIKYENAAYHCLQQNAHPKGPDFSRAPQQAN |
| Other | - | MNMQWYRLGLSASLALVLAACASSGGSSNGNWQNIGTTSNGNIKVAVDKSSIKSNGQLVTFRDRKVISKLSEEKFANTPAYKTALADWEIHCGNKTYRLTALQLLNERGQTVTNQRYTATNLRPMSVMSGSITEKQYELVCGKKL |
| Other | - | MIKETLMRPIFLSFVLFPILITACSTPDKSARWENIGTISNGNIHTYINKDSVRKNGNLMIFQDKKVVTNLKQERFANTPAYKTAIAEWEIHCNNKTYRLSSLQLFDTKNTEISTQNYTASSLRPMSILSGTLTEKQYETVCGKKL |
| Other | - | MGSKSIIILLMAAGIGLGACSQGNTQQEVSTADNVSRATPTAKAATATPKTAEDVVINGNNTVQTQAVSGQNIEVTGDSNTATFTGHSQEFNLTGSDNVVVLENVKTIEVTGDNNTVTWRGSTPTVTNLGQHNVIERAK |
| Other | - | MIRKTAMMLLLAVGTGLGACSSGETHSELSGSDSIPSATGEASRASHAAQQVESATSASSAEDIVINGSNTVQTLALQRQNIELTGDDNTATFTGQGRECSVTGSNNVVALEDVKAIEVTGDNNTVTWRGGTPTVTNLGQHNVIERAK |
| Other | - | MLKISLAAAVLLTVSGCATQNDWGAVGGSKSDAVVRLSYELGALDKHTPDESDGLKLAKLRCGYWGFSGAKPFDVETKTCADEQCRSYTITKEYQCVD |
| Other | - | MKKRTKLLFATCVMLVVSGCATRSDWEAVSGSKSDGVVTLRYELVASKAYLPDEPEAIALAKQRCGYWGFSSAKAFDTETQHCSDEECRTYIITKDYQCIN |
| Other | - | MFLVGCNGAAQSVRHATAQQERCVDVPRSDVAPGARNEAFHSARAGVEAVPVAATLGAASPTAPLTPPSSPPLEAQSPQQPIVCSCSA |
| Other | - | MSRAWAVLLCMFLVGCNGAAQSVRHATAQQERCVDVPRSDVAPGARNEAFHSARAGVEAVPVAATLGAASPTAPLTPPSSPPLEAQSPQQPIVCSCSA |
| Other | - | MKKIRQTLKQFGLVLILGTALSFSACSTGTKDGDTNVEDSGAKDKDPENMNNTETEVSGRPDSLDMDKNKTYQKVDPDSGARDADNDGKVDQ |
| Other | - | MKKILLSINRFSLVLLLGAALQLTACSTGTKEGDTNVEDGGAKDKNPEKLNTNEPENSAAADSIRNSDDPDANKTYQKVDPDNGTRDADNDGQADQ |
| Other | - | MRSFYATLAISCSALTACSTTPPCEDILEVKRQEQECKRLSQVINNPKNPQQALTARQRFEAECENLRYYRDDYDTICKGSNTPIGNVETKRKDP |
| Other | - | MKFKYGIAPALLALIVGCSSKSPCEDVLEVKRQELECKRLQAASNSKNLQQAGVAKSRFKAECENLRYYRDDYDTICKGEQKPIGEPQTAPAVKQD |
| Other | - | MYYCRWLVLVALLGGCKDSTAPATPVEIPEQRVVATVYVVNASSGSANCYRTPDSNAPSVATLRNGQLVDLVAEQEGIIQRGTQFWLHVYPRLSHRPSCYLNVDSLVPVS |
| Other | - | MRGLFYFFTTLITILLTACTDNGNLLNIADISSRPIATVYKVNTAAIDYAPIECRRSPAKNAEVMIRLYNGSLVDLVALEEGMLKRDDRYWLHVYPPLSHRPSCYLDVRYLVPHA |
| Other | - | MIMKKSGLVIAATAAAIFLSGCATGARNIPENTPLCEASCPPPCVTPCNTCKCMSQCKQVCPPKYKKHRHRRCRTAENATNQMNG |
| Other | - | MKKTGLVIAAAAAAMILSGCATGARNIPENSPICETPCAAPCNTCKCMSSCKSACPTKKHKRHKKHRRCSAENTQTQMSA |
| Other | - | MHAIRIRTMIATASLVSLGLLSACSKEAEPSANASGAMSATENSGGMADGAAMTSDNSMAADGNSVAAGNAMGDSMQPSDSMQKNSH |
| Other | - | MPVLRNRLMVVVFAATGLVLLSGCNSTSGGSSNNEAASGPSDNGGAMADGAAMMSSNGSMAADENAMSAGSENGGSMQSTDGMQKNSQ |
| Other | - | MKKYNMMMAAAVASLTLAACSKPAEAPKADATAKADAKTAADPAKSEKCYGIAMAGKNDCANKTGTHSCAGQAKTDNDPNEWKYVDKGTCAGLGGKTA |
| Other | - | MSRHPLPLPGFAAALAGAAMLAACGSGGKVVQEPTTEANGKGTEIATREKCFGVALKGHNDCKAGPGTTCAGTATVDYQGNAWKYVDTGSCEASGGSLIERAGNTPPAAQKG |
| Other | - | MRTLLALAFVALTAACGGGSDEAADPPQLRQGSVQVMSSERVLVDITVSAPIKLGKNELAVDFPSRPNTELYGVSALMPAHGHGSPAPTIERTDEGFLVHDVVLYMSGRWELHFQIRVDGRDDEAVVVVDVP |
| Other | - | MRLPARLANVLLSVLLACACSPGSEQGTSPDAGRDTASGLVRLEAGFEGTLQLRGNMLRIHVTDTAGTPVEATRVSVSLWMPGHGHGAPAPAVTREARGDYLATVDFTMPGTWTVTIQVDTEGRSDTLELSVEAP |
| Other | - | MATKMGKTLAFTAISGLVAGLAACGGSAPPAETPEAPAAADAPAAEGGEKAGCSGAKEGEEKAGCSGAKDGEKGGCSAKNGCSGAK |
| Other | - | MATKMGKTLAFTAISGLMAGLAGCGGATPPADAAAPAPEAPAAEGAEKASCSGTKEGGEKSGCKAGSCGGKKEGAEPAAEGEKASCSGAK |
| Other | - | MNRTIQRARFALAGATVAIVAGCGGGGNDDASTPPQQVPASASASVDGFISYLKSLLGSDADRSEPADVGAFTPPLDETGEPQKVE |
| Other | - | MNKLQLALIVVSSAVITACGGSDDEVMGTPPVVVVPPVTAEVPPSASATEAGLFAYLDALFKASADGLEPVDLSTFNPALSDTTEPMAIG |
| Other | - | MRNLAIASVTALALGLAACSAETEQKADAAAEAAGDDIEANANAAGEVIENAADDVEASVDEAAKDLDAEADKAGDKIEKEWDEAKAEAKEEAQ |

| | | |
|---|---|---|
| Other | - | MRTFAIVSLAALALGVSACSAETENKTDAAAEAAGDDIEANVDAAGEAIDDTMDDVEASVDQAAKDLDAEADKAGDKIEKEWDDAKAEVRKETD |
| Other | - | MNPLKSNFYGIIIGLITALLMSCNESKSTTKENTKEYKRALQKLKNTDSIKPEPLDTSSSEEMNALKKQLKEDQYGEWPIPNTVK |
| Other | - | MKFKLISVGAAVLVLGGCASLMKETVSKRAPFDLNCDAGSITIQQLGYRTYGVSGCDKRATYVLQGPCSGPGSQCLAVMNSNVDEG |
| Other | - | MKKTFIAIAAAATALSFSSCQNPYGPGATNTQRDATTGALIGGIAGAVIGNQSGRSLEGAAVGAALGGAGGAAVGSTKDKQQNYYR |
| Other | - | MNSKLKTSLVVAASALLGLSACGGGSAQSSESTGETASSGSETTTETTTTESTGDATGTQASCGAGSCGGAAGESTGEAPAEGSGT |
| Other | - | MLKKVLVVSALALALGGCETARQDRMAGGALIGGGGGALIGGLATNSVGGAVAGGLVGAAAGAIIADATRPGRCYYHDRYGRRHYVRCR |
| Other | - | MKLFTRSLFATPLIAAATLGLSACNSPAPADNGTTNDLKLVNEEAPLDANLSTDGELSGNAGSLDAVPTGNETALGNAADGNASVGNTTAH |
| Other | - | MKPVLFVSLILTLATSLTACDNSSNTDKGTEVSAQLKETGAKVSEPAENFSATVATEAEQVSQSAQEATSDTTEIIEDNLESIQENVENNP |
| Other | - | MKRIVIVCSLLFILIITGCNNLSTPKISEDEAQSILLEEHTKHIGKVKIISVSHKGEEYVVQWENKENCKNGTDYVHDQTGEITKGEVSIC |
| Other | - | MHHLKLFCLAFLFTFLVGCSNERIEISNDVKLFASIPDGAREFEVEVLIIIPKGEVLKVNSRDYMKDFMVIGVTYRDVEGFVIFDSRKMQLIQN |
| Other | - | MKRHSASLAVAGTIVAVLSGCAAPQESLPLPSPGPQACNAAASQSLIGSQVGAVSFAADANVRVVCTTCPTTRDYRPDRLNVRFDQATGRIEKVDCG |
| Other | - | MKKILIVVVSSYLVLTSCSKATNKNAIVVRDCTGTYLQIDGKDYQVCNLEKVAAFQDRTAVTATYKKLAECNGSAKDAIVCMMLHSNEGWIEVTDIK |
| Other | - | MKKTSSLIALAGALALAACQDSASQNSAAPAAQQQPAAAAKAGEGKCGEGKCGGAASAAAGKAGEGKCGEGKCGGASAAASKAGEGKCGEGKCGGEKK |
| Other | - | MKKVAVGGFAVLLSACASMAVTDQRLEQNTAFALGLDVGDFTIANRMDEGLKTTYSVKTKAGKQYNCYVMGTIGITGKNVSDAMCNEKGKPAVNPLTGK |
| Other | - | MKTAKFLAVIILSIFILGCGNSDPGPLAGTWQMSGLMPMTVQFRKGETETMGIIEKVSYKISGNDVLVTYEDGIMKGTTMRYTIADKDTVKTELGLLQRVK |
| Other | - | MNKISIAALAGALSLAACGEAQKPSQPEAPKTTEAPAATSAPAKAAEGKCGEGKCGGASAPASGAKAAEGKCGEGKCGGASAPAKSAEGKCGEGKCGGAASASAK |
| Other | - | MLTMMTMKTRSMQPSRRTAAAGLLLALTLAAGCGGDPHRGDGDGGSNGGSNGGGDNGSLPDRVSTAAEFLAYVQQVIARFSADTEPLDVDRLGAPTADRDEAADV |
| Other | - | MKPVITGLTAIALFGSLAACGGTDQPAENLTANDVNAMMAPPELPAVNDIMADPVTEAEPAAETQTPAAPAPAAKPTPTKPVPAAPKPKPAEPAEDPHAGHDMNNM |
| Other | - | MRSVGTTTSRGARRALHRGSAAGALALLLLAGCAGDAGSAPGPEQLGVLFSEHAAVVLGGAARFARDAGGHGFVARTPGGFEASLPGHGEHAIALRLPGGARAHGA |
| Other | - | MKSAHVILFITTTLLIGCNNFLPENYVKTNINNSHAINAIDHKRINTTDCKDADDWYLDGYRVGKSFSTQKKEMLDQRLGFCHLTSKKLPKKFRTNWEKGFSVGSRG |
| Other | - | MKTSALFAAGLAALFASACTSTPKVDVRQPGDRNMSCAQLEAEMEALDDIQEEAENNQGVNTANVAAVVFFWPAAVGNYMEADRAMDLAEERQDHLMDIYTEKNCDG |
| Other | - | MKLRYLLLVGLAVPLTACSVHVGGHRPPPYYAPPPHVEWRWDPGIDAYVVIGWPHLYYRDHIYYRWHDNHWYSSGRHDGPWAKGPKGVPPGLNKKYGKPGRGRGGGGY |
| Other | - | MKIKVLSAVMLSVLLSGCAGQMAVSNATMKFNMDAVDNRYARGGLTILMAPVYAVTTVADYGLFNPIEFWTGENILTDKKSIYDMKGKNYIEINDDLDESLKTAPIKLD |
| Other | - | MSLNKGKYLATATTVLFCMLLSGCASPPEHPQLVEADLLFAALQSKSESITLVPQETQAAFVPLAQAYILSNRNRSDPRIEELSILAKKRIALAEQLISEKLALGCNGSC |
| Other | - | MYLRYLALAGVLVMATGCVEERVVHERRGPPPRAEHVEVIAPQAPPERIIEEEPAPRPGYIWSRGYWRWNGNRYVPVHGHWEAVRPGYRYQHPYWENRGDGWHYHVGVWVN |
| Other | - | MSKSNSLKGIVFVLFSFLMLGCTSSYESVTQVDDTKAFILLTGNFENASLQINTNSPITLTDSIETFMLDGKKVAKFEVTAGTNTVKVFKNGALVVNRKLYVTNGNSFEVNVQ |
| Other | - | MRKGLVFIAIAAFVYACGGSSAPAESNSSAGAKAVAAVAKPDGEKIYKQYCVTCHGLYGDMGASGAYNLQESALTLEERINVITNGRKAMTAFEALLDEKEIKAVAKYTMKLGE |
| Other | - | MKIKGLIIIGAVLLLSACNSTQGGLYWGSYSDTLHNYKQEPGDTTRQRHVKTLNDIIKTSDKRGTRVPPGVLIELAVMEIESGSSENADALLNREMSLYPESRTLVLELKKRNGA |
| Other | - | MRRILLFLVITGLLTACDNSPKPPSERVTAALPADPALARVYDTSCKLCHANPASGAPLRGDRAAWSPRVARGADTLLDHSINGYNGMPPMGLCMQCSEEQFLALISFMAGVELQ |
| Other | - | MILNNLKVIFIFVTCLTLSACSLFDIQQLDDKTKIVKIYDAFEHIEQCQFINELVGSEGTWYSYLFVSNKDLTLGSINDLKNQANAMGANAVHIQYSLDFNTSVTFFAQAYDCAQ |
| Other | - | MTLLQVINKYSGITIALALLLSCNHVSEQKTDADSTARVQTFKTKDGWAYSIFLGKKEFIRQLYIPCIQGQIPFETDSQALKAGTLVLNKLKDHQIPSLNLKELEENHLLPAKDVNNNNK |
| Other | - | MRHHIAATAAVALAVGVALTGCTTEQRRDLLGDVAAEGLRVAAEDAFASAGFPIEDQLDCQVEDVGESEIAADCTGTTQQGADVTFTGSYDTGDTDFTDGVQGTFTGSVEGTEVFSSDCLGCGG |
| Other | - | MAQSIAKMLKALLVSGLLFLTSAVTACGYAPQMAWADGLPVTTMIGVTSSGTSAWIALSGLDVSQDFDPNKQERIQGTTDMMEESMLVPGGIPSVSKNVEQSLERASGLITGETQEQMMKQSSD |
| Other | - | MGHGVIRTLVAGGALAASAAALLAGCGGGGNSAATCGDAKKAYERYIGGVRAVSAADAAQWRQPTEQLAANLSGLADKASDDRLKSALKDQADKLRAAAGTVATGDVEQLNATLAAAPTALGTACG |

| | | |
|---|---|---|
| Other | - | MHTRSDSPRRGLSLFFTGLLFTGLALVSIAACSSQQPATQGTSMAAGERRENDCSTGPLLECPEGQLDGCANGTTTEHRCVEAGRGRCLDIRAALVKCLDGEILTHDGCPEPAYMQRCAPRSAQLP |
| Other | - | MKTSYFLKFIPISFFTLAAVICLLSSSCSPAYEEIEVVYQATDGQRITAVYHNPTNDESEFSVTLKFPSGQSVTLNQGAAASGVRYTDDKTLVWWTKGGGAIMMKPDGKNDWEITGTYTEIPMHHNR |
| Other | - | MTMLSGKYTLLGSSTLLLTGCVFVPHTEVSFDEYCGVYKESTSIDMAPRAASNLVFHELELVGGLALALSNGVVKAQQHQAYKEACLANGIDIDAEAKLSADSVRSQVKQSHYCASQASDLCEIERRR |
| Other | - | MRLRQHVRRSVTAAFVAAPFAVLVLSGCETLDQAAATVDKAGVCVDALQAANFTPDVNNPDISAQNAADAAQKLSDLAGQTTDVTLQDALEAMSSTISQMNVGDLNPQAIADWTQQKADLYQSLSSACR |
| Other | - | MIAGTAKRAAAAAALTIVAAPALSACGGSDLDGAYYDKSGRITIDGGSVTYHTLGCESTGKSAVIINDKAKKTGELNDAGDQVIWSGGGGTKPITVSKNGDTIDIDGKRYAVMDEKEAMDGYKHMCGQN |
| Other | - | MANRLLITVFILLFMQGCKPTQQQTVFYGQPYVYVSMFELLVHTERFNWRDFIIRGYLADGYLYLSPEQAKHRDTSYAIAIHPDLSSPAINFSDFNEKWVFINGRLLFEYDSKAYAGYSTLVVREISSRD |
| Other | - | MFRTLLMLCVCIVLTACSGTPSDTLIEESVAQQKTVSNMIRVVSAEKLNGWKDQEFYVADVRYELEFLTDYKTFSESLKDETPDSLVGSFFSGFGLLALSMQYGKFEKGQKVTERAEFRFRDTENGWQLAD |
| Other | - | MNAYLLKAIALTFVIALTSACTTTEKYVSTQDQLQQQLNTLATLQRKITTANTEDERNNLLMEQQRAIQQSINTLEQIQQSKRDANIECIEREKSVPLGDETCHEVESLDDTRITMMVALMQQLLLRTQGH |
| Other | - | MRGFGLRLAGLLTAFVMAGCAGNQNGEGDTPPQVLQVTIQNDGAITQQPRINLVPEGGGAMTILVGRLTTLGVETLTVRRPDLGGTYRLQAQATGGYTVNSPIFSTRGNERLFWDLRRNVVSVRSLTDDSAQ |
| Other | - | MKRFFLLSSVLALASCASFINMHRMHAAQSKLQATRTLQQSGQWDAALEMAERMHSSVAKTIQSAPTQKSPGGKVDLRPLLTGWESGPFAALKTALQKHDAKASTAAFSSLRQQCMNCHTVIGKADLRLKEIP |
| Other | - | MLALSVFLAACATSPKATTYDQLGGASGIEGIVDALLEKIVEDERINFQFADADIVRLRSMLIEQFCAESDGPCTYSGLSMQESHAGRNIDDAQFNALVEDLIEVMTVRKVPVGTQNRLLKRLAPMHGDIVEP |
| Other | - | MRYQRLPALGAALFALGLLSGCFEAKETAPEQRHSSLSNFQQFIQKTGGLKSLNTFQKRRPEINSEESIGVIYDGARTQGEVMRFRDKQLAKEMLPALKAVRQRLGESENCTLRGYFIACGEEKFVDVFKKWE |
| Other | - | MNLRWLAVSGLLLLPLAGCAIANDDAAASEVSFSSIAGTSTAPSAAPIDVVDLAAPGTICGSVAAAQDDAPVVIRSGSANCVRVLRAARAYSSAAAHADGQAVTVETGGWRCTATVSRGPATCRAGDASFSVG |
| Other | - | MKKRNLSIGGFALALLLAGCGTPYEQCVRQVSKESRVLKSAIAESRLRISRGYAIHRQTVSYRVPSICHRHDHKMHRPAPYPCGRTEYTTIETPVPIDVAAERRKLASYQKQLVQAEAATKRGIRQCEAQYPKEG |
| Other | - | MASLRPRTLFCRLPLALLLCAALGACMSERPPIGLPSAQSIQFDGVHALGPNCASIALPSTIGDPDMVEHPSIPFGCATYSNLAAQLARPADIVQPTPYGGTNGVVAERALSRYNAMPEPHAAPDAAPATTNVGH |
| Other | - | MKRFILVAMLLTVAACERKQETAEVPPSTVPPETTQMMPPVTIAPAPTVPPATTPETAPPTTQPSSGSSAPAGSSGSEGAMGSSGAAGAGEYTVSAGDTLSGIAREHNVTSGDLANWNNIQNPNRIHKGQTLRLTAP |
| Other | - | MKTFYAILLTFIVALAGCNGNKVNLSSLKPAATRTVGKLTIVLLNKSGELMQGQNEFVIQFKDDQGQPADVGDVQLGSNMSMPGMSPMSGDAEITPAGQTGVYKVTSNFAMSGAWHFTLSWSGPYGQGHTTFNNNVR |
| Other | - | MTGPSRLAALALVTAAAGLAACQREPAGGPLIATVDNSAAIALLQSVNMQAQTCWMRSKDRDFAAYRLIPELDTRVGKPRILLVQARAAQGLPQLVIEAHGNPVQLTTYGPLTGKPLSGRINTDVTRWATGGTGCKA |
| Other | - | MTISSARLSLHRSLPYIGIVCLFLAGGCATSPSERQGQRSDIEIGTTTKADVLKRYGAPDLVQMLPDGEIATYRPSASPQAKPTVSVPTVQAGPAGMMTTQNQTVEPGLGKNTRSSNRLQTELQIRYDQQGVVRELIQ |
| Other | - | MKKTVCLLFVFFLGCAGRPDSTAVRISLVNNNRSIRFKGLDPAIMGEVSRNASPAVWENLIPVYRMPADTDLKDYQPIQHGVYQVKDSVIVFTPDTPFVKGHTYFLRYFRFEGGTKPWDFIGGKKKLGSVPHQDLVVRD |
| Other | - | MRAKPAVAAVVLTVSLTLAGCGAGSSSDSAGKARGVERQADGARGEKGGEDGGAAVAATGAPGAEGKAADGRNPAVVGAQVIRTAELTVEVRDARKALGTARTAAEKAGGLVRNETTERVEDDEVTSTLVLRVPQDAYDA |
| Other | - | MRRMMLLLAASVVAACQAPMPAANPQMAWVDFSTPFPNDRLLLAERLDKQRLSDGRFFEVSPGRHELIVRFDYEVTGGGGMGMMGGPTVRVCYLTIHYEHFEAGQRYVLEGRSMAMTPEARLYNAKREIVAEASDYYCLM |
| Other | - | MSRVITKSVAAAALCVTLSACGSFFSDAQSRHETPTLPPGTPNPIALAKLDGLWTGTGKADGLTVRIARAGSSTPRALYGFRGEQPIEPQCSAADATLNCTLPAGLTTTYTANADGSVDFTAKGPNPGDELIAAKLQRAQ |
| Other | - | MMISKSWAAVTGRGGLAFGLLAILALAGCATPQQRAAEKDDNLAAAGFVIRPANTPQRQAMLRSLPPNRIVQRAHGNTVSFVYADPLVCNCLYIGTQQAYDAYRRYMQQKQLADEQQVTAETWSDAGWDWGPWGPWPWGP |
| Other | - | MTPRWIATLALPLFAGLSACAMPPGPGGAPSLVGTEWRLEDLGGAGVLDRVPATLAFPEAGRVAGNGSCNRFFGSYTLVQDRIAFGQMGMTRMACVGAVGEQENRYMAALQKAQRVQVQGSTLTLSVEGMDKPLRFVRTKP |
| Other | - | MNNMIKALAASLTVALALAACATPVQRTASGEIPPARIYIKSMTEQAAGLSQVELSRNGGLLNNELLELAINDVVLAQIAGGEHLSIWLKPGSYDFSVKRINTLTEGSAPTQHKLTLQVNQSGAYKIQISSELRGLSIQLMK |
| Other | - | MTKGNIYLILAAAGLALVGCNKAESPSEVQHDVTNAQAEAQRDVTDAQSDARESMADAQKDVADAQADNDADDVADQARDASETAAQGDFKVAVARAEATHKIAIEKCEALQDNAQEDCKARADADLENAKRAAEARRDGAG |

| | | |
|---|---|---|
| Other | - | MNRLLHTVVFFPLILLSACATGTVVSKSPEAKSPIPSGKSRIVVYRTQIVGMAVQPSVKVDGQKTGLCSPNGLFYVYVTPGVHEVSATTETTDTATVETRAGQTTYIECSVTTGIFVGRPSIIEVTESRAKPKISGLSFTGQY |
| Other | - | MTRQQRTPMVRGGIALVLTASVLTGLTACGSKNDADKFAEKLRTKGYGKVHVSADIERKGGKKRTVAYDAHVLVNTDADPQTCDVELENDVRSSGRGLVDYFDVDEVRDARGSGHEVEDDRTWPDNPTLTQLRAELVEHSIDC |
| Other | - | MRSRARRVLLWAVISGFTALGCEGATPDPPTGEPDGGTSLPDAGPGSEEPSGCPVVPAPTACPSPAPHYPDVAPIFDRRCVVCHSGSPGGPWSLADYGHVADWQDTIRTNVRDCTMPPADAGVPMTLEERLAILTWIRCGLPR |
| Other | - | MNHYLKRFIPALAVLGLTACAGMHGQHAHHGPLDTAPPGGEYAQVSELVPLPAFIPGLGTLYVQPATLPAGPFLAYDRDDRLVSTIYMIPLDDLNARKNFPELATAERKVVRTELAYNAGHPGVAEPHYHIVLWHVPPAEAKLK |
| Other | - | MPRFMLNRSAALLALAAAAVLPLASCANAGTNSSTTQKDVMTSGVGPMLEEIYASRKPVRFEQIDVSDIVAKYIPPSAEKSTVLETFGKSPTSKIVENTPDKIVVRDNKGQAMLDPDARSIVMTFFLDADGKVTKVDAVHIKNQ |
| Other | - | MRKITLLSLITITTLFLLGCDAQKTEQASAGSISLANPASTYCVAQGGTLSIESEEDGQVGYCTLPSGERVEEWQLYKRDHKESNEDIEVTEMLHTPNPAAEYCLKLGGTSDLSSGNCILPNGEKIDQWQLFRRDHLFIDPQNR |
| Other | - | MRKMMTAGLLAISGVLAASMGCIAPPADEPIADEGVAAEVEEAAAEPTGEAADELSVCGGGLVCQWAFASGAPQCEQYPSGPRLYCCPSGYGLVNNTCVPKCGSGLQCSGSSIPGSHSCTQRDTGSSVIHCCESGQRISGGRCVW |
| Other | - | MNMRKASPRSLWQRLGLATLLAGLLVLGGCMNQPTHAPDSRTAVIAPNSGTLTSQQVFVTLLTTLAERGFIIERADPDLERLDASYAARPPLALEAWVSEVNGQIRLSVSGDSNGADIAPGRLDNLLVEVAVALDARTVPVIGAP |
| Other | - | MRYLLLFILCISLWSCSTSKTTSKKEPISKDALAQLQDKRWVLQKLPDTTVPKMTKQIYILFDKNGVEVKGFGGCNGFGGSYKTDDKGLHLSNMISTQMWCDYGPVEQKFMHAIQDCDNFIITGDDMQLLKNNKPVAYFTAVYLNN |
| Other | - | MTNTFHATLLATLAFIISIALSACAQTKALDQGKPGKAFVADTSVIAIFNDRSKINFSSEELSLVDSILQQCINDYDLMNISAYKRQYVPAINDKGEKVVWVNCFCGTSRNWKKDLIVVNDGGSCYFNVTINLTSHKYENLIVNGVA |
| Other | - | MTNAIRPSNPVARLAIGLLLGTAFGCGEAPEPYRAIQGRVTYQGKPVPHAVVVFESTAQELTIAADADEQGIYKAIRSVNVPGLPIGSYRVAVTPPLFYPGLGPVGKNPGPPTREDIPASYRESATSGLTIDVNKEGSDASFDIEMK |
| Other | - | MKKALLLASSVLLGSVLVGCQTSEDSTTMTQTTPAAMKTVTGTLAYRERIALPENAVVTVTLQDVSLMDAPAKVIATQTFETKGKQVPFAFELAYDSAQIDARHTYSVSARIELNGKLRFITDTHYGVITDDNNTNQVDLKLVGVSAN |
| Other | - | MNKTLPNFLLISSLALIFLAGCSQDKSNEPVSGRWYTPNQLALGKTIFADNCAACHGTNAQSIPNWQQSLADGSYPPPPLNGSAHAWHHPLNLLVRTIEEGGAAVGGKMPAFKNVLNKQEQLAAIAFFQSYWDDRIYQIWKENGGLTK |
| Other | - | MRIPLTLSARPALLLSLLLGACAGGPPQRSSVNPNDPSLRRGEALVRHGAGSDALSPVAPAPAVQDLGAQAGFGRAWTASSVRASVYLFNTYQEARVAEDWLKAHVPEGLRGAGTVNGDLLVWATADATDEAGRAILENLIGSFAGEE |
| Other | - | MRNWTMAILPLALGMTVTACNGTDHDGKPQGKSEAGAPAAAGPVTTQSYALTGFTGVEVAGPDDVTIRQGDNFSISARGRADVIDRLEIKVDGTRLVIGRKRDGFSFSSRDDDDLDIAITMPRLDALRLTGSGSIDADAAGGMPLKPS |
| Other | - | MHSPLRSASFVVLLFALEGCSWLPSWLGGAPAVVNTTIASPLLIEDAGLLATRDRRAEIGIRLNNASSRPLWVGVRVQTPGGRTDCVILKELPPQAHQLFTCPQTKVFADVDYAVVIDSYRDPGLNERLAPLQTLLRFNANDLNTIGQP |
| Other | - | MDIRTISATLALLGSGLTLGGCNKAQTDATEVPGNAGAAGEKSCKGHADSDKSCGGHAEGEKSCGGEKGCGGEKGCSGVKGDTSCAASKPTETAATDAAPAADPATTATATAPADAAATPATTTTEKKKKKPAAKKAKGEAACGEGTCG |
| Other | - | MKHYQITAAMMVLMTSGCASIVSKSSYPVTIRSEPQGASVLIKDKHGVAIHKGTTPFTVTLASGKGYFGPGDYTLDFELSGYQPHHTVMQAGVNPWYLGNIFIGGAIGFLIVDPLTGAMWKLADQKSVTLNAQATDIIPPLASPAETSQP |
| Other | - | MKKRYRVFLALLVMFATGACLSKMSSHEMLSLQQLKWKHRIVVTTLVDNDAVNALAEQIKKNSDFFDERKLIVVVKVNDMLVEIKQNIMIEGNNKTQFLNRLQNNNTILIGLDGGTKATYPALNWQEIFADIDSMPMRRNEIERNIKDNK |
| Other | - | MRPNLSIDKIAGALAVLGTSVLMGACGGESKPAEAPVSGTEVAPAGEKAGEHKAGEANCSADHKGSASCGAGKGSASCGAGKADPAAAAPATTPAAAAPAAAAPAAADAKADSKAPAAAPTGSSKKPATPPAAGKKAGGASCGAGTCSAKK |
| Other | - | MKISTKLLTIMFVPVFIVACGGGGGGGGGGSSTPAPTPTVETIDNTQDIVAERNFSFDVGETITISMNYQGSADGALHLYTKAAFTTENGVVVADPMSRITTIYPTRTNEVELEVNGNWSQLYAQWVPMSANESEKNWVVSLNQSNNNYHLDF |
| Other | - | MHSFAKPGFACVVALLLSSLAGCASTGANSPSARPVLYPNASLNRVGDAQGRAEADACMARAVSAGLTPDEKNNAVARGAGVGAATGAVASAVGALITGRGGEGVVRAGAAGAAVGGSAGAVQGAFRNDRPSSTYRNFVQRCLGDRGFEVIGWN |
| Other | - | MFKKALVLALPVVLAGCSAPKFTAEPIAQSNQSKEITIVKDDATREVFMDSLQEWCLDTARKCTVVSDGTAPKTDELTLTYVSRWSWDLRTFIADAKINAYKNDQKVGHVEFKASNNGNLDKFGDDTKRIESMIQILFGEQTISDAQQKIKSGEI |

| | | |
|---|---|---|
| Other | - | MKKIIPSLVLIGACLCGCAESAAPAQPGPARPPVVMDEGTPWTVDDVTYTVTWVTLPHGYGGDPAPHVEINVHVDNHGKNYAHFPDPSAVYDGGATAVAGWTQPPSGAEIAPGHTADFRSSFPLRMPGGTLRVRVLAFTGDVEELGYWIGTVHSG |
| Other | - | MKKLLLVCGLSALLIGCSDNEVGDVSLGIFTLKDIKLNSLVDPLVPGVTCHVASIEADLSLADPSDSSISCRQTSEITPEMIAQINKSKDGEVIFTKSKSIFFKSMKIRRIFDQKHQTLMYLSYSTKETSGSFKHSLSTVPLWGTKAYVEPAQQK |
| Other | - | MHLMLLGSMLGLALLLAGCASSPRPDRQPAFRTLSENQSSDIAIHALGLVGTPYRYGGNTPESGFDCSGLIGYVVYKASAGVAPPRTVAQLVDFGGSIDASEARTGDLVIFGGSRPFHAGIFVGEGRFVHAPSTGGRVRLDRLNNPYWQRQGQAFRRP |
| Other | - | MRARAAHALVTAGTAVFVVACDGTASSGDAAGPAVEPTITVRVRGLTVEAEVADSDAERRQGLMFXTHLPPGQGMIFDFGGETTSGFYMYRTVLPLSIMFVRDGRVVGVREMTPCPGDDPSACPVYYPDGVYTHAVEAPAHTFAGVVVGDPVAMGSD |
| Other | - | MAWRLTLLALLLSLLSSCGFHLRGNLPLSQFPAIYVQSEAHSELAALLKQRFSQNQVELLGSYQQDRPALQLVRDTLERRTLSLFANGQVAEYELIYKVEYRIQLPGEQEQFYQFELYRDYQDDPNQALAKAQELELLLSELRQQAANRIIRQLARLG |
| Other | - | MHRSIPISLLLATMLAACAQTGQHPAAAVQPTPAPAQARPAPAAAPAAATTSTSVAPTPAPVVQADEPNVVAFATGSDKLDDAASAKITDLVEQARQAKKVEVLGSAQVNGKFNKKLALSRAFAVKRALAHAGVRPGKIMVRYSTIESRDVATITFKP |
| Other | - | MNLKAIGLVLLSALHVSGCGSDAHAGNEIETPPVEGGKNTCTGFGSPHINVFIRESLNSDVLIENASVRVVMESENEESTVEPVFISSDDRNEDTKTGAYHAILEHNNLAFDVSIVAFAEGYYPFVTKNISFEVNTGCGASNDLTYDVYLCPVGTTCL |
| Other | - | MHKHVFSLFVMLLFIHACATMGASQKNAESPASPPLTSILDSPGLKAFKRTLNQIVLERNPGTGQSQHFFVSKYSENESLAYMFWQEKKLLWIMSLGATDEESWLGVRYPSSGQLIDLKNSVVASVDDVGGSSYLVTKEWAAERLFDTVVHGDLIVISQ |
| Other | - | MKLFNIILVLLVVSGCKNSNTSENKLSNAIKVIGAVKAQEEQEAIDAMEDTKAMETLDWQGTYFGITDCANCDGIETELVLKDNEYILSLKPTNTDEEKFVQKGTFTWKGSIIILEDIIGSASMYKIEENQVKQVYYIDNKIQGEDWKGYILKKMTNSK |
| Other | - | MMRVTQRSLLSAGFCLMLAACGGKALAPPPPPTIVQLAVSADAGVNPDAKGRATPVVVRYYLLGNAGPFEAADFFSLFDGDEKALGATMVSREEVTLRPGDSVSSRLAPMQEAKALGVFVAFRDPNKTQWRAVVPVPANKTTAYKVSVLRDRVTITPAP |
| Other | - | MRYALLAGFAAIALSACSQGADEAPADPVENAWVRLPAVDGRPAAAYFTLHGGDAGDTLIAVDSERVASIELHETTMEGGAMRMRPIMSVDIPAGETIAFEPGGRHAMLFGVDPEVTAGTNLTLHFRFDSGRDVSIEAATIDAGDDAPAQESGQDHGAH |
| Other | - | MRIVGVLGCLLVLGACGSEPEVGCTMIGTPVGIGLDVAMPEVRTATMQVCWDDKCVDPVVELHPSSSAGPQTCTGTAPTDSCGVSVVPTGGRNGFADVPDLPTAPVRVTLRLSDANGVVLVDEVLTATPKPAYPNGVECGGQGPQTGLVVSAAGVVTERT |
| Other | - | MKIMRLAAAAALLTLAACSTSPTAPAPAASSPATRSDLTNGQGMIGGGTRAEP |
| Other | - | MKAVIIILASATLFTGCVTHIPVGHHHGHSVIVKKKHGHHRHHHRGHRKHIRIIR |
| Other | - | MQKNIYTRYLSSCLFVGCAHSPQSHSSLCSRGFAPLPSRSILKSIGYDTISIQLT |
| Other | - | MRRVPAGLGVLPALTGCSGLQEAAQDEGQDGRKASVKGGVTSVGLAFDGKDRVGAC |
| Other | - | MKSIILVLVSVALLSGCQLTRVEGEVDDMEVKVGTKENNDDNGKFCPPGQKKKGKC |
| Other | - | MNKLSILLALAAALAVAACDNNIPPTKAPGTADQYPQVDPAPNLTSGGDQSGTPPAQ |
| Other | - | MKLLAAFLALAILSACAARARVPPAPVAAPPAAVQPAVEPLPAGVKPLTVEQDEQISQG |
| Other | - | MTPVRPIFTAFACAALLALGACAARKPSEAEVLTVACSAGIAEACGYLSRGRSVTGTPL |
| Other | - | MKRIMRTWGLLAGLLWLSGCVAYVPYGHRPAGYYAPHGGYGYDAGPRHGGGWHHRHRHHR |
| Other | - | MRGALPLLLILAASGCGDRRTFDDRFNDTQANLQKRARTLDEQLNHQADETANNQQAPTGR |
| Other | - | MKTRGLWLLAALALASCSDDSPPPTPEQPQGLSSCLESPTQLARPPSGQLPCELLPPGFGT |
| Other | - | MRILIPWALSALLLAGCSALPTASERQAHATGLAASKGWLPSHIPAGNFDLMAYAPPSLAAW |
| Other | - | MTQFKRLKRAKAGAIVALLSLLCGCTLGGAAAGGYAGNRLTHGSAVGTVGGAVAGGIVGHEVGEHW |
| Other | - | MMFRKLLAAAGCACALAGCSAAVAVADATVTVVATGVKVTAKTVGAVADAVIPDGEDDAAQSAPQD |
| Other | - | MVNAALKKITPLLVLAVMLAGCQSHEGPAEKAGKKIDNAASQVKKDVGNATDKAGKKIEDAGNSIKNN |
| Other | - | MTKSMTRSLFLAATFAGVVGLAGCAGNGYDSGASTSTDASAQAPMTSDGSTAAPGTTAQPMPTSPAPTAQP |
| Other | - | MNKKIKIGGLSLLTAAFFIFNTGCKKRETTLDSKPALHKLNGTLSLSTKNLKSYHYAYYGNEKIELNSELPKCL |
| Other | - | MTKKECNATHDLMNRSPAIAAALVAALTSGCVQVSAPEEPIVIELNINVQQTVDVNLQEDVENLIENNPELFPQ |
| Other | - | MKKITLMFMVVWGLSGCGGSGDDHASNNTQTQSDPVLRETLTLTNQSQSSFSQSEPKSLKDISEASIDNKAEPLTVKF |
| Other | - | MRLVSLGTRWTGAVWAPALTAGCTKNDNPVSAELLRGPGHADNASSHGGLGPAPSPNAGTAPPGAVPLDPAGKRIRLRC |
| Other | - | MAKKMTIVALPLAFALTTSIVSGCSNEESHPKTKQTPASKQQTSTSERKLQQYAKDPDAKKTDEDFDLVGVVKSVKPLKKSH |
| Other | - | MKTTLIVLLSLCILSACQTADVIDTAGHVLTPTEIRHDGAPNEVVRFGVNACGHLKAQCRVEHYQEWPALFRDVGCRCDRSH |
| Other | - | MIRTSGIALAALYLIGCAGDSPPTSGECRALLQSRTATRAEMVAFIVRAKRDLDAAPRNRIDSHTEGLGLCIGALMRKAQSIK |

| | | |
|---|---|---|
| Other | - | MRFRMVIAPCILMLGLGLGGCTRSCEDLAAEARAKSADAMHMAADAYADGASTEKIAAKMSNLQKDMIDLRNEMAGKNCPMTF |
| Other | - | MKLSFKSLVLFSAFAAFATTSCSERQQENTEAATENAVDQAGDATSEAANEVQADMATEPGDTAVVQNKEADKLVEKVPATPQN |
| Other | - | MTRSSVVGSVLVLGLAIALAGCTTVSGPGGAVARQGAGGGAGGGEGVARRVAKDSASEKARARITAQRPAAAPSRRMMVIGAGW |
| Other | - | MTTRLSTMLALSLVLTLVGCNQQNFQYDARYQLTEVDGVLYRLNQKTGQVDRLEQDRFVPMKEPLPDTMADAVQGQAEQESDGE |
| Other | - | MKPTHLIPAALLLALAACSKVTVANYDKLRAGQSYDEVQQLLGKPSECDDVLSARNCVWRSGKATVNVSFVGGQVILFTAQGLR |
| Other | - | MRLFAALPCLTLLATLAACTGDAPASNELQLENDGGGKFSGKAGPEWTGAELKQEAATSVCGGAEPATFKLSRKKDVWSFKGKC |
| Other | - | MFRRPGRAFAAVGLLLLLSGCGQTGPLYMPPPEAPAAPPAAESTPPAQPAQ |

**Table S7. Protein sequences of predicted autoinducing peptide-lipoproteins.** Sequences are grouped by cluster as shown in the SSN in Fig. S27).

| Cluster | Annotation | Sequence |
|---|---|---|
| I | AIP | MKKALAITLSAIGAVLSTVATTGCAFVLFDEPEMPRSLID |
| I | AIP | MKKALAIALSAVGAVLGTVATTGCIFVLFDEPEMPRSLID |
| I | AIP | MKKALAIALSAIGAVLGTVATTGCILALFDEPEMPRSLID |
| I | AIP | MKKRVAKLLVALGMLATASASVGCLWWSLDEPKALKGMD |
| I | AIP | MKKALAIALSAIGAVLGTVATTGCFAVLLDEPEMPRSLID |
| I | AIP | MKKRVAAFVSAIALLATGAASMGCAWILVDEPNNIKAIKD |
| I | AIP | MKKRLSQLLMAVAMLATGAASLGCWWWSVEEPDSVGVFND |
| I | AIP | MKKFLSVALAAIGAVIGTVGTSGCFVLLLDEPEMPASLID |
| I | AIP | MKLFATIAAAIGSLITATATTGCIMVVFDEPEMPESML |
| I | AIP | MKLFATITATLGAFLATIATAGCVIVYVDEPEMPESML |
| I | AIP | MKKLLAMILSGVGVLAAGAATTACVIVLIDEPSMPNKMIEK |
| I | AIP | MKKAFTYVLTAIAILATGAASMGCAIALTDEPKAPKSMMD |
| I | AIP | MKKLLSIGLAAIALLASTASSVGCMWIIIDEPKALKNMD |
| I | AIP | MKKLLSIGLAAIALLASTAASVGCTWVLIDEPKALKNMD |
| I | AIP | MKKALAIALSAIGAILGTVATTGCVVVLLDEPEMPKSLID |
| I | AIP | MKKKLAGVVAAVAMMLTGAASMGCMIFLVDEPVAPRDLID |
| I | AIP | MKLFATIAAAIGSLIAATATTGCIMVVFDEPEMPESML |
| I | AIP | MKKRLASILAAVAIMATGAASMGCIWIYWDEPNSYNLFCD |
| I | AIP | MKKALAIALSAIGAIIGTVATTGCVVVLFDEPEMPKSLID |
| I | AIP | MKKKLAGVVAAVAMMLTGAASMGCMFILIDEPVAPNALID |
| I | AIP | MKKALAITLSAIGAVLGTVATSGCIFAIFDEPEMPRSLID |
| I | AIP | MKKALAIALSAVGAILGTVATSGCIMVLFDEPEIPRSLID |
| I | AIP | MKKALAITLSAIGAVLGTVATSGCIMVLLDEPEMPRSLID |
| I | AIP | MKKALAITLSVIGAVLGTVATTGCLVVLFDEPEMPKSLID |
| I | AIP | MKKLLSISLAAIALLASTAASVGCMWILIDEPEALKNMD |
| I | AIP | MKKALAIALSAIGAVLGTVATTGCIFVLFDEPEMPRSLID |
| I | AIP | MKKALAIALSAIGAVLGTVATSGCIFGLLDEPEMPKSLID |
| I | AIP | MKLFATITATLGAFLATIATAGCVIIYVDEPEIPESML |
| I | AIP | MKKLLAMVLSGVGILAAGAASIACPMLIIDEPEMPKEMIER |
| I | AIP | MKLFATITATIGAFLATVATSGCILAYIDEPEMPESLL |
| I | AIP | MKKLLSIGLAAIALLASTAASVGCTWILIDEPKALKNMD |
| I | AIP | MKLFASIAAFVGSLIAATATTGCIFVVFDEPEMPESLL |
| I | AIP | MKKFLSIALAAIGAVVGTVATSGCIALLLDEPEMPNSLIG |
| I | AIP | MKKALAIALSAIGAVLGTVATSGCIFGLLDEPEMSRSLID |
| I | AIP | MKKFLSIALATIGAVIGTVAYSGCAFLLLDEPEMPDSLID |
| I | AIP | MKKALAITLSAIGAVLGTVATTGCLVVLLDEPEMPKSLID |
| I | AIP | MKKFLSIALAAIGAVVGTVATSGCFVLLLDEPEMPASLID |
| I | AIP | MKKFLSIALAAIGAVVGTVATSGCVLALFDEPEMPASLID |
| I | AIP | MKALFVSLISAVGALVAASATSGCILVWIDEPTMPKSLLER |
| I | AIP | MKKALAIALSAIGAVLGTVATTGCVMVLFDEPDMPKSLID |
| I | AIP | MKKALAITLSAIGAVLGTVATTGCFLALLDEPEMPKSLID |
| I | AIP | MKKLLSIGLAAIALLASTAASVGCMWILIDEPKALKNMD |
| I | AIP | MKLIAKITAFLGALVATTATTGCITVILDEPEMPETLL |
| I | AIP | MKKRMAKLLAAVAMLATASASVGCAWFCVEEPKALKNMD |
| I | AIP | MKKRVAKLLVALGMLATASASVGCIWWTVDEPKALKGMD |
| I | AIP | MKKLLSIGLAAIALLASTAASVGCMWVIIDEPKALKNMD |
| I | AIP | MKKRVAAFVSAIALLATGAASMGCVWMLIDEPNNIKAIKD |
| I | AIP | MKKFLSIALATIGAVVGTVATSGCIIMLLDEPEMPDSLID |

| | | |
|---|---|---|
| I | AIP | MKLFASIAAFVGSLIAATATTGCVFIVFDEPEMPESLL |
| I | AIP | MKLFATIAAAIGSLIAATATTGCILAVFDEPEMPESML |
| I | AIP | MKKRVAAFVSAIALLATGAASMGCVWLLVDEPSNIKAIKD |
| I | AIP | MKKFLSIALAAIGAVVGTVASSGCVLALLDEPEMPASLID |
| I | AIP | MKKALAIALSAVGAVLGTVATSGCIMVLFDEPEIPRSLID |
| I | AIP | MKKRLSQLLMAVAMLATGMASTGCIWFILEEPNSIDTFCE |
| I | AIP | MKLFATIAATVGAFLATISTAGCLIIYVDEPEMPESML |
| I | AIP | MKKRVAKLLVALGMLATASASVGCVWWTVDEPKALKGMD |
| I | AIP | MKLIASILASVGALVASIGTVGCPALICDEPKMPKSLLNK |
| I | AIP | MKKALAITLSAVGAVLGTVATTGCVVIFLDEPEMPKSLID |
| I | AIP | MKKALAIALSAIGAVLGTVATTGCVVLLFDEPEMPRSLID |
| I | AIP | MKKRVAAFVSAIALLATGAASMGCAWMFVDEPNNIKAIKD |
| I | AIP | MKKVLAIALSAVGAILGTVATSGCVMVFLDEPEMPRSLID |
| I | AIP | MKKLLSIGLAAIALLASTAASVGCTWIIIDEPKALKNMD |
| I | AIP | MKKFLSIALAAIGAVVGTLATSGCLFLLLDEPEMPVSLID |
| I | AIP | MKLIAKITAFLGALVAATATTGCITVILDEPEMPETLL |
| I | AIP | MSKVFAKIAAALGGIMAYTTVGACSWGWLEEPEMPKSLIK |
| I | AIP | MKKFLSIALAAIGAVVGTVATSGCFLLLLDEPEMPASLID |
| I | AIP | MKYLSSLLAVLGTLAATLGTQGCSYLICDEPKMPKSLLNK |
| I | AIP | MKKALAIVLSAIGAVLGTVATTGCIFVLFDEPEMPRSLID |
| I | AIP | MKKLFAMVLSGLGILAAGAATSACVCLVIDEPEMSKEMIER |
| I | AIP | MKLFYSILAAIGAVSATIGTQGCVVLLLDEPKMPKSLLNK |
| I | AIP | MKALFVSAISALGALLAATATSGCAIVFIDEPTMPKSMIER |
| I | AIP | MKKRVAAFVSAIALLATGAASMGCIWILADEPNNIKAIKD |
| I | AIP | MKKLFAMVLSGLGILAAGAATSACLCLVLDEPEMSKEMIER |
| I | AIP | MKKRMAKLLAAVAMLLTTTASVGCVLFIVEEPKALKNMD |
| I | AIP | MKKALAIALSAVGAFLGTVATSGCLFVLLDEPEMPRSLID |
| I | AIP | MKLIAKITAFLGALVAATATTGCITIILDEPEMPETLL |
| I | AIP | MKKRVAKLLVALGMLATASASVGCVWWSLDEPKALKGMD |
| I | AIP | MKKRLSQLLMAVAMLATGAASVGCVWWFYDEPDAVNIFIND |
| I | AIP | MKKRVAKLLVALGMLATASASVGCIWWAVDEPKALKGMD |
| I | AIP | MKLFATIAATIGSLIAATATTGCIMVVFDEPEMPESML |
| I | AIP | MKLFATIAAAIGSLIAATATTGCIVAVFDEPEMPESML |
| I | AIP | MKLITSILASVGALVASIGTVGCPALICDEPKMPKSLLNK |
| I | AIP | MKKFLSIALAAIGAVVGTVAYSGCAFLLLDEPEMPASLID |
| I | AIP | MKKRVAKLLVALGMLATASASVGCIWWTIDEPKALKGMD |
| I | AIP | MKKRVAKLLVALGMLATVSASVGCVWWALDEPKALKGMD |
| I | AIP | MKLIAKITAFLGALVAATATTGCITVILDEPEMPKTLL |
| I | AIP | MKKLLSIGLAAIALLASTAASVGCTWVIIDEPKALKNMD |
| I | AIP | MKKRLSQLLMAVAMLATGATSVGCFWVVLEEPDSLDLFKE |
| I | AIP | MKLFYSILATIGAVSATIGTQGCSYFLVDEPKMPKSLLNK |
| I | AIP | MKKLFSVALAAVALLASTAASVGCMWLFIDEPEAPKFMD |
| I | AIP | MKKALAITLSAIGAVLGTVATTGCVMVIFDEPEMPRSLID |
| I | AIP | MTKRVAKLLVALGMLATASASVGCVWWSLDEPKALKGMD |
| I | AIP | MKKLFAMVLSGLGILAAGAATSACLLIVLDEPEMSKEMIER |
| I | AIP | MKKALAIALSAIGAIIGTVATTGCILALFDEPEMPRSLID |
| II | AIP | MKKLMSIARKATTTLAALALVIATTSVPGACYHWFAQPVEPEELRKFMNDK |
| II | AIP | MKKLMSIARKATTTLAALALVIATTSVPSACYHWFAQPVEPEELRKLVNDK |
| II | AIP | MKKLGKLFKLGISLIAALSAIAAMGISVNACHLWFAQPKVPKELDAFRNENK |
| II | AIP | MKKTSKVLRMAASLMAALSAVAAMGISVNACHFWFAQPKVPQGLDKFRSEK |
| II | AIP | MKKLMSIAKKASTALAALALVVATTSVPSACFHWFAQPVEPEELRNYVNRK |

| | | |
|---|---|---|
| II | AIP | MKKLMSIAKKASTALAALALVVATTSVPSACFHWFAQPVEPEELRNYVSKK |
| II | AIP | MSIAKKASTALAALALVVATTSVPSACFHWFAQPVEPEELRNYVNKK |
| II | AIP | MKKLMSIAKKASTALAALALVVATTSVPIACSHWFAQPVEPEELRNYVNKK |
| II | AIP | MKKTSKVLRMAASLMAALSAVAAMGISVNACHFWFAQPKVPQGLDKFRSKE |
| II | AIP | MKKLMSIAKKASTALAALALVVATTSVPSACFHWFAQPVEPEELRNFVNNK |
| II | AIP | MKKIILNKCIFVMSNLVLFIATTSISQCCNTQIYQPSVDEDLKNKIIQANKK |
| II | AIP | MKKLMSIAKKASTALAALALVVATTSVPSACFHWFTQPVEPEELRNYVNKK |
| II | AIP | MKNLMKIARKTSFVLAAMALVVATSSVASCCYHWFAQPVEPEELRKFVGEN |
| II | AIP | MKKLMSIAKKASTALAALALVVATTSVPSACFHWFAQPVEPEELRKYVNKK |
| II | AIP | MKKLMSIAKKASTALAALALVVATTSVPSACFHWFAQPVEPEELRNYVNKK |
| II | AIP | MKKLMSIAKKASTALASLALVVATTSVPSACFHWFAQPVEPEELRNYVNKK |
| II | AIP | MKKLMSIAKKASTALAALALVVATTSVPSACYHWFAQPVEPEELRNYVNKK |
| II | AIP | MKKLMSIAKKASTALAALALVVATTAVPSACFHWFAQPVEPEELRNYVNKK |
| II | AIP | MKKLMSIAKKASTALAALALVVATTSVPSACYHWFAQPVEPEELRKYVNKK |
| II | AIP | MKKLMKLAVKASTVLAAMAMFVATSSVSACCHCCFAQPVEPEELRNLVNNK |
| II | AIP | MKKLMSIAKKASTALAALALVVATTSVPSACFHWFAQPVEPEELRNYVNKE |
| II | AIP | MKKLMSIAKKASTALAALALVVATTSVPSACFHWFAQPVV |
| II | AIP | MKNLMKIARKTSSVLAAMALVAATSSVASCCYHWFAQPVEPEELRKFVGEN |
| II | AIP | MKNLMKIARKTSSVLAAMALVVATSSVASCCYHWFAQPVEPEELRKFVGEN |
| II | AIP | MKKLMSIAKKASTALAALVVATTSVPSACFHWFAQPVEPEELRNYVNKK |
| II | AIP | MKKLMSIAKKASTALAALALVVATTAVPSACFHWLAQPVEPEELRNYVNKK |
| III | AIP | MKNKVLRSLVKVLTVAAFVCASTSSLCNCYEPEKPASLR |
| III | AIP | MKYRILNSMVKVLTLVAFVCAMSPSRCNCYEPEKPASLR |
| III | AIP | MKNKVLRSLVKVLTVAAFVCASTSSLCNCYEPEKPVSLR |
| III | AIP | MKNKVLRSLVKVLTVAAFVCASTSSLFNCYEPEKPASLR |
| III | AIP | MKYRILNSMVKVLTLVAFVCAASPSHSNCYEPEKPASLR |
| IV | AIP | MKKLNKNLLTLFAALTTVVATTVATSACIWFTHQPEEPKSLRDE |
| IV | AIP | MKKLNKNLLTLFAALTTVIATTVATSACIWFTHQPEEPKSLRDE |
| IV | AIP | MKKKLFMTVATVATLVASFVATSACWACFYQPEEPKSLRDE |
| IV | AIP | MKKKILMSLAAISTFIASIVATSACTWYFYQPEEPECLRDK |
| IV | AIP | MKRRILMVVATAATLIASIVSTSACIWGHYQPEEPKCLREE |
| IV | AIP | MKKKILMGIAAVATVFASVISTSACYFSFYQPEEPECLREE |
| V | AIP | MEKFKQYSLKTFAALLSFVAISSVNSACAVIFGQEKEPDSLAKYKKC |
| V | AIP | MSKKDFGKKFLNHLASLALIAGVISSSTACRFVLHQPAMPDKMKKLLGEK |
| V | AIP | MKKEKFIKSFANVLAAVAVTVTSGVATTGCFAFLHQPELPKGAEKLRKFK |
| V | AIP | MQKFKNFIMKFGSSFAVLALIIGISTSNSACGMIFHQPKEPAAMNKFKF |
| V | AIP | MKKMMLFLSKFTCIMAMALAVLSVNSTCGFTAYQPDVPESLNHR |
| V | AIP | MQKIKNFIMKFGSSFAALALIIGITTSNSACGMIFHQPEEPAAMNKFKF |
| V | AIP | MRKKGFGKKFLNHLASLALIAGVISSSTACRFVLHQPAMPDKMKKLFSEK |
| V | AIP | MQKFKNFIMKFGSSFAALALIIGISTSNSACLMVFHQPKEPAAMNKFKF |
| V | AIP | MEKFKQYSLKTVAALLSFVAILSVNSACAVIFGQEKEPDSLAKYKKY |
| V | AIP | MKKEKFIKGFANVLAAVAVTVTSGVATTGCFAFLHQPELPKGAEKLRKFK |
| V | AIP | MQKIKNFIMKFGSSFAALALIVGISTANSACDIIWHQPKEPTAMNKFKF |
| V | AIP | MKKIMLLLSKVTCIMAMALAVLSVNSTCGFTAYQPDVPESLEHD |
| V | AIP | MKKSMNFIKKLSTVVVTCLTLVLTINANTASCFILNEPEEPKNIEKFKMFK |
| V | AIP | MEKFKQYSLKTVAALLSFVAISSVNSACAVIFRQEKEPDSLAKYKKY |
| V | AIP | MQKIRNFIMKFGSSFAALALIIGISTSNSACGMIFHQAKEPEAMNKFKI |
| V | AIP | MKKEKFIKGFANVLAAVAVTVTSGVAITGCFTFLHQPELPKGAEKLRKFK |
| V | AIP | MKKFKNIIKKISPAIVGCLTLVLTINANTASCFILNEPKEPKTIEKFKLRK |
| V | AIP | MKKMMLFLSKFTCIMAMTLAVLSVNSTCGFTAYQPDVPESLNHR |
| V | AIP | MEKFKQYSLKTVAALLSFVAILSVNSACAVIFGQEKEPDSLAKYKKC |

| | | |
|---|---|---|
| V | AIP | MQKIKNFIMKFGSTFAALALIVGISASNSACNILWHQPEEPAAMNKFKF |
| V | AIP | MRKKGFGKKILNHLASLALIAGVISSSTACRFVLHQPAMPDKMKKLLGEK |
| V | AIP | MEKFKQYSLKTFAALLSFVAISSVNSACAVIFGQEKEPDSLSKYKKY |
| V | AIP | MKKEKFIKGFANVLAAVAVTVTSGVATTGCFTFLHQPELPKGAEKLRKFK |
| V | AIP | MRKKGFGKKFLNHLASLALIAGVISSSTACRFVLHQPAMPDKMKKLLGEK |
| V | AIP | MLIFGKFTCVMAMALAILSVNSTCGFTAYQPDVPESLNNE |
| V | AIP | MKKTKNIIKKISPVIVGCLTLVLTINANTASCFILNEPKEPNLINKFKIFK |
| V | AIP | MKKIKNLIKRISPVLTACLTLILAINANTASCFILNEPKEPNEIQKFKIFK |
| V | AIP | MLKKLTIQSAKILAVLALAVGTISGQVACSGVYYQPKMPEKMKK |
| V | AIP | MKKKGFGKKFLNHLASLALIAGVISSSTACRFVLHQPAMPDKMKKLLGEK |
| V | AIP | MVYNFKKICLSLFARLLTLLTIIGVNSACNIVYGQPNEPQSLARYKKR |
| V | AIP | MKKIKSLLKKFSPIMVSCLTMVLAINANTASCFLINEPQEPSSIENFKKFK |
| V | AIP | MKKIKRFLSIISPAITGCLTLALAINANSASCFIINQPKEPETIKKFKMFK |
| V | AIP | MKNCKQYFLKTFAFLLSFLAISSVNSACAVLFGQEKEPDSLARYKKC |
| V | AIP | MKKGKKFFLNSFAFLMSFLAISSVNSACAIVFGQEEEPESLKRYKKIK |
| V | AIP | MEKFKQYSLKTVAALLSFVAISSVNSTCAVIFGQEKEPDSLAKYKKY |
| V | AIP | MEKFKQYSLKTVAALLSFVAISSVNSACAVIFGQEKEPDSLAKYKKY |
| V | AIP | MKKIILKYGGMLAALAMVFTTLTVNSTCTWMTYQEELPETAKKLRKF |
| V | AIP | MEKFKQYSLKTVAALLSFVAISSVNSACAVIFGQEKEPDSLSKYKKY |
| V | AIP | MQKFKNFIMKFGSSFAALALIIGISTSNSACMMVFHQPKEPAAMNKFKF |
| V | AIP | MKNGKKFFLNSFAFLMSFLAISSVNSACAIIFGQEEEPESLKRYKKIK |
| V | AIP | MTYNLKKLMLNAFAGLLTLLAIVGVNSACNIVFGQPEEPESLSRYKKIKG |
| V | AIP | MEKFKQYFLKTVATLLSFVAISSVNSACAVIFGQEKEPDSLAKYKKC |
| V | AIP | MEKFKQYFLKTVAALLSFVAISSVNSACAVIFGQEKEPDSLAKYKKY |
| V | AIP | MKTHNKKWMLIFGKFTCVMAMALAILSVNSTCGFTAYQPDVPESLNNE |
| V | AIP | MKKIILKYGGMLAALAMVFTTLTVNSTCTWMTYQEELPDTAKKLRKF |
| VI | LptM chaperone | MSKREKMKKLLSVFLLSAVCALVSGCGVKGPLYFPEKEPAQQQVK |
| VI | LptM chaperone | MKKLLSILLLSAFCTLATGCGVKGPLYFPEKEQPQNTQNTQ |
| VI | LptM chaperone | MKKLLSILLLSTFCTLATGCGVKGPLYFPEKEQPQNTQ |
| VI | LptM chaperone | MKKLLSILLLSAFCTLSTGCGVKGPLYFPEKEQPQNTQKTQ |
| VI | LptM chaperone | MKKLLSILLLSAFCTLATGCGVKGPLYFPEKEQPQSTQKTQ |
| VI | LptM chaperone | MKKLLSILLLSAFCTLATGCGVKGPLYFPEKEQPQNTQKTQ |
| VI | LptM chaperone | MKKLLSILLLGAFCTLATGCGVKGPLYFPEKEQPKNTQ |
| VI | LptM chaperone | MKKLISVFLLVTLCALSTGCGVKGPLYFPEQDSAQKTNK |
| VI | LptM chaperone | MKKLISLFLLVTLCALSSGCGVKGPLYFPEQDTAQQATK |
| VI | LptM chaperone | MKKLLSILVLSTICIMATGCGVKGPLYFPEKEQAQQTK |
| VI | LptM chaperone | MKKVLSILLLSAFCTLATGCGVKGPLYFPEKEQPQNTQKTQ |
| VI | LptM chaperone | MKKLLSILLLSAFCTLATGCGVKGPLYFPEKEQPQNTQ |
| VI | LptM chaperone | MKKLLSVLLLSAFCTLATGCGVKGPLYFPEKEQPQNTQKTQ |
| VI | LptM chaperone | MKKLLSILVLSAICILATGCGVKGPLYFPEKEQAQQTK |
| VI | LptM chaperone | MKKLLSILLLSVFCTLATGCGVKGPLYFPEKEQPQNMQKTQ |
| VI | LptM chaperone | MKKLLSILLLSTFCTLATGCGVKGPLYFPEKEQPQNTQKTQ |
| VI | LptM chaperone | MKKLLSILLLGAFCTLVTGCGVKGPLYFPEKEQPQNTQKTQ |

| | | |
|---|---|---|
| VI | LptM chaperone | MKKFISLLLLGAICVLSTGCGVKGPLYFPEQEQASNN |
| VI | LptM chaperone | MKKLISVFLLITLCALSTGCGVKGPLYFPEQDSAQKTSK |
| VI | LptM chaperone | MKKLISVFLILTLCALSTGCGVKGPLYFPEQDSAQKTSK |
| VI | LptM chaperone | MKKLLSILLLGAFCTLATGCGVKGPLYFPEKEQPQKTQKTQ |
| VI | LptM chaperone | MKKLLSILLLSAFCTLAAGCGVKGPLYFPEKEQPQNTQKTQ |
| VI | LptM chaperone | MKKLLSILLLSAFCTLATGCGVKGPLYFPEKEQLQNTQKTQ |
| VI | LptM chaperone | MKKLLSILLFSAFCTLATGCGVKGPLYFPEKEQPQNTQKTQ |
| VI | LptM chaperone | MKKLLSILVLSAICIVATGCGVKGPLYFPEKEQAQQTK |
| VI | LptM chaperone | MKKLISVFLLVTLCALSIGCGVKGPLYFPEQDSAQKTSK |
| VI | LptM chaperone | MKKLLSILLLGAFCTLATGCGVKGPLYFPEKEQPQNAQKTQ |
| VI | LptM chaperone | MKKVLSILLLGAFCTLATGCGVKGPLYFPEKEQPQNTQKTQ |
| VI | LptM chaperone | MKKLLSILVLSAVCIVATGCGVKGPLYFPEKEQAQQTK |
| VI | LptM chaperone | MKKLLSILLLSAFCTLATGCGVKGPLYFPEKEQPQNTQKHNKN |
| VI | LptM chaperone | MKKLLSILVLSAICIAATGCGVKGPLYFPEKEQAQQTK |
| VI | LptM chaperone | MKKLLSILVLSAVCIAATGCGVKGPLYFPEKEQAQQTK |
| VI | LptM chaperone | MKKLLSILVLSAICIMATGCGVKGPLYFPEKEQAQQTK |
| VI | LptM chaperone | MKKLISVFLLVTLCTLSTGCGVKGPLYFPEQDSAQKTTK |
| VI | LptM chaperone | MKKLLSILLLGAFCTLATGCGVKGPLYFPEKEQPQNTQKTQ |
| VI | LptM chaperone | MKKLLSILLLGAFCTLATGCGVKGPLYFPEKEKPQNTQKTQ |
| VI | LptM chaperone | MKKLISVFLLVTLCALSTGCGVKGPLYFPEQDSAQKTSK |
| VI | LptM chaperone | MKKLLSILLLSAFCTLATSCGVKGPLYFPEKEQPQNTQKTQ |
| VI | LptM chaperone | MKKLLSILLLSAFCTLATGCGVKGPLYFPEKEQSLNTQKTQ |
| VI | LptM chaperone | MKKLISVFLLITLCVLSTGCGVKGPLYFPEQDSAQKTSK |
| VI | LptM chaperone | MKKLLSILLLSAFCALATGCGVKGPLYFPEKEQPQNTQKHNKN |
| VI | LptM chaperone | MKRSVSILLLGAFCTLATGCGVKGPLYFPEKEQPQKTQKTQ |
| VII | TolC-subunit | MNTSKIIACGATLLLSSCGIYTSYEPQTSVPENLYGEEVAEAVSNKL |
| VII | TolC-subunit | MNTLKIIACGATLLLSSCGIYTSYEPQTSVPENLYGEEVTEAVSK |
| VII | TolC-subunit | MNNMNTSKIIACGATLLLSSCGIYTSYEPQTSVPENLYGEEVTEAVSK |
| VII | TolC-subunit | MNNMNTSKIIACGATLLLSSCGIYTSYEPQTSVPENLYDEEVTEAVSK |
| VII | TolC-subunit | MNNMNTSKIIACGATLLLSSCGIYSSYEPQTSVPENLYGEEVTEAVSK |
| VII | TolC-subunit | MKKQIITLTVAALTLSSCGIYTQYKPATEVPDNL |
| VII | TolC-subunit | MKKLIISLAAILALSSCGIYSKYKPVTEIPD |
| VIII | AIP | MNKWIYRVCTKLAACAFLAAIVSVGTASCNGMYQPEIPSQLK |
| VIII | AIP | MKKIMYQVCTKLAACAFLAAIVSVGTASCNGLYQPKMPDKLVK |
| VIII | AIP | MKKIIYQVCTRLAACAFLAAVVSVGTASCGGLYQPEVPSKPVK |

| | | |
|---|---|---|
| VIII | AIP | MKKIIYQVCTKLAACAFLAAIVSVGTASCNGMYQPAVPKQLNKQS |
| Other | N/A | MTKSINVFSAILLNLFNLVSKSGCSSMWGEPDYPEELL |
| Other | N/A | MSEKLRKFLVMAAGVLVILETTVISGCRANISGTESDRMVEYTVICGDEIPEELKSR |
| Other | N/A | MRKYIQIISFLLAAVTLSGCENWLDVKPETEVNG |
| Other | N/A | MRKNFLWIMATAIIFTSSCAKAPEEVKKGK |
| Other | N/A | MRKFKSLITVLAITVTSAALFTGCGNTGSNDSNSSSQLQDESK |
| Other | N/A | MRIKRLLYAIATILPFLFLCSCYEEQEPQQEKQDKEK |
| Other | N/A | MQKLLNCFSTLLINIFGLVSNTSCAGYYGEPDYPEELLK |
| Other | N/A | MPMERHEQTNSIKKMDYRNMKGIFLAAALAAVFTTGCWAQQPASPDFVTQKN |
| Other | N/A | MNMKKIASVVLALALSVSVFAACGSNGNSSASSTACLLYTSDAADEL |
| Other | N/A | MNLKKIFFSAVTVSVLCALTGCDYIEEGKPESSLLKQQEEH |
| Other | N/A | MNKKNFSGGITFALVVSLSIVTSGCSSNIDDPTDAQNLC |
| Other | N/A | MNKFSIVLTLLLCGSCALALDPNLEKN |
| Other | N/A | MNIKVDLKKLFAAAVSAIMCLSAGGCYLLPDEEEVLDAP |
| Other | N/A | MNHLSSLIRLVFLPIVLIACNSTVEKPQEDKLQELQKK |
| Other | N/A | MNDYQKKVLSLALSVSLLFNASCNKDKVIYNLPPEPIKSEEPKELKLKPKKIDNE |
| Other | N/A | MNDYQKKVLSIALSVSLLFNASCNKGTVVYNLPPEPIKSEEPKELKIKPKKIDEE |
| Other | N/A | MMKKYLFLLLIGAASLTACGDDDDLVVPELDKLT |
| Other | N/A | MKTFFKTSCKKIAAILPAIALFFAVSPCVGKIYEPKLPEQLK |
| Other | N/A | MKTELARKLLSGLSLWVLLFALSSCGSKMDEDAAAAYFDESPAVTLYATCFFDEEQPNLKADFSVKF C |
| Other | N/A | MKSKIFYACLFAASVLTSCSDFLEEKPKGVL |
| Other | N/A | MKRNLKLALLMAFCLTATACGKTAEKPAEKRLKSKP |
| Other | N/A | MKQKLAVLLSLVMTISLTACSSAGTYSKSSTFASDDVK |
| Other | N/A | MKNLVKNFIVKHSKLLCSLMVVSAYASLTSCRFLYFQPEVPEGLDKIVK |
| Other | N/A | MKNLVKNFIVKHSKLLCSLMVVSAYASLTSCRFLYFQPEIPEGLDKIVK |
| Other | N/A | MKNLKNIFTVALIATCMSGCTGFLDIEPETTLTGKNFYTRQYGIDKS |
| Other | N/A | MKNLKKYIIIDILASVIIYLSGCASLTYNPHSQTEKVKVTLKNK |
| Other | N/A | MKNLKKYIIIAILASVIIYLSGCASLTYNPHSQPEKVKVTLKNK |
| Other | N/A | MKNLFHIAKKTFTWVTVTMAVSLFNGCSDFLDKQP |
| Other | N/A | MKNKFTHFLMAGALVCWSSYTLHAQTEVPMSVKR |
| Other | N/A | MKNKALVWMCGLLSAIMAFTSCTNFLEEDPKTFLSPSEY |
| Other | N/A | MKNKAHVWMCGLLSATMAFTSCTNFLEEDPKTF |
| Other | N/A | MKLSNKLNSLLVAPLAIALFAACSTDENLSDGG |
| Other | N/A | MKLSLVKYILLFCGVIPLLTSCESFLDKQETEDLTFE |
| Other | N/A | MKLKNILLCASVLGGLALSSCTGNFLDEDRNPNSLDP |
| Other | N/A | MKLKKLVPFVISVLMIFQVCVPFTFADEDPP |
| Other | N/A | MKLKKIIACAAAFAMTATCFAGCGSTSDSAGAK |
| Other | N/A | MKKYLFLLLIGAASLTACGDDDDPVVPELNKLTKS |
| Other | N/A | MKKVLVSSMLILGGLFSACSGFLDEDPKSKIPEEEAYKSEKLV |
| Other | N/A | MKKVILIAVALLGALTVSSCGPGYQPLSEEEWKKQQK |
| Other | N/A | MKKSKFKLFAALLLAAAMTVLSLSSACLLYTSDAADE |
| Other | N/A | MKKSDLFKIGVLLVATTLGTTGCSFGEDEKKPEIVVDPAEKTIE |
| Other | N/A | MKKSDLFKIGVLLMATTLGTTGCSFGEDEKKPEIVVDPAEKQ |
| Other | N/A | MKKRKIAMLVMTGVLLTGCMGTNYLEEGVSQLEEKQ |
| Other | N/A | MKKNKLLTALALASACLLYTSDAADE |
| Other | N/A | MKKLTALLLALVMALSLTACSFVMLPEEQEPPADPKQE |
| Other | N/A | MKKLLALLMALMMCCTAFAFAEEEAEVPAVEMNVSFEA |
| Other | N/A | MKKLILFVSKNLCNVAVFAAAFAVTTSCSRWFYQEELDEQVMSLNRYRD |
| Other | N/A | MKKLILFVSKNLCNVAVFAAAFAVTASCSRWFYQEELDEQVMSLNRYRD |
| Other | N/A | MKKLFLFPVMLLFALTITACSNSPNGFEEPETCLLYTSDAADE |

| | | |
|---|---|---|
| Other | N/A | MKKKYFACAMGLVLALSLAACGEEEVVVENTNEEVTQEVQTQEVQTQASCCIVYQPKAPKELERYRKTK |
| Other | N/A | MKKKMGAFILCFALLGAGCVFAEEEKTSP |
| Other | N/A | MKKKLAFKLATLMCAVTSIGQIATCAAPNSLLLWGEPTPPMKKEDK |
| Other | N/A | MKKKFIYSFMIAAAAAAFSSCSNFLEERPTDAFDEETAF |
| Other | N/A | MKKIYRLLLTFMPALVLYTGCYAFFQEKIPMDISKDSSSLTDLKVP |
| Other | N/A | MKKINLLSALFLCTILASCASENTSSSSSYSSSSTISSS |
| Other | N/A | MKKILLALLTSCALVSCEGYFDQLPKTELPS |
| Other | N/A | MKKILFAFMAMYCFIACDPSDSFADNE |
| Other | N/A | MKKFLSVLLMVAGLMSCGDDDKPFIPELNKR |
| Other | N/A | MKKFLSLILILMLMTLSTGCAKEEEPECLSGTCPFYPTETLKVN |
| Other | N/A | MKKFLSLILILMLMALSTGCAKEEEPECLSGTCPFYPTETLKVN |
| Other | N/A | MKIKSILKLGTLALLTAALPACSFLDTDPQINPDDGYYNS |
| Other | N/A | MKHLSSILLLAFVCTMFASCYDEPDPAEREHDSNLIGE |
| Other | N/A | MKFKNIMKSCLSLCVMLVAYCSVNTACTLLVYQDELPEGVQRMREAKFK |
| Other | N/A | MFNLKYIENMKTELARKLLSGLSLWVLLFALSSCGSKMDEDAAAAYFDESPAVTLYATCFFDEEQPNLKADFF |
| Other | N/A | MFKIAKKLTVIMLSASIIFSCSGIIAHAEGIKELLEKQMKK |

### Table S8. Strains and plasmids used in this study.

| Strain/plasmid name | Purpose |
|---|---|
| 5alpha (NEB) | E. coli strain for cloning plasmids |
| BL21 (DE3) | E. coli strain for protein expression |
| pACYCDuet-1:His-ChrA | For expression of His-ChrA |
| pACYCDuet-1:His-DybA | For expression of His-DybA |
| pACYCDuet-1:His-ChsA | For expression of His-ChsA |
| pACYCDuet-1:His-HymA | For expression of His-HymA |
| pRSFDuet-1:His-ArlA | For expression of His-ArlA |
| pACYCDuet-1:His-MelA1 | For expression of His-MelA1 |
| pCDFDuet-1:ChrH | For expression of untagged ChrH |
| pCDFDuet-1:DybH | For expression of untagged DybH |
| pCDFDuet-1:ChsH | For expression of untagged ChsH |
| pCDFDuet-1:HymH | For expression of untagged HymH |
| pRSFDuet-1:MCS1_His-ArlA_MCS2_ArlH | For coexpression of His-ArlA with untagged ArlH |
| pCDFDuet-1:MelH | For expression of untagged MelH |
| pCDFDuet-1:MCS1_MelH_MCS2_MelHc | For coexpression of untagged MelH and MelHc |
| pCDFDuet-1:MCS1_His-MelH_MCS2_MelHc | For coexpression of His-MelH and untagged MelHc |
| pET28a:His-TEV-DybH | For expression of His-DybH |
| pRSFDuet-1:MCS1_His-DybA_MCS2_DybH | For coexpression of His-DybA and untagged DybH |
| pRSFDuet-1:MCS1_DybA-His-embed_MCS2_DybH | For coexpression of DybA (His-tag embedded at residue 21) with untagged DybH |
| pRSFDuet-1:NedA-C-His | For expression of C-terminally His-tagged NedA |
| pCDFDuet-1: MCS1_NedB_MCS2_NedC | For expression of untagged NedB and NedC |

**Table S9. Sequences and accession IDs of proteins characterized in this study.** Bolded sequences indicate added purification tags. Codon opt. = codon optimized for *E. coli.*

| Protein name | NCBI accession no. | Organism | Protein sequence (bold indicates added tag) | Coding sequence | Codon opt.? |
|---|---|---|---|---|---|
| DybA | WP_157488328.1 | Dyadobacter crusticola strain DSM 16708 | M**GSSHHHHHH**KIS<br>KPVLQAVAVAVALS<br>TITACADMDIKPEK<br>DKKAKTRTLDPCPA<br>CGMG | ATGGGCAGCAGCCATCACCATCATCACCACAAAATCTCAA<br>AACCAGTACTTCAGGCCGTTGCCGTTGCAGTTGCACTGAG<br>CACAATTACCGCTTGCGCCGACATGGATATTAAACCTGAA<br>AAGGATAAAAAAGCGAAGACGCGCACCCTGGATCCTTGC<br>CCGGCCTGCGGAATGGGTTGA | No |
| DybH | WP_031530742.1 | Dyadobacter crusticola strain DSM 16708 | MSQIFSTVACNLDA<br>HILQASIPLFEMERV<br>EAIEWSFDALFKVD<br>VIPEWFTDFIKEFSD<br>QNRLIGHGVYFSLF<br>SGKWSAEQQKWL<br>DQLSKLCDQFQFD<br>HITEHFGFMTGSDF<br>HKGAPLGIPFNAST<br>LAIGKDRLARIQNA<br>CNCPVGLENLAFSY<br>SLDEVKRHGDFLA<br>QLVESVNGFIILDLH<br>NLYCQCHNFELDFI<br>DFLSFYPLERVREM<br>HISGGSWDTQIPGR<br>VVRRDTHDEAVPS<br>EVFEYLRQAIPRCP<br>NLKFVVMEQLGTAL<br>DTAEKQAGFQKDF<br>LEMDRIVKSVGATL<br>PDSVKNFQPLSQQ<br>VQIGPPIEDLTLHQ<br>QQMQLSEILEKARD<br>LHEAKNLLLASDLS<br>NTDWHIERWEPHM<br>LQTAMNIAQKWKA<br>GF | ATGTCGCAAATATTCTCCACAGTCGCCTGTAACCTCGACG<br>CGCACATTCTGCAAGCGTCTATTCCATTGTTTGAAATGGA<br>GCGAGTGGAGGCTATTGAATGGTCTTTCGATGCGCTCTTC<br>AAAGTGGATGTGATACCGGAATGGTTTACTGATTTTATAAA<br>AGAATTCAGTGACCAGAACAGGCTGATTGGTCACGGCGTT<br>TACTTTTCGCTGTTTTCGGGAAAATGGTCGGCAGAGCAGC<br>AAAAATGGCTGGATCAGCTCAGCAAGCTCTGCGACCAGTT<br>TCAGTTTGATCACATCACGGAGCATTTCGGCTTTATGACC<br>GGTTCTGACTTCCACAAAGGCGCGCCGCTCGGCATTCCC<br>TTCAATGCATCTACATTAGCAATCGGAAAAGACAGGCTGG<br>CGCGCATTCAGAATGCGTGTAATTGTCCGGTTGGATTGGA<br>AAACCTGGCTTTTTCCTATTCACTGGATGAGGTAAAACGG<br>CATGGAGACTTTTTGGCCCAACTGGTGGAAAGTGTCAACG<br>GGTTCATCATCCTCGACCTGCACAACCTGTATTGCCAATG<br>CCATAATTTTGAGCTCGACTTCATCGATTTCCTTTCGTTCT<br>ATCCACTTGAACGCGTGCGGGAAATGCATATCTCCGGCG<br>GGAGCTGGGACACACAGATTCCGGGAAGGGTAGTACGGC<br>GCGACACCCACGACGAGGCAGTACCATCGGAAGTATTCG<br>AATACCTCAGGCAGGCAATCCCGCGCTGCCCTAACTTGAA<br>GTTTGTAGTAATGGAGCAGCTGGGCACAGCACTCGATACA<br>GCTGAAAAGCAGGCTGGGTTTCAAAAGGACTTTCTTGAAA<br>TGGACCGCATTGTCAAATCCGTCGGAGCAACCCTGCCCG<br>ATTCAGTAAAGAACTTTCAGCCTCTTTCACAACAGGTGCAA<br>ATTGGCCCGCCGATCGAAGACCTTACTTTGCACCAGCAG<br>CAAATGCAATTGTCGGAAATCCTGGAAAAAGCACGGGACC<br>TGCATGAGGCAAAAAACCTCTTGCTTGCGTCTGACCTCAG<br>CAATACAGACTGGCACATTGAACGTTGGGAGCCGCACAT<br>GCTGCAAACAGCGATGAACATTGCGCAGAAATGGAAGGC<br>AGGATTTTGA | No |
| ChrA | WP_034727518.1 | Chryseobacterium sp. JM1 | M**GSSHHHHHHSQ**<br>MKIPALVMASLLAV<br>SVSGQTTKPVKKG<br>TKSVKKVKKMDSP<br>KTVKAEPTKVVKRD<br>TILKHGGGCPACG<br>MG | ATGGGCAGCAGCCATCACCATCATCACCACAGCCAGATG<br>AAGATTCCGGCTCTGGTGATGGCCAGCCTGCTGGCGGTA<br>TCAGTGAGCGGGCAAACCACGAAACCGGTCAAAAAAGGC<br>ACGAAAAGTGTTAAAAAAGTGAAAAAAATGGATAGCCCAA<br>AAACCGTTAAAGCAGAACCCACCAAAGTCGTTAAACGTGA<br>CACGATCCTGAAACATGGCGGCGGTTGTCCGGCGTGCGG<br>TATGGGGTGA | Yes |
| ChrH | WP_034727520.1 | Chryseobacterium sp. JM1 | MKKPLLGLAMMPE<br>ADFVSAILPLLQTQ<br>SVDVLEWSFDTLY<br>DVEEPEWLSGLLD<br>FYSDNSRLLGHGV<br>YYSLFDARWMERQ<br>EIWLKKLKEEVKRR<br>KYNHITEHFGFMNT<br>ENFHQGVPLPVPLL<br>PKTLQIGKDRLSRL<br>QDAVEIPVGVENLA<br>FSFSMDDVKEQGE<br>FLDRLVEDIDGFLIL<br>DLHNIYCQSCNFKV<br>DMMEIINLYPLEKV<br>NEIHLSGGSWQES<br>AYGKKPVRRDTHD<br>DRIPEEILNLLPEVLI<br>QCHPEYVIIERLGH<br>TLNTEVAKQIFFDD<br>FNRVKKILELSDYP | ATGAAGAAACCTCTCCTTGGATTAGCCATGATGCCGGAAG<br>CTGACTTTGTTTCCGCTATTCTTCCGCTTTTGCAAACGCAA<br>AGTGTTGACGTACTCGAGTGGAGCTTCGATACGCTGTATG<br>ACGTCGAAGAGCCGGAATGGCTGTCCGGACTGCTTGACT<br>TCTATTCAGACAATTCACGCCTGTTGGGACACGGGGTGTA<br>CTATTCATTATTCGACGCCCGCTGGATGGAGCGCCAAGA<br>GATTTGGCTCAAGAAGCTTAAAGAAGAGGTGAAGCGCCG<br>CAAGTACAATCATATCACCGAGCATTTTGGCTTCATGAACA<br>CTGAGAATTTCCACCAAGGCGTACCTCTTCCTGTACCACT<br>TCTGCCGAAGACGCTGCAAATTGGGAAGGATCGCCTGTC<br>TCGTCTCCAGGACGCCGTGGAGATCCCGGTGGGTGTCGA<br>GAATCTGGCATTCTCTTTCAGCATGGATGACGTCAAGGAG<br>CAAGGTGAATTCTTGGACCGTTTGGTGGAAGACATCGACG<br>GGTTCCTCATCCTCGATCTCCACAATATTTACTGCCAGAG<br>CTGCAACTTCAAGGTGGACATGATGGAGATCATCAATTTA<br>TATCCGCTCGAGAAGGTCAACGAGATCCACCTCTCCGGC<br>GGCTCTTGGCAAGAGTCGGCATACGGCAAGAAACCTGTG<br>CGCCGTGATACGCATGACGACCGTATCCCTGAAGAGATT<br>CTGAATTTATTACCCGAGGTTTTAATTCAGTGCCATCCTGA<br>GTACGTAATTATTGAACGCTTAGGCCACACACTTAATACA<br>GAGGTGGCCAAGCAAATTTTCTTCGATGACTTTAACCGCG | Yes |

| | | | | | |
|---|---|---|---|---|---|
| | | | VGEEKIWNRQNTVHSKPVEDLLLYAEQTELTRLLFDGNTTESIKNRDFHYFKPESWDEEMITTAQQIIKKWNPY | TAAAGAAAATCTTGGAGTTATCAGATTACCCAGTCGGAGAAGAGAAAATTTGGAATCGCCAGAATACCGTCCACTCGAAGCCTGTTGAAGATCTCCTCTTGTATGCTGAGCAAACTGAATTAACCCGTTTACTTTTCGATGGTAATACGACGGAATCTATTAAGAATCGCGATTTTCATTATTTCAAGCCTGAATCCTGGGACGAGGAGATGATCACAACTGCGCAACAGATCATTAAGAAGTGGAACCCGTACTAA | |
| ChsA | WP_072362158.1 | Chitinophaga sancti strain DSM 784 | M**HHHHHH**KLSKSLLSAIMIGIAVQTTVVSCSKDEQVKPKKADQANKQSESKPVDNPDSCPACGMG | ATGCATCATCATCATCATCATAAACTGAGCAAAAGCCTGCTGAGCGCGATTATGATTGGTATTGCGGTTCAGACCACCGTGGTGAGCTGTAGCAAAGATGAACAGGTGAAACCGAAAAAAGCGGATCAGGCGAACAAACAGAGCGAAAGCAAACCGGTGGATAATCCGGATAGCTGCCCGGCGTGCGGCATGGGCTAA | Yes |
| ChsH | WP_072362160.1 | Chitinophaga sancti strain DSM 784 | MPKILSAVACNLDANILAACLPLFEESRVEAIEWSFDALYKVKEVPGWFRELLSAFSDENRLIGHGVFFSLFSGKWLPEQEAWLSHLQQTATEFSFDHITEHFGFMTGEDFHHGAPLNIPYSSSTLNIGKDRLKRIYHACGRPVGLENLAFSYSLDEVKRHGTFLEQLLEPVNGFIILDLHNLFCQLKNFDLDFNTLISLYPLDKVREIHISGGSWDDSAAAPDRSIRRDTHDESVPPEVFQLLEMTIPQCPNLKYVVLEQLGNGLQSTASKQGFYNDFLQMQEIAQQHNYKEGNPFLPLESLPIITGAAIEDLRLYQQQLELSAILENAGTYAEAMQQLQQSSLAHSDWKIEEWEPYMIETAVKIARKWKK | ATGCCGAAAATTCTGAGCGCGGTGGCCTGCAATCTGGATGCAAACATTCTGGCGGCGTGCCTGCCGCTGTTTGAAGAAAGCCGCGTTGAAGCCATTGAATGGAGCTTTGATGCCCTGTATAAAGTGAAAGAAGTGCCGGGCTGGTTTCGCGAACTGCTGAGCGCCTTTAGCGATGAAAACCGCCTGATTGGCCATGGCGTGTTTTTTTCACTGTTTTCGGGCAAATGGCTGCCGGAACAGGAAGCGTGGCTGAGCCATCTGCAGCAGACCGCGACCGAATTTAGCTTCGATCACATCACCGAACATTTCGGCTTCATGACCGGCGAAGATTTCCACCATGGCGCCCCGCTGAATATCCCGTACAGCAGCAGCACCCTGAACATTGGCAAAGATCGTCTGAAACGCATTTATCATGCGTGCGGTCGCCCGGTTGGCCTGGAAAATCTGGCCTTCAGCTATAGCCTGGATGAAGTGAAACGCCATGGCACCTTTCTGGAACAGCTGCTGGAACCGGTGAACGGCTTCATTATTCTGGATCTGCATAACCTGTTTTGTCAGCTGAAAAATTTTGATCTGGATTTTAATACCCTGATTTCACTGTATCCGCTGGATAAAGTGCGTGAAATTCATATTAGCGGCGGTAGCTGGGATGATAGCGCGGCGGCGCCGGATCGTAGCATTCGCCGCGATACCCACGATGAAAGCGTTCCGCCGGAGGTTTTTCAGCTGCTGGAAATGACCATTCCACAGTGCCCGAACCTGAAATATGTGGTGCTGGAACAGCTGGGCAACGGCCTGCAGTCTACCGCGTCCAAACAGGGCTTCTATAACGATTTTCTGCAGATGCAGGAAATTGCCCAGCAGCACAATTATAAAGAAGGCAACCCGTTCCTGCCGCTGGAAAGCCTGCCGATTATTACCGGCGCGGCGATTGAAGATCTGCGTCTGTATCAGCAGCAGCTGGAGCTGTCAGCGATTCTGGAAAATGCGGGTACCTATGCGGAAGCCATGCAGCAGCTGCAGCAGTCGAGCCTGGCCCATAGCGATTGGAAAATTGAAGAATGGGAACCGTACATGATTGAAACCGCGGTGAAAATTGCGCGCAAATGGAAAAAATAA | Yes |
| HymA | WP_170934670.1 | Hymenobacter gelipurpurascens strain DSM 11116 | M**HHHHHH**KLSQAILGAVLVGLTAQTTGCIKKSDPTPKEEQGKSGKKSPEIPYNCPGCGLG | ATGCATCATCATCATCATCATAAGCTCAGCCAAGCCATATTAGGCGCAGTATTAGTGGGTCTGACAGCTCAAACAACCGGTTGTATAAAGAAGTCTGATCCTACTCCAAAGGAAGAACAAGGAAAGAGCGGTAAGAAGAGCCCGGAGATTCCGTACAATTGTCCAGGATGCGGACTGGGGTAA | Yes |
| HymH | WP_088841642.1 | Hymenobacter gelipurpurascens strain DSM 11116 | MRTVAPPDNANERAIYSSIACNLDANILTAAFPLLEQGKVEALEWSFDALFWAEQVPDWFTELLQAYSAQHRLVGHGVYFSLLSGRWTAEQQQWLQHLKELTRRFSFDHVTEHFGFFTGQNFHAGAPLPIPYGSNTLRIGQDRLRRMQEACGCPVGLENLAFAYSLEEVKRHGEFLEQLVEPVNGFLILDLHNVFCQLHNFSVPYEELMALYPLDRVREIHISGGSWEASGQVPGLQIRRDTHDEAVPEEVFQLLYNTMPRCPNLKYVVLEQLGTGLQTEDSRAHFQSDFHRMQALVNQHRSTTSIRAVQPFLPIHSAVAGPVAE | ATGAGAACTGTGGCTCCACCGGATAACGCTAACGAACGTGCGATTTATTCATCAATCGCGTGTAATCTTGATGCGAATATCCTCACCGCTGCGTTCCCTCTGCTTGAACAAGGCAAAGTTGAAGCGTTAGAGTGGTCATTCGACGCACTTTTCTGGGCAGAACAGGTTCCAGACTGGTTCACAGAGCTCTTACAAGCATATTCGGCCCAGCACCGCTTAGTAGGTCACGGTGTTTATTTTAGCTTATTAAGCGGCAGATGGACCGCGGAACAACAACAATGGCTCCAGCATCTTAAAGAATTGACACGACGTTTCAGTTTTGATCATGTAACCGAACATTTCGGTTTCTTCACAGGGCAAAACTTCCACGCAGGTGCCCCATTGCCGATTCCCTACGGATCTAATACGCTCCGTATCGGCCAGGATCGATTACGACGCATGCAAGAAGCATGTGGTTGCCCAGTCGGGCTGGAGAATTTAGCATTCGCATACAGCCTGGAAGAGGTTAAGCGCCATGGTGAATTCTTGGAGCAATTGGTTGAACCTGTCAATGGGTTCTTGATCTTGGATTTACATAATGTCTTTTGTCAACTGCATAATTTTAGCGTACCTTATGAGGAATTGATGGCTTTGTACCCATTAGACCGTGTCCGTGAGATACACATCAGCGGCGGATCATGGGAAGCTAGTGGCCAAGTTCCCGGGCTGCAAATACGACGTGACACGCACGACGAAGCAGTTCCTGAGGAAGTTTTCCAACTTTTGTATAATACGATGCCTCGTTGCCCGAATCTTAAGTACGTCGTATTAGAACAACTCGGCACCGGTCTGCAAACAGAAGATTCAAGAGCGCATTTTCAATCAGATTTCCACCGAATGCAAGCTTTAGTAAATCAGCATCGTAGTACAACATCCATACGAGCGGTGCAACCCTTCTTGCCGATTCACAGCGCCGTAGCT | Yes |

| | | | | | |
|---|---|---|---|---|---|
| | | | DAQLHEQQQQLSH<br>ILETAPSYEDAQRQ<br>LQASALAYTAWKV<br>EEWEPHMLETAISI<br>AQKWKPQPR | GGGCCTGTGGCAGAGGACGCACAACTTCATGAACAACAA<br>CAACAACTGTCTCATATTCTTGAGACCGCACCTTCTTATGA<br>GGATGCACAACGCCAACTGCAAGCCTCGGCTCTCGCGTA<br>TACAGCCTGGAAGGTAGAAGAATGGGAGCCGCACATGCT<br>TGAGACTGCTATTTCCATCGCACAAAAGTGGAAACCACAA<br>CCACGTTGA | |
| ArlA | WP_25744<br>8965.1 | Archangium<br>lipolyticum strain<br>CY-1 | M**GSSHHHHHH**NQ<br>NLREKATQLLDVAS<br>NTQGMKSSSPLPC<br>PACGMMMALPESS<br>PQLVTPATR | ATGGGCAGCAGCCATCACCATCATCACCACAACCAGAATC<br>TCCGTGAGAAGGCCACCCAGCTCCTCGATGTGGCGAGCA<br>ACACCCAGGGAATGAAGTCGAGCAGTCCTCTTCCCTGTC<br>CGGCCTGCGGCATGATGATGGCGCTGCCCGAGTCCAGCC<br>CCCAACTCGTGACCCCGGCCACCCGCTGA | No |
| ArlH | WP_25744<br>8967.1 | Archangium<br>lipolyticum strain<br>CY-1 | MPPRVGLNLLTDD<br>AFREAVRPLFAEGL<br>VAALEWDIDDSWG<br>FSTRALPGWTERM<br>LDLYAEEGALYGH<br>GVWLSVLTAAWQP<br>RQEAWIERLAQEC<br>RRRRYRHVSEHFG<br>FTAAGPFTRSTMLP<br>LPYCTAAVDIGRDR<br>LERLRAATGGPVGL<br>EVLANTLAPVDALH<br>QGPFLDAVLTPTEG<br>FLVLDVHNVWTQA<br>VNTGLPPELLLETY<br>PLECVREIHLSGGS<br>WGRALGAGDSRPV<br>RLDSHDGPLLSPVL<br>SLLRRALALCPHCE<br>VVIVERRSETLESE<br>EVRAGWRAQYRAV<br>VQLVAEASATPVSP<br>VRVSASAPPPELLD<br>TEELARYQRELVTA<br>LVEESESRVILQRL<br>RQGIAGARLAPYLE<br>TFDPRMVELLAILA<br>SRWSAFDPARATP | ATGCCGCCTCGCGTGGGTCTGAATCTTCTGACCGACGAC<br>GCCTTCCGCGAGGCGGTACGGCCGCTCTTCGCCGAGGG<br>GCTGGTCGCCGCGCTCGAGTGGGACATCGACGATTCCTG<br>GGGATTCTCCACCCGTGCACTCCCCGGCTGGACCGAGCG<br>TATGCTCGACCTCTATGCGGAGGAGGGAGCCTTGTACGG<br>CCACGGCGTGTGGCTGTCGGTGCTGACCGCCGCCTGGC<br>AGCCTCGGCAGGAGGCGTGGATCGAGCGTCTCGCCCAG<br>GAGTGCCGCCGCCGCCGCTACCGCCACGTCTCCGAGCA<br>CTTCGGCTTCACCGCCGCGGGGCCGTTCACCCGCAGCAC<br>CATGCTGCCCCTGCCGTATTGCACCGCCGCCGTGGACAT<br>CGGCCGCGACCGGCTCGAACGGCTGCGCGCCGCCACCG<br>GCGGCCCCGTCGGCCTCGAGGTGCTGGCCAACACCCTC<br>GCCCCCGTCGATGCGCTCCACCAAGGTCCGTTCCTCGAT<br>GCCGTGCTGACCCCCACCGAGGGCTTCCTGGTGCTCGAC<br>GTCCACAACGTGTGGACCCAGGCGGTCAACACCGGCCTG<br>CCTCCAGAGCTCCTGCTCGAGACCTATCCGCTGGAGTGC<br>GTCCGGGAAATCCACCTCTCGGGTGGGAGCTGGGGCCG<br>CGCGCTCGGCGCCGGGGACTCCCGTCCCGTCCGGCTCG<br>ACAGTCACGACGGACCCCTCCTGTCACCCGTGCTGTCCC<br>TGCTCCGCCGGGCGCTCGCCCTCTGCCCTCATTGCGAGG<br>TCGTCATCGTCGAGCGGCGAAGCGAGACCCTCGAGTCAG<br>AGGAAGTGCGCGCCGGGTGGCGCGCCCAGTACCGCGCC<br>GTGGTCCAGCTGGTGGCCGAGGCCAGCGCAACCCCCGT<br>CAGCCCGGTCCGTGTCTCCGCCTCCGCGCCGCCCCCCG<br>AGCTGCTCGACACGGAGGAGCTCGCCCGGTACCAGCGC<br>GAGCTGGTCACGGCCCTCGTCGAGGAGTCCGAGAGCCG<br>GGTTATCCTGCAGCGTCTGCGCCAGGGTATCGCCGGCGC<br>ACGGCTCGCGCCCTACCTCGAGACCTTCGATCCGCGCAT<br>GGTCGAGCTGCTGGCCATCCTCGCGTCACGCTGGTCCGC<br>CTTCGACCCCGCTCGAGCCACTCCCTGA | No |
| MelA1 | MCC6978<br>278.1 | Candidatus<br>Melainabacteria<br>bacterium isolate<br>SJ512 | M**GSSHHHHHHSQ**<br>**M**KNKSAKVLTSLFA<br>LGLTTGAVPARADK<br>AAVETAEASVKMFE<br>QRSGTNNSSYTSLL<br>MYLAGVYLANDRE<br>KDAEIIYNKAIFNLK<br>SKPDRKAEVPALML<br>NWAMLLASNKGAS<br>KEKAEKALSDGLKL<br>ANDLPAASKERINY<br>LIGTINFYNVIGKPA<br>EKQARIKAADEHLA<br>TLEKNEKLNNEEIT<br>NVAANLVKLAEIQT<br>FPMPVMRLRYQPP<br>VFQVVPDNSPDKP<br>NTVRAKEFKSAEAF<br>QLRAIKQYDRLPET<br>VPWRIEAHRKLILW<br>YRSLGQTKQEEFQI<br>QQLGKIMHTTDRD<br>KLFPQPAPCPACG<br>MG | ATGGGCAGCAGCCATCACCATCATCACCACAGCCAGATG<br>AAAAACAAATCAGCCAAAGTTCTAACTTCGCTATTCGCGCT<br>CGGTCTGACCACTGGCGCCGTGCCCGCGCGCGCAGATA<br>AAGCAGCGGTGGAAACAGCCGAAGCAAGCGTAAAAATGT<br>TCGAGCAACGCTCAGGCACGAACAATTCGTCTTATACCAG<br>CCTACTGATGTACCTGGCTGGGGTATATCTGGCGAACGAC<br>AGAGAGAAAGACGCTGAGATCATATACAACAAAGCAATCT<br>TCAATCTCAAATCGAAACCGGATAGGAAAGCTGAAGTGCC<br>CGCCTTGATGCTGAACTGGGCAATGCTGCTGGCATCGAA<br>CAAAGGTGCATCTAAAGAAAAAGCGGAAAAAGCATTGTCG<br>GATGGACTGAAACTCGCAAACGACTTGCCGGCCGCCTCC<br>AAAGAGCGGATCAATTATCTGATAGGAACAATCAATTTCTA<br>CAACGTAATTGGCAAACCTGCCGAGAAGCAAGCCCGCAT<br>CAAAGCAGCCGATGAGCATCTTGCCACGCTGGAGAAAAA<br>CGAAAAACTAAACAATGAAGAAATCACCAATGTCGCAGCG<br>AATCTAGTGAAACTGGCAGAGATTCAAACGTTTCCCATGC<br>CGGTCATGCGTTTGCGCTACCAGCCTCCGGTGTTTCAAGT<br>AGTGCCGGACAACTCGCCGGACAAACCGAACACAGTGCG<br>AGCAAAGGAGTTCAAGTCGGCGGAAGCTTTCCAATTGCG<br>CGCGATAAAACAATACGATCGTTTGCCGGAAACGGTGCC<br>CTGGCGCATCGAAGCGCACAGAAAGTTGATTCTCTGGTAT<br>CGCAGCCTGGGTCAGACGAAACAGGAAGAGTTTCAAATT<br>CAGCAGTTAGGCAAAATCATGCACACTACAGATCGCGACA<br>AACTGTTTCCCCAGCCGGCACCCTGCCCGGCTTGCGGCA<br>TGGGTTAG | No |

| | | | | | |
|---|---|---|---|---|---|
| MelH | MCC6978279.1 | Candidatus Melainabacteria bacterium isolate SJ512 | MQKSKPKLGLSLMPTEDFQMATEQLFEEERVESLEWSFDFTWNGVVVDPWALELIDKYSEANALTGHGVNLSPLSARFSKRQEEWLARAREEFKTRKYVHASEHFGFSEAGPIKHGAPLSVPMDAASLKVGKEMLKRYADATGCPVGLENLAFAFSINDVKKQGHFIDELISSVNGFLLLDLHNIFCQIANFGISELELLNSYPLDKVREIHISGGSWSPSISGKRAAVRRDTHDDGVPQEVFNLAALALKLCPNIEFVIFERLGNTMFSLEAQSEFRDDFESMEQILEFCYA | ATGCCCACCGAGGACTTCCAAATGGCAACGGAACAGTTATTCGAGGAAGAGCGCGTGGAGAGCTTGGAGTGGTCGTTCGACTTTACGTGGAATGGAGTCGTTGTGGACCCCTGGGCATTAGAGCTGATCGATAAATACAGCGAGGCTAACGCTCTTACGGGACACGGTGTTAATCTGTCCCCGCTGTCTGCCCGTTTCTCTAAGAGACAAGAGGAGTGGCTCGCTCGTGCTCGCGAGGAATTCAAGACCCGCAAATACGTGCACGCTTCTGAACACTTTGGGTTCTCCGAGGCTGGTCCTATCAAGCACGGAGCTCCCCTGTCTGTTCCGATGGACGCGGCCAGCCTTAAGGTTGGTAAGGAAATGTTAAAGAGATATGCCGACGCTACCGGGTGTCCCGTTGGACTTGAAAATCTGGCGTTCGCTTTCTCTATAAACGACGTTAAGAAACAAGGTCACTTCATTGATGAGCTGATCAGTTCTGTAAACGGATTCCTGTTATTGGACTTACATAACATTTTCTGCCAAATTGCCAATTTCGGAATAAGTGAGCTGGAGCTTCTTAATAGCTATCCGTTAGATAAGGTCCGGGAGATACATATATCTGGTGGTTCATGGTCACCCAGTATTTCTGGTAAACGTGCGGCCGTGAGAAGAGATACCCACGATGACGGGGTGCCTCAGGAAGTGTTTAACCTGGCAGCGTTGGCGTTGAAACTGTGTCCGAACATCGAGTTTGTGATATTCGAGCGACTTGGAAACACTATGTTTTCTTTAGAGGCACAGTCCGAATTCCGGGATGACTTTGAATCAATGGAACAAATTCTTGAATTCTGTTACGCGTAA | Yes |
| $MelH_c$ | MCC6978280.1 | Candidatus Melainabacteria bacterium isolate SJ512 | MPDTLKLQEFQDALNELLASGKSHAEIMEELNSNPQFESYREYISEFDPDMVEVARELMGKWAQRVGE | ATGCCCGACACTCTGAAGCTCCAGGAATTTCAGGATGCATTGAACGAACTCCTCGCTTCGGGTAAGAGCCACGCGGAGATTATGGAAGAGTTAAATAGTAACCCGCAGTTCGAGTCATACCGGGAGTATATATCGGAGTTTGACCCCGACATGGTGGAAGTTGCCCGAGAACTCATGGGTAAATGGGCCCAACGGGTAGGTGAATGA | Yes |
| NedA | WP_085365104.1 | Neisseria dentiae strain CCUG 53898 310003 | MKTTSTLMALAGALVLAGCQVEPAQSSSPEVSVQQKAGEGKCGAASQAGKAAEGKCGEGKCGSK**HHHHHH** | ATGAAAACCACGTCGACCCTGATGGCACTGGCAGGGGCGCTGGTGCTGGCGGGCTGCCAAGTCGAACCAGCGCAGTCGTCCAGTCCTGAGGTTAGCGTGCAGCAGAAAGCAGGCGAAGGCAAATGCGGCGCAGCCTCGCAGGCGGGTAAGGCGGCGGAGGGAAAATGTGGTGAAGGTAAATGCGGCTCGAAGCATCACCATCATCACCACTGA | Yes |
| NedB | WP_085365102.1 | Neisseria dentiae strain CCUG 53898 310003 | MNTLHGAGLGYKRSMAADFLQLDRNNSPIRFIEIAPENWLRMGGAARKQFDEVAERFPVACHGLSLSLGGQDPLQLDFLKQIKAFLRQYRIGFFSEHLSYCSHHGHIYDLLPLPFTEESVRHTAARIRAVQDILEMRIAVENTSYYAHNPIAEMDEAEFLNAVVREADCDIHLDINNIYVNAVNHGIVAPRDYIDRTDLARVSYMHMAGHDEKTENLLIDTHGQPVCDDVWDLFAYACRRLPHSVPTLLERDSNFPPFAELEAEVARIAAIQQQAEKERHAAA | ATGAACACTCTGCAcGGCGCTGGGCTGGGTTATAAACGTTCTATGGCAGCAGATTTCCTGCAGCTGGATCGCAACAATTCTCCGATTCGCTTTATCGAGATTGCCCCGGAAAATTGGTTGCGTATGGGCGGCGCGGCCCGCAAGCAATTCGACGAAGTGGCCGAGCGTTTTCCGGTGGCTTGCCAcGGTCTGAGCCTGTCCCTGGGAGGCCAGGACCCGTTACAACTGGATTTCCTGAAACAGATCAAAGCGTTTTTGCGCCAGTACCGtATCGGTTTCTTTTCAGAACATCTGTCCTACTGTTCCCACCAcGGACACATCTACGATCTGCTGCCGCTTCCGTTCACGGAAGAGTCGGTACGTCACACAGCTGCCCGCATTCGCGCGGTGCAGGACATTTTGGAGATGCGTATCGCCGTGGAAAATACATCTTACTATGCACACAATCCCATCGCGGAGATGGATGAAGCGGAGTTCCTGAACGCCGTGGTGCGTGAGGCGGATTGCGACATCCACTTAGATATTAATAATATTTACGTGAATGCCGTCAACCACGGCATCGTGGCCCCACGCGACTACATTGACCGCACGGATCTGGCCCGTGTGAGTTATATGCACATGGCAGGCCATGACGAGAAGACTGAGAATTTGTTAATCGACACCCACGGCCAACCGGTTTGTGATGACGTGTGGGACTTATTTGCCTACGCGTGTCGCCGCCTGCCTCACTCTGTGCCAACCTTACTGGAACGCGATAGCAATTTTCCGCCTTTTGCCGAATTAGAAGCCGAGGTAGCTCGCATCGCCGCTATCCAACAGCAGGCCGAGAAAGAACGTCATGCGGCGGCGTAA | Yes |
| NedC | WP_085365101.1 | Neisseria dentiae strain CCUG 53898 310003 | MQPHNPSPSARAQAELADHVRNPALPAPAGIAPERLAVYTRLVRNNLKSFLDLCFSDSSLMLDPAQWQGWQNRFLIEARPESPFFNDIPAQFLAYLNRLPEHDRPSENILAMMDFETALLHAETARQPDSDGRWSEHSVLSWAPAARLQQYPCDFVSSGLAQINAGACHVLSWRNRRNEVYYRIVEDTDL | ATGCAACCGCATAACCCAAGTCCGTCGGCGCGTGCGCAAGCCGAGCTGGCAGATCATGTTCGCAACCCTGCACTGCCCGCCCCGGCGGGGATTGCACCGGAACGCCTGGCGGTATATACCCGCCTGGTGCGTAACAATCTTAAATCGTTCCTGGATCTGTGCTTCTCTGATTCCAGCCTGATGCTGGATCCCGCGCAATGGCAGGGCTGGCAGAATCGCTTTCTGATCGAGGCACGCCCCGAAAGCCCGTTTTTTAATGATATTCCGGCGCAGTTTCTTGCcTATTTAAACCGTTTGCCTGAACATGATCGTCCGTCAGAGAACATTCTTGCGATGATGGATTTCGAAACCGCTCTGTTGCATGCGGAGACGGCGCGCCAGCCTGATTCTGATGGCCGTTGGAGCGAGCATAGCGTCCTGAGCTGGGCCCCGGCCGCACGTCTGCAGCAATATCCGTGTGATTTCGTGAGCTCCGGCCTGGCACAGATCAAcGCGGGCGCGTGTCACGTTCTGTCCTGGCGTAACCGTCGCAACGAAGTCTATTATCGTATC | Yes |

| | | | | |
|---|---|---|---|---|
| | | | FLLQHFQSQNDTFA RLLESLQSLLPGQD IEGRLKTAVGGWV EAGVLLTAE | GTGGAAGATACCGATCTGTTTCTGCTGCAGCAcTTTCAGT CACAGAACGACACTTTTGCGCGCCTGCTGGAAAGCCTGC AGTCCTTATTACCGGGTCAAGATATTGAAGGTCGTCTTAA AACGGCCGTCGGTGGGTGGGTTGAAGCCGGCGTGCTGC TGACCGCGGAGTAA |

### Table S10. Primers and gblocks used in this study.

| Primer/gblock name | Sequence (5'-3') | Purpose |
|---|---|---|
| pACYC:DybA insert F | ATCACCACAAAATCTCAAAACCAGTACTTCAGGCC | For cloning DybA into pACYCDuet |
| pACYC:DybA insert R | GCAAGCTTTCAACCCATTCCGCAGGCC | For cloning DybA into pACYCDuet |
| pACYC:DybA plasmid F | AATGGGTTGAAAGCTTGCGGCCGCATAATG | For cloning DybA into pACYCDuet |
| pACYC:DybA plasmid R | GTTTTGAGATTTTGTGGTGATGATGGTGATGGCTGC | For cloning DybA into pACYCDuet |
| pRSF:DybAH insert F | ATCACCACAAAATCTCAAAACCAGTACTTCAGGCC | For cloning DybA and DybH into pRSFDuet |
| pRSF:DybAH insert R | AGCCTAGGTTAATTATCAAAATCCTGCCTTCCATTTCTGCG | For cloning DybA and DybH into pRSFDuet |
| pRSF:DybAH plasmid F | GATTTTGATAATTAACCTAGGCTGCTGCCACCGC | For cloning DybA and DybH into pRSFDuet |
| pRSF:DybAH plasmid R | GTTTTGAGATTTTGTGGTGATGATGGTGATGGCTGC | For cloning DybA and DybH into pRSFDuet |
| pACYC:ChsA/HymA plasmid F | AACAGAAAGTAATCGTATTGTACACG | For cloning precursor peptides into pACYCDuet |
| pACYC:ChsA/HymA plasmid R | ATTTCCTAATGCAGGAGTCGCATAAG | For cloning precursor peptides into pACYCDuet |
| pACYC:MelA1 plasmid F | AGCTTGCGGCCGCATAATGCTTA | For cloning precursor protein into pACYCDuet |
| pACYC:MelA1 plasmid R | CTGCTGCCCATGGTATATCTCCTTATTAAAG | For cloning precursor protein into pACYCDuet |
| pRSF:ArlAH plasmid F | CAATAACTAGCATAACCCCTTGGGG | For cloning in ArlA and ArlH gblock into pRSFDuet |
| pRSF:ArlAH plasmid R | TATCTCCTTATTAAAGTTAAACAAAATTATTTCTACAGG | For cloning in ArlA and ArlH gblock into pRSFDuet |
| pCDF:DybH insert F | TTTAATAAGGAGATATACATGTCGCAAATATTCTCCACAGTCG | For cloning modifying enzyme into pCDFDuet |
| pCDF:DybH insert R | CGCAAGCTTTCAAAATCCTGCCTTCCATTTCTGC | For cloning modifying enzyme into pCDFDuet |
| pCDF:DybH plasmid F | CAGGATTTTGAAAGCTTGCGGCCGCATAATGC | For cloning modifying enzyme into pCDFDuet |
| pCDF:DybH plasmid R | GCGACATGTATATCTCCTTATTAAAGTTAAACAAAATTATTTCT | For cloning modifying enzyme into pCDFDuet |
| pCDF:ChsH/HymH plasmid F | GCATAATGCTTAAGTCGAACAGAAAGTA | For cloning modifying enzyme into pCDFDuet |
| pCDF:ChsH plasmid R | CGGCATGTATATCTCCTTATTAAAGTTAAACAAAATTATTTC | For cloning modifying enzyme into pCDFDuet |
| pCDF:HymH plasmid R | GTTCTCATGTATATCTCCTTATTAAAGTTAAACAAAATTATTTC | For cloning modifying enzyme into pCDFDuet |
| pCDF:MelH+Hc plasmid F | AATGACTAGGCTGCTGCCACCG | For cloning modifying enzyme into pCDFDuet |
| pCDF:MelH+Hc plasmid R | GGGCATGTATATCTCCTTATTAAAGTTAAACAAAATTATTTC | For cloning modifying enzyme into pCDFDuet |
| pCDF:MelH+Hc N-term fix F | AGAAGTCGAAACCCAAACTAGGACTTTCTCTCATGCCCACCG AGGAC | For cloning in missed N-terminal residues |
| pCDF:MelH+Hc N-term fix R | AGTTTGGGTTTCGACTTCTGCATGGTATATCTCCTTATTAAAG TTAAACAAAATT | For cloning in missed N-terminal residues |
| pCDF:MelH plasmid F | ACATATGCCCGAATGACTAGGCTGCTG | For deleting MelHc to only express MelH |
| pCDF:MelH plasmid R | CTAGTCATTCGGGCATATGTATATCTCCTTCTTATAC | For deleting MelHc to only express MelH |
| pCDF:His-MelH+Hc F | GGCAGCCATCATCATCATCATCACTCTCAGAAGTCGAAACCC AAACTAG | For adding His-tag to MelH |

| | | |
|---|---|---|
| pCDF:His-MelH+Hc R | ATGATGATGATGGCTGCCCATGGTATATCTCCTTATTAAAGTT<br>AAACAAAATTA | For adding His-tag to MelH |
| pRSF:ArlA F | TACATATGCCGACTCCCTGATAATTTACCTAGGC | For removing ArlH from plasmid |
| pRSF:ArlA R | AGGGAGTCGGCATATGTATATCTCCTTCTTATAC | For removing ArlH from plasmid |
| pRSF:DybAH no His F | ATAAGGAGATATACCATGAAAATCTCAAAACCAGTACTTCAG | To remove N-terminal His tag from DybA |
| pRSF:DybAH no His R | GAGATTTTCATGGTATATCTCCTTATTAAAGTTAAACAAAATTA<br>TTTCTAC | To remove N-terminal His tag from DybA |
| pRSF:DybAH His embed F | ATCACCATCATCACCACGATATTAAACCTGAAAAGGATAAAAA<br>AGCGAAGACGC | Inserting His tag to position 27 (following signal peptide) |
| pRSF:DybAH His embed R | GTGGTGATGATGGTGATGCATGTCGGCGCAAGCGGTAATT | Inserting His tag to position 27 (following signal peptide) |
| pRSF:NedA C-His plasmid F | AACAGAAAGTAATCGTATTGTACACG | For cloning in NedA gblock into pRSFDuet |
| pRSF:NedA C-His plasmid R | ATTTCCTAATGCAGGAGTCGCATAAG | For cloning in NedA gblock into pRSFDuet |
| pCDF:NedBC plasmid F | CAATAACTAGCATAACCCCTTGGGG | For cloning NedBC gblock into pCDFDuet |
| pCDF:NedBC plasmid R | TCCTAATGCAGGAGTCGCATAAGGGAG | For cloning NedBC gblock into pCDFDuet |
| ChsA gblock (codon optimized) | TGCGACTCCTGCATTAGGAAATTAATACGACTCACTATAGGG<br>GAATTGTGAGCGGATAACAATTCCCCTGTAGAAATAATTTTGT<br>TTAACTTTAATAAGGAGATATACCATGCATCATCATCATCATCA<br>TAAACTGAGCAAAAGCCTGCTGAGCGCGATTATGATTGGTATT<br>GCGGTTCAGACCACCGTGGTGAGCTGTAGCAAAGATGAACA<br>GGTGAAACCGAAAAAAGCGGATCAGGCGAACAAACAGAGCG<br>AAAGCAAACCGGTGGATAATCCGGATAGCTGCCCGGCGTGC<br>GGCATGGGCTAAAAGCTTGCGGCCGCATAATGCTTAAGTCGA<br>ACAGAAAGTAATCGTATTGTACACG | Precursor peptide |
| ChsH gblock (codon optimized) | TTGTTTAACTTTAATAAGGAGATATACATGCCGAAAATTCTGA<br>GCGCGGTGGCCTGCAATCTGGATGCAAACATTCTGGCGGCG<br>TGCCTGCCGCTGTTTGAAGAAAGCCGCGTTGAAGCCATTGAA<br>TGGAGCTTTGATGCCCTGTATAAAGTGAAAGAAGTGCCGGGC<br>TGGTTTCGCGAACTGCTGAGCGCCTTTAGCGATGAAAACCGC<br>CTGATTGGCCATGGCGTGTTTTTTTCACTGTTTTCGGGCAAAT<br>GGCTGCCGGAACAGGAAGCGTGGCTGAGCCATCTGCAGCAG<br>ACCGCGACCGAATTTAGCTTCGATCACATCACCGAACATTTC<br>GGCTTCATGACCGGCGAAGATTTCCACCATGGCGCCCCGCT<br>GAATATCCCGTACAGCAGCAGCACCCTGAACATTGGCAAAGA<br>TCGTCTGAAACGCATTTATCATGCGTGCGGTCGCCCGGTTGG<br>CCTGGAAAATCTGGCCTTCAGCTATAGCCTGGATGAAGTGAA<br>ACGCCATGGCACCTTTCTGGAACAGCTGCTGGAACCGGTGAA<br>CGGCTTCATTATTCTGGATCTGCATAACCTGTTTTGTCAGCTG<br>AAAAATTTTGATCTGGATTTTAATACCCTGATTTCACTGTATCC<br>GCTGGATAAAGTGCGTGAAATTCATATTAGCGGCGGTAGCTG<br>GGATGATAGCGCGGCGGCGCCGGATCGTAGCATTCGCCGCG<br>ATACCCACGATGAAAGCGTTCCGCCGGAGGTTTTTCAGCTGC<br>TGGAAATGACCATTCCACAGTGCCCGAACCTGAAATATGTGG<br>TGCTGGAACAGCTGGGCAACGGCCTGCAGTCTACCGCGTCC<br>AAACAGGGCTTCTATAACGATTTTCTGCAGATGCAGGAAATTG<br>CCCAGCAGCACAATTATAAAGAAGGCAACCCGTTCCTGCCGC<br>TGGAAAGCCTGCCGATTATTACCGGCGCGGCGATTGAAGATC<br>TGCGTCTGTATCAGCAGCAGCTGGAGCTGTCAGCGATTCTGG<br>AAAATGCGGGTACCTATGCGGAAGCCATGCAGCAGCTGCAG<br>CAGTCGAGCCTGGCCCATAGCGATTGGAAAATTGAAGAATGG<br>GAACCGTACATGATTGAAACCGCGGTGAAAATTGCGCGCAAA<br>TGGAAAAAAATAAAAGCTTGCGGCCGCATAATGCTTAAGTCGA<br>ACAGAAAGTA | Modifying enzyme |
| HymA gblock (codon optimized) | TGCGACTCCTGCATTAGGAAATTAATACGACTCACTATAGGG<br>GAATTGTGAGCGGATAACAATTCCCCTGTAGAAATAATTTTGT<br>TTAACTTTAATAAGGAGATATACCATGCATCATCATCATCATCA<br>TAAGCTCAGCCAAGCCATATTAGGCGCAGTATTAGTGGGTCT<br>GACAGCTCAAACAACCGGTTGTATAAAGAAGTCTGATCCTACT<br>CCAAAGGAAGAACAAGGAAAGAGCGGTAAGAAGAGCCCGGA<br>GATTCCGTACAATTGTCCAGGATGCGGACTGGGGTAAAAGCT<br>TGCGGCCGCATAATGCTTAAGTCGAACAGAAAGTAATCGTATT<br>GTACACG | Precursor peptide |
| HymH gblock (codon optimized) | TTGTTTAACTTTAATAAGGAGATATACATGAGAACTGTGGCTC<br>CACCGGATAACGCTAACGAACGTGCGATTTATTCATCAATCG | Modifying enzyme |

| | | |
|---|---|---|
| | CGTGTAATCTTGATGCGAATATCCTCACCGCTGCGTTCCCTCT<br>GCTTGAACAAGGCAAAGTTGAAGCGTTAGAGTGGTCATTCGA<br>CGCACTTTTCTGGGCAGAACAGGTTCCAGACTGGTTCACAGA<br>GCTCTTACAAGCATATTCGGCCCAGCACCGCTTAGTAGGTCA<br>CGGTGTTTATTTTAGCTTATTAAGCGGCAGATGGACCGCGGA<br>ACAACAACAATGGCTCCAGCATCTTAAAGAATTGACACGACGT<br>TTCAGTTTTGATCATGTAACCGAACATTTCGGTTTCTTCACAG<br>GGCAAAACTTCCACGCAGGTGCCCCATTGCCGATTCCCTACG<br>GATCTAATACGCTCCGTATCGGCCAGGATCGATTACGACGCA<br>TGCAAGAAGCATGTGGTTGCCCAGTCGGGCTGGAGAATTTAG<br>CATTCGCATACAGCCTGGAAGAGGTTAAGCGCCATGGTGAAT<br>TCTTGGAGCAATTGGTTGAACCTGTCAATGGGTTCTTGATCTT<br>GGATTTACATAATGTCTTTTGTCAACTGCATAATTTTAGCGTAC<br>CTTATGAGGAATTGATGGCTTTGTACCCATTAGACCGTGTCCG<br>TGAGATACACATCAGCGGCGGATCATGGGAAGCTAGTGGCC<br>AAGTTCCCGGGCTGCAAATACGACGTGACACGCACGACGAA<br>GCAGTTCCTGAGGAAGTTTTCCAACTTTTGTATAATACGATGC<br>CTCGTTGCCCGAATCTTAAGTACGTCGTATTAGAACAACTCGG<br>CACCGGTCTGCAAACAGAAGATTCAAGAGCGCATTTTCAATC<br>AGATTTCCACCGAATGCAAGCTTTAGTAAATCAGCATCGTAGT<br>ACAACATCCATACGAGCGGTGCAACCCTTCTTGCCGATTCAC<br>AGCGCCGTAGCTGGGCCTGTGGCAGAGGACGCACAACTTCA<br>TGAACAACAACAACAACTGTCTCATATTCTTGAGACCGCACCT<br>TCTTATGAGGATGCACAACGCCAACTGCAAGCCTCGGCTCTC<br>GCGTATACAGCCTGGAAGGTAGAAGAATGGGAGCCGCACAT<br>GCTTGAGACTGCTATTTCCATCGCACAAAAGTGGAAACCACA<br>ACCACGTTGAAAGCTTGCGGCCGCATAATGCTTAAGTCGAAC<br>AGAAAGTA | |
| MelA1 gblock (not codon optimized) | ATAAGGAGATATACCATGGGCAGCAGCCATCACCATCATCAC<br>CACAGCCAGATGAAAAACAAATCAGCCAAAGTTCTAACTTCGC<br>TATTCGCGCTCGGTCTGACCACTGGCGCCGTGCCCGCGCGC<br>GCAGATAAAGCAGCGGTGGAAACAGCCGAAGCAAGCGTAAA<br>AATGTTCGAGCAACGCTCAGGCACGAACAATTCGTCTTATAC<br>CAGCCTACTGATGTACCTGGCTGGGGTATATCTGGCGAACGA<br>CAGAGAGAAAGACGCTGAGATCATATACAACAAAGCAATCTT<br>CAATCTCAAATCGAAACCGGATAGGAAAGCTGAAGTGCCCGC<br>CTTGATGCTGAACTGGGCAATGCTGCTGGCATCGAACAAAGG<br>TGCATCTAAAGAAAAAGCGGAAAAAGCATTGTCGGATGGACT<br>GAAACTCGCAAACGACTTGCCGGCCGCCTCCAAAGAGCGGA<br>TCAATTATCTGATAGGAACAATCAATTTCTACAACGTAATTGG<br>CAAACCTGCCGAGAAGCAAGCCCGCATCAAAGCAGCCGATG<br>AGCATCTTGCCACGCTGGAGAAAAACGAAAAACTAAACAATG<br>AAGAAATCACCAATGTCGCAGCGAATCTAGTGAAACTGGCAG<br>AGATTCAAACGTTTCCCATGCCGGTCATGCGTTTGCGCTACC<br>AGCCTCCGGTGTTTCAAGTAGTGCCGGACAACTCGCCGGACA<br>AACCGAACACAGTGCGAGCAAAGGAGTTCAAGTCGGCGGAA<br>GCTTTCCAATTGCGCGCGATAAAACAATACGATCGTTTGCCG<br>GAAACGGTGCCCTGGCGCATCGAAGCGCACAGAAAGTTGATT<br>CTCTGGTATCGCAGCCTGGGTCAGACGAAACAGGAAGAGTTT<br>CAAATTCAGCAGTTAGGCAAAATCATGCACACTACAGATCGC<br>GACAAACTGTTTCCCCAGCCGGCACCCTGCCCGGCTTGCGG<br>CATGGGTTAGAGCTTGCGGCCGCATAATGCTTA | Precursor protein |
| MelH + MelHc gblock (codon optimized) | TTGTTTAACTTTAATAAGGAGATATACATGCCCACCGAGGACT<br>TCCAAATGGCAACGGAACAGTTATTCGAGGAAGAGCGCGTGG<br>AGAGCTTGGAGTGGTCGTTCGACTTTACGTGGAATGGAGTCG<br>TTGTGGACCCCTGGGCATTAGAGCTGATCGATAAATACAGCG<br>AGGCTAACGCTCTTACGGGACACGGTGTTAATCTGTCCCCGC<br>TGTCTGCCCGTTTCTCTAAGAGACAAGAGGAGTGGCTCGCTC<br>GTGCTCGCGAGGAATTCAAGACCCGCAAATACGTGCACGCTT<br>CTGAACACTTTGGGTTCTCCGAGGCTGGTCCTATCAAGCACG<br>GAGCTCCCCTGTCTGTTCCGATGGACGCGGCCAGCCTTAAG<br>GTTGGTAAGGAAATGTTAAAGAGATATGCCGACGCTACCGGG<br>TGTCCCGTTGGACTTGAAAATCTGGCGTTCGCTTTCTCTATAA<br>ACGACGTTAAGAAACAAGGTCACTTCATTGATGAGCTGATCA<br>GTTCTGTAAACGGATTCCTGTTATTGGACTTACATAACATTTTC<br>TGCCAAATTGCCAATTTCGGAATAAGTGAGCTGGAGCTTCTTA<br>ATAGCTATCCGTTAGATAAGGTCCGGGAGATACATATATCTGG<br>TGGTTCATGGTCACCCAGTATTTCTGGTAAACGTGCGGCCGT<br>GAGAAGAGATACCCACGATGACGGGGTGCCTCAGGAAGTGT<br>TTAACCTGGCAGCGTTGGCGTTGAAACTGTGTCCGAACATCG | Modifying enzymes |

| | | |
|---|---|---|
| | AGTTTGTGATATTCGAGCGACTTGGAAACACTATGTTTTCTTT<br>AGAGGCACAGTCCGAATTCCGGGATGACTTTGAATCAATGGA<br>ACAAATTCTTGAATTCTGTTACGCGTAAAAGCTTGCGGCCGCA<br>TAATGCTTAAGTCGAACAGAAAGTAATCGTATTGTACACGGCC<br>GCATAATCGAAATTAATACGACTCACTATAGGGGAATTGTGAG<br>CGGATAACAATTCCCCATCTTAGTATATTAGTTAAGTATAAGAA<br>GGAGATATACATATGCCCGACACTCTGAAGCTCCAGGAATTT<br>CAGGATGCATTGAACGAACTCCTCGCTTCGGGTAAGAGCCAC<br>GCGGAGATTATGGAAGAGTTAAATAGTAACCCGCAGTTCGAG<br>TCATACCGGGAGTATATATCGGAGTTTGACCCCGACATGGTG<br>GAAGTTGCCCGAGAACTCATGGGTAAATGGGCCCAACGGGT<br>AGGTGAATGACTAGGCTGCTGCCACCG | |
| ArlAH gblock (not codon optimized) | AATTTTGTTTAACTTTAATAAGGAGATATACCATGGGCAGCAG<br>CCATCACCATCATCACCACAACCAGAATCTCCGTGAGAAGGC<br>CACCCAGCTCCTCGATGTGGCGAGCAACACCCAGGGAATGA<br>AGTCGAGCAGTCCTCTTCCCTGTCCGGCCTGCGGCATGATGA<br>TGGCGCTGCCCGAGTCCAGCCCCCAACTCGTGACCCCGGCC<br>ACCCGCTGAAAGCTTGCGGCCGCATAATGCTTAAGTCGAACA<br>GAAAGTAATCGTATTGTTCACGGCCGCATAATCGAAATTAATA<br>CGACTCACTATAGGGGAATTGTGAGCGGATAACAATTCCCCA<br>TCTTAGTATATTAGTTAAGTATAAGAAGGAGATATACATATGCC<br>GCCTCGCGTGGGTCTGAATCTTCTGACCGACGACGCCTTCCG<br>CGAGGCGGTACGGCCGCTCTTCGCCGAGGGGCTGGTCGCC<br>GCGCTCGAGTGGGACATCGACGATTCCTGGGGATTCTCCAC<br>CCGTGCACTCCCCGGCTGGACCGAGCGTATGCTCGACCTCT<br>ATGCGGAGGAGGGAGCCTTGTACGGCCACGGCGTGTGGCTG<br>TCGGTGCTGACCGCCGCCTGGCAGCCTCGGCAGGAGGCGT<br>GGATCGAGCGTCTCGCCCAGGAGTGCCGCCGCCGCCGCTAC<br>CGCCACGTCTCCGAGCACTTCGGCTTCACCGCCGCGGGGCC<br>GTTCACCCGCAGCACCATGCTGCCCCTGCCGTATTGCACCGC<br>CGCCGTGGACATCGGCCGCGACCGGCTCGAACGGCTGCGC<br>GCCGCCACCGGCGGCCCCGTCGGCCTCGAGGTGCTGGCCA<br>ACACCCTCGCCCCCGTCGATGCGCTCCACCAAGGTCCGTTC<br>CTCGATGCCGTGCTGACCCCCACCGAGGGCTTCCTGGTGCT<br>CGACGTCCACAACGTGTGGACCCAGGCGGTCAACACCGGCC<br>TGCCTCCAGAGCTCCTGCTCGAGACCTATCCGCTGGAGTGC<br>GTCCGGGAAATCCACCTCTCGGGTGGGAGCTGGGGCCGCGC<br>GCTCGGCGCCGGGGACTCCCGTCCCGTCCGGCTCGACAGTC<br>ACGACGGACCCCTCCTGTCACCCGTGCTGTCCCTGCTCCGC<br>CGGGCGCTCGCCCTCTGCCCTCATTGCGAGGTCGTCATCGT<br>CGAGCGGCGAAGCGAGACCCTCGAGTCAGAGGAAGTGCGC<br>GCCGGGTGGCGCGCCCAGTACCGCGCCGTGGTCCAGCTGG<br>TGGCCGAGGCCAGCGCAACCCCCGTCAGCCCGGTCCGTGTC<br>TCCGCCTCCGCGCCGCCCCCCGAGCTGCTCGACACGGAGGA<br>GCTCGCCCGGTACCAGCGCGAGCTGGTCACGGCCCTCGTCG<br>AGGAGTCCGAGAGCCGGGTTATCCTGCAGCGTCTGCGCCAG<br>GGTATCGCCGGCGCACGGCTCGCGCCCTACCTCGAGACCTT<br>CGATCCGCGCATGGTCGAGCTGCTGGCCATCCTCGCGTCAC<br>GCTGGTCCGCCTTCGACCCCGCTCGAGCCACTCCCTGATAAT<br>TTACCTAGGCTGCTGCCACCGCTGAGCAATAACTAGCATAAC<br>CCCTTGGGG | Precursor peptide and modifying enzyme |
| NedA gblock (codon optimized) | TGCGACTCCTGCATTAGGAAATTAATACGACTCACTATAGGG<br>GAATTGTGAGCGGATAACAATTCCCCTGTAGAAATAATTTTGT<br>TTAACTTTAATAAGGAGATATACCATGAAAACCACGTCGACCC<br>TGATGGCACTGGCAGGGGCGCTGGTGCTGGCGGGCTGCCAA<br>GTCGAACCAGCGCAGTCGTCCAGTCCTGAGGTTAGCGTGCA<br>GCAGAAAGCAGGCGAAGGCAAATGCGGCGCAGCCTCGCAGG<br>CGGGTAAGGCGGCGGAGGGAAAATGTGGTGAAGGTAAATGC<br>GGCTCGAAGCATCACCATCATCACCACTGAAAGCTTGCGGCC<br>GCATAATGCTTAAGTCGAACAGAAAGTAATCGTATTGTACACG | Precursor peptide |
| NedBC gblock (codon optimized) | CTCCCTTATGCGACTCCTGCATTAGGAAATTAATACGACTCAC<br>TATAGGGGAATTGTGAGCGGATAACAATTCCCCTGTAGAAATA<br>ATTTTGTTTAACTTTAATAAGGAGATATACCATGAACACTCTGC<br>ACGGCGCTGGGCTGGGTTATAAACGTTCTATGGCAGCAGATT<br>TCCTGCAGCTGGATCGCAACAATTCTCCGATTCGCTTTATCGA<br>GATTGCCCCGGAAAATTGGTTGCGTATGGGCGGCGCGGCCC<br>GCAAGCAATTCGACGAAGTGGCCGAGCGTTTTCCGGTGGCTT<br>GCCACGGTCTGAGCCTGTCCCTGGGAGGCCAGGACCCGTTA<br>CAACTGGATTTCCTGAAACAGATCAAAGCGTTTTTGCGCCAGT<br>ACCGTATCGGTTTCTTTTCAGAACATCTGTCCTACTGTTCCCA | Modifying enzymes |

| | |
|---|---|
| | CCACGGACACATCTACGATCTGCTGCCGCTTCCGTTCACGGA AGAGTCGGTACGTCACACAGCTGCCCGCATTCGCGCGGTGC AGGACATTTTGGAGATGCGTATCGCCGTGGAAAATACATCTTA CTATGCACACAATCCCATCGCGGAGATGGATGAAGCGGAGTT CCTGAACGCCGTGGTGCGTGAGGCGGATTGCGACATCCACT TAGATATTAATAATATTTACGTGAATGCCGTCAACCACGGCAT CGTGGCCCCACGCGACTACATTGACCGCACGGATCTGGCCC GTGTGAGTTATATGCACATGGCAGGCCATGACGAGAAGACTG AGAATTTGTTAATCGACACCCACGGCCAACCGGTTTGTGATG ACGTGTGGGACTTATTTGCCTACGCGTGTCGCCGCCTGCCTC ACTCTGTGCCAACCTTACTGGAACGCGATAGCAATTTTCCGC CTTTTGCCGAATTAGAAGCCGAGGTAGCTCGCATCGCCGCTA TCCAACAGCAGGCCGAGAAAGAACGTCATGCGGCGGCGTAA AAGCTTGCGGCCGCATAATGCTTAAGTCGAACAGAAAGTAAT CGTATTGTACACGGCCGCATAATCGAAATTAATACGACTCACT ATAGGGGAATTGTGAGCGGATAACAATTCCCCATCTTAGTATA TTAGTTAAGTATAAGAAGGAGATATACATATGCAACCGCATAA CCCAAGTCCGTCGGCGCGTGCGCAAGCCGAGCTGGCAGATC ATGTTCGCAACCCTGCACTGCCCGCCCCGGCGGGGATTGCA CCGGAACGCCTGGCGGTATATACCCGCCTGGTGCGTAACAAT CTTAAATCGTTCCTGGATCTGTGCTTCTCTGATTCCAGCCTGA TGCTGGATCCCGCGCAATGGCAGGGCTGGCAGAATCGCTTT CTGATCGAGGCACGCCCCGAAAGCCCGTTTTTTAATGATATT CCGGCGCAGTTTCTTGCCTATTTAAACCGTTTGCCTGAACATG ATCGTCCGTCAGAGAACATTCTTGCGATGATGGATTTCGAAAC CGCTCTGTTGCATGCGGAGACGGCGCGCCAGCCTGATTCTG ATGGCCGTTGGAGCGAGCATAGCGTCCTGAGCTGGGCCCCG GCCGCACGTCTGCAGCAATATCCGTGTGATTTCGTGAGCTCC GGCCTGGCACAGATCAACGCGGGCGCGTGTCACGTTCTGTC CTGGCGTAACCGTCGCAACGAAGTCTATTATCGTATCGTGGA AGATACCGATCTGTTTCTGCTGCAGCACTTTCAGTCACAGAAC GACACTTTTGCGCGCCTGCTGGAAAGCCTGCAGTCCTTATTA CCGGGTCAAGATATTGAAGGTCGTCTTAAAACGGCCGTCGGT GGGTGGGTTGAAGCCGGCGTGCTGCTGACCGCGGAGTAACT AGGCTGCTGCCACCGCTGAGCAATAACTAGCATAACCCCTTG GGG |
